# Analysis of 329,942 SARS-CoV-2 records retrieved from GISAID database

**DOI:** 10.1101/2021.08.04.454929

**Authors:** Maria Zelenova, Anna Ivanova, Semyon Semyonov, Yuriy Gankin

## Abstract

**Background:** The 31st of December 2019 was when the World Health Organization received a report about an outbreak of pneumonia of unknown etiology in the Chinese city of Wuhan. The outbreak was the result of the novel virus labeled as SARS-CoV-2, which spread to about 220 countries and caused approximately 3,311,780 deaths, infecting more than 159,319,384 people by May 12th, of 2021. The virus caused a worldwide pandemic leading to panic, quarantines, and lockdowns – although none of its predecessors from the coronavirus family have ever achieved such a scale. The key to understanding the global success of SARS-CoV-2 is hidden in its genome.

**Materials and Methods:** We retrieved data for 329,942 SARS-CoV-2 records uploaded to the GISAID database from the beginning of the pandemic until the 8th of January 2021. To process the data, a Python variant detection script was developed, using *pairwise2* from the BioPython library. Pandas, Matplotlib, and Seaborn, were applied to visualize the data. Genomic coordinates were obtained from the UCSC Genome Browser (https://genome.ucsc.edu/). Sequence alignments were performed for every gene separately. Genomes less than 26,000 nucleotides long were excluded from the research. Clustering was performed using HDBScan.

**Results:** Here, we addressed the genetic variability of SARS-CoV-2 using 329,942 worldwide samples. The analysis yielded 155 genome variations (SNPs and deletions) in more than 0.3% of the sequences. Nine common SNPs were present in more than 20% of the samples. Clustering results suggested that a proportion of people (2.46%) were infected with a distinct subtype of the B.1.1.7 variant. The subtype may be characterized by four to six additional mutations, with four being a more frequent option (G28881A, G28882A, and G28883С in the N gene, A23403G in S, A28095T in ORF8, G25437T in ORF3a). Two clusters were formed by mutations in the samples uploaded predominantly by Denmark and Australia, which may indicate the emergence of “Danish” and “Australian” variants. Five clusters were linked to increased/decreased age, shifted gender ratio, or both. According to a correlation coefficient matrix, 69 mutations correlate with at least one other mutation (correlation coefficient greater than 0.7). We also addressed the completeness of the GISAID database, where between 77% and 93% of the fields were either left blank or filled incorrectly. Metadata mining analysis has led to a hypothesis about gender inequality in medical care in certain countries. Finally, we found ORF6 and E as the most conserved genes (96.15% and 94.66% of the sequences totally match the reference, respectively), making them potential targets for vaccines and treatment. Our results indicate areas of the SARS-CoV-2 genome that researchers can focus on for further structural and functional analysis.

## Introduction

A virus that appeared in Wuhan in December 2019 was not initially expected to cause a worldwide crisis and a deadly pandemic. It was soon recognized as a coronavirus, a single-stranded positive-sense RNA virus belonging to a Coronaviridae family, whose members have already frightened humankind before. First discovered in the 1960s, two Coronaviridae family members (CoV-229E and CoVOC43) did not present a global threat (Qi et al., 2020, Yin C et al., 2020). However, Severe Acute Respiratory Syndrome Coronavirus (SARS-CoV, 2002/2003) and the Middle East Respiratory Syndrome Coronavirus (MERS-CoV, 2012) changed public opinion: SARS-CoV left ∼8,098 people infected and ∼774 dead; MERS-CoV caused ∼2,494 infections, leading to ∼858 deaths. The SARS-CoV-2 exceeded the predecessors, infecting more than 159,319,384 people worldwide and causing more than 3,311,780 deaths in about 220 countries and territories by May 12th, 2021 (reported by WHO). The World Health Organization declared a SARS-CoV-2 - related pandemic and public health emergency on the 30th of January 2020 (Weber et al., 2020., Walls et al., 2020). The worst outcomes of the COVID-19, the disease caused by SARS-CoV-2, are currently associated with old age (65 and older), male gender, smoking, and comorbidities such as diabetes, cardiovascular disorders, and hypertension (de Sousa et al., 2020). At present, over a year and a half later, the reasons for SARS-CoV-2 high transmissibility are still elusive. Most researchers focus on the viral genome, its evolution, and its mutations (Kaur et al., 2020). Studies of single nucleotide polymorphisms (SNPs) are especially beneficial in revealing heavily mutated genomes and understanding the viral changing pattern (Yin C et al., 2020). Since common knowledge of SARS-CoV-2 proteins’ functioning, signaling pathways, protein-protein, and protein-host cell interactions keeps rapidly accumulating due to its novelty, there is an urgent need to explore the SARS-CoV-2 changes. A relationship between SNPs and their consequences can be predicted from genotyping, protein analysis, and transmission tracking (Wang et al., 2020).

### Describing the viral sequence

SARS-CoV-2 genome was first sequenced in January 2020, a month after COVID-19 became a worldwide alert (Zhou et al., 2020., Zhu et al., 2020). The genome consists of 29903 nucleotides (GenBank accession number MN908947). Its length and overall genetic contents carry little surprise since it has long been established that coronaviruses have ones of the largest genomes amid all RNA viruses (varying from ∼26 to ∼32 kb in length) (Kaur et al., 2020). Although many mutations have currently been found in the viral genome (Wang et al., 2020), only a few of them are high-frequency: 119 SNPs exceed the 0.3% threshold, according to Yuan et al., 2020. Based on the mutations, eight distinct viral clades have been reported by GISAID and twelve by Nextstrain (by March 2021). Specific SARS-CoV-2 variants caused the most concern. A “British variant” (also B.1.1.7, 20I/501Y.V1 or Variant of Concern 202012/01) caused a travel ban in December 2020 because of its increased transmissibility (Chand et al., 2020). A “South African” variant (B.1.351/ 501Y.V2) is thought to be more abundant in healthy young people and result in a more severe disease course in those cases (Tegally et al., 2020). A “Brazilian” variant (P.1) is presumed to be more infectious (Faria et al., 2021). The most recent B.1.617.2 (delta) variant struck India in March 2021 and quickly became the most reported variant (Lopez Bernal et al., 2021). Most frequently, mutations are found in SARS-CoV-2 sequences coding for spike (S) protein, RNA-dependent RNA polymerase (RdRp), and nucleoprotein (N). Despite a vast amount of knowledge accumulating daily, the exact consequences of most viral mutations are unknown (Yin 2020). Current updates on the positions and functions of viral regions are presented in Table 1.

**Table 1.** SARS-CoV-2 genes, their genomic positions, length, and function as assumed to date (functions according to NCBI Gene, Miorin, et al., 2020, Zhou et al., 2021, Park 2020).

| **Viral gene** | **Genomic position**<br>**(According to UCSC Genome Browser)** | **Gene length** | **Presumable main function** |
| --- | --- | --- | --- |
| ORF1ab | 266-21555 | 21290 | Codes for polyproteins PP1ab and PP1a which allow for viral replication, transcription, and other functions |
| S | 21563-25384 | 3822 | Provides cell entry |
| ORF3a | 25393-26220 | 828 | Activates the NLRP3 inflammasome; may contribute to virus replication and pathogenesis |
| E | 26245-26472 | 228 | Facilitates virion assembly within cells |
| M | 26523-27191 | 669 | Facilitates virion assembly within cells |
| ORF6 | 27202-27387 | 186 | Likely promotes viral replication |
| ORF7a | 27394-27759 | 366 | Likely interacts with immune cells |
| ORF7b | 27756-27887 | 132 | The structural component of the SARS-CoV-2 virion |
| ORF8 | 27894-28259 | 366 | Downregulates MHC-I |
| N | 28274-29533 | 1260 | Packages viral genome inside the capsid |
| ORF10 | 29558-29674 | 117 | Not identified |

Although any results of genomic variation analysis obtained using a bioinformatic approach should be considered with caution until experimental confirmation (Kaur et al., 2020, Yin 2020), bioinformatics plays an important role in unraveling the viral mysteries. Overall, SARS-CoV-2 genome mutations are hypothesized to impact viral transmissivity, case fatality risk, and numerous other features. In this paper, we describe our research aimed at analyzing 329,942 viral FASTA sequences obtained from human hosts to observe mutational changes and explore the accompanying data.

## Materials and Methods

Data for 329,942 SARS-CoV-2 genomes isolated from human hosts were retrieved from the GISAID database, along with additional information (records from the 24th of December 2019 until the 8th of January 2021). Custom code for revealing insertions, deletions, and SNPs was used alongside the pairwise2 local tool (https://biopython.org/docs/1.78/api/Bio.pairwise2.html) from the BioPython library (Python version 3.7, BioPython version 1.78; https://biopython.org/). Alignments were done for every viral gene separately, except ORF1ab which was not considered in the present research. Every gene was aligned to a reference sequence, and final positions were calculated on a reference genome (accession number MN908947.3) (Wu et al., 2020). Genomic positions were retrieved from the UCSC genome browser (see Table 1). We used Pandas (version 1.2; https://pandas.pydata.org/), Matplotlib (version 3.3; https://matplotlib.org/), and Seaborn (version 0.11; https://seaborn.pydata.org/installing.html) to visualize the data. Cluster analysis was executed using HDBScan (version 0.8; https://hdbscan.readthedocs.io/en/latest/) and visualized with t-SNE (t-distributed stochastic neighbor embedding; sklearn version 0.23; https://scikit-learn.org/stable/modules/generated/sklearn.manifold.TSNE.html). The occurrence cut off (0.3% or 989 records) was selected as the starting point for clustering parameters search, resulting in the minimum cluster size being set at 2000, and “minimum samples” at 5. Clustering was performed using data on SNPs and deletions whose frequency exceeded 0.3% in the present research. The clustering parameters that yielded a minimum number of clusters, subject to the condition of at least 989 records in one cluster, were determined as suitable for the research. Only sequences more than 26,000 nucleotides long were included in the study, since the smaller sequences did not allow us correctly align all genes of interest. Data filtering was completed as follows: 100% match to a reference genome was required to consider a sequence highly conservative, more than 99% match - to consider it moderately conservative, alignments in a range from 99% to 93% match were marked as low conservative. The genomes that were less than 93% similar to the reference sequence contained low-quality sequences and were excluded from further analysis. As these cutoffs were determined experimentally and we considered all the viral genes separately, we were free from simply deleting all the records containing ambiguous/unidentified symbols (“N”, “Y” etc.). Instead, examining genes separately increased the number of sequences that could be used in the research. If unidentified symbols were determined in the aligned gene, and their count was not equal to the count of SNPs, the sequence was included in the research. Statistical significance was measured using a t-test and Bonferroni correction (for two parameters – age and gender). Correlation was measured using Pearson correlation coefficient.

## Results

By the 8th of January 2021, the GISAID database had SARS-CoV-2 records deposited by 142 countries. Even though more than 329,000 records had been uploaded up until then, these data has limited research potential due to several significant problems. First, some of the uploaded sequences are dramatically smaller than the reference sequence (e.g. <5000 nucleotides) or contain an enormous (more than 7% of each gene of interest) number of ambiguous letters (Fig.1. represents the sequence size range obtained for the data used in current research; the smallest sequences were mostly obtained by Sanger sequencing). Another weakness is the lack of automation/control in terms of data entry to the system. That drawback led to numerous misspellings and data variants, along with missing information. Thus, the “collection date” field may include a year, a month, and a date, contain only the year, or, for some records, have a wrong year (e.g., 2002 instead of 2020). “Gender” and “Patient age” parameters are filled only for 23.3% and 23.1% of the records, respectively. The least informative for research is “Patient status,” which is not only filled for just 6.9% but also contains hardly interpretable data. Records’ bias is another problem. The prevalent number of genomes is uploaded by the United Kingdom (45.3%), USA (18.3%), Denmark (6.7%), and Australia (5.1%), with other countries’ input ranging from 3 to less than 1 percent of all records.

**Fig.1.**
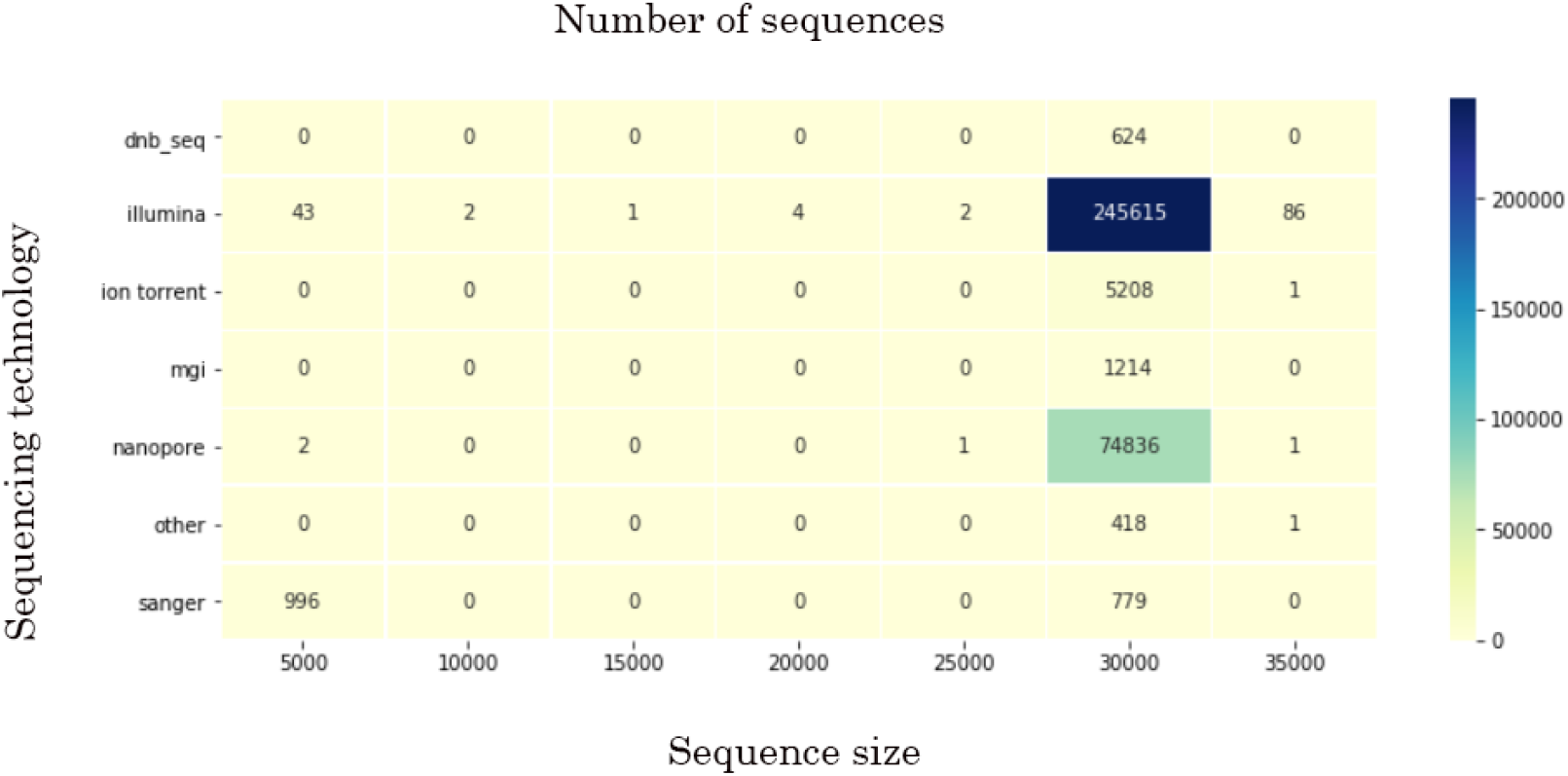
The sequence size range data obtained for the data used in current research.

Mean age was determined as 48 (confidence intervals (95% CI): 47.8, 48.1). Although gender values for a studied cohort equal 52% of males and 48% of females (95% CI: 0.51, 0.52), mean gender values in some countries significantly declined from these numbers. Most gender inequality among records was noted in Saudi Arabia (80% males among 446 gender-filled records, p << 0.001), Singapore (75% among 1584 gender-filled records, p << 0.001), and Bangladesh (68% among 586 gender-filled records, p << 0.001) in terms of male prevalence, and South Africa (64% of females among 2591 gender-filled records, p << 0.001), Lithuania (61% among 193 gender-filled records, p << 0.001) and Russia (57% among 1545 gender-filled records, p << 0.001) in terms of female prevalence. The highest mean age was revealed in records submitted by the United Kingdom (59.6, p << 0.001) and France (59.5, p << 0.001), the lowest – by United Arab Emirates (35.6, p << 0.001), Gambia (37, p << 0.001), Oman (37.3, p << 0.012) and Bangladesh (38.5, p << 0.001). Only the countries which submitted more than 100 parameter-filled records are mentioned above. For full data, see Supplement 1. The records’ bias also affected the patients’ status. Some countries presumably uploaded the records with predominantly one or another status (e.g., out of all records uploaded by Brazil, 40% contained patient status “Dead”).

### Genomic data

The data were considered for every viral gene separately, except for ORF1ab, which was not considered in the present research. While filtering the data to include only good-quality sequences (Table 2), we encountered an obscure phenomenon concerning an ORF7b gene. Nearly 11,290 (out of 329,942) FASTA records were featured by a similar pattern consisting of 52 “N”s (for most, genomic coordinates: 27757 - 27808). Sixty percent of that data was obtained using Nanopore sequencing (although 22.7% of all the data was acquired by that sequencing technology). Besides sequencing technology, the problem may derive from a particular assembly method, more precisely – from choosing a wrong method or unsuitable parameters, such as k-mer size. Assembly method data are present in 45.9% of all records (while sequencing technology – 99.9%). For records where sequences contain stretches with 52 “N”s, the “assembly method” is filled for 23.5%. Since we could not estimate the assembly method and its parameters, we investigated the most prevalent methods among records containing stretches with 52 “N”s. Although further research is limited due to too many variations created by manual system entry, the top assembly method was either “mapping consensus” or numerous variations of ARCTIC.

**Table 2.**
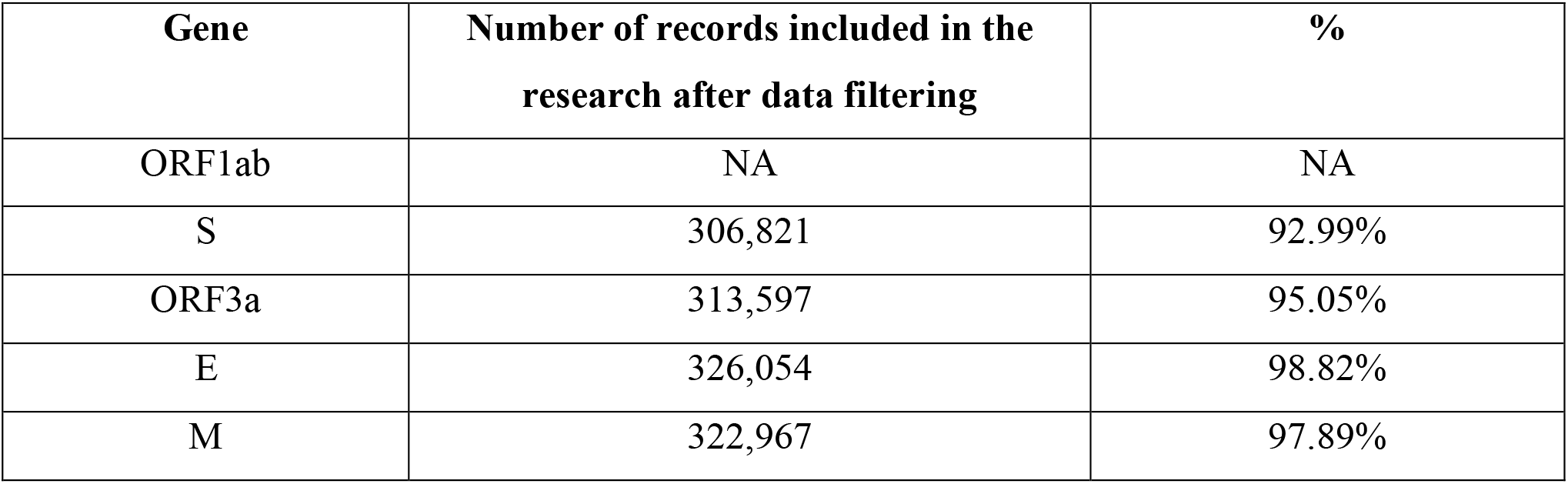
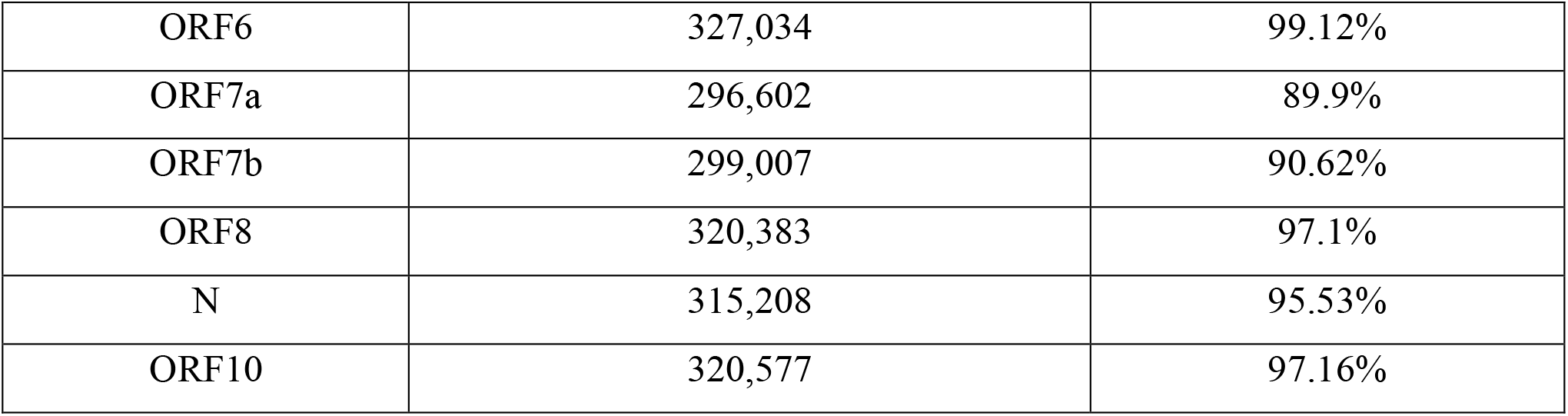
Number of records included in the research after data filtering, except for ORF1ab, which was not considered in the present research.

### Conservation

Analyzing the conservation of the genes allowed us to get some insights into their importance for the virus and potential treatment (Table 3).

**Table 3.**
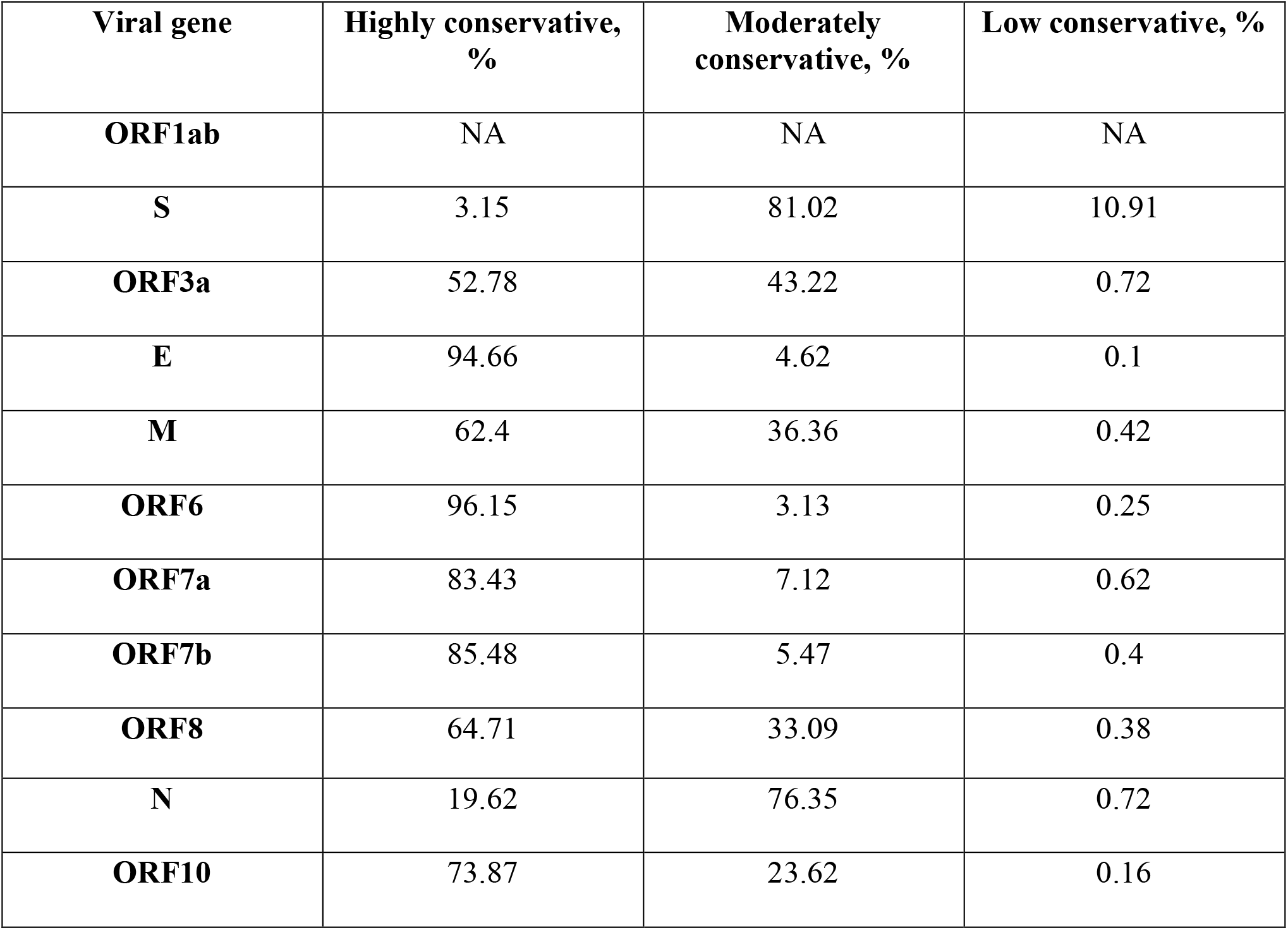
Comparing the conservation of viral genes.

| **Viral gene** | **Highly conservative, %** | **Moderately conservative, %** | **Low conservative, %** |
| --- | --- | --- | --- |
| **ORF1ab** | NA | NA | NA |
| **S** | 3.15 | 81.02 | 10.91 |
| **ORF3a** | 52.78 | 43.22 | 0.72 |
| **E** | 94.66 | 4.62 | 0.1 |
| **M** | 62.4 | 36.36 | 0.42 |
| **ORF6** | 96.15 | 3.13 | 0.25 |
| **ORF7a** | 83.43 | 7.12 | 0.62 |
| **ORF7b** | 85.48 | 5.47 | 0.4 |
| **ORF8** | 64.71 | 33.09 | 0.38 |
| **N** | 19.62 | 76.35 | 0.72 |
| **ORF10** | 73.87 | 23.62 | 0.16 |

**Table 4.** Mutations that are most different in terms of gender.

| SNP | Amino acid change | Mean gender (binary format, 1 is male) | Gender filled records | Country(ies) uploaded most of the records for SNP | Gender filled records among country's uploads for SNP |
| --- | --- | --- | --- | --- | --- |
| **S** |  |  |  |  |  |
| T24910C | T1116T | 0.41 | 367 | UK | 61 |
| T24088C | G842G | 0.31 | 49 | UK | 44 |
| C23929T | Y789Y | 0.76 | 1744 | Singapore, India | 943 and 506 |
| T22020C | M153T | 0.42 | 216 | Japan | 0 (most gender data for the SNP are uploaded by Russia and Lithuania, mean gender 0.43 and 0.4) |
| G24781T | K1073N | 0.40 | 63 | UK | 22 |
| **ORF3a** |  |  |  |  |  |
| G25617T | K75N | 0.39 | 72 | UK | 48 |
| G25757T | R122I | 0.3 | 10 | Denmark | 9 |
| G25947T | Q185H | 0.39 | 140 | UK | 16 |
| **M** |  |  |  |  |  |
| C26645T | N41N | 0.39 | 152 | UK | 38 |
| **ORF7a** |  |  |  |  |  |
| C27513T | Y40Y | 0.42 | 99 | UK | 13 |
| **ORF8** |  |  |  |  |  |
| A28095T | K68@ | 0.29 | 24 | UK | 8 |
| **N** |  |  |  |  |  |
| C28311T | P13L | 0.74 | 1886 | Singapore | 938 |

### Insertions and deletions

No insertions with a frequency greater than 0.3% were found. Two deletions were identified in the S gene: 21765-ATACATG>A with 4.67% frequency and 21991-TTTA>T with 2.94% frequency.

### SNPs

Analyzing genomic data considering the date axis, we were not merely able to determine the most frequent mutations but also reveal their changes through the year (Supplement 2 contains data on SNPs occurring with more than 0.3% frequency among 329,942 viral genomes. Supplement 3 contains charts representing changes by month for each mutation).

#### Clustering

HDBScan yielded 43 clusters (based on data on SNPs and deletions with a frequency greater than 0.3%; Fig.2). Some data did not fit any cluster.

**Fig.2.**
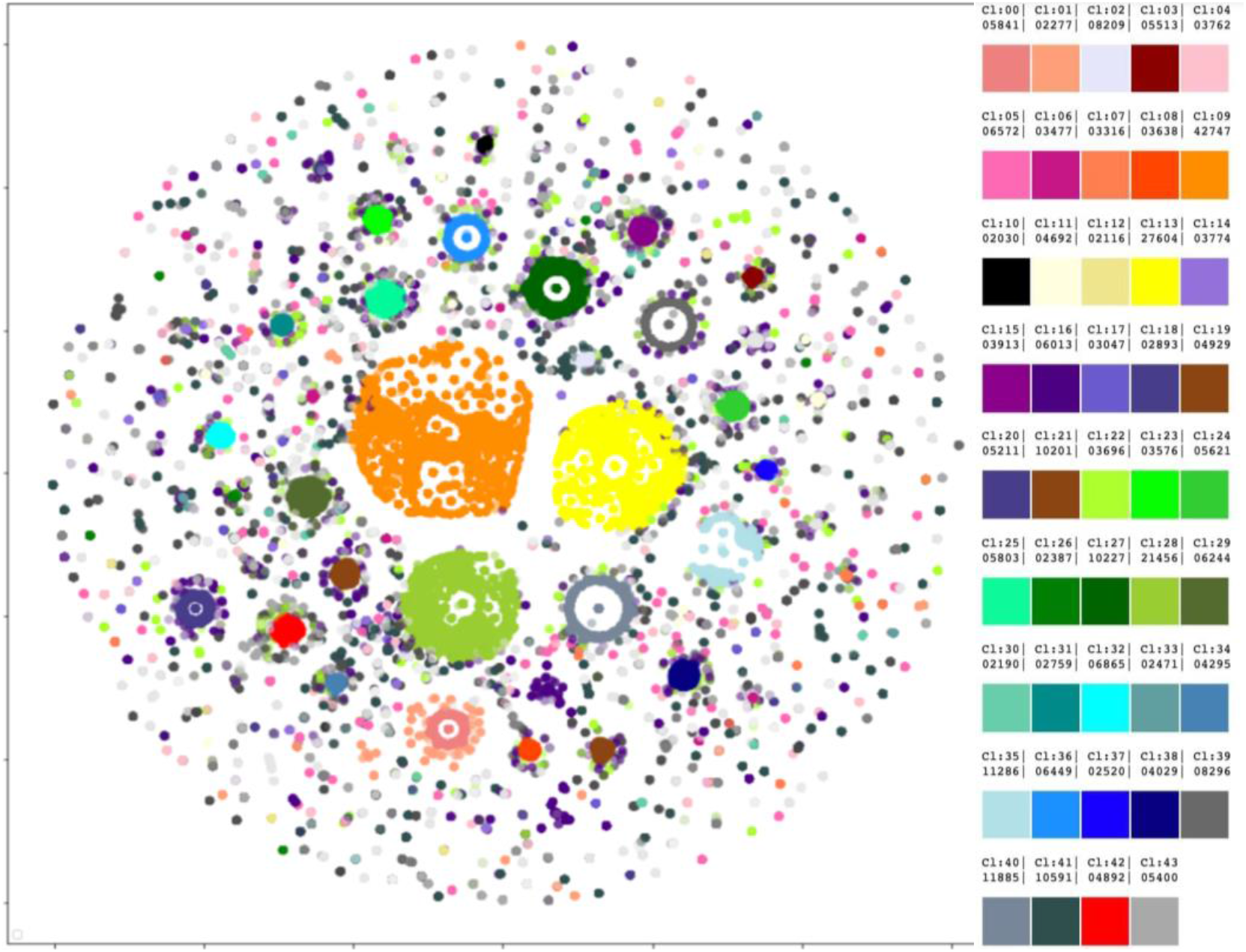
Forty-three clusters were revealed by HDBScan. Legend on the right contains cluster numbers and color schemes.

A number of the forty-three clusters grouped by concomitant mutations present interesting data. Cluster #0 (size regarding all studied genomes - 1.77%) contains all mutations from a “British variant”, except an SNP in the M gene (ORF1ab mutations were not considered due to the specificity of the research), in 100% records of the cluster. Four mutations are present in the cluster with 100% frequency - G28881A, G28882A, and G28883C in the N gene and A23403G – in the S. Cluster #1 also demonstrates interesting results. Containing 0.69% of all records, it also has the aforementioned mutations from the “British variant” and the following mutations: A28095T in ORF8 (frequency in the cluster - 49.98%), G28881A, G28882A, and G28883C in the N gene, A23403G – in S (100% each), and G25437T in ORF3a (31.58%). Cluster #20 shows significantly different parameters in terms of age and gender. The cluster includes one mutation in ORF3a (G26144T) and is characterized by a mean age of 57 and a gender ratio of 50.46 males to 49.54 females. One cluster featured by the increased mean age is #25. It is characterized by the age of 53 and can be described by 5 mutations occurring with different frequency: A23403G (99%), G25563T (87%), C27964T (87%), C28977T (10%), and C23731T (2%). One cluster is featured by the mean age of 43 (cluster #34). The cluster is represented by 9 mutations: C28869T (100%), C27964T (100%), A23403G (100%), G25563T (100%), G25907T (100%), C28472T (99%), G29402T (23%), A22255T (17%), G23593T (4%). Two clusters, #13 and #39, show an altered male to female ratio. Cluster #13 is featured by 54.8% of males and 3 mutations: A23403G (100%), G25563T (100%), C26735T (5%); cluster #39 is characterized by 46.31% of males and 8 mutations: A23403G (99%), G22992A (99%), G23401A (99%), G28881A (99%), G28882A (99%), G28883C (99%), C27059T (7%), C22480T (6%). Mutations found in samples uploaded mainly by Denmark and Australia form two clusters, each containing 8 mutations. The following mutations are found in clusters from Denmark and Australia, respectively: C26735T (100%), T26876C (100%), G25563T (100%), C25710T (100%), G29399A (100%), A23403G (99%), G22992A (99%), C27434T (13%); A23403G (99%), G22992A (99%), G23401A (99%), G28881A (99%), G28882A (99%), G28883C (99%), C27059T (7%), C22480T (6%).

#### Concomitant mutations

According to a correlation coefficient matrix, 69 mutations have correlations with at least one other mutation (Fig.3; larger resolution and lower cutoff may be found in Supplement 4). In total, 160 pairs with a correlation coefficient greater than 0.7 were found (Supplement 5).

**Fig.3.**
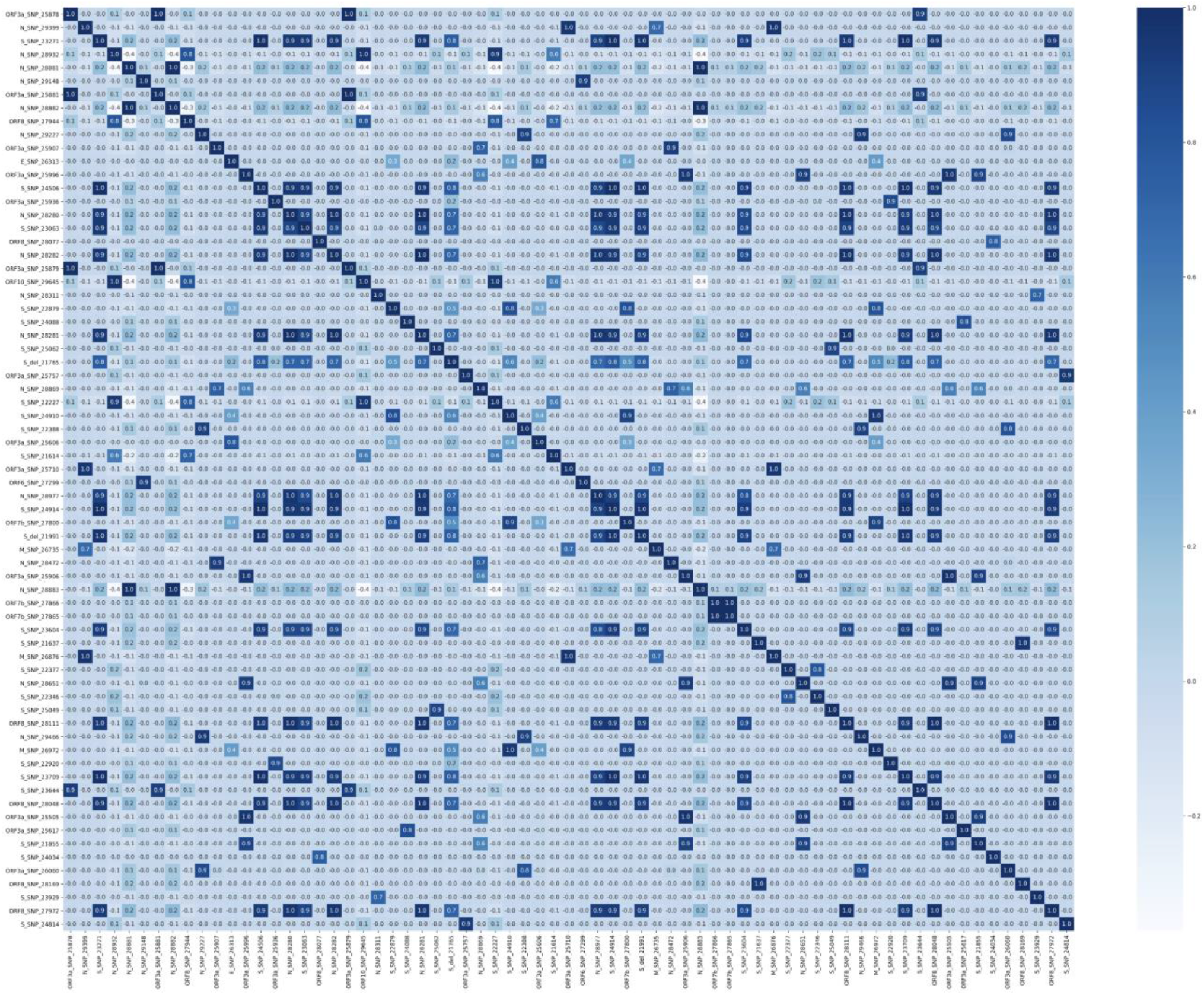
Correlation coefficient matrix based on mutations with a frequency greater than 0.3% and correlation coefficient starting from 0.7.

## Discussion

The statistical and bioinformatic analysis of 329,942 records obtained from the GISAID database yielded data concerning many areas, from database design and medical care issues to genomic mutations and their probable effects. The abovementioned results are discussed below.

### 1. Treatment targets: conservative sites

At the moment, one of the most promising treatment and vaccine targets is the S protein, which enables the virus to enter human cells and is already targeted in such vaccines as Gam-COVID-Vac (Sputnik V), Oxford/AstraZeneca, Pfizer/BioNTech, and Moderna (Dai et al., 2020). However, the S gene has dramatically changed since the reference genome was first published – only 3.15% of the analyzed sequences totally match the reference sequence. Viral genes that changed least during the pandemic are ORF6 and E (96.15% and 94.66% of the sequences have 100% match to the reference sequence, respectively). Although E protein acts together with an M protein in order to accomplish a virion assembly within the cells (V’kovski et al., 2021), the gene has changed dramatically less compared to M (62.4% of the sequences are highly conservative). According to these data, ORF6 and E are extremely perspective targets for treatment/vaccine development. At the moment, the E gene is only used as one of two qRT–PCR targets in SARS-CoV-2 detection assays by Roche (cobas® SARS-63 CoV-2 test). However, it is already known that the E protein of SARS-CoV-2 is highly immunogenic (Bhattacharya et al., 2021, Tilocca et al., 2020). Researchers have attempted drug discovery concerning both E and ORF6. One group determined a drug-binding site of E’s transmembrane domain using a solid-state NMR spectroscopy (Mandala et al. 2020). ORF6 is able to suppress both primary interferon production and interferon signaling. It is thought that SARS-CoV-2 with deleted ORF6 may be discussed in terms of intranasal live-but-attenuated vaccine invention (Yuen et al., 2020). Since ORF6 is one of three proteins causing the highest toxicity when overexpressed in human 293 T cells, and it also interacts with nucleopore proteins (RAE1, XPO1, RANBP2, and nucleoporins), treatment with an XPO1 inhibitor, Selinexor, was considered. Selinexor was found to reduce ORF-6-induced toxicity in human 293 T cells (Lee et al., 2021). Other groups found that Gliclazide and Memantine may inhibit E protein’s channel activity and Belachinal, Macaflavanone E, and Vibsanol B may inhibit the protein’s function (Tomar et al., 2020; Gupta et al., 2020).

### 2. Сlustering

According to clustering results, it may be proposed that, although “British variant” mutational contents may not be expanded due to the absence of the concomitant mutations in the general cohort, there is a proportion of people who got infected with its distinct subtype. The subtype may be characterized by four to six additional mutations, with four being a more frequent option (G28881A, G28882A, and G28883С in the N gene, A23403G – in S, A28095T in ORF8, G25437T in ORF3a). Both clusters containing the “British variant” mutations are also the most recent, with a mean upload time of 1.2 months ago (count back from the 8th of January 2021). A mutation in ORF3a (G26144T) that forms a cluster and is featured by increased age (57) and significantly different males to females ratio (50.46:49.54), has presumably disappeared from the population and was last noted in the uploads in September 2020. Due to increased age among patients carrying the virus with the mutation, it may be proposed to have increased virulence. Two clusters of mutations are found to be associated with increased or decreased patient age (57 and 43), while two other clusters are featured by shifted male:female ratio compared to mean in the studied group: increased proportion of males in one (54.8%), and of females – in the other (46.31%). Distinct clusters are formed by mutations in samples uploaded predominantly by Denmark and Australia (8 mutations in each), which lets us speculate on the existence of so-called “Danish” and “Australian” variants.

### 3. Concomitant mutations

Current research shows that some mutations often present together with one or more others. In total, 160 pairs of mutations with a correlation coefficient greater than 0.7 were found. Most studies in this direction focus on certain concomitant mutations. For example, D614G is often considered together with P323L. Some researchers suggest inability of D614G to cause viral success when presented alone (Ilmjärv et al., 2020; Wang et al., 2020). T85I is noted to co-occur with Q57H, and P504L – with Y541C (Wang et al., 2020). Also, R203K and G204R in the N gene were found to occur together with high frequency (Rahman M. S. et al., 2021) which is confirmed in our research. G28881A is concomitant with G28882A and G28883С (r = 0.998), but with no other mutation. Variants of concern (e.g. B.1.1.7, B.1.351, P.1) contain co-occurring mutations, as well. However, to our knowledge, there are no publications analyzing concomitant mutations on a large scale. Therefore, our work shows this subject as a potentially fruitful ground for novel research.

### 4. The most frequent mutations

The most frequent mutation in the analyzed genes is a mutation in the S gene - A23403G (D614G), which is found in 94.15% of all studied genomes and in 99.9% of genomes uploaded in December 2020. D614G is considered to be more infectious than the ancestral form, but not associated with increased disease severity (Yurkovetskiy L. et al., 2020., Korber et al., 2020). Mutations with more than 20% frequency were found in different genes. In S, it’s C22227T (A222V) with 22.25%. It was found in 53.8% of all uploaded sequences in November 2020 and assumed to influence viral transmissivity and antigenicity (Hodcroft et al., 2020; Bartolini et al., 2020). The mutation might, however, enhance the ability of the protein to interact with the environment (Lon et al., 2021). The M gene is also featured by a frequent mutation - C26801G (L93L) is found in 21.82%, which constitutes between 53.4% and 43.2% of all uploads from November 2020 to December 2020. The assumed consequences of the mutation are yet to be described. The ORF3a gene has a G25563T (Q57H) mutation, found in 21.41% of the genomes. The N gene is featured by 4 mutations with a frequency greater than 20%: G28881A (R203K), G28882A (R203R), G28883С (R203R), and C28932T (A220V). Interestingly, Q57H and R203K were found to cause substantial changes in protein structures (RMSD ≥ 5.0 Å). The mutations are also thought to affect binding affinity of intraviral protein interactions (Wu et al., 2021). Last, one most frequently occurring mutation found in ORF10, G29645T (V30L), is present in 22.03% of uploads in a general group, and 44.6% of all uploads from December 2020. At the moment, it is proposed that ORF10 may not be a protein-coding gene with its premature termination not affecting viral fitness or transmissivity (Pancer et al, 2020).

### 5. Disappearing mutations potentially decrease viral fitness

Only three mutations have not been noted in the uploads since the beginning of December 2020: G26144T (G251V) and G25979T (G196V) in ORF3a, which were last uploaded around September 2020 and early December 2020, respectively, and a C28836T (S188L) in the N gene, which was last seen around early to middle November. G251V is revealed to result in the loss of a phosphatidylinositol-specific phospholipase X-box domain and a creation of a serine protease cleavage site (Issa et al, 2020). Another work states that G251V and G196V might influence virulence, infectivity, ion channel activity, and viral release (Coronaviridae Study Group of the International Committee on Taxonomy of Viruses, 2020). Might disappearing mutations impact viral fitness or human survival? The data is yet incomplete. However, in the present research G26144T (G251V) was found to create a cluster on its own; the mutation is featured by increased age (57) and increased proportion of women compared to the general cohort.

### 6. Novel mutations

The most recent mutation in the current analysis is A28111G (Y73C) in ORF8, which appeared in the uploaded data about early September 2020. The mutation is included in a B.1.1.7 (“British variant”) mutations’ list. In total, B.1.1.7 is featured by 23 mutations (Rambaut et al., 2020) and is preliminarily reported as possibly associated with an increased risk of death (Frampton D. et al, 2021). We detected 13/14 mutations not located in the ORF1ab region and associated with the variant in the analyzed data. A T26801C mutation in the M gene was not found among mutations with a frequency greater than 0.3%. Our data yielded two mutations in the same position (freq > 0.3%): C26801G and C26801T. The discrepancy could occur due to the differences in the reference sequences. This, however, cannot be verified as Rambaut et al. did not specify the reference sequence number. We have also considered two other variants that appeared lately - B.1.351 (a variant from South Africa) and P.1 (a variant from Brazil). Nevertheless, out of 8 and 14 non-ORF1ab mutations, respectively, only 2 and 3 were detected in our analysis among highly-present mutations. Consequently, it can be speculated that either a “British variant” has more transmissivity compared to the other two variants, or this result is due to a bias because of the number of the uploads.

### 7. GISAID database drawbacks lead to its severely limited research value

We have revealed that the major drawback of letting the users manually fill the fields of the records led to a loss of approximately 77% to 93% of the data, depending on the parameter. The absence of quality control for genomic data yielded a presence of many sequences significantly shorter or longer than the reference genome (ranging from <5000 to 34000 nt). Many laboratories uploading the data do so significantly later than the sample collection date, some even a year later, which may distort the bioinformatic analysis. Certain laboratories indicate a month and a year, or only a year, of sample collection, omitting the day, or day and month. An important analysis factor is that most data are uploaded by the United Kingdom, which creates an overall data bias towards the UK statistics. As time is a crucial factor in a pandemic, a database update can be recommended in order to increase its value and quality.

### 8. Gender inequality in the uploaded data may reflect medical care availability issues

The cohort studied in the current research is represented by 52% of males and 48% of females (mean values; gender was not indicated for a subset of records). However, among records uploaded by Saudi Arabia, Singapore, and Bangladesh men were present in 80%, 75%, and 68% of the records, respectively. Official statistics, male to female: Saudi Arabia - 58%:42%; Bangladesh 51%:49%, Singapore 52%:48% (https://data.worldbank.org/). While Saudi Arabia is known for limiting access to medical care for women without a male guardian (World Report 2020), Singapore, on the contrary, was ranked high (11th among 162 countries) for gender equality by the United Nations Development Programme last year (Human Development Report). The answer to this discrepancy most probably lies in the dormitories for migrants. In December of 2020, the Ministry of Health of Singapore declared that the majority of all COVID-19 cases occurred in migrant worker dormitories (“Measures to contain the COVID-19 outbreak in migrant worker dormitories”). Although Bangladesh has shown significant improvement in moving towards gender equality (according to Human Development Report 2020), a medical access problem for rural areas persists. Estimating the rates of female inequality concerning medical care, a paper from the National Institute of Medical Health states that female patients were about half in number compared to male patients (Nuri et al., 2019). Our research also highlights possible issues in terms of health care for males: South Africa, Lithuania, and Russia uploaded 64%, 61%, and 57% of female records, respectively (the top three countries are considered for a shift in male to female ratio for both genders). Official statistics, male to female: South Africa 49:51, Lithuania 46:54, and Russia 46:54 (https://data.worldbank.org/). There are no data on limited medical care options for men in South Africa, Lithuania, or Russia. Thus, it can be speculated that the current lack of male patients may derive from a strong idea of masculinity (e.g., men must be strong and health complaints mean weakness) (Colvin, 2017). Russian Minister of Healthcare noted in his speech, dated August 3rd, 2020, that Russian males address outpatient hospitals 2.5 times less often than females, with inpatient estimates being about the same for both genders (reported by TASS). One more explanation is that more people working in the areas related to abundant social contact (e.g., medicine, education) in these countries are women. We suppose that this distribution may also be considered in terms of hospitalization criteria and sex differences between distinct age groups, and therefore leave this question to be still open for discussion.

### 9. Gender and age-related mutations

Although mean age across gender-filled records in our cohort was determined as 48 and mean gender as 52% of males and 48% of females, some mutations were characterized by increased or decreased age and shift in male to female ratio. In total, two mutations in the S gene exhibited a decline in more than 10 points (years or percent) for both age and gender parameters. A T24814C (D1084D) could not be further analyzed due to the lack of data. Denmark uploaded most of the records for the SNP (99.28%), but none of them contained patient age or gender. Data for both age and gender relies only on 5 records and cannot reflect the situation. A G23311C (E583D) was predominantly uploaded by the UK (97.1%), so it may be considered with respect to the other UK statistics. Among the records containing the SNP, the numbers (27% males and 73% of females) were obtained using 140 gender-filled records. In total, gender among records uploaded by the UK (6275 records) is 50/50, however for the current SNP a solely UK number is 20:80, males to females. (107 records). The patient age for the SNP is 61 (139 records), among only UK records – 68 (mean age in the UK is 59). We have not found data on the mutation with respect to age/gender. The only interesting message was an article stating that this mutation co-mutates with infectivity-enhancing S protein mutations, such as D614G, which cannot yet explain our finding (Wang R. et al., 2020.). Beside the aforementioned data, there are 12 mutations that are 10 points different in terms of gender (Table 5), and 2 – in terms of age.

Consequently, only C23929T (Y789Y) and C28311T (P13L) may further be considered due to the number of gender-filled records. P13L is presumably associated with decreased deaths and significant changes altering the protein structure (Huang et al., 2020; Wu et al., 2020). Age-related changes were noted for the mutations in the S (A22255T) and E (T26424C) genes, with characteristic ages of 38 and 62, respectively. For A22255T, 97.31% of the sequences were uploaded by the USA and the total age-filled records’ number for the SNP is 122, most uploaded by the USA. The mean patient age for the USA is 49. For T26424C mutation, 97.96% of the sequences are uploaded by the UK, only 47 records are age-filled, most uploaded by the UK, where the mean patient age is 59. Increased age has been linked to the worst outcomes in those suffering from COVID-19. The mortality risk increases from 0-0.1% for children and adolescents under the age of 19 to 4.3–10.5% for the age group of 75–84 years. The most dramatic consequences are seen for individuals from 85 and older (up to 27.3% case fatality rate). Older patients get hospitalized more often (median age 74 vs median age of 43 for individuals in the outpatient care) and suffer from concomitant health issues (e.g. cardiovascular disorders, diabetes) which increase mortality rates by itself (Dowd et al., 2020, Promislow, 2020, Korber et al., 2020). Interestingly, it has been repeatedly noted that men seem to suffer from COVID-19 more severely than women (Conti, Younes, 2020), with males proposedly being hospitalized more often than females (e.g. Korber et al., 2020 report 67% of males versus 33% of females). Some mutations (for example, C27964T in ORF8) have been found to have gender dependence with a presumed ratio of 2:1 (Wang et al., 2020). Although the reasons why males seem to be more severely affected are not yet clear, there are certain hypotheses on the topic. For instance, is it known that a primary way of SARS-CoV-2 entrance to the body is through its connection to angiotensin-converting enzyme 2 (ACE2), a part of the human renin-angiotensin-aldosterone system (RAAS) (Zhou et al., 2020), and males show greater overall RAAS activity compared to females (Zalucky et al., 2014). Also, as increased mortality risk is associated with cardiovascular diseases (Yang et al., 2020), the greater percentage of these disorders and thrombosis in men may contribute to fatality increase among males. A higher case fatality rate could also result from the fact that in general, among intubated patients, men are more likely to acquire ventilator-associated pneumonia (Cook et al., 1998, Ahmed, Dumanski, 2020).

## Conclusions

In this paper, we have analyzed 329,942 SARS-CoV-2 records obtained from the GISAID database. We addressed the quality of the uploaded records, gender distribution, gene conservation, SNPs, insertions and deletions, clusters, and a correlation coefficient matrix. Our research shows that mutations occurring with high frequency (>0.3%) are not abundant and constitute 155 changes concerning all genes (except ORF1ab, which was not considered in a current work). Many mutations present with concomitant changes, which may alter their consequences for the virus or a human host. A big number of co-occurring mutations creates grounds for research on their meaning, as well as a probability of the occurrence in terms of novel mutations and concomitant variants. Conservation analysis suggests ORF6 and E genes as prospective treatment/vaccine targets due to their high conservation. Clustering allows speculations on the existence of a subtype of a B.1.1.7 variant, and the possible existence of variants specific to Denmark and Australia. Taken together, our results describe the genetic variability of SARS-CoV-2 and may be used for further research in different scientific areas.

## Acknowledgments

Authors thank Tatiana Tatarinova, Dallas Dorsey and Isaiah Knox from the University of La Verne, Nika Tsutskiridze, Tsotne Khetsuriani, Revaz Mgeladze, Nona Kuloshvili, Tinatin Mekvabishvili from Quantori, Artem Artemov from MedUni Wien and Alexander Mikov from Amedart for insightful comments and discussions.

## References

1. Qi, Furong, Shen Qian, Shuye Zhang, and Zheng Zhang. 2020. “Single Cell RNA Sequencing of 13 Human Tissues Identify Cell Types and Receptors of Human Coronaviruses.” Biochemical and Biophysical Research Communications 526 (1): 135–40.

2. Yin, Changchuan. 2020. “Genotyping Coronavirus SARS-CoV-2: Methods and Implications.” Genomics 112 (5): 3588–96.

3. World Health Organization: https://covid19.who.int/

4. Weber, Stefanie, Christina Ramirez, and Walter Doerfler. 2020. “Signal Hotspot Mutations in SARS-CoV-2 Genomes Evolve as the Virus Spreads and Actively Replicates in Different Parts of the World.” Virus Research 289 (November): 198170.

5. Walls, Alexandra C., Young-Jun Park, M. Alejandra Tortorici, Abigail Wall, Andrew T. McGuire, and David Veesler. 2020. “Structure, Function, and Antigenicity of the SARS-CoV-2 Spike Glycoprotein.” Cell 183 (6): 1735.

6. Sousa, Eric de, Dário Ligeiro, Joana R. Lérias, Chao Zhang, Chiara Agrati, Mohamed Osman, Sherif A. El-Kafrawy, et al. 2020. “Mortality in COVID-19 Disease Patients: Correlating the Association of Major Histocompatibility Complex (MHC) with Severe Acute Respiratory Syndrome 2 (SARS-CoV-2) Variants.” International Journal of Infectious Diseases: IJID: Official Publication of the International Society for Infectious Diseases 98 (September): 454–59.

7. Kaur, Navpreet, Rimaljot Singh, Zahid Dar, Rakesh Kumar Bijarnia, Neelima Dhingra, and Tanzeer Kaur. 2021. “Genetic Comparison among Various Coronavirus Strains for the Identification of Potential Vaccine Targets of SARS-CoV2.” Infection, Genetics and Evolution: Journal of Molecular Epidemiology and Evolutionary Genetics in Infectious Diseases 89 (April): 104490.

8. Wang, Rui, Yuta Hozumi, Changchuan Yin, and Guo-Wei Wei. 2020. “Decoding SARS-CoV-2 Transmission and Evolution and Ramifications for COVID-19 Diagnosis, Vaccine, and Medicine.” Journal of Chemical Information and Modeling 60 (12): 5853–65.

9. Zhou, Peng, Xing-Lou Yang, Xian-Guang Wang, Ben Hu, Lei Zhang, Wei Zhang, Hao-Rui Si, et al. 2020. “Addendum: A Pneumonia Outbreak Associated with a New Coronavirus of Probable Bat Origin.” Nature 588 (7836): E6.

10. Zhu, Na, Dingyu Zhang, Wenling Wang, Xingwang Li, Bo Yang, Jingdong Song, Xiang Zhao, et al. 2020. “A Novel Coronavirus from Patients with Pneumonia in China, 2019.” The New England Journal of Medicine 382 (8): 727–33.

11. Wang, Rui, Jiahui Chen, Kaifu Gao, Yuta Hozumi, Changchuan Yin, and Guowei Wei. 2020. “Characterizing SARS-CoV-2 Mutations in the United States.” Research Square, August. https://doi.org/10.21203/rs.3.rs-49671/v1.

12. Yuan, Fangfeng, Liping Wang, Ying Fang, and Leyi Wang. 2020. “Global SNP Analysis of 11,183 SARS-CoV-2 Strains Reveals High Genetic Diversity.” Transboundary and Emerging Diseases, November. https://doi.org/10.1111/tbed.13931.

13. Chand, Meera, Hopkins, Susan, Dabrera, Gavin, Achison, Christina, Barclay, Wendy, Ferguson, Neil, Volz, Erik, Loman, Nick, Rambaut, Andrew, Barrett, Jeff. 2020. Investigation of novel SARS-COV-2 variant: Variant of Concern 202012/01 (Report). Public Health England. p. 2.

14. Tegally, Houriiyah, Eduan Wilkinson, Marta Giovanetti, Arash Iranzadeh, Vagner Fonseca, Jennifer Giandhari, Deelan Doolabh, et al. 2020. “Emergence and Rapid Spread of a New Severe Acute Respiratory Syndrome-Related Coronavirus 2 (SARS-CoV-2) Lineage with Multiple Spike Mutations in South Africa.” medRxiv. https://www.medrxiv.org/content/10.1101/2020.12.21.20248640v1.full.

15. Faria, Nuno R., Ingra Morales Claro, Darlan Candido, L. A. Moyses Franco, Pamela S. Andrade, Thais M. Coletti, Camila A. M. Silva, et al. 2021. “Genomic Characterisation of an Emergent SARS-CoV-2 Lineage in Manaus: Preliminary Findings.” Virological. https://www.icpcovid.com/sites/default/files/2021-01/Ep%20102-1%20Genomic%20characterisation%20of%20an%20emergent%20SARS-CoV-2%20lineage%20in%20Manaus%20Genomic%20Epidemiology%20-%20Virological.pdf.

16. Lopez Bernal, Jamie, Nick Andrews, Charlotte Gower, Eileen Gallagher, Ruth Simmons, Simon Thelwall, Julia Stowe, et al. 2021. “Effectiveness of Covid-19 Vaccines against the B.1.617.2 (Delta) Variant.” The New England Journal of Medicine, July. https://doi.org/10.1056/NEJMoa2108891.

17. Miorin, Lisa, Thomas Kehrer, Maria Teresa Sanchez -Aparicio, Ke Zhang, Phillip Cohen, Roosheel S. Patel, Anastasija Cupic, et al. 2020. “SARS-CoV-2 Orf6 Hijacks Nup98 to Block STAT Nuclear Import and Antagonize Interferon Signaling.” Proceedings of the National Academy of Sciences of the United States of America 117 (45): 28344–54.

18. Zhou, Ziliang, Chunliu Huang, Zhechong Zhou, Zhaoxia Huang, Lili Su, Sisi Kang, Xiaoxue Chen, et al. 2021. “Structural Insight Reveals SARS-CoV-2 ORF7a as an Immunomodulating Factor for Human CD14+ Monocytes.” iScience 24 (3): 102187.

19. Park, Matthew D. 2020. “Immune Evasion via SARS-CoV-2 ORF8 Protein?” Nature Reviews. Immunology 20 (7): 408.

18. Wu, Fan, Su Zhao, Bin Yu, Yan-Mei Chen, Wen Wang, Zhi-Gang Song, Yi Hu, et al. 2020. “A New Coronavirus Associated with Human Respiratory Disease in China.” Nature 579 (7798): 265–69.

19. Dai, Lianpan, and George F. Gao. 2021. “Viral Targets for Vaccines against COVID-19.” Nature Reviews. Immunology 21 (2): 73–82.

20. V’kovski, Philip, Annika Kratzel, Silvio Steiner, Hanspeter Stalder, and Volker Thiel. 2021. “Coronavirus Biology and Replication: Implications for SARS-CoV-2.” Nature Reviews. Microbiology 19 (3): 155–70.

21. Bhattacharya, Shreya, Arundhati Banerjee, and Sujay Ray. 2021. “Development of New Vaccine Target against SARS-CoV2 Using Envelope (E) Protein: An Evolutionary, Molecular Modeling and Docking Based Study.” International Journal of Biological Macromolecules 172 (March): 74–81.

22. Tilocca, Bruno, Alessio Soggiu, Maurizio Sanguinetti, Gabriele Babini, Flavio De Maio, Domenico Britti, Alfonso Zecconi, Luigi Bonizzi, Andrea Urbani, and Paola Roncada. 2020. “Immunoinformatic Analysis of the SARS-CoV-2 Envelope Protein as a Strategy to Assess Cross-Protection against COVID-19.” Microbes and Infection / Institut Pasteur 22 (4-5): 182–87.

23. Mandala, Venkata S., Matthew J. McKay, Alexander A. Shcherbakov, Aurelio J. Dregni, Antonios Kolocouris, and Mei Hong. 2020. “Structure and Drug Binding of the SARS-CoV-2 Envelope Protein Transmembrane Domain in Lipid Bilayers.” Nature Structural & Molecular Biology 27 (12): 1202–8.

24. Yuen, Chun-Kit, Joy-Yan Lam, Wan-Man Wong, Long-Fung Mak, Xiaohui Wang, Hin Chu, Jian-Piao Cai, et al. 2020. “SARS-CoV-2 nsp13, nsp14, nsp15 and orf6 Function as Potent Interferon Antagonists.” Emerging Microbes & Infections 9 (1): 1418–28.

25. Lee, Jin-Gu, Weiliang Huang, Hangnoh Lee, Joyce van de Leemput, Maureen A. Kane, and Zhe Han. 2021. “Characterization of SARS-CoV-2 Proteins Reveals Orf6 Pathogenicity, Subcellular Localization, Host Interactions and Attenuation by Selinexor.” Cell & Bioscience 11 (1): 58.

26. Singh Tomar, Prabhat Pratap, and Isaiah T. Arkin. 2020. “SARS-CoV-2 E Protein Is a Potential Ion Channel That Can Be Inhibited by Gliclazide and Memantine.” Biochemical and Biophysical Research Communications 530 (1): 10–14.

27. Gupta, Manoj Kumar, Sarojamma Vemula, Ravindra Donde, Gayatri Gouda, Lambodar Behera, and Ramakrishna Vadde. 2021. “In-Silico Approaches to Detect Inhibitors of the Human Severe Acute Respiratory Syndrome Coronavirus Envelope Protein Ion Channel.” Journal of Biomolecular Structure & Dynamics 39 (7): 2617–27.

28. Ilmjärv, Sten, Fabien Abdul, Silvia Acosta-Gutiérrez, Carolina Estarellas, Ioannis Galdadas, Marina Casimir, Marco Alessandrini, Francesco Luigi Gervasio, and Karl-Heinz Krause. 2020. “Epidemiologically Most Successful SARS-CoV-2 Variant: Concurrent Mutations in RNA-Dependent RNA Polymerase and Spike Protein.” medRxiv. https://www.medrxiv.org/content/10.1101/2020.08.23.20180281v1.abstract.

29. Rahman, M. Shaminur, M. Rafiul Islam, A. S. M. Rubayet Ul Alam, Israt Islam, M. Nazmul Hoque, Salma Akter, Md Mizanur Rahaman, Munawar Sultana, and M. Anwar Hossain. 2021. “Evolutionary Dynamics of SARS-CoV-2 Nucleocapsid Protein and Its Consequences.” Journal of Medical Virology 93 (4): 2177–95.

30. Yurkovetskiy, Leonid, Xue Wang, Kristen E. Pascal, Christopher Tomkins-Tinch, Thomas P. Nyalile, Yetao Wang, Alina Baum, et al. 2020. “Structural and Functional Analysis of the D614G SARS-CoV-2 Spike Protein Variant.” Cell 183 (3): 739–51.e8.

31. Yurkovetskiy, Leonid, Xue Wang, Kristen E. Pascal, Christopher Tomkins-Tinch, Thomas P. Nyalile, Yetao Wang, Alina Baum, et al. 2020. “Structural and Functional Analysis of the D614G SARS-CoV-2 Spike Protein Variant.” Cell 183 (3): 739–51.e8.

32. Korber, Bette, Will M. Fischer, Sandrasegaram Gnanakaran, Hyejin Yoon, James Theiler, Werner Abfalterer, Nick Hengartner, et al. 2020. “Tracking Changes in SARS-CoV-2 Spike: Evidence That D614G Increases Infectivity of the COVID-19 Virus.” Cell 182 (4): 812–27.e19.

33. Hodcroft, Emma B., Moira Zuber, Sarah Nadeau, Katharine H. D. Crawford, Jesse D. Bloom, David Veesler, Timothy G. Vaughan, et al. 2020. “Emergence and Spread of a SARS-CoV-2 Variant through Europe in the Summer of 2020.” medRxiv : The Preprint Server for Health Sciences, November. https://doi.org/10.1101/2020.10.25.20219063.

34. Bartolini, Barbara, Martina Rueca, Cesare Ernesto Maria Gruber, Francesco Messina, Emanuela Giombini, Giuseppe Ippolito, Maria Rosaria Capobianchi, and Antonino Di Caro. 2020. “The Newly Introduced SARS-CoV-2 Variant A222V Is Rapidly Spreading in Lazio Region, Italy.” medRxiv. https://www.medrxiv.org/content/10.1101/2020.11.28.20237016v1.abstract.

35. Lon, J. R., B. Xi, B. Zhong, Y. Zheng, P. Guo, Z. Chen, and H. Du. 2021. “Molecular Dynamics Simulation Study of Effects of Key Mutations in SARS-CoV-2 on Protein Structures.” bioRxiv. https://www.biorxiv.org/content/10.1101/2021.02.03.429495v1.abstract.

36. Wu, Siqi, Chang Tian, Panpan Liu, Dongjie Guo, Wei Zheng, Xiaoqiang Huang, Yang Zhang, and Lijun Liu. 2021. “Effects of SARS-CoV-2 Mutations on Protein Structures and Intraviral Protein-Protein Interactions.” Journal of Medical Virology 93 (4): 2132–40.

37. Pancer, Katarzyna, Aleksandra Milewska, Katarzyna Owczarek, Agnieszka Dabrowska, Michał Kowalski, Paweł P. Łabaj, Wojciech Branicki, Marek Sanak, and Krzysztof Pyrc. 2020. “The SARS-CoV-2 ORF10 Is Not Essential in Vitro or in Vivo in Humans.” PLoS Pathogens 16 (12): e1008959.

38. Issa, Elio, Georgi Merhi, Balig Panossian, Tamara Salloum, and Sima Tokajian. 2020. “SARS-CoV-2 and ORF3a: Non-Synonymous Mutations and Polyproline Regions.” bioRxiv. https://doi.org/10.1101/2020.03.27.012013.

39. Coronaviridae Study Group of the International Committee on Taxonomy of Viruses. 2020. “The Species Severe Acute Respiratory Syndrome-Related Coronavirus: Classifying 2019 - nCoV and Naming It SARS-CoV-2.” Nature Microbiology 5 (4): 536–44.

40. Rambaut, Andrew, Nick Loman, Oliver Pybus, Wendy Barclay, Jeff Barrett, Alesandro Carabelli, Tom Connor, Tom Peacock, David L. Robertson, Erik Volz, on behalf of COVID-19 Genomics Consortium UK (CoG-UK). Preliminary genomic characterisation of an emergent SARS-CoV-2 lineage in the UK defined by a novel set of spike mutations. https://virological.org., 2020. https://virological.org/t/preliminary-genomic-characterisation-of-an-emergent-sars-cov-2-lineage-in-the-uk-defined-by-a-novel-set-of-spike-mutations/563

41. Frampton, Dan, Tommy Rampling, Aidan Cross, Heather Bailey, Judith Heaney, Matthew Byott, Rebecca Scott, et al. 2021. “Genomic Characteristics and Clinical Effect of the Emergent SARS-CoV-2 B.1.1.7 Lineage in London, UK: A Whole-Genome Sequencing and Hospital-Based Cohort Study.” The Lancet Infectious Diseases, April. https://doi.org/10.1016/S1473-3099(21)00170-5.

42. World Report 2020: https://www.hrw.org/world-report/2020/country-chapters/saudi-arabia

43. Human Development Report - Singapore: http://hdr.undp.org/en/countries/profiles/SGP

44. Measures to contain the COVID-19 outbreak in migrant worker dormitories: https://www.moh.gov.sg/news-highlights/details/measures-to-contain-the-covid-19-outbreak-in-migrant-worker-dormitories

45. Human Development Report 2020 - Bangladesh: http://hdr.undp.org/sites/all/themes/hdr_theme/country-notes/BGD.pdf

46. Nuri, Nazmun Nahar, Malabika Sarker, Helal Uddin Ahmed, Mohammad Didar Hossain, Fekri Dureab, Faith Agbozo, and Albrecht Jahn. 2019. “Overall Care-Seeking Pattern and Gender Disparity at a Specialized Mental Hospital in Bangladesh.” Materia Socio-Medica 31 (1): 35–39.

47. Colvin, Christopher J. 2017. “Gender, Health and Change in South Africa: Three Ways of Working with Men and Boys for Gender Justice.” Recherches Sociologiques et Anthropologiques : RS & A 48 (1): 109–24.

48. Russian Minister of Healthcare’s speech reported by TASS: https://tass.ru/nacionalnye-proekty/7926683

49. Huang, Chaolin, Yeming Wang, Xingwang Li, Lili Ren, Jianping Zhao, Yi Hu, Li Zhang, et al. 2020. “Clinical Features of Patients Infected with 2019 Novel Coronavirus in Wuhan, China.” The Lancet 395 (10223): 497–506.

50. Dowd, Jennifer Beam, Liliana Andriano, David M. Brazel, Valentina Rotondi, Per Block, Xuejie Ding, Yan Liu, and Melinda C. Mills. 2020. “Demographic Science Aids in Understanding the Spread and Fatality Rates of COVID-19.” Proceedings of the National Academy of Sciences of the United States of America 117 (18): 9696–98.

51. Promislow, Daniel E. L. 2020. “A Geroscience Perspective on COVID-19 Mortality.” The Journals of Gerontology. Series A, Biological Sciences and Medical Sciences 75 (9): e30–33.

52. Korber, B., W. Fischer, S. G. Gnanakaran, and H. Yoon. 2020. “Spike Mutation Pipeline Reveals the Emergence of a More Transmissible Form of SARS-CoV-2.” BioRxiv. https://www.biorxiv.org/content/10.1101/2020.04.29.069054v2.abstract.

53. Conti, P., and A. Younes. 2020. “Coronavirus COV-19/SARS-CoV-2 Affects Women Less than Men: Clinical Response to Viral Infection.” Journal of Biological Regulators and Homeostatic Agents 34 (2): 339–43.

54. Zhou, Peng, Xing-Lou Yang, Xian-Guang Wang, Ben Hu, Lei Zhang, Wei Zhang, Hao-Rui Si, et al. 2020. “Addendum: A Pneumonia Outbreak Associated with a New Coronavirus of Probable Bat Origin.” Nature 588 (7836): E6.

55. Zalucky, A. A., D. D. M. Nicholl, M. C. Mann, B. R. Hemmelgarn, T. C. Turin, J. M. Macrae, D. Y. Sola, and S. B. Ahmed. 2014. “Sex Influences the Effect of Body Mass Index on the Vascular Response to Angiotensin II in Humans.” Obesity 22 (3): 739–46.

56. Yang, Jing, Ya Zheng, Xi Gou, Ke Pu, Zhaofeng Chen, Qinghong Guo, Rui Ji, Haojia Wang, Yuping Wang, and Yongning Zhou. 2020. “Prevalence of Comorbidities and Its Effects in Patients Infected with SARS-CoV-2: A Systematic Review and Meta-Analysis.” International Journal of Infectious Diseases: IJID: Official Publication of the International Society for Infectious Diseases 94 (May): 91–95.

57. Cook, D. J., and M. H. Kollef. 1998. “Risk Factors for ICU-Acquired Pneumonia.” JAMA: The Journal of the American Medical Association 279 (20): 1605–6.

58. Ahmed, Sofia B., and Sandra M. Dumanski. 2020. “Sex, Gender and COVID-19: A Call to Action.” Canadian Journal of Public Health. Revue Canadienne de Sante Publique 111 (6): 980–83.

