## Supplement 2 for "Analysis of 329,942 SARS-CoV-2 records retrieved from GISAID database"

**Supplement 2. SNPs occurring with more than 0.3% frequency among 329,942 viral records obtained from GISAID. “Months ago” are counted back from January 8, 2021. We use “@” to indicate a stop codon.**

| **#** | **Genomic**  **change** | **Amino**  **acid**  **change** | **Frequency**  **%** | **Most records**  **containing SNP are uploaded by** | **% for**  **the**  **country*** | **SNP**  **age** | **SNP**  **gender** | **Uploaded months ago, min** | **Uploaded months ago, max** |
| --- | --- | --- | --- | --- | --- | --- | --- | --- | --- |
| **S** | | | | | | | | | |
| 1. | A23403G | D614G | 94.15% | UK | 46.46% | 48 | 0.51 | 0.2 | 12.5 |
| 2. | C22227T | A222V | 22.25% | UK | 80.73% | 50 | 0.48 | 0.2 | 10.3 |
| 3. | C21614T | L18F | 10.51% | UK | 95.53% | 51 | 0.47 | 0.2 | 10.6 |
| 4. | G22992A | S477N | 6.1% | Australia | 52.67% | 45 | 0.47 | 0.4 | 11.6 |
| 5. | C24334T | A924A | 4.69% | UK | 98.89% | 48 | 0.44 | 0.5 | 10.1 |
| 6. | G23401A | Q613Q | 3.5% | Australia | 99.97% | 43 | 0.46 | 0.7 | 11.6 |
| 7. | C23604A | P681H | 3.31% | UK | 83.74% | 46 | 0.50 | 0.2 | 10.1 |
| 8. | A23063T | N501Y | 3.14% | UK | 93.12% | 44 | 0.45 | 0.2 | 9.3 |
| 9. | C23709T | T716I | 2.88% | UK | 95.99% | 50 | 0.47 | 0.2 | 9.9 |
| 10. | G24914C | D1118H | 2.84% | UK | 96.77% | 50 | 0.45 | 0.2 | 10.1 |
| 11. | C23271A | A570D | 2.84% | UK | 96.76% | 48 | 0.45 | 0.2 | 11.9 |
| 12. | T24506G | S982A | 2.83% | UK | 96.78% | 50 | 0.45 | 0.2 | 3.7 |
| 13. | C23731T | T723T | 2.39% | UK | 74.55% | 46 | 0.46 | 0.4 | 10.5 |
| 14. | C22879A | N439K | 2.13% | UK | 52.44% | 47 | 0.46 | 0.3 | 10.0 |
| 15. | C22444T | D294D | 1.84% | UK | 53.94% | 41 | 0.55 | 0.3 | 11.1 |
| 16. | T24910C | T1116T | 1.52% | UK | 46.17% | 50 | 0.41 | 0.4 | 9.3 |
| 17. | C21637T | P25P | 1.4% | UK | 97.42% | 51 | 0.46 | 0.5 | 10.0 |
| 18. | C21575T | L5F | 1.39% | UK | 53.21% | 46 | 0.50 | 0.4 | 11.2 |
| 19. | C22388T | L276L | 1.35% | UK | 92.29% | 48 | 0.44 | 0.2 | 10.2 |
| 20. | C21855T | S98F | 1.19% | Denmark | 25.75% | 45 | 0.55 | 0.4 | 11.6 |
| 21. | G22346T | A262S | 1.17% | UK | 80.56% | 48 | 0.46 | 0.5 | 10.7 |
| 22. | C22377T | P272L | 0.86% | UK | 72.69% | 49 | 0.45 | 0.4 | 9.8 |
| 23. | T24088C | G842G | 0.76% | UK | 99.31% | 48 | 0.31 | 0.7 | 6.4 |
| 24. | T24076C | G838G | 0.73% | USA | 97.54% | 44 | 0.54 | 0.4 | 9.8 |
| 25. | C23929T | Y789Y | 0.67% | Singapore | 45.75% | 40 | 0.76 | 0.6 | 10.6 |
| 26. | C22480T | F306F | 0.65% | Australia | 86.77% | 40 | 0.47 | 0.8 | 10.6 |
| 27. | G25049T | D1163Y | 0.63% | UK | 82.86% | 46 | 0.51 | 0.5 | 9.9 |
| 28. | G25062T | G1167V | 0.59% | UK | 89.47% | 44 | 0.55 | 0.6 | 10.0 |
| 29. | T22020C | M153T | 0.55% | Japan | 82.62% | 45 | 0.42 | 0.3 | 12.5 |
| 30. | G21624T | R21I | 0.55% | UK | 95.06% | 53 | 0.48 | 0.7 | 10.1 |
| 31. | T24814C | D1084D | 0.55% | Denmark | 99.28% | 66 | 0.6 | 0.7 | 10.3 |
| 32. | G21800T | D80Y | 0.54% | UK | 65.93% | 47 | 0.46 | 0.6 | 10.4 |
| 33. | G23311C | E583D | 0.54% | UK | 97.1% | 61 | 0.27 | 0.4 | 9.8 |
| 34. | C24034T | N824N | 0.42% | USA | 59.66% | 50 | 0.46 | 0.4 | 12.1 |
| 35. | G24368T | D936Y | 0.41% | UK | 39.82% | 51 | 0.53 | 0.5 | 10.5 |
| 36. | C23644T | A694A | 0.39% | UK | 89.07% | 47 | 0.47 | 0.5 | 9.1 |
| 37. | G22051T | A163A | 0.38% | UK | 96.83% | 52 | 0.58 | 0.7 | 10.5 |
| 38. | C23525T | H655Y | 0.37% | UK | 86.64% | 46 | 0.56 | 0.5 | 11.7 |
| 39. | C23248T | F562F | 0.36% | Denmark | 57.99% | 41 | 0.52 | 0.7 | 10.0 |
| 40. | C23625T | A688V | 0.35% | UK | 73.03% | 41 | 0.44 | 0.5 | 10.4 |
| 41. | C22088T | L176F | 0.35% | UK | 82.94% | 45 | 0.59 | 0.7 | 10.2 |
| 42. | G21724T | L54F | 0.34% | UK | 35.49% | 47 | 0.61 | 0.5 | 10.5 |
| 43. | G23593T | Q677H | 0.33% | USA | 33.14% | 46 | 0.52 | 0.3 | 10.4 |
| 44. | G24781T | K1073N | 0.33% | UK | 82.32% | 53 | 0.40 | 0.5 | 10.1 |
| 45. | A22920T | Y453F | 0.33% | Denmark | 96.12% | 33 | 0.77 | 0.9 | 8.8 |
| 46. | G22205C | D215H | 0.33% | UK | 88.55% | 53 | 0.48 | 0.4 | 11.4 |
| 47. | C23707T | P715P | 0.32% | USA | 33.77% | 49 | 0.45 | 0.5 | 10.6 |
| 48. | G25088T | V1176F | 0.32% | Brazil | 68.01% | 49 | 0.53 | 0.6 | 10.5 |
| 49. | A22320G | D253G | 0.31% | USA | 91.2% | 48 | 0.48 | 0.6 | 9.5 |
| 50. | A22255T | I231I | 0.3% | USA | 97.31% | 38 | 0.55 | 0.4 | 6.9 |
| **ORF3a** | | | | | | | | | |
| 51. | G25563T | Q57H | 21.41 | USA | 52.99% | 48 | 0.53 | 0.2 | 11.6 |
| 52. | C25710T | L106L | 2.75 | UK | 29.5% | 46 | 0.50 | 0.4 | 10.5 |
| 53. | G26144T | G251V | 2.02 | UK | 68.06% | 57 | 0.50 | 3.5 | 12.5 |
| 54. | G25907T | G172V | 1.88 | USA | 94.39% | 43 | 0.54 | 0.3 | 9.5 |
| 55. | C26060T | T223I | 1.66 | UK | 79.32% | 39 | 0.53 | 0.2 | 10.4 |
| 56. | G25996T | V202L | 1.12 | Denmark | 26.0% | 45 | 0.55 | 0.4 | 10.0 |
| 57. | A25505G | Q38R | 1.06 | Denmark | 27.25% | 45 | 0.55 | 0.4 | 9.8 |
| 58. | G25906C | G172R | 1.03 | Denmark | 28.2% | 45 | 0.55 | 0.4 | 9.8 |
| 59. | C25614T | S74S | 0.99 | UK | 94.65% | 49 | 0.55 | 0.5 | 10.5 |
| 60. | G25617T | K75N | 0.88 | UK | 97.79% | 56 | 0.39 | 0.6 | 10.5 |
| 61. | G25429T | V13L | 0.83 | UK | 78.41% | 53 | 0.55 | 0.7 | 10.5 |
| 62. | G25757T | R122I | 0.59 | Denmark | 92.22% | 40 | 0.3 | 0.7 | 9.5 |
| 63. | C25889T | S166L | 0.43 | UK | 58.82% | 49 | 0.44 | 0.4 | 10.6 |
| 64. | C25904T | S171L | 0.42 | UK | 43.46% | 42 | 0.49 | 0.2 | 11.4 |
| 65. | G25979T | G196V | 0.41 | Spain | 29.98% | 51 | 0.48 | 1.2 | 10.9 |
| 66. | G25494T | T34T | 0.41 | UK | 70.64% | 48 | 0.53 | 0.4 | 10.1 |
| 67. | G25552T | A54S | 0.40 | UK | 75.67% | 47 | 0.55 | 0.4 | 10.5 |
| 68. | G25606T | A72S | 0.37 | UK | 73.4% | 41 | 0.57 | 0.5 | 10.1 |
| 69. | G25720T | A110S | 0.36 | UK | 77.38% | 47 | 0.59 | 0.4 | 9.8 |
| 70. | A25881T | V163V | 0.35 | UK | 87.68% | 48 | 0.47 | 0.5 | 10.3 |
| 71. | T25878C | S162S | 0.35 | UK | 87.41% | 48 | 0.46 | 0.5 | 9.8 |
| 72. | G25879A | V163I | 0.35 | UK | 87.58% | 48 | 0.47 | 0.5 | 9.5 |
| 73. | G25437T | L15F | 0.34 | UK | 76.33% | 45 | 0.44 | 0.4 | 10.5 |
| 74. | C25936T | H182Y | 0.31 | Denmark | 88.92% | 44 | 0.59 | 0.9 | 10.2 |
| 75. | G25947T | Q185H | 0.31 | UK | 71.25% | 56 | 0.39 | 0.5 | 11.5 |
| **E** | | | | | | | | | |
| 76. | T26424C | S60S | 1.231391 | UK | 97.96% | 62 | 0.44 | 0.5 | 6.9 |
| 77. | C26313T | F23F | 0.394720 | UK | 65.27% | 48 | 0.48 | 0.5 | 10.0 |
| **M** | | | | | | | | | |
| 78. | C26801G | L93L | 21.82 | UK | 81.5% | 49 | 0.48 | 0.2 | 10.3 |
| 79. | C26735T | Y71Y | 5.07 | UK | 38.98% | 44 | 0.55 | 0.3 | 11.4 |
| 80. | T26876C | I118I | 2.58 | UK | 30.37% | 45 | 0.50 | 0.4 | 10.1 |
| 81. | T26972C | R150R | 1.56 | UK | 47.11% | 50 | 0.45 | 0.4 | 9.5 |
| 82. | C26681T | F53F | 0.5 | UK | 46.49% | 48 | 0.51 | 0.3 | 11.4 |
| 83. | C26801T | L93L | 0.44 | Denmark | 46.95% | 41 | 0.52 | 0.4 | 10.5 |
| 84. | A26530G | D3G | 0.43 | Denmark | 26.37% | 50 | 0.51 | 0.7 | 11.5 |
| 85. | C27046T | T175M | 0.43 | UK | 42.06% | 50 | 0.59 | 0.7 | 10.7 |
| 86. | C27059T | Y179Y | 0.34 | Australia | 68.86% | 52 | 0.41 | 0.6 | 10.5 |
| 87. | C27002T | D160D | 0.33 | Denmark | 55.41% | 40 | 0.58 | 0.7 | 10.7 |
| 88. | C26645T | N41N | 0.30 | UK | 74.39% | 49 | 0.39 | 0.4 | 9.9 |
| **ORF6** | | | | | | | | | |
| 89. | T27384C | D61D | 0.61 | UK | 48.11% | 47 | 0.43 | 0.4 | 12.2 |
| 90. | T27299C | I33T | 0.36 | Brazil | 62.2% | 46 | 0.47 | 0.7 | 10.5 |
| **ORF7a** | | | | | | | | | |
| 91. | C27434T | T14I | 0.59% | UK | 37.3% | 45 | 0.45 | 0.5 | 10.5 |
| 92. | C27513T | Y40Y | 0.51% | UK | 76.4% | 47 | 0.42 | 0.4 | 10.2 |
| **ORF7b** | | | | | | | | | |
| 93. | C27800A | A15A | 1.65 | Denmark | 44.93% | 49 | 0.43 | 0.4 | 10.0 |
| 94. | C27769T | S5L | 1.14 | UK | 97.82% | 53 | 0.44 | 0.5 | 10.0 |
| 95. | A27865T | H37L | 0.38 | UK | 95.7% | 53 | 0.51 | 0.8 | 9.9 |
| 96. | T27866A | H37Q | 0.38 | UK | 96.2% | 56 | 0.56 | 0.8 | 9.4 |
| **ORF8** | | | | | | | | | |
| 97. | C27944T | H17H | 15.92 | UK | 84.51% | 50 | 0.5 | 0.2 | 12.3 |
| 98. | C27964T | S24L | 4.08 | USA | 94.7% | 48 | 0.5 | 0.3 | 10.5 |
| 99. | C27972T | Q27@ | 2.82 | UK | 94.69% | 51 | 0.46 | 0.2 | 10.1 |
| 100. | G28048T | R52I | 2.79 | UK | 95.34% | 50 | 0.46 | 0.2 | 10.5 |
| 101. | A28111G | Y73C | 2.76 | UK | 95.99% | 50 | 0.46 | 0.2 | 3.7 |
| 102. | T28144C | L84S | 2.68 | USA | 39.66% | 51 | 0.52 | 0.3 | 12.6 |
| 103. | A28169G | E92E | 1.35 | UK | 98.84% | 49 | 0.46 | 0.5 | 9.8 |
| 104. | C28087T | A65V | 1.0 | UK | 64.46% | 50 | 0.46 | 0.3 | 10.1 |
| 105. | C28253T | F120F | 0.67 | UK | 27.69% | 41 | 0.46 | 0.2 | 12.5 |
| 106. | A28133T | T80T | 0.51% | Denmark | 37.55% | 43 | 0.59 | 0.4 | 9.9 |
| 107. | A28095T | K68@ | 0.38% | UK | 94.87% | 45 | 0.29 | 0.2 | 10.0 |
| 108. | G28077T | V62L | 0.37% | UK | 43.8% | 43 | 0.47 | 0.3 | 11.3 |
| 109. | G28001T | P36P | 0.34% | UK | 79.44% | 48 | 0.58 | 0.5 | 10.2 |
| 110. | G28077C | V62L | 0.31% | USA | 77.83% | 51 | 0.49 | 0.7 | 12.0 |
| **N** | | | | | | | | | |
| **#** | **Genomic**  **change** | **Amino**  **acid**  **change** | **Frequency**  **%** | **Most records**  **containing SNP are uploaded by** | **% for**  **the**  **country*** | **SNP**  **age** | **SNP**  **gender** | **Uploaded months**  **ago, min** | **Uploaded months**  **ago, max** |
| 111. | G28881A | R203K | 35.551445 | UK | 49.37% | 46 | 0.49 | 0.2 | 11.6 |
| 112. | G28882A | R203R | 35.460712 | UK | 49.44% | 46 | 0.49 | 0.2 | 11.6 |
| 113. | G28883С | R203R | 35.458174 | UK | 60.31% | 46 | 0.50 | 0.3 | 10.2 |
| 114. | C28932T | A220V | 22.0% | UK | 80.45% | 49 | 0.49 | 0.2 | 10.3 |
| 115. | C28854T | S194L | 5.88% | USA | 36.69% | 42 | 0.53 | 0.3 | 12.5 |
| 116. | C28977T | S235F | 2.99% | UK | 88.94% | 50 | 0.45 | 0.2 | 9.8 |
| 117. | C28869T | P199L | 2.95% | USA | 59.56% | 44 | 0.53 | 0.3 | 10.0 |
| 118. | T28282A | D3E | 2.78% | UK | 95.96% | 50 | 0.44 | 0.2 | 9.9 |
| 119. | A28281T | D3V | 2.78% | UK | 95.96% | 50 | 0.44 | 0.2 | 9.9 |
| 120. | G28280C | D3H | 2.78% | UK | 95.95% | 50 | 0.44 | 0.2 | 9.9 |
| 121. | G28975C | M234I | 2.62% | Denmark | 29.7% | 46 | 0.50 | 0.4 | 9.7 |
| 122. | G29399A | A376T | 2.61% | Denmark | 29.47% | 46 | 0.50 | 0.4 | 9.7 |
| 123. | C28472T | P67S | 1.82% | USA | 93.11% | 43 | 0.53 | 0.3 | 10.5 |
| 124. | C29366T | P365S | 1.59% | UK | 57.72% | 55 | 0.43 | 0.4 | 10.8 |
| 125. | C28725T | P151L | 1.4% | Japan | 94.53% | 41 | 0.49 | 0.7 | 10.2 |
| 126. | C29466T | A398V | 1.37% | UK | 93.39% | 47 | 0.45 | 0.4 | 10.2 |
| 127. | G29227T | S318S | 1.36% | UK | 94.43% | 46 | 0.56 | 0.6 | 10.9 |
| 128. | C29386T | D371D | 1.31% | Denmark | 52.43% | 50 | 0.46 | 0.4 | 9.7 |
| 129. | G29402T | D377Y | 1.31% | USA | 37.74% | 47 | 0.48 | 0.3 | 10.3 |
| 130. | C28657T | D128D | 1.28% | UK | 50.99% | 55 | 0.49 | 0.6 | 10.7 |
| 131. | C28651T | N126N | 1.15% | Denmark | 25.06% | 45 | 0.55 | 0.4 | 10.1 |
| 132. | C29095T | F274F | 0.91% | UK | 46.8% | 47 | 0.53 | 0.3 | 12.2 |
| 133. | C28887T | T205I | 0.9% | USA | 44.29% | 42 | 0.49 | 0.2 | 11.6 |
| 134. | C28311T | P13L | 0.88% | Singapore | 34.07% | 40 | 0.74 | 0.3 | 10.6 |
| 135. | G28975T | M234I | 0.84% | Japan | 48.68% | 42 | 0.52 | 0.3 | 10.5 |
| 136. | G29179T | P302P | 0.81% | UK | 49.65% | 49 | 0.50 | 0.5 | 10.7 |
| 137. | G28580T | D103Y | 0.75% | UK | 91.34% | 52 | 0.47 | 0.7 | 10.5 |
| 138. | C28863T | S197L | 0.62% | Spain | 51.68% | 56 | 0.49 | 0.4 | 10.7 |
| 139. | C28821A | S183Y | 0.6% | USA | 88.87% | 41 | 0.44 | 0.6 | 10.1 |
| 140. | G29427A | R385K | 0.49% | UK | 69.9% | 42 | 0.48 | 0.6 | 10.2 |
| 141. | C28706T | H145Y | 0.44% | UK | 80.28% | 46 | 0.44 | 0.5 | 11.5 |
| 142. | G28878A | S202N | 0.41% | United Arab  Emirates | 20.28% | 40 | 0.65 | 0.3 | 11.8 |
| 143. | G28851T | S193I | 0.38% | UK | 62.91% | 55 | 0.53 | 0.4 | 11.8 |
| 144. | T29148C | I292T | 0.37% | Brazil | 59.25% | 45 | 0.48 | 0.7 | 10.6 |
| 145. | G28842T | S190I | 0.37% | USA | 73.73% | 49 | 0.50 | 0.7 | 10.5 |
| 146. | T28759C | P162P | 0.37% | UK | 53.0% | 48 | 0.51 | 0.4 | 10.2 |
| 147. | G28378T | A35A | 0.36% | UK | 37.7% | 46 | 0.56 | 0.4 | 10.9 |
| 148. | C28836T | S188L | 0.33% | UK | 89.69% | 56 | 0.37 | 1.4 | 10.5 |
| 149. | G28300T | Q9H | 0.32% | UK | 63.25% | 51 | 0.55 | 0.4 | 10.5 |
| 150. | G29044A | K257K | 0.32% | UK | 64.57% | 42 | 0.5 | 0.7 | 9.8 |
| 151. | C28310A | P13T | 0.31% | UK | 95.07% | 55 | 0.55 | 0.5 | 10.7 |
| **ORF10** | | | | | | | | | |
| 152. | G29645T | V30L | 22.03 | UK | 80.85% | 50 | 0.48 | 0.2 | 10.3 |
| 153. | C29614T | C19C | 0.31 | UK | 39.88% | 49 | 0.47 | 0.4 | 10.2 |

***of all records containing the SNP, the country from the previous column uploaded this percent of the records
