## Supplementary figures and images for "Analysis of 329,942 SARS-CoV-2 records retrieved from GISAID database"

### output_19_7.png

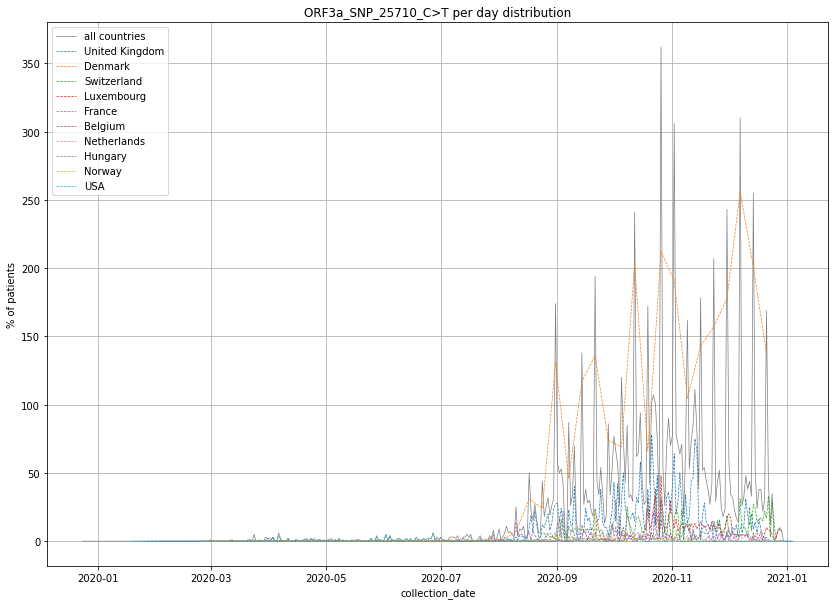

### output_19_19.png

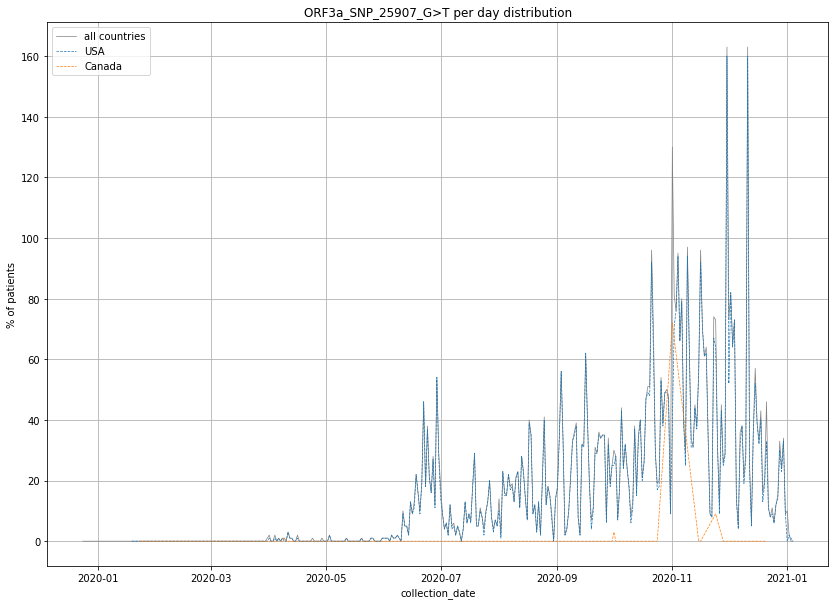

### output_19_32.png

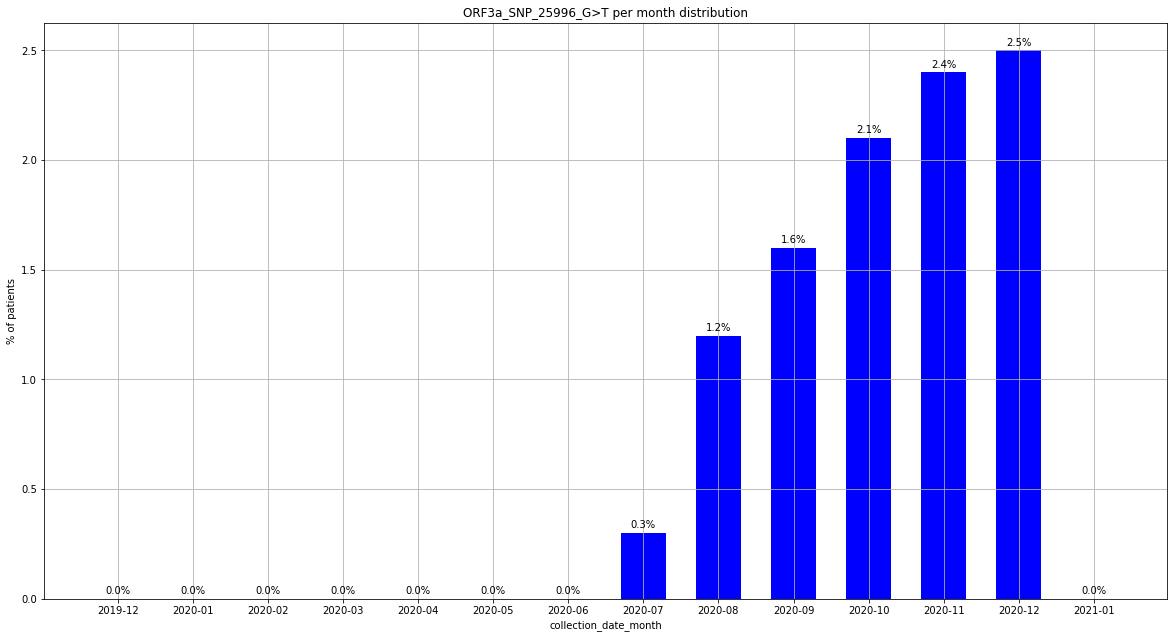

### output_19_67.png

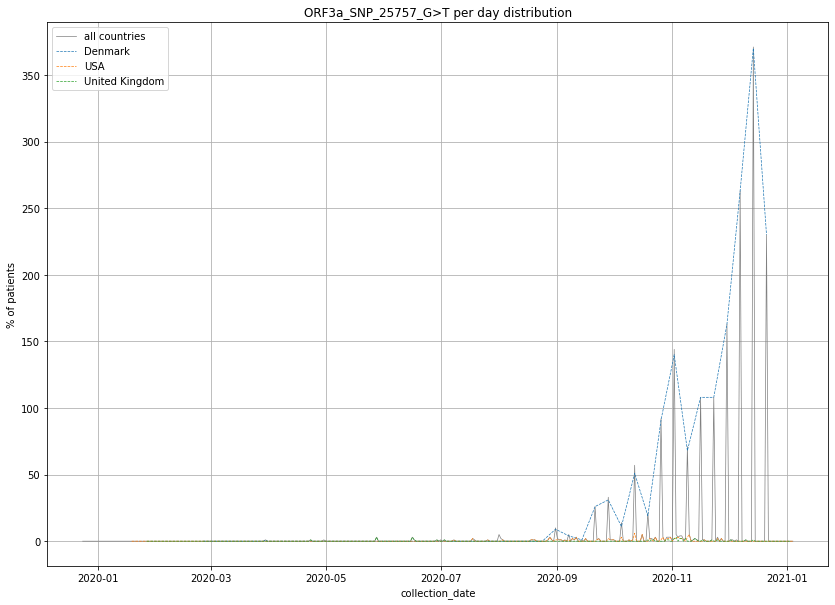

### output_19_85.png

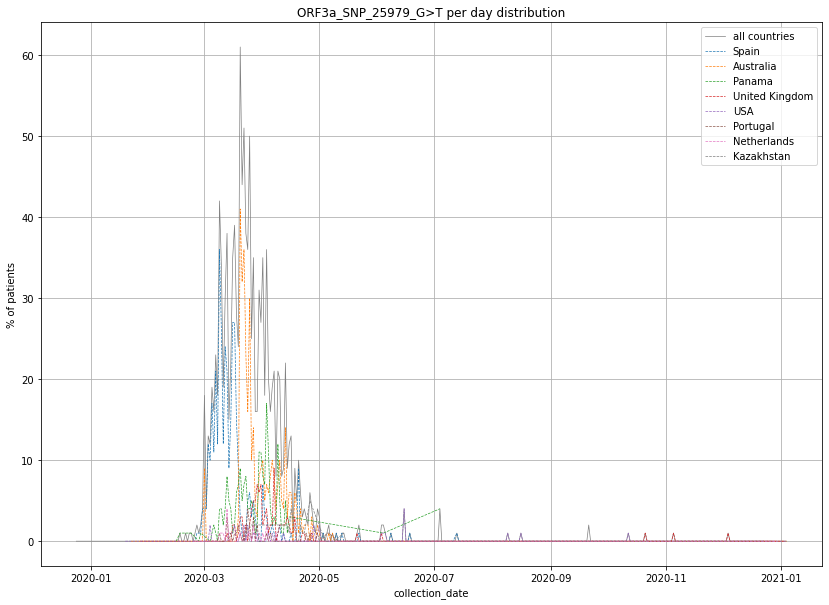

### output_19_92.png

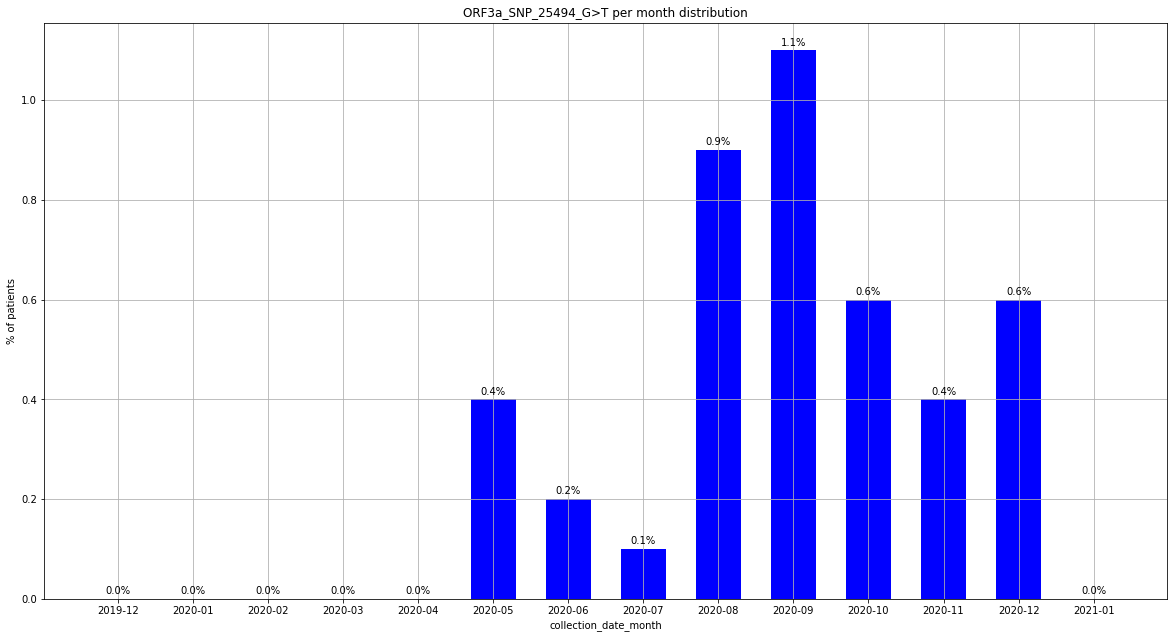

### output_19_104.png

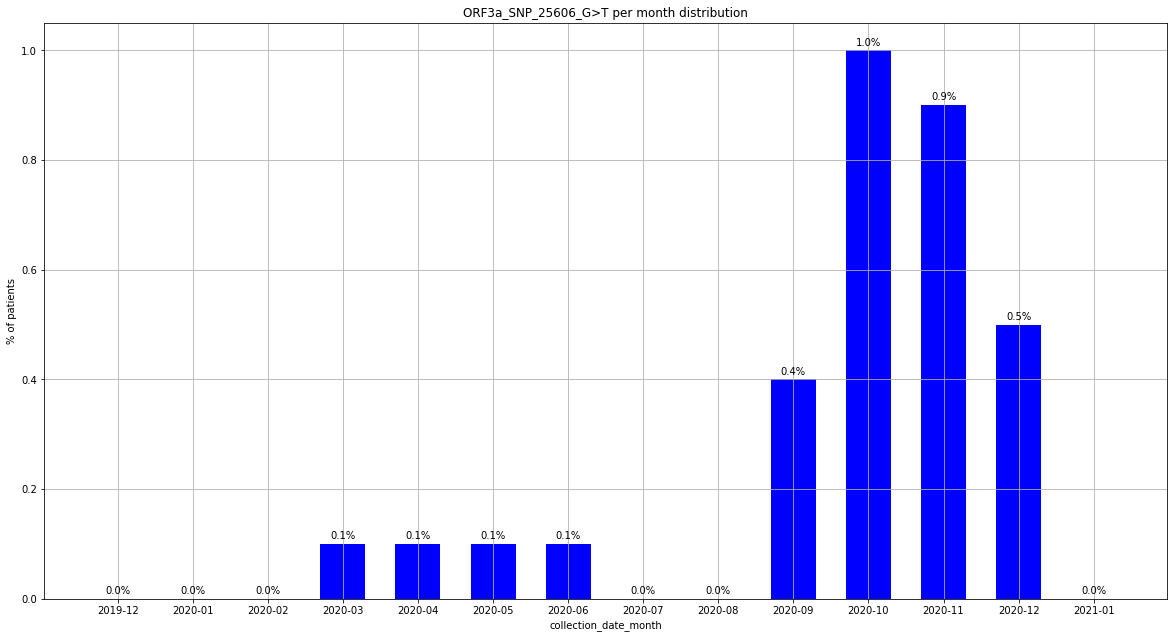

### output_19_115.png

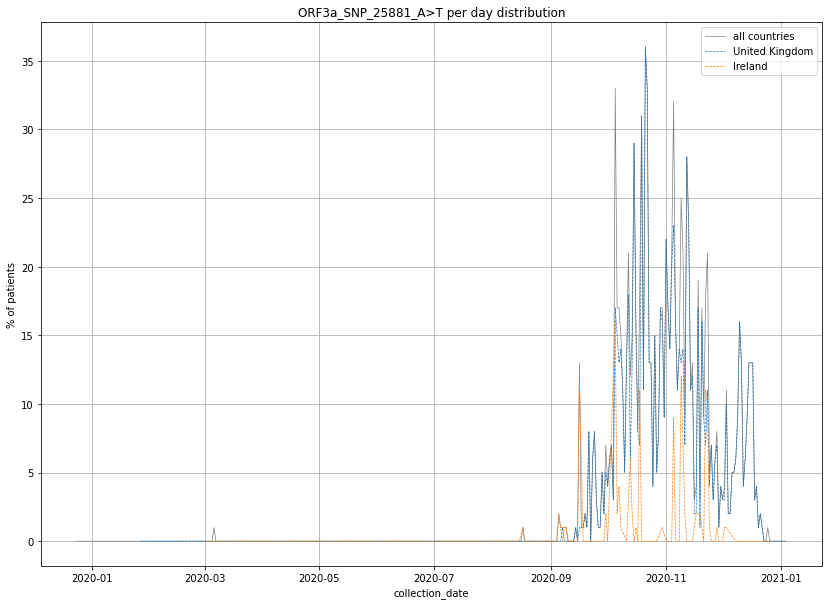

### output_19_116.png

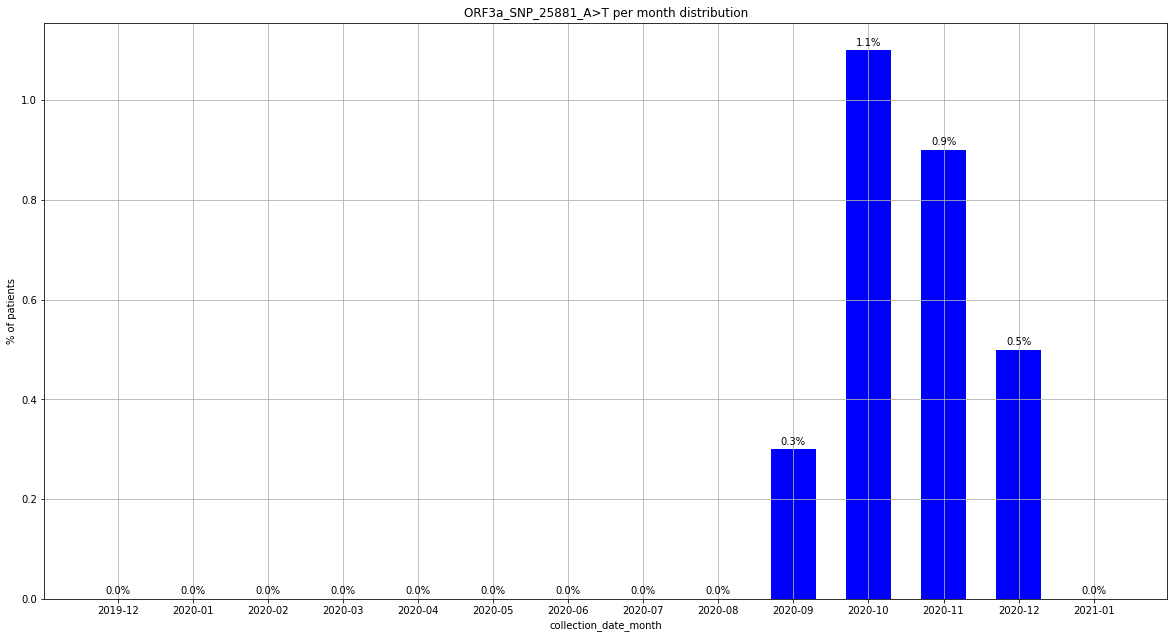

### output_19_145.png

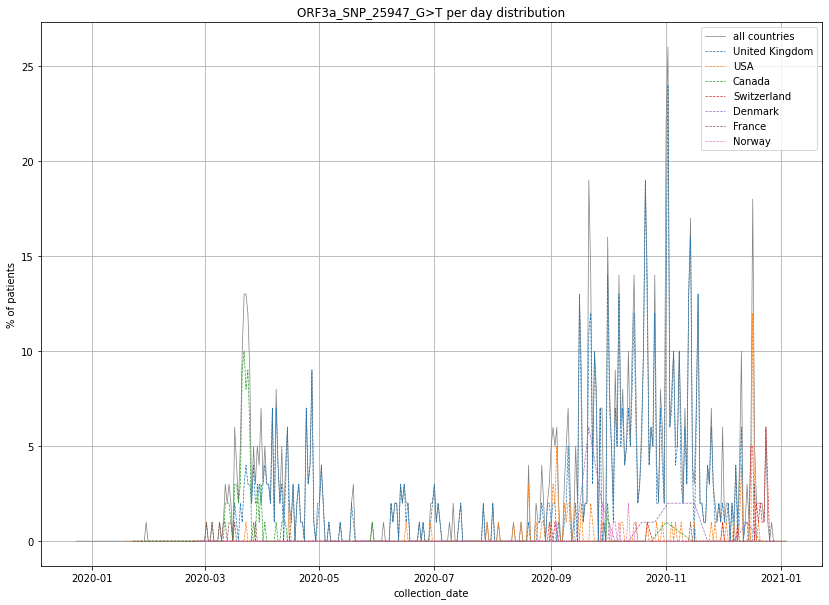

### output_20_2.png

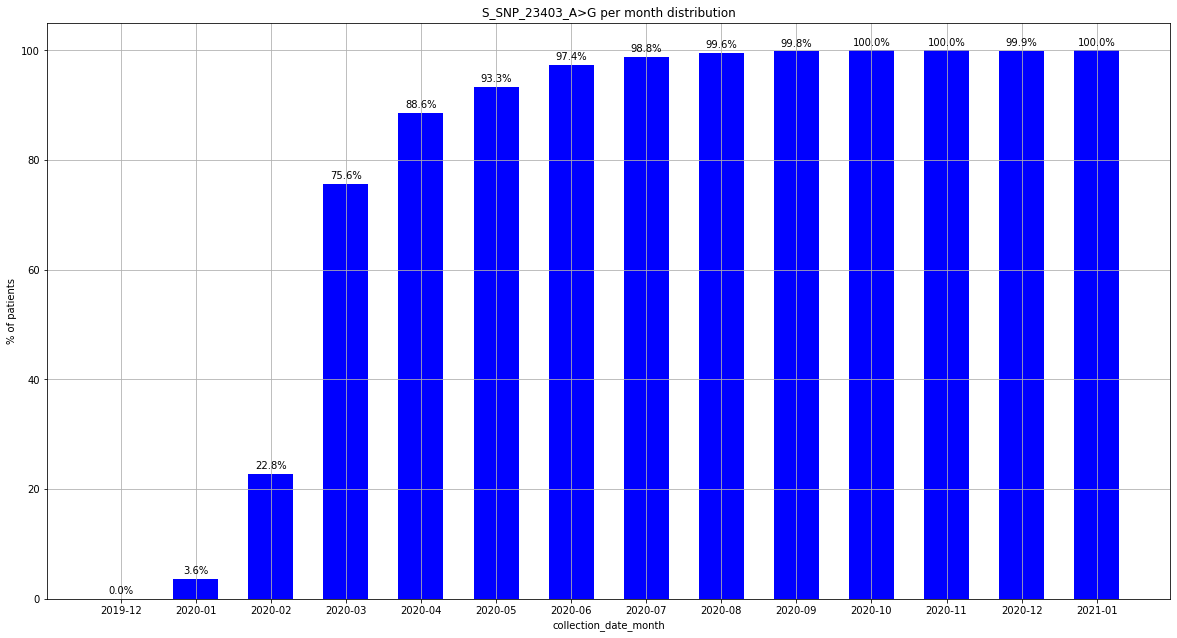

### output_20_7.png

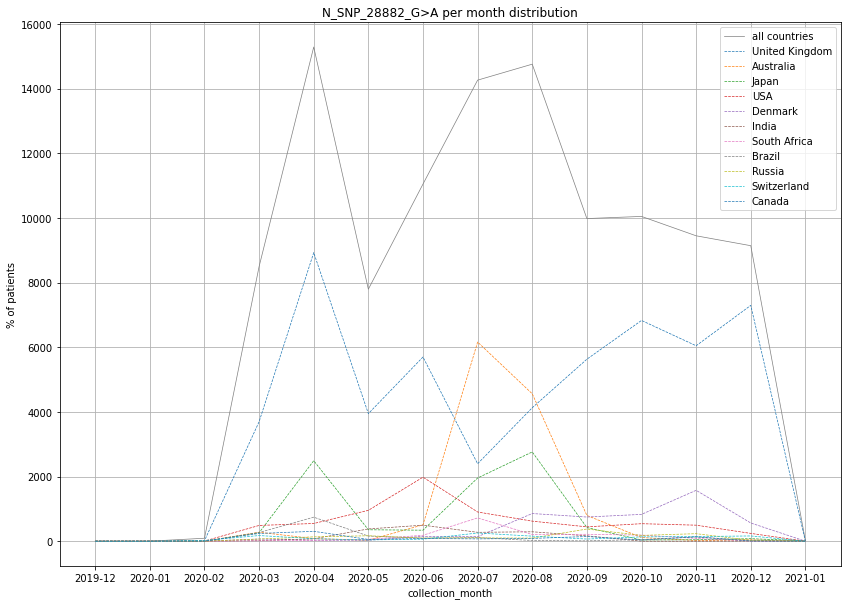

### output_20_7.png

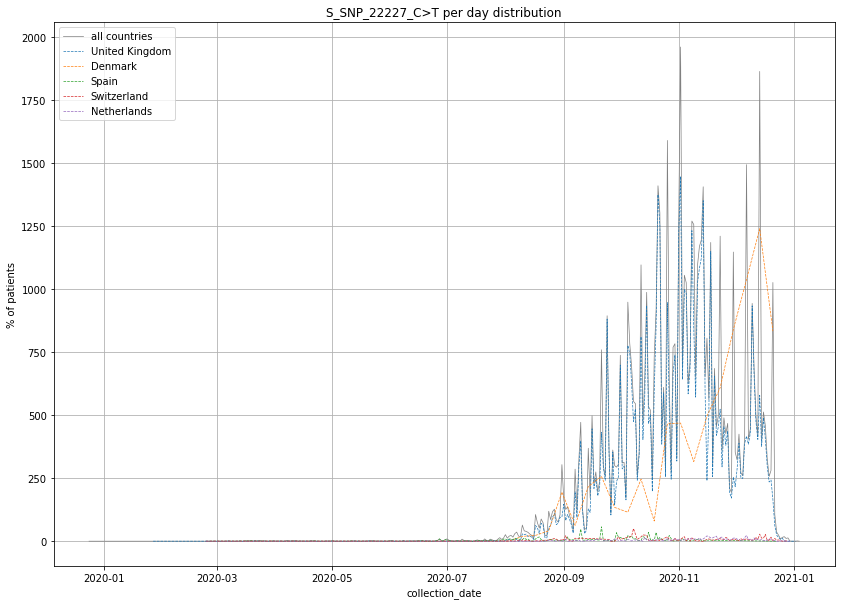

### output_20_8.png

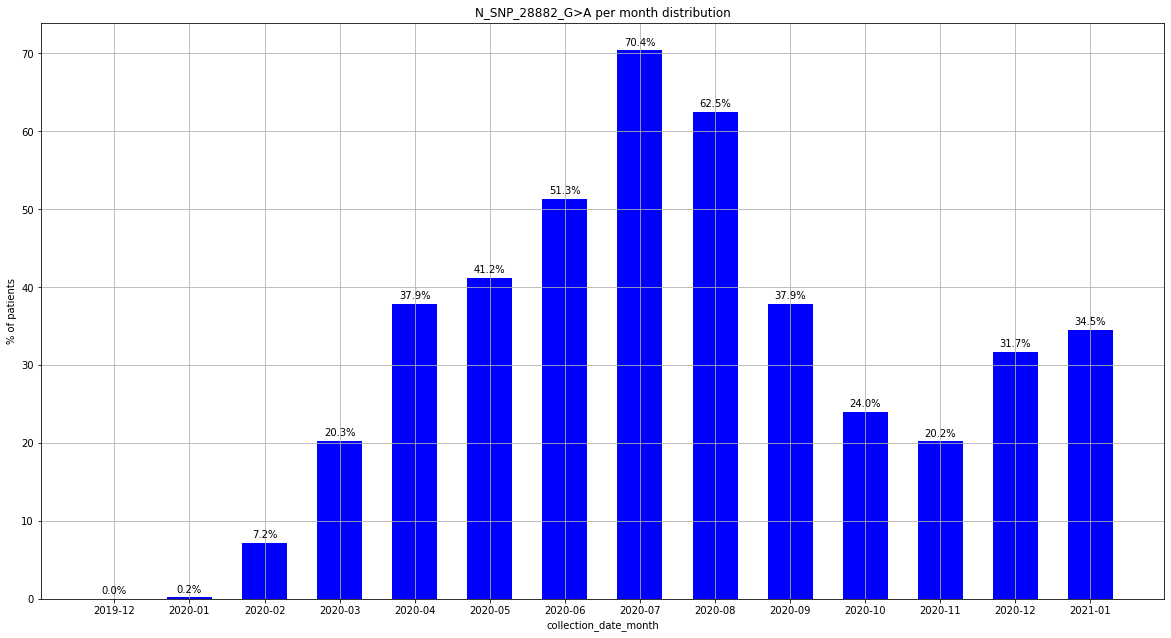

### output_20_13.png

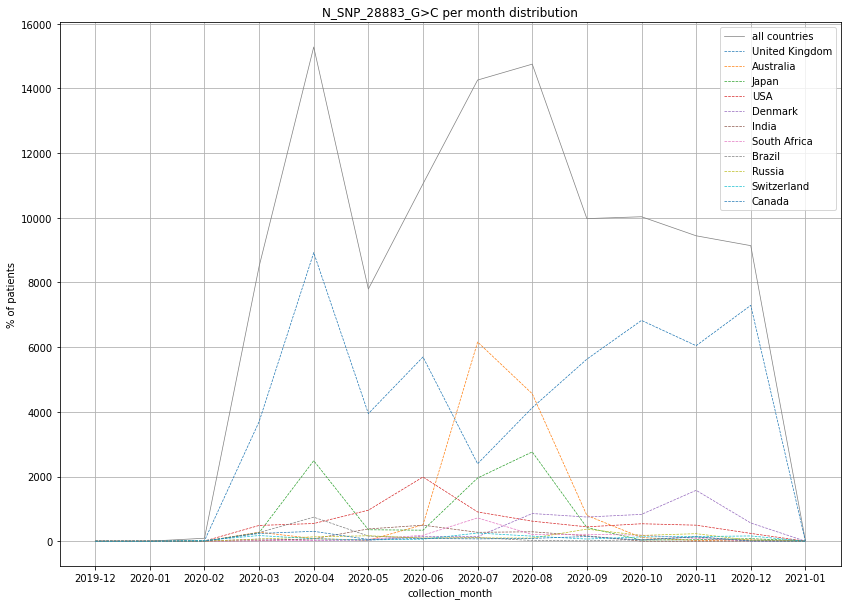

### output_20_14.png

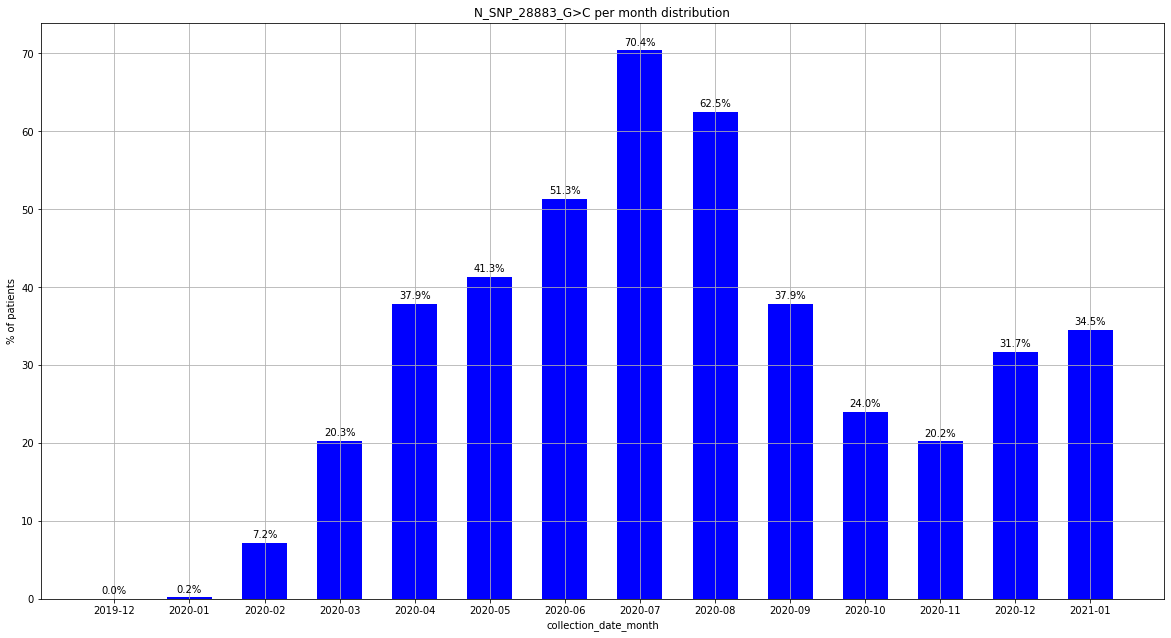

### output_20_14.png

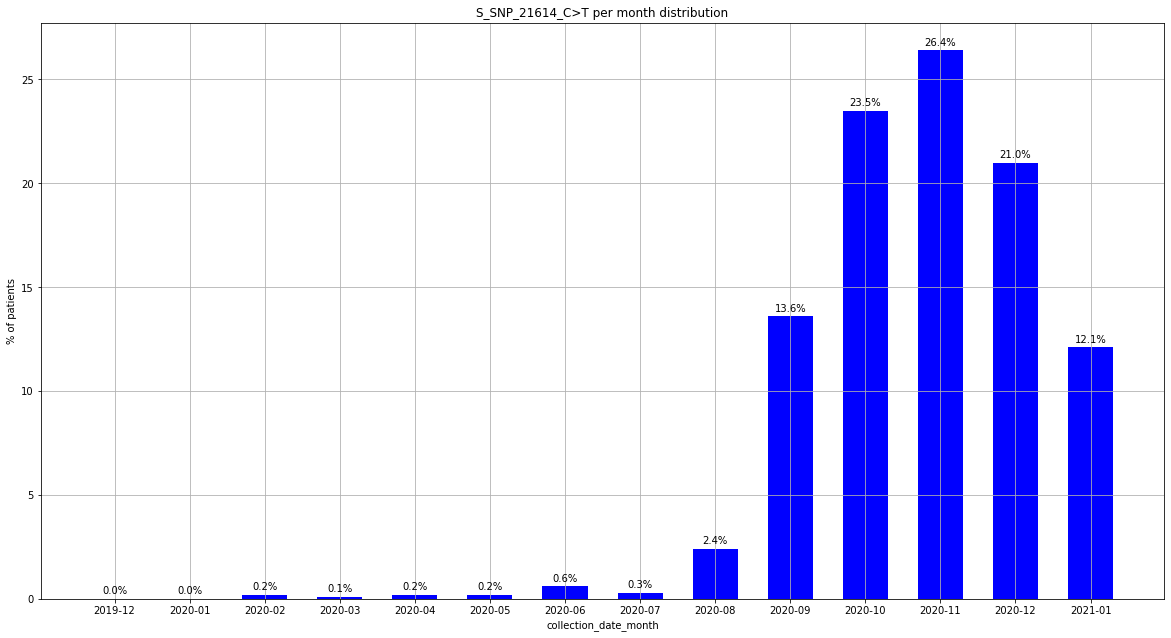

### output_20_19.png

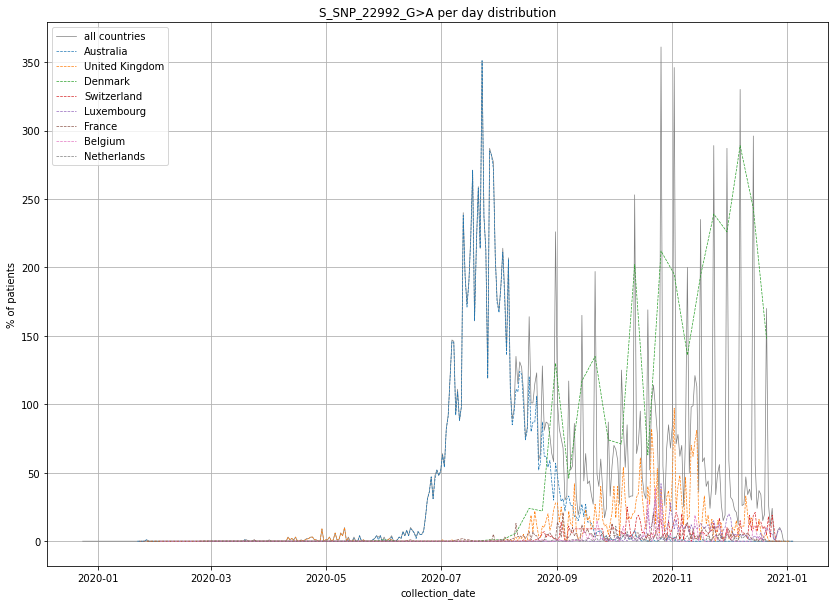

### output_20_19.png

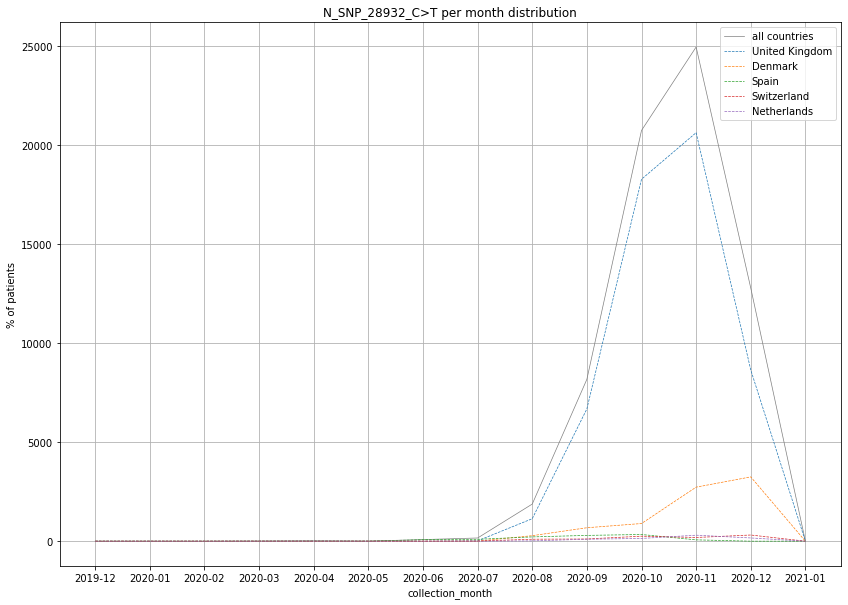

### output_20_20.png

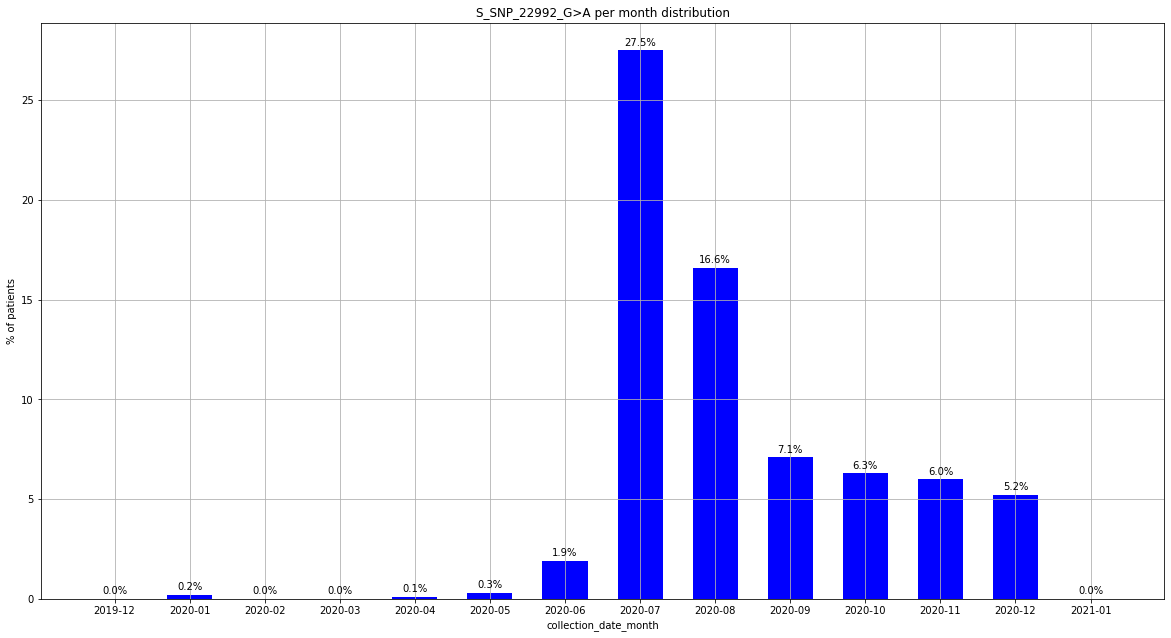

### output_20_20.png

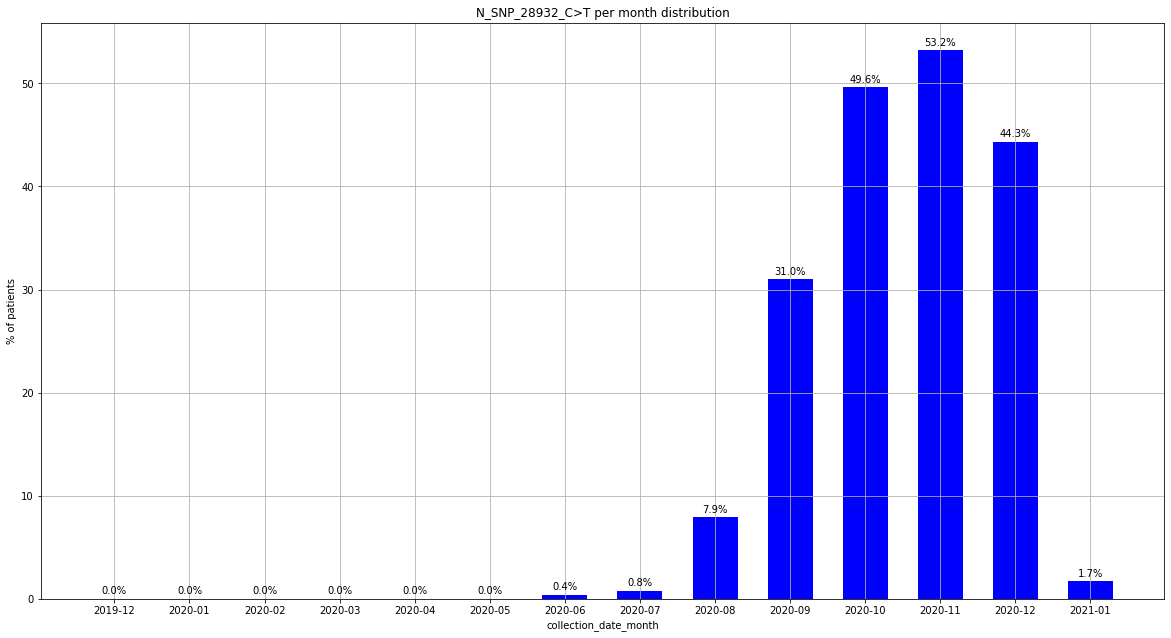

### output_20_25.png

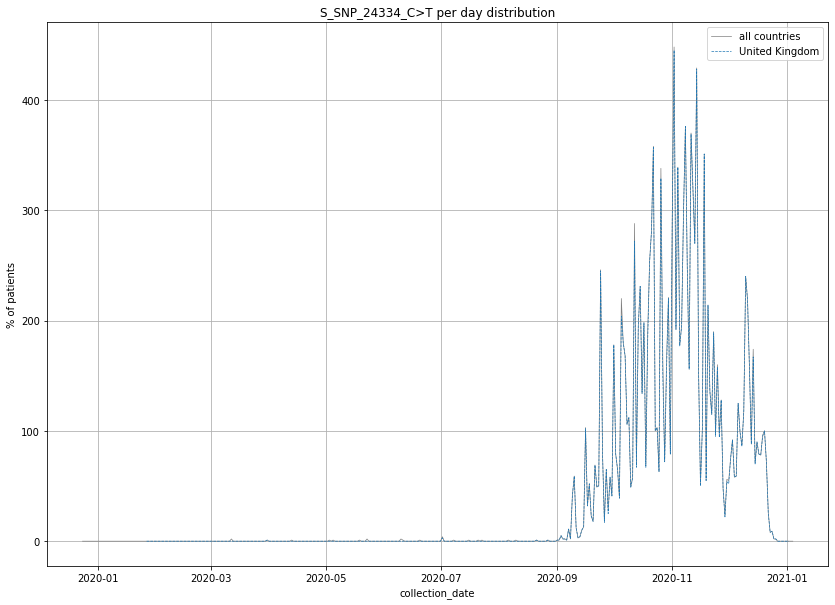

### output_20_25.png

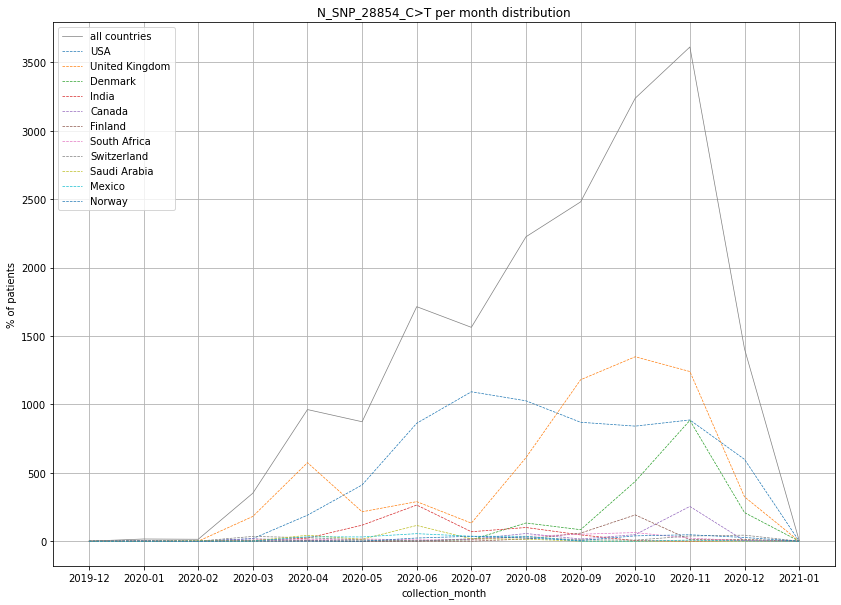

### output_20_26.png

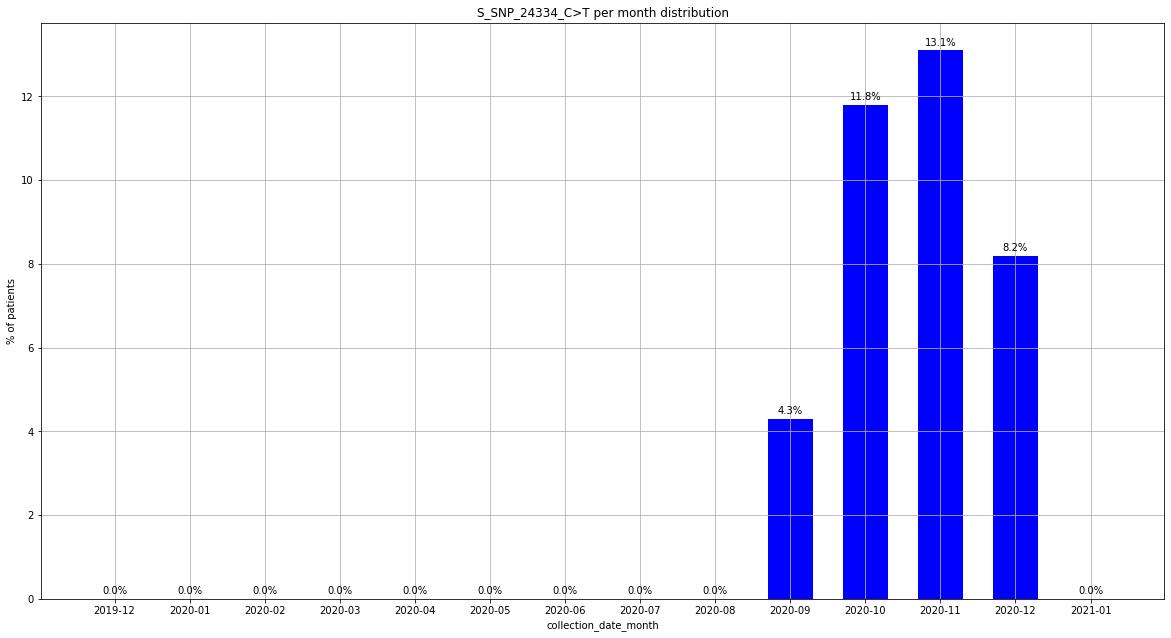

### output_20_32.png

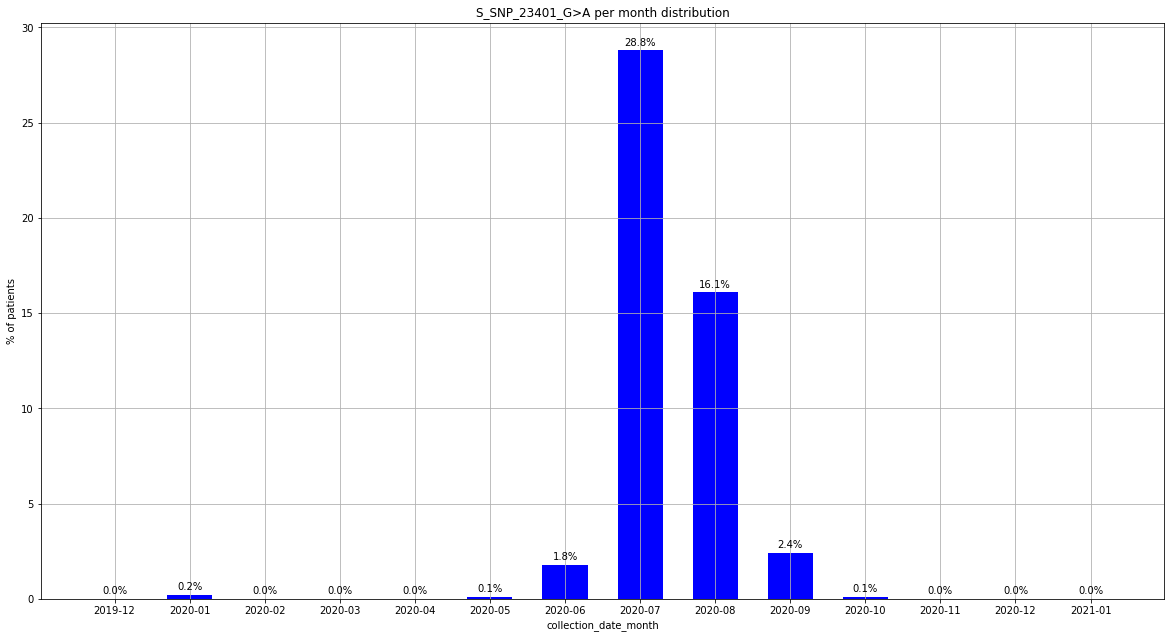

### output_20_37.png

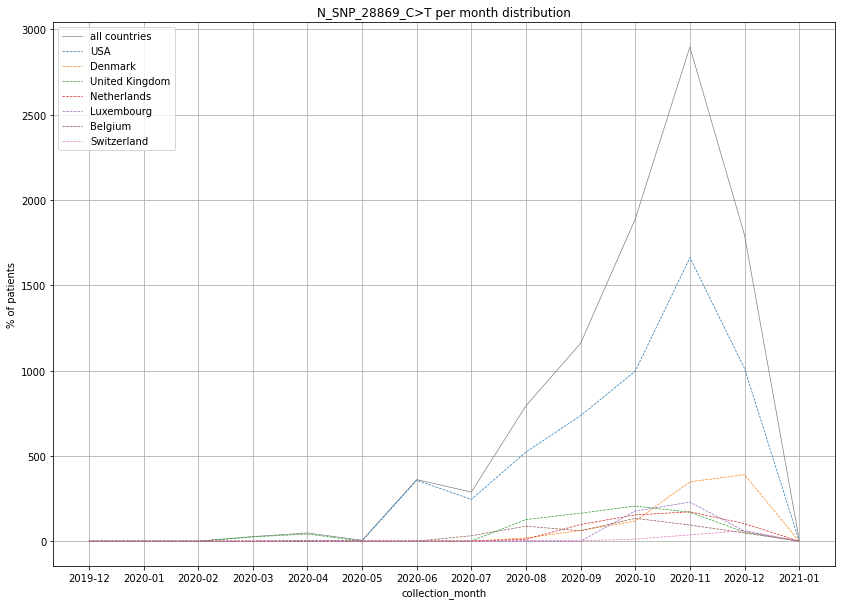

### output_20_37.png

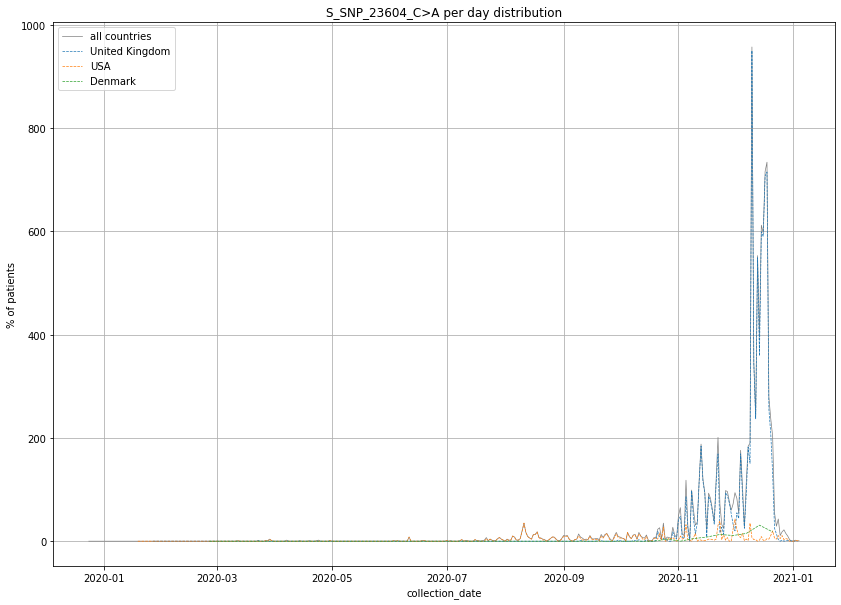

### output_20_43.png

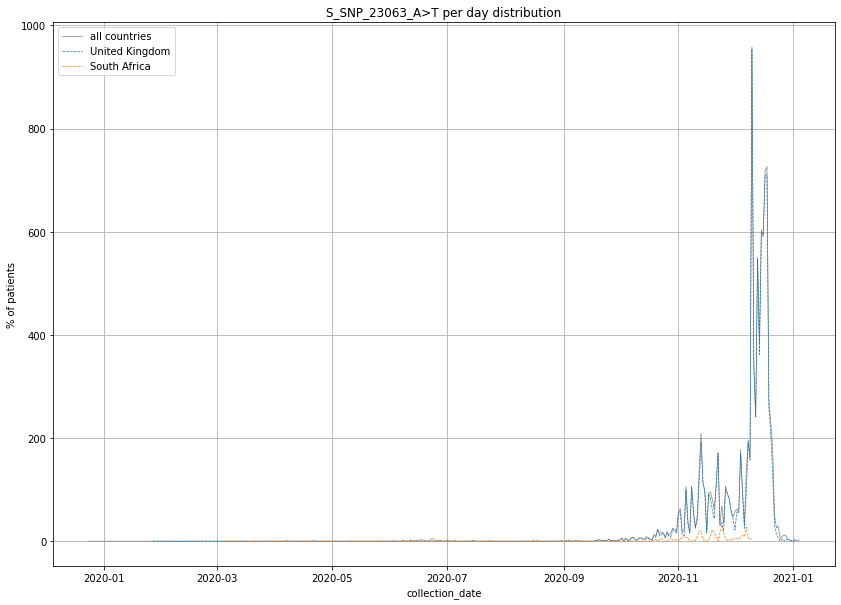

### output_20_43.png

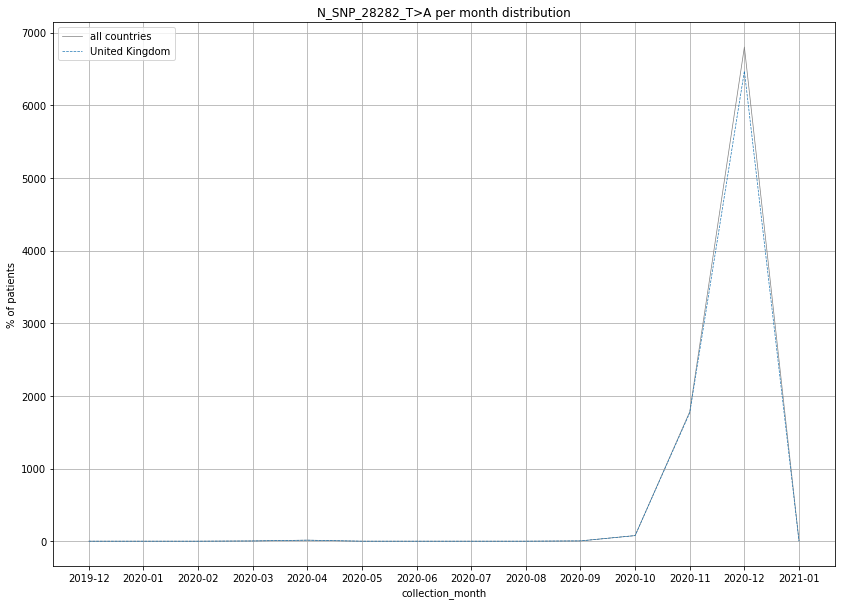

### output_20_50.png

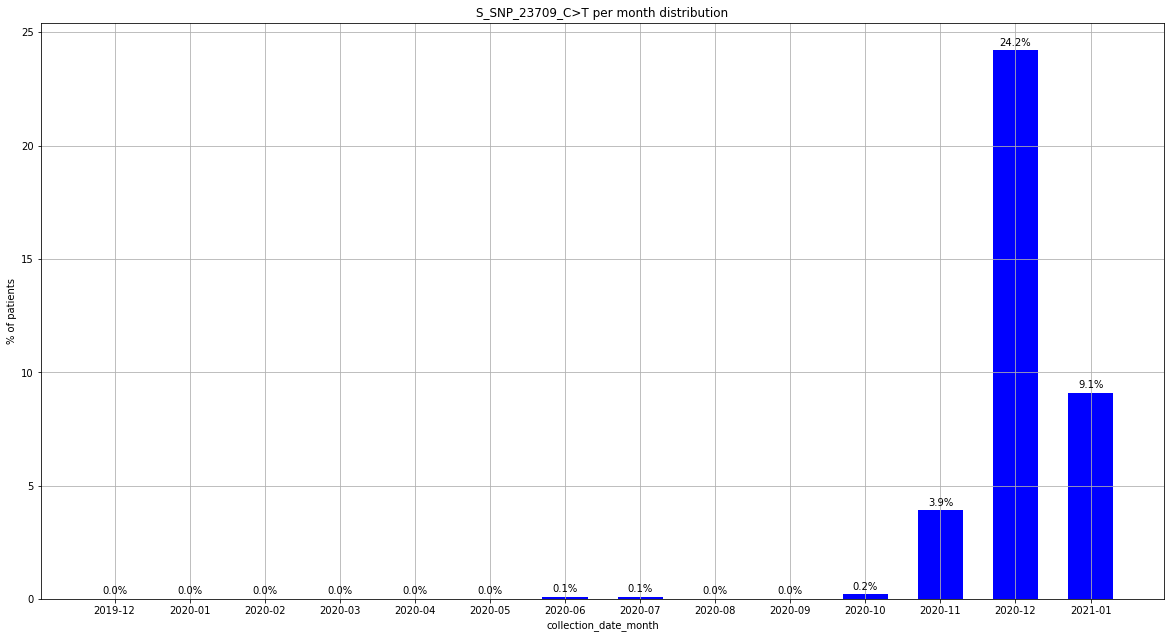

### output_20_56.png

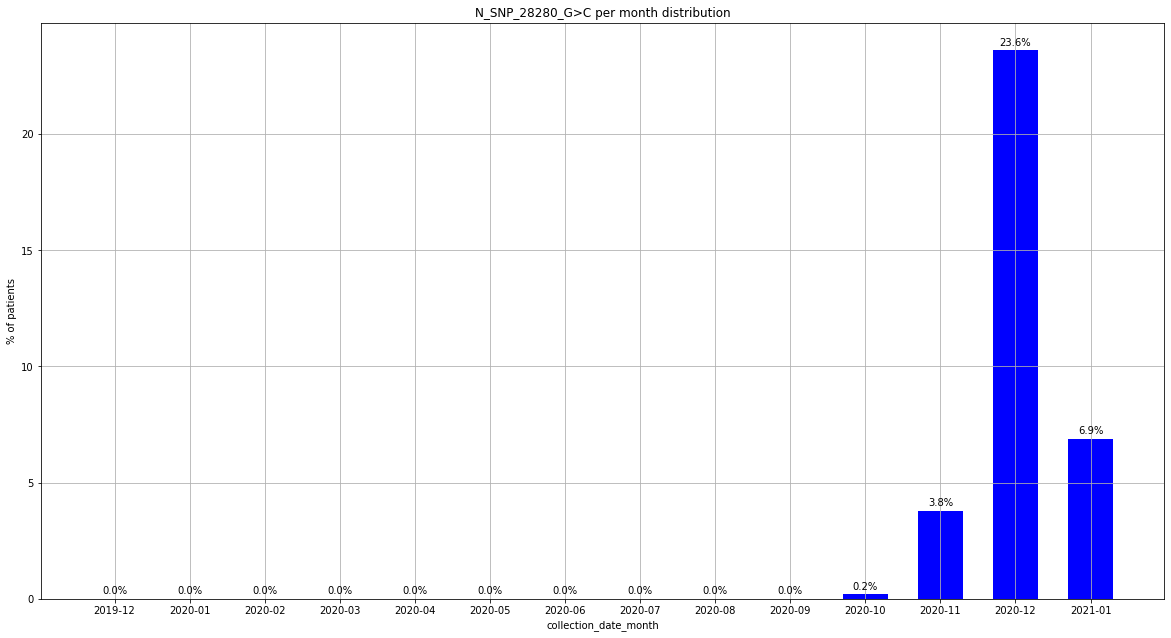

### output_20_61.png

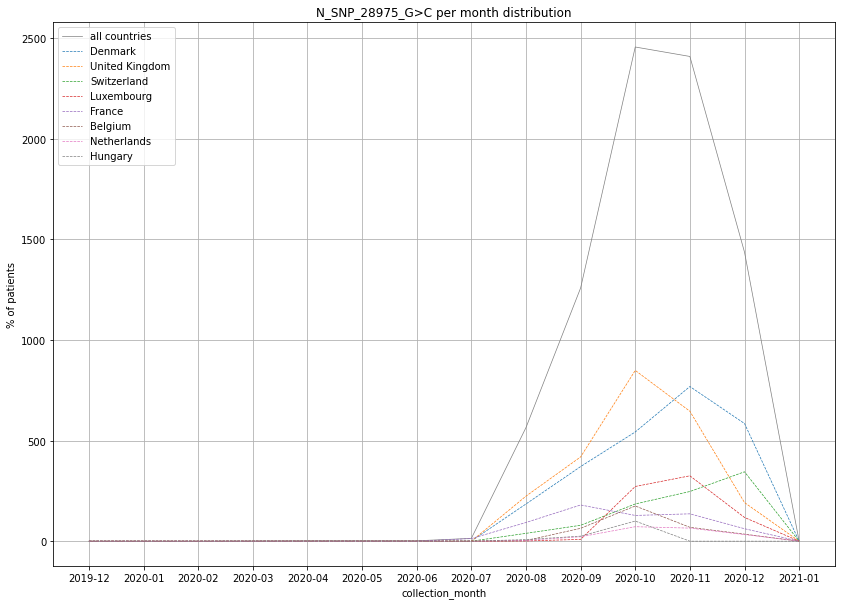

### output_20_61.png

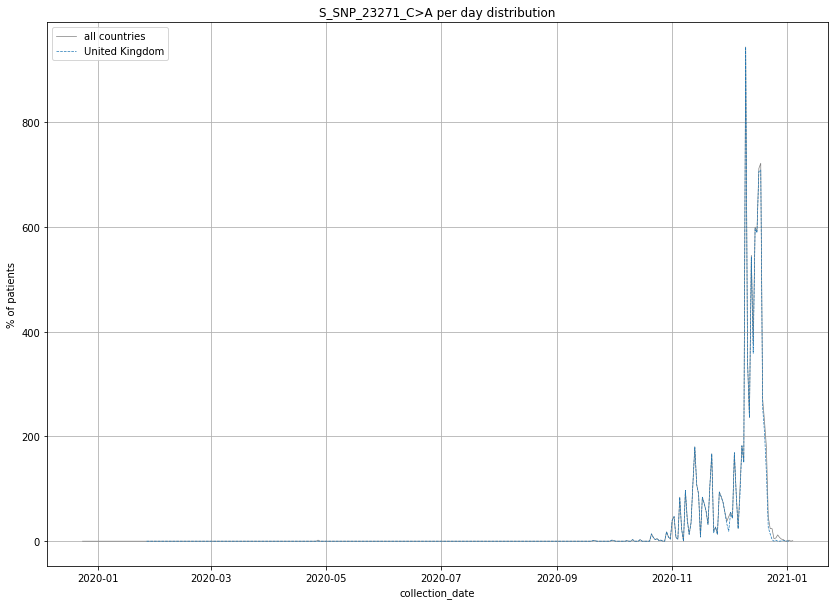

### output_20_73.png

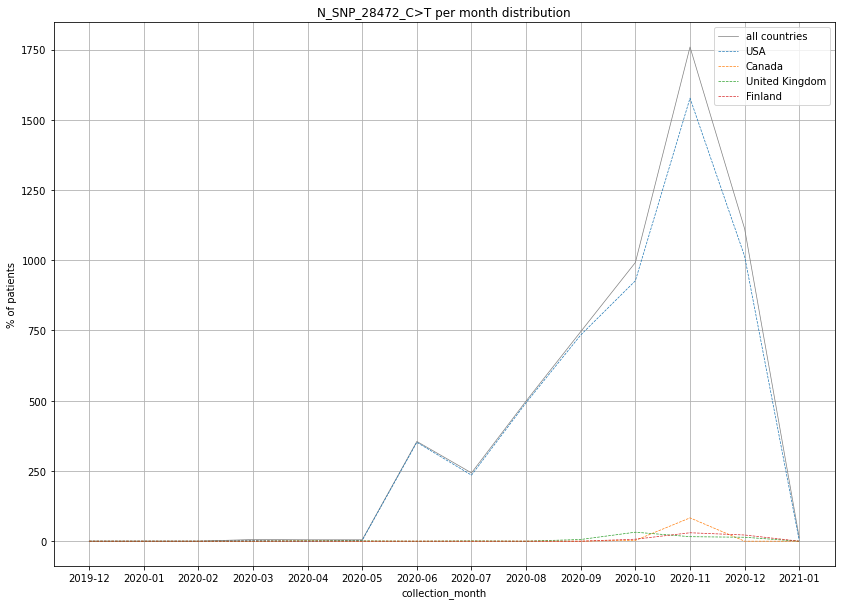

### output_20_74.png

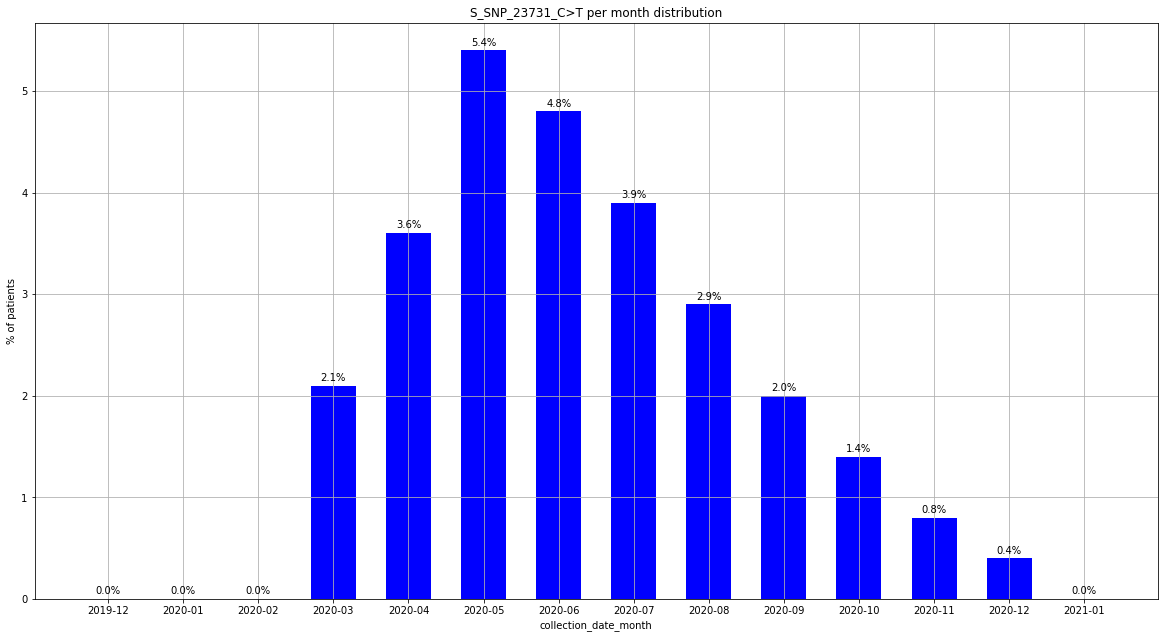

### output_20_79.png

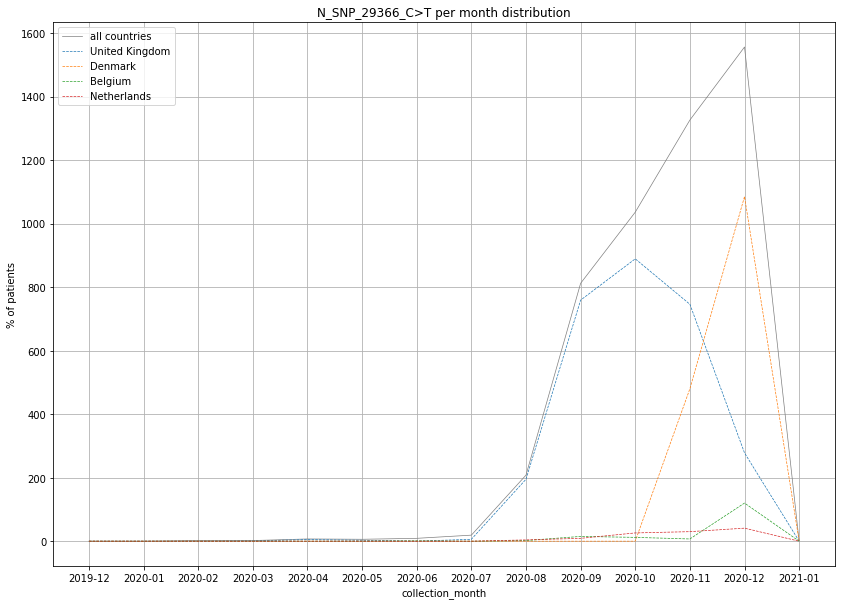

### output_20_79.png

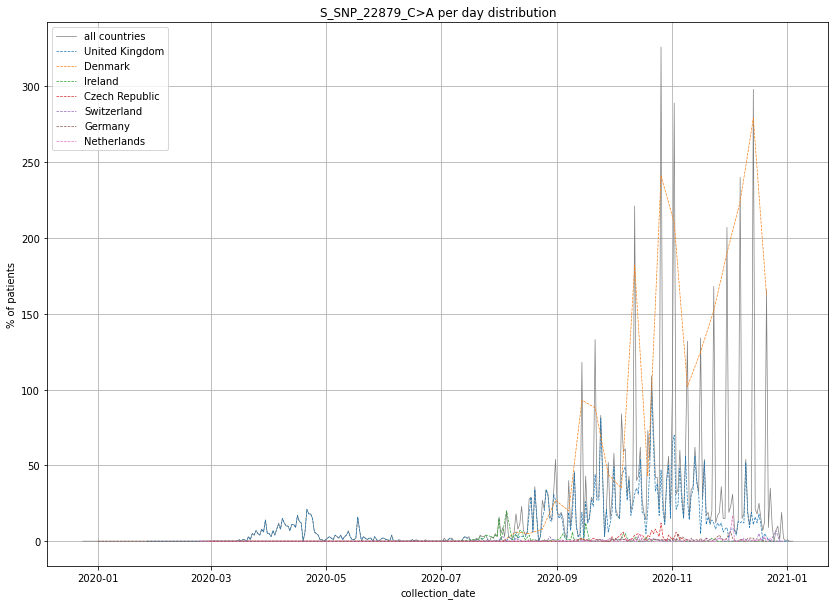

### output_20_86.png

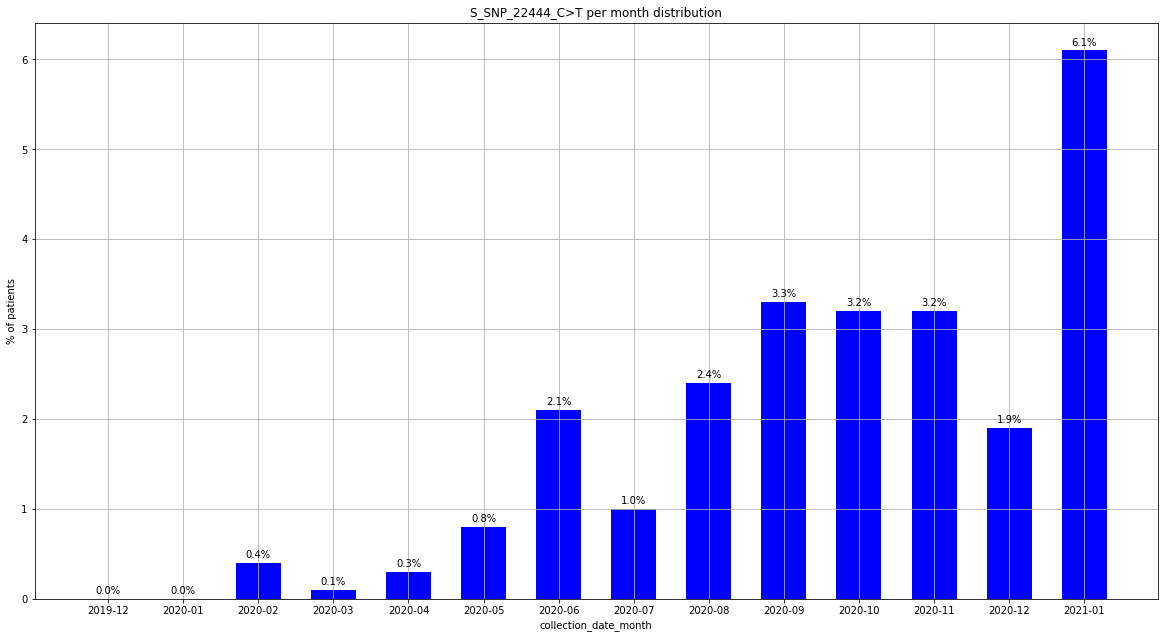

### output_20_91.png

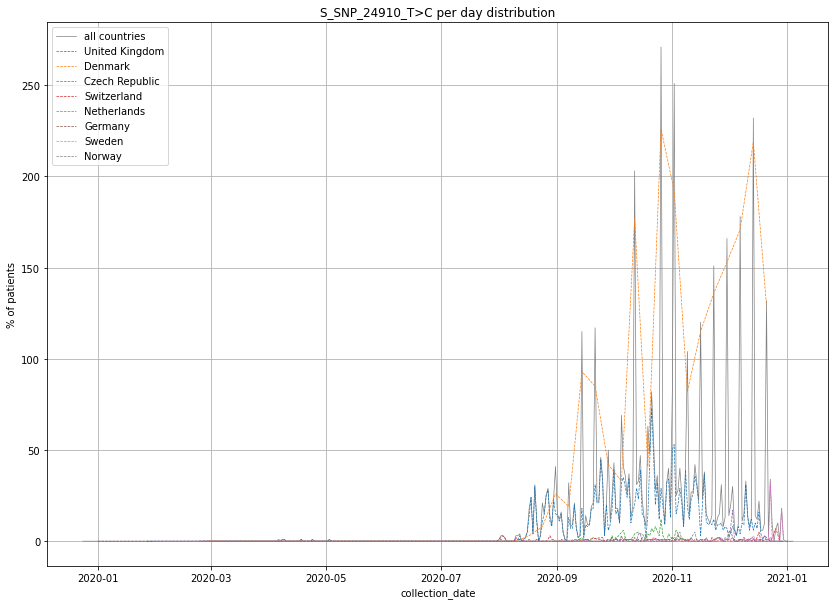

### output_20_92.png

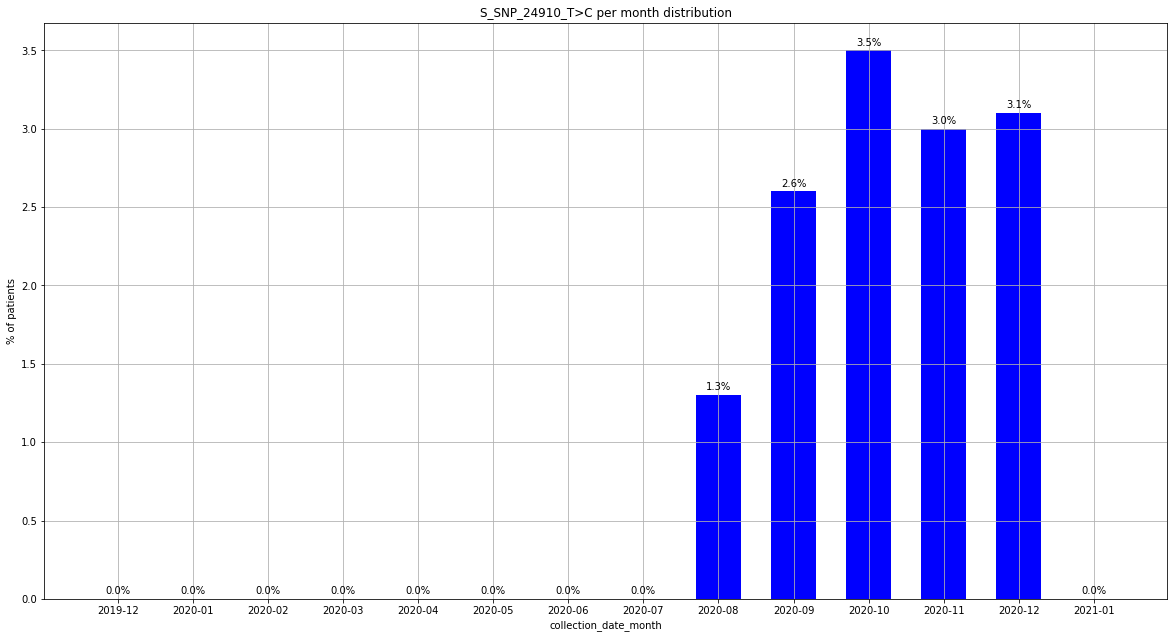

### output_20_92.png

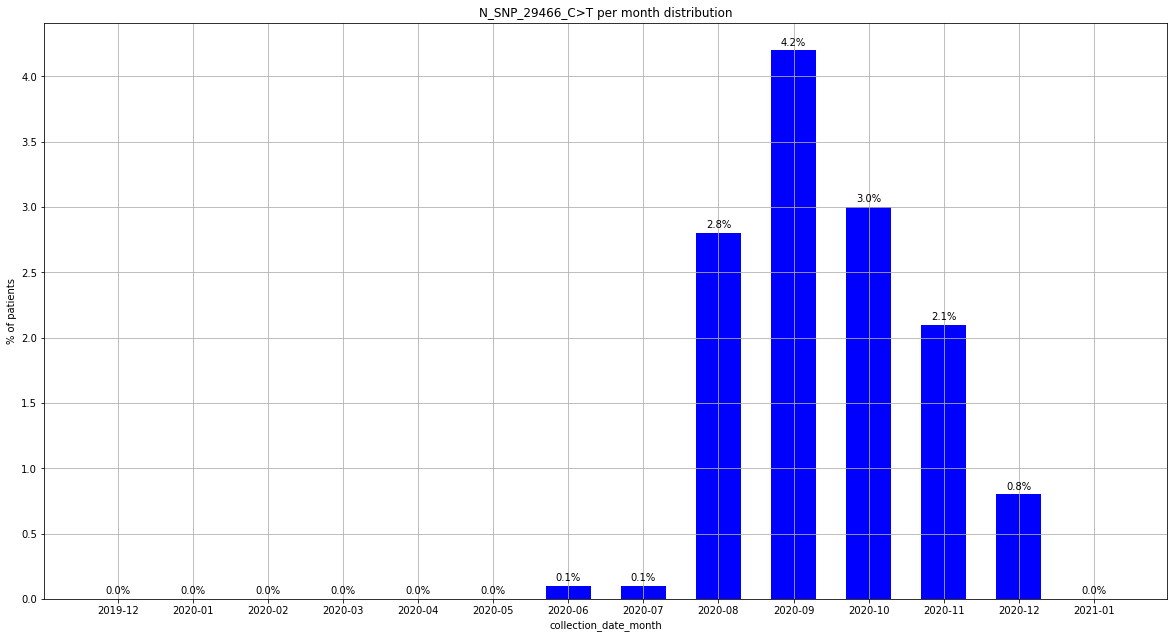

### output_20_97.png

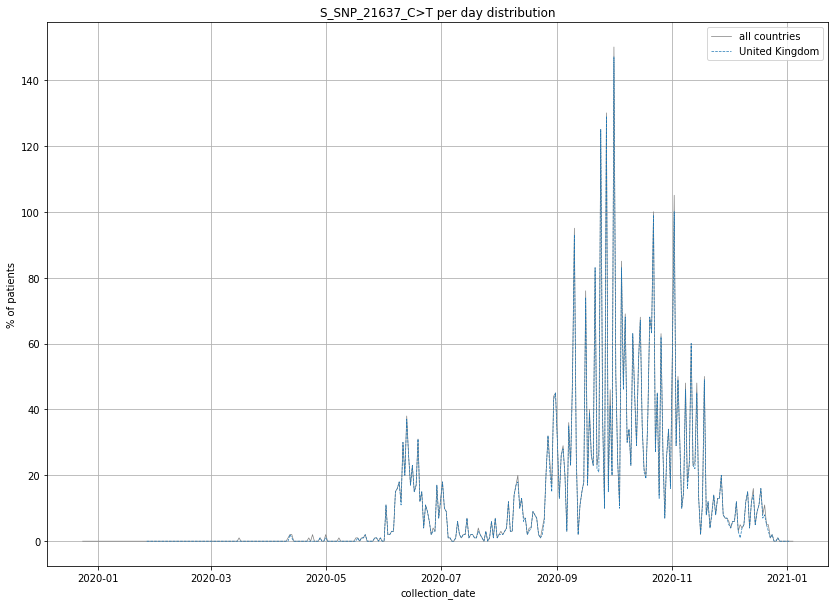

### output_20_98.png

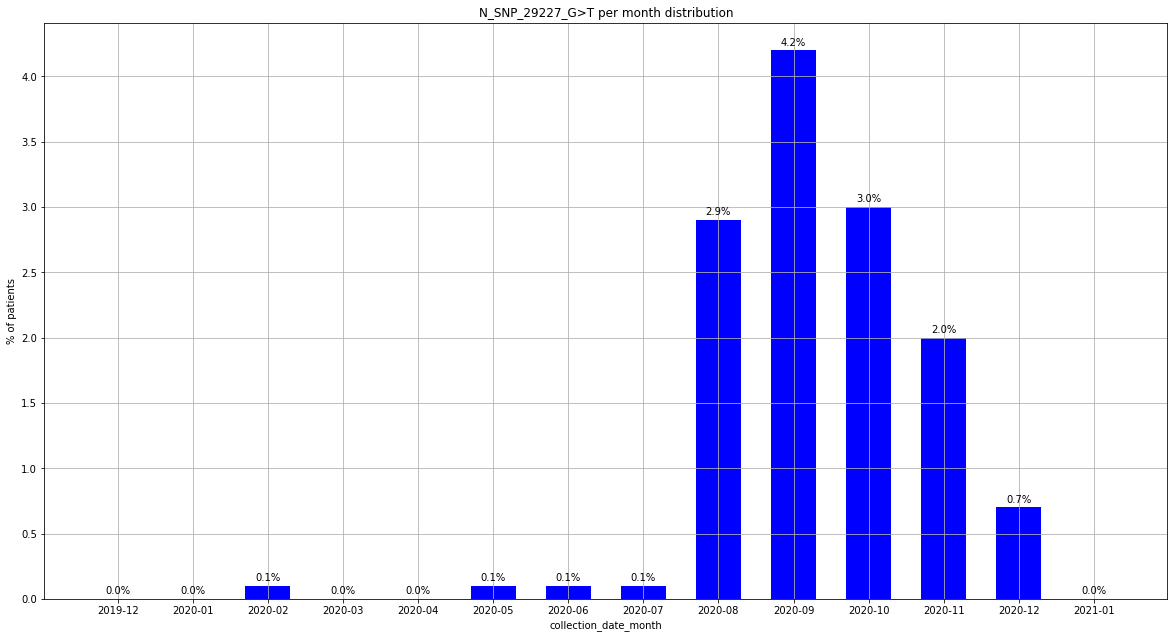

### output_20_98.png

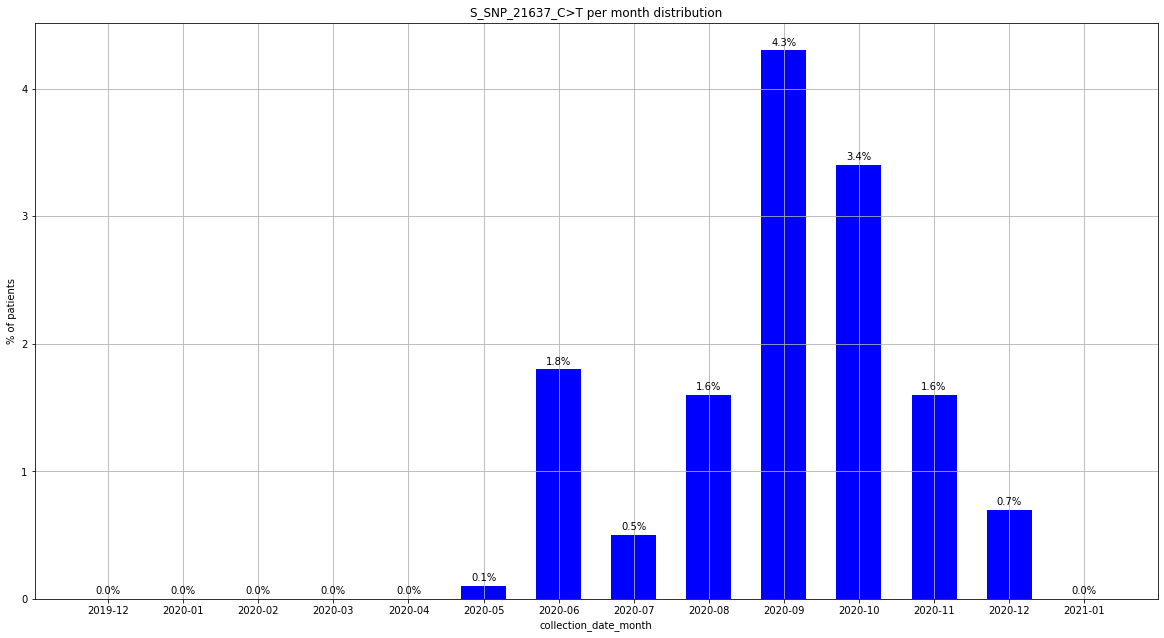

### output_20_103.png

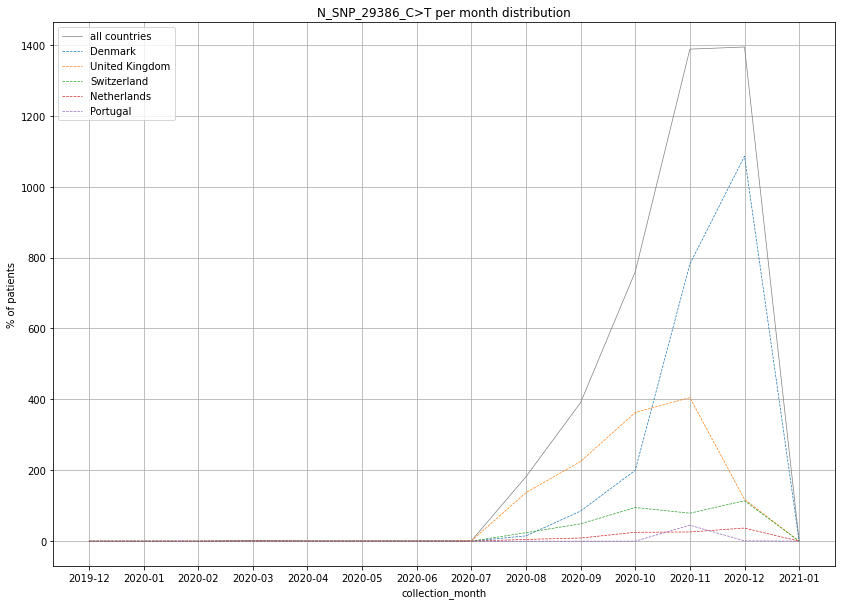

### output_20_104.png

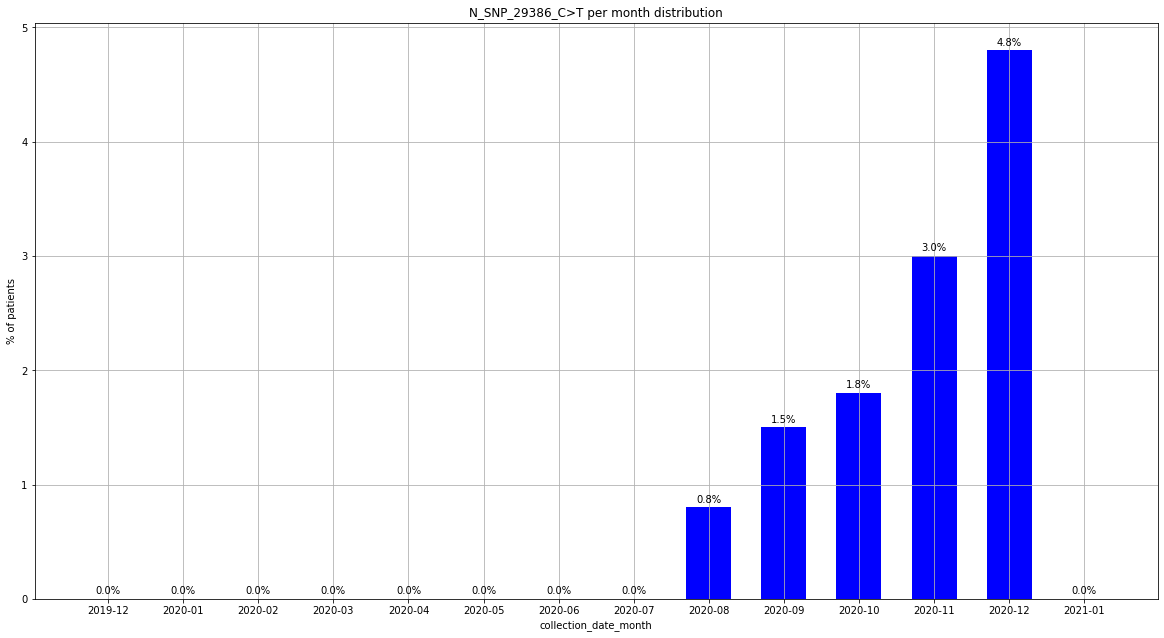

### output_20_104.png

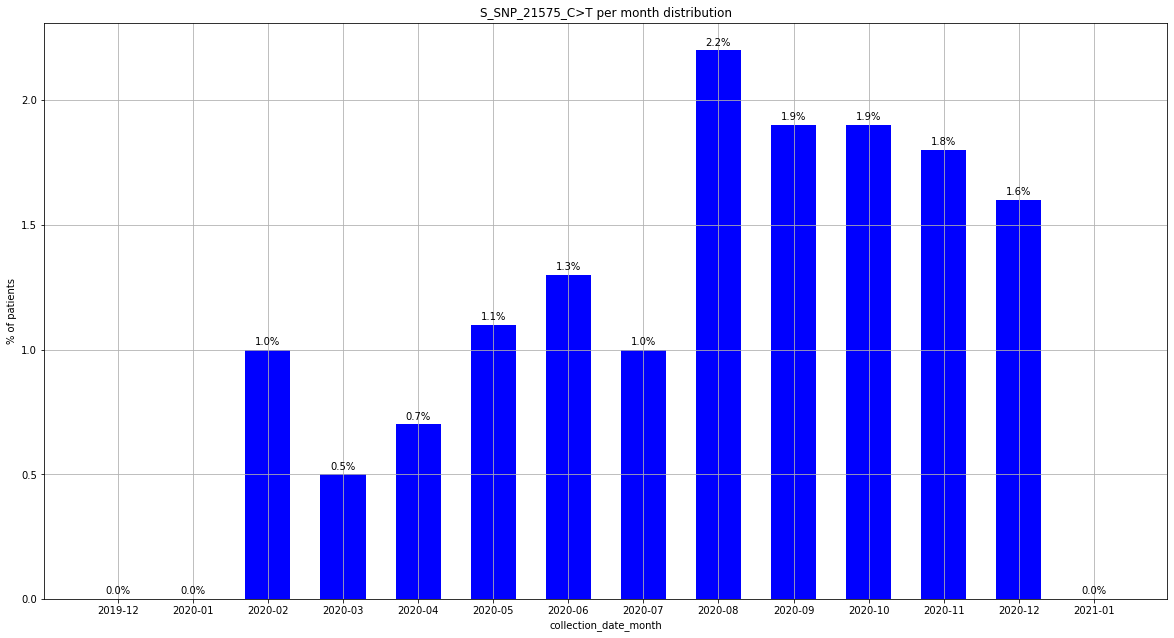

### output_20_115.png

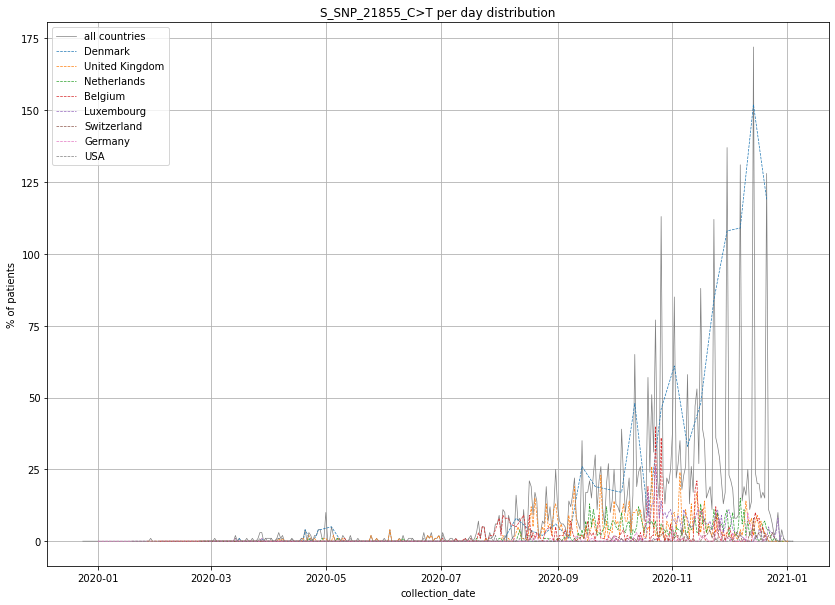

### output_20_115.png

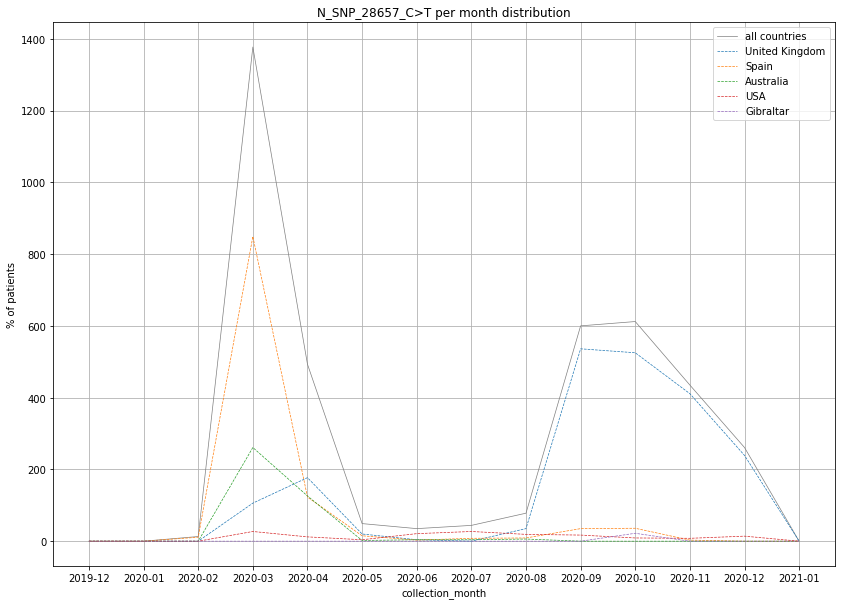

### output_20_116.png

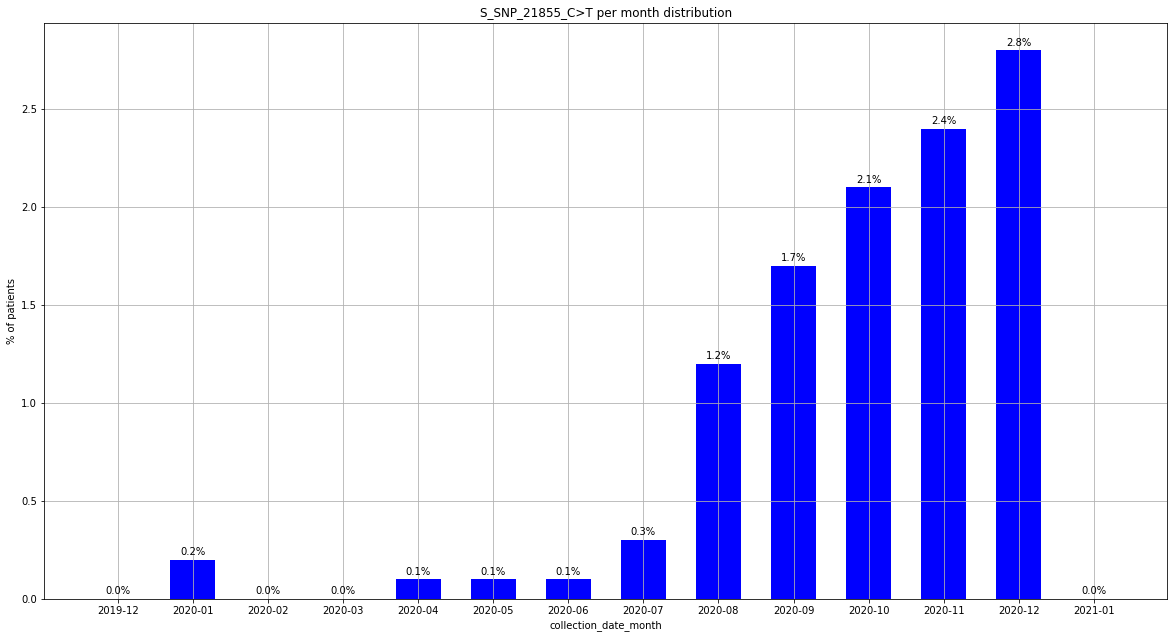

### output_20_121.png

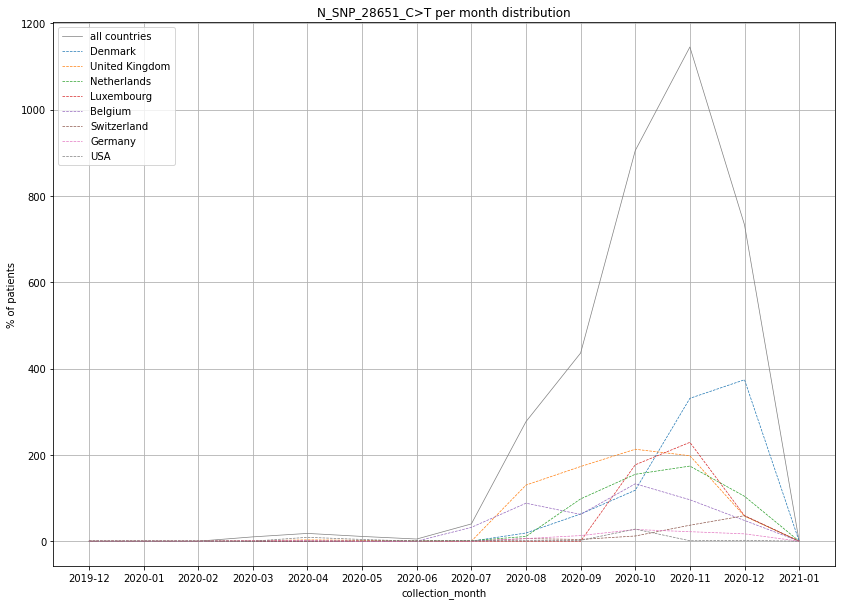

### output_20_122.png

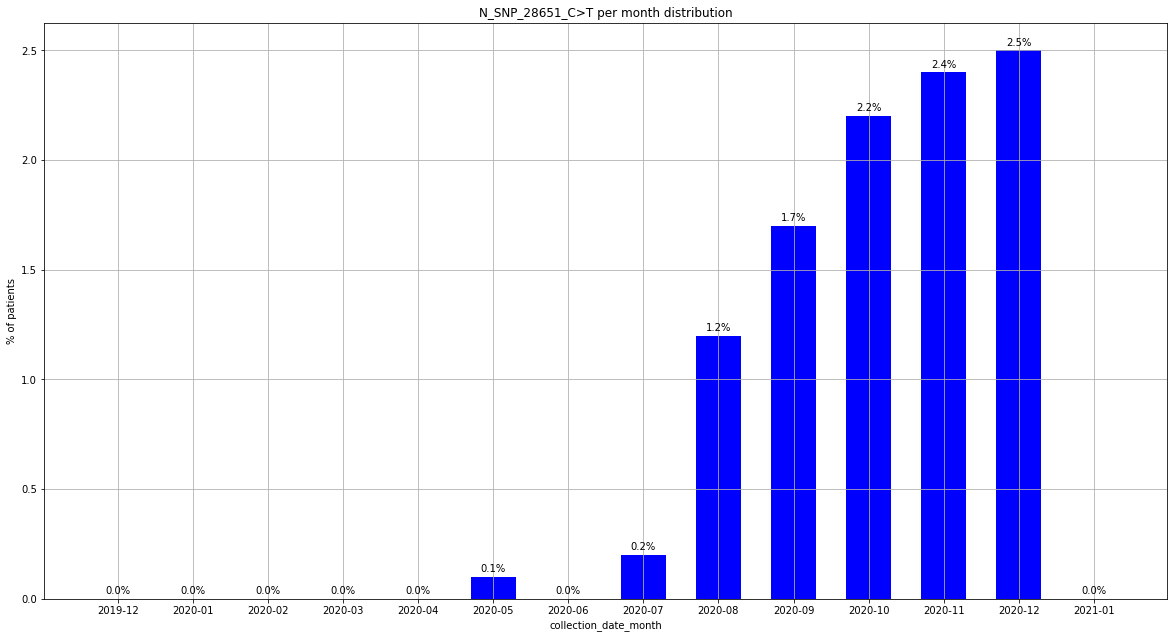

### output_20_127.png

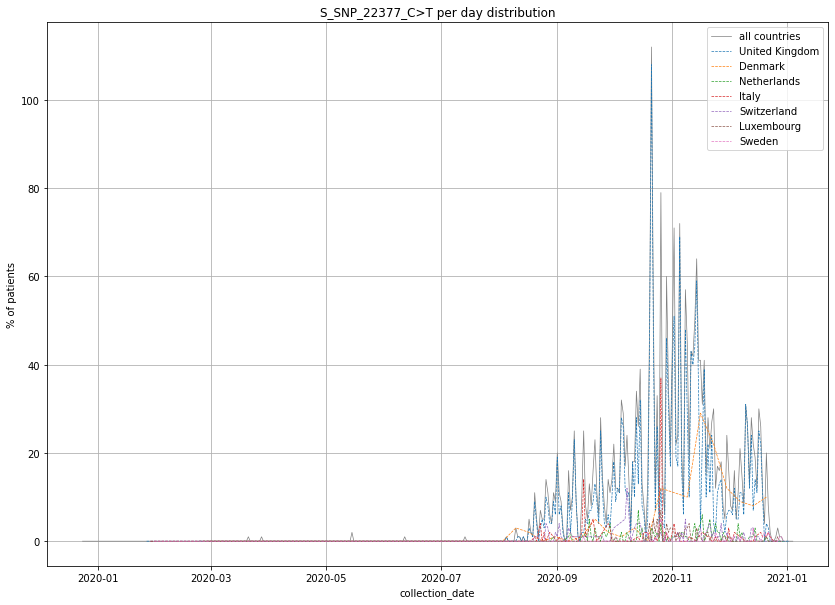

### output_20_128.png

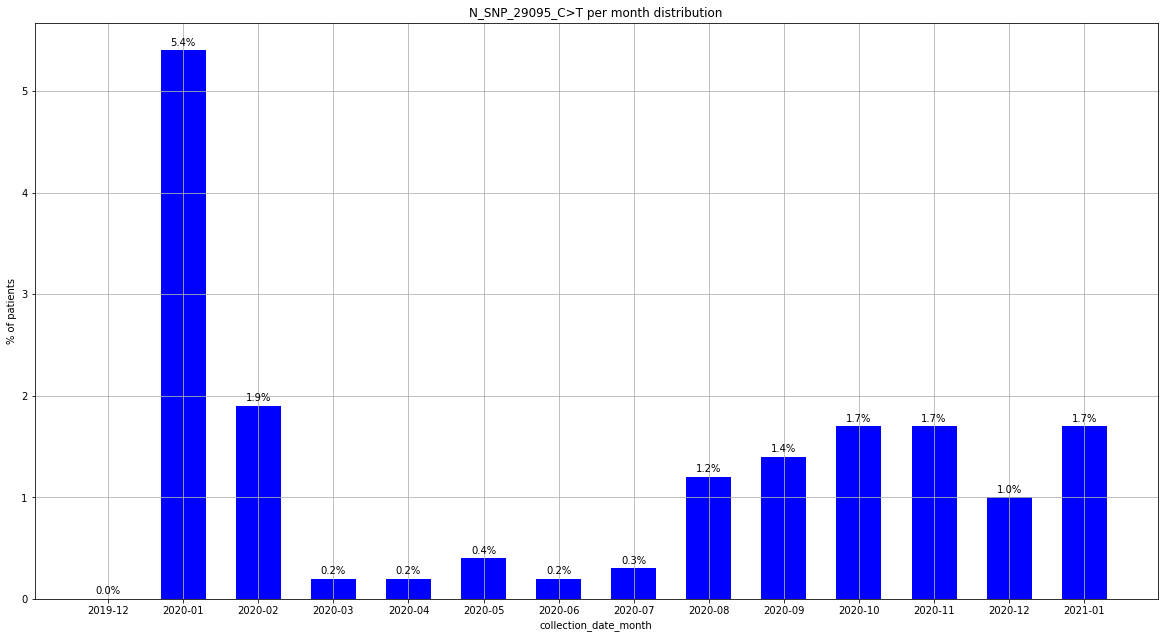

### output_20_128.png

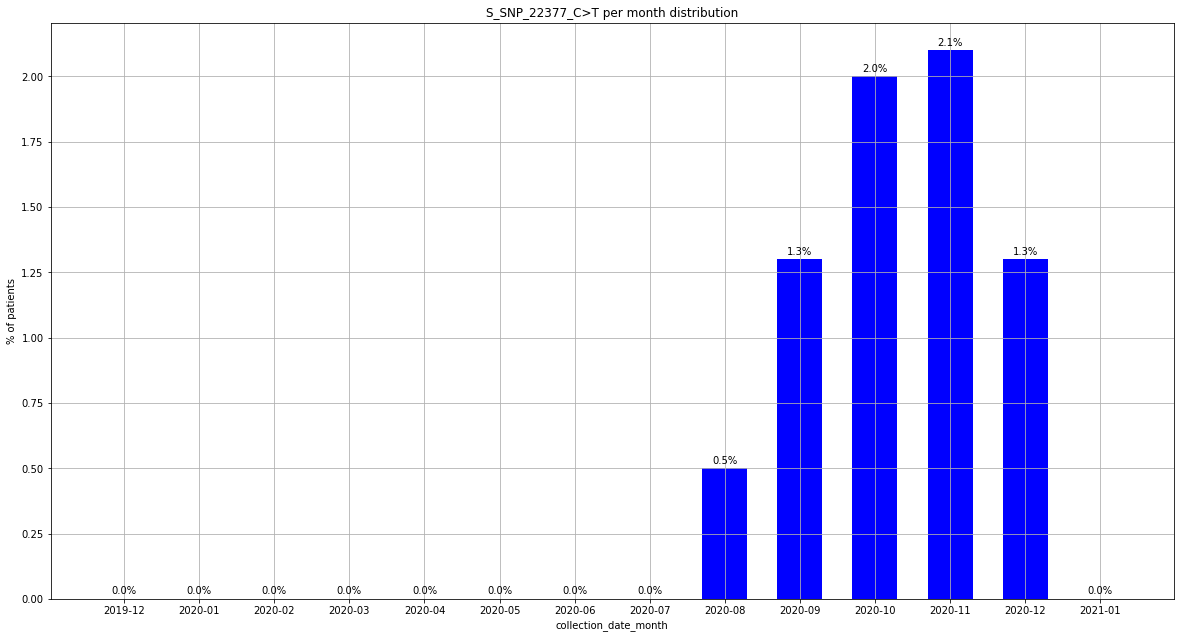

### output_20_133.png

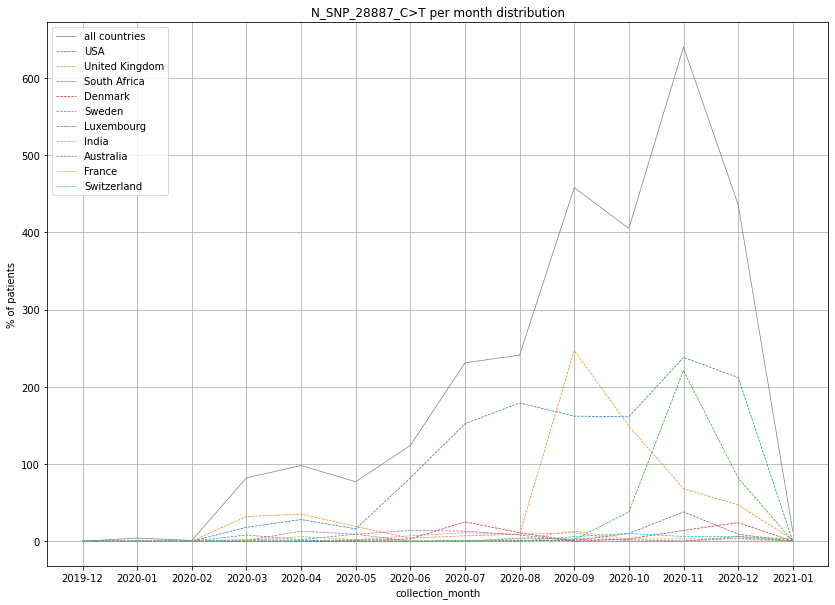

### output_20_133.png

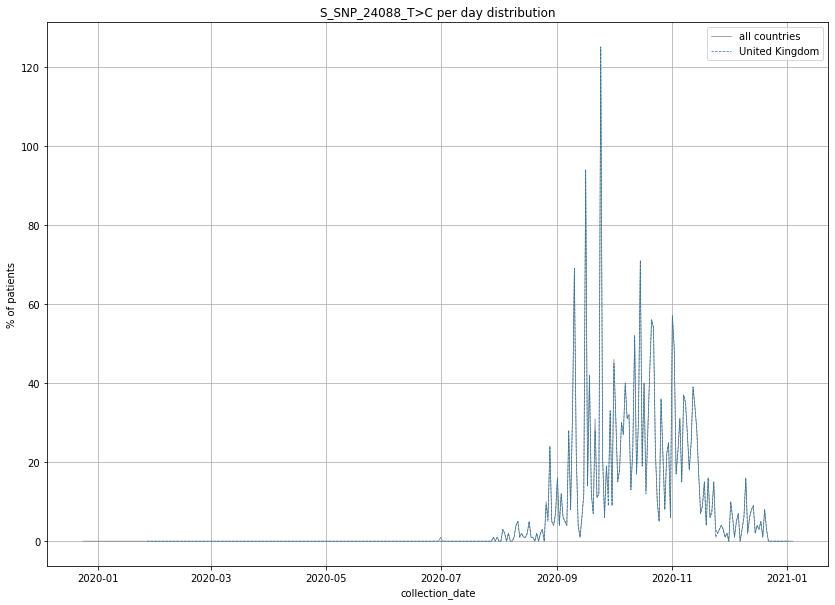

### output_20_134.png

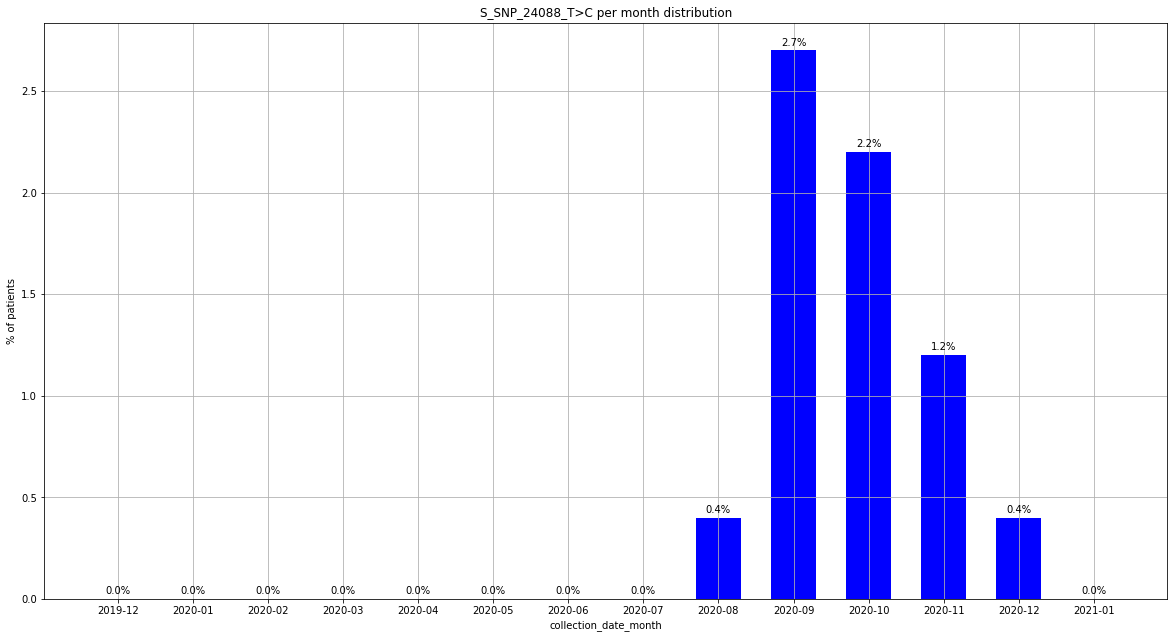

### output_20_140.png

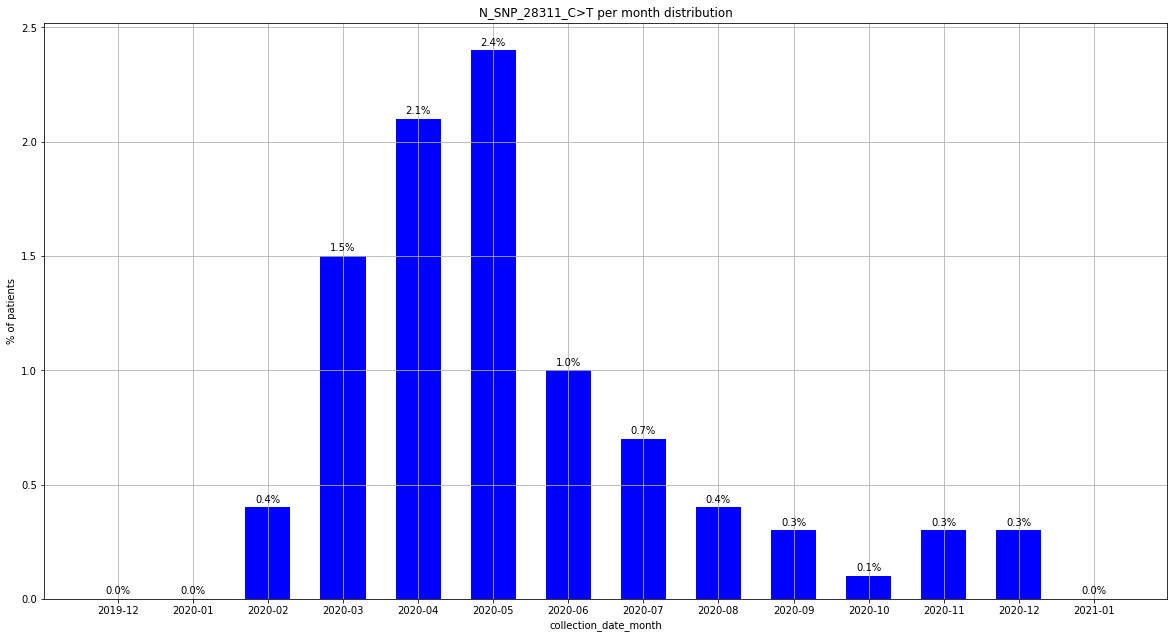

### output_20_151.png

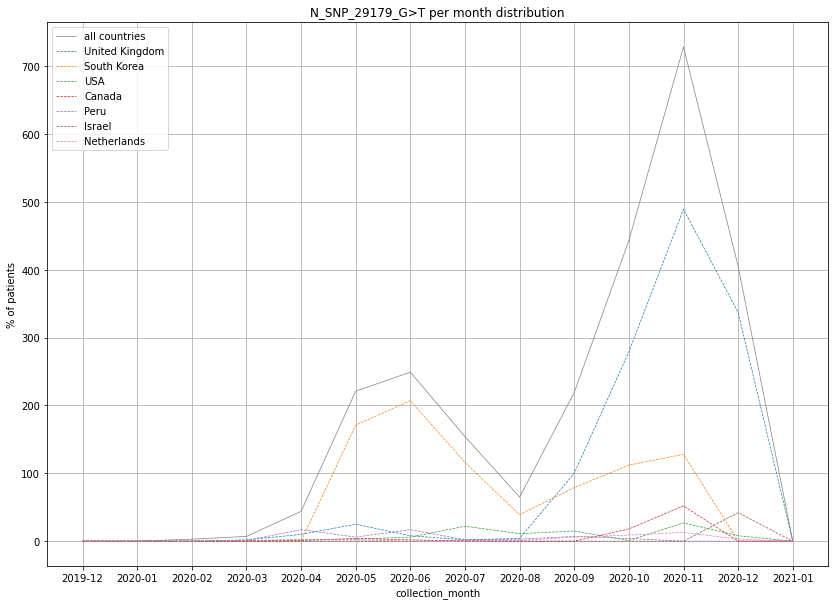

### output_20_152.png

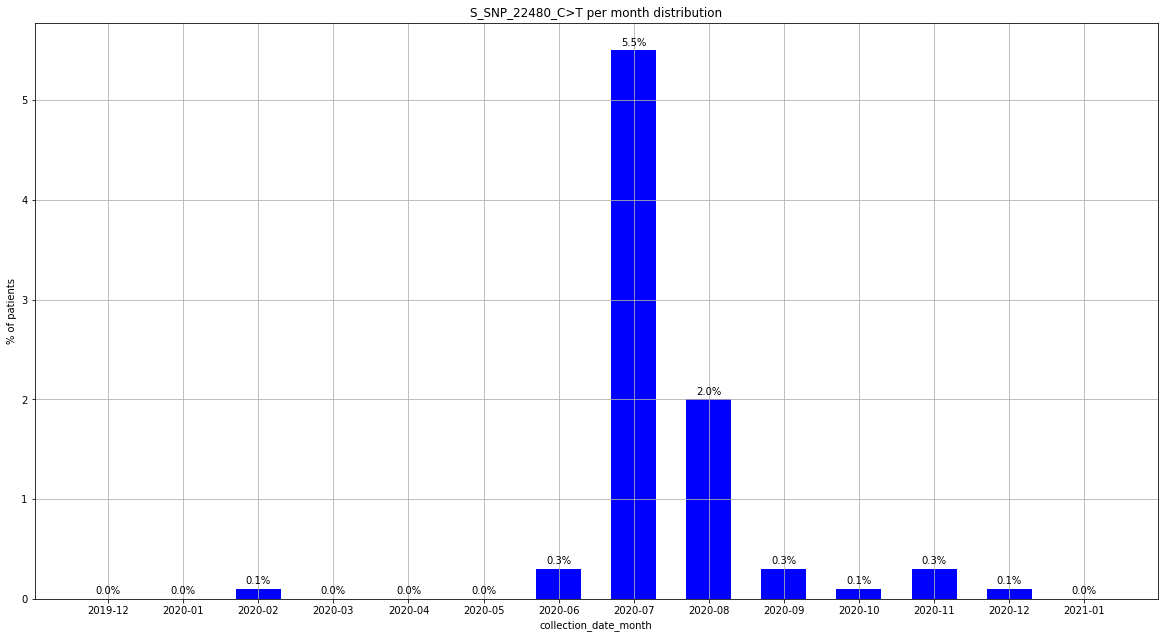

### output_20_157.png

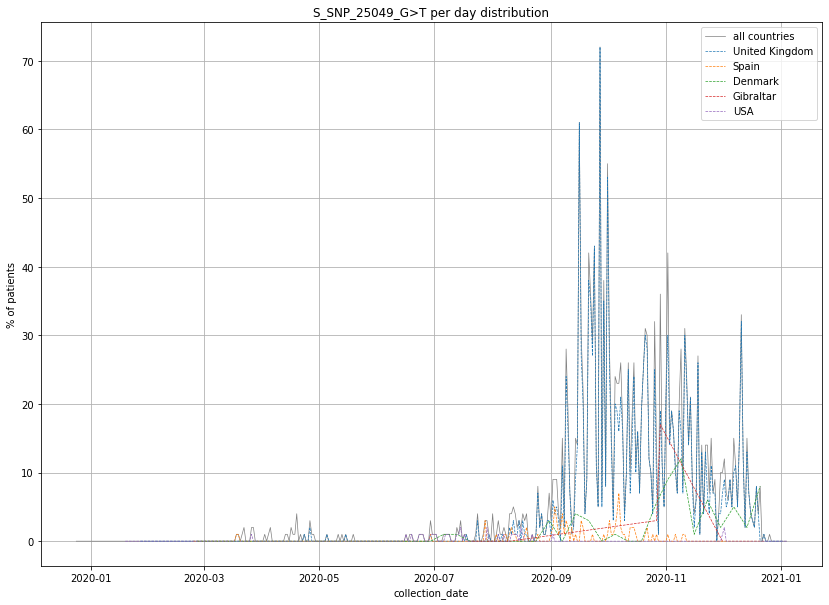

### output_20_158.png

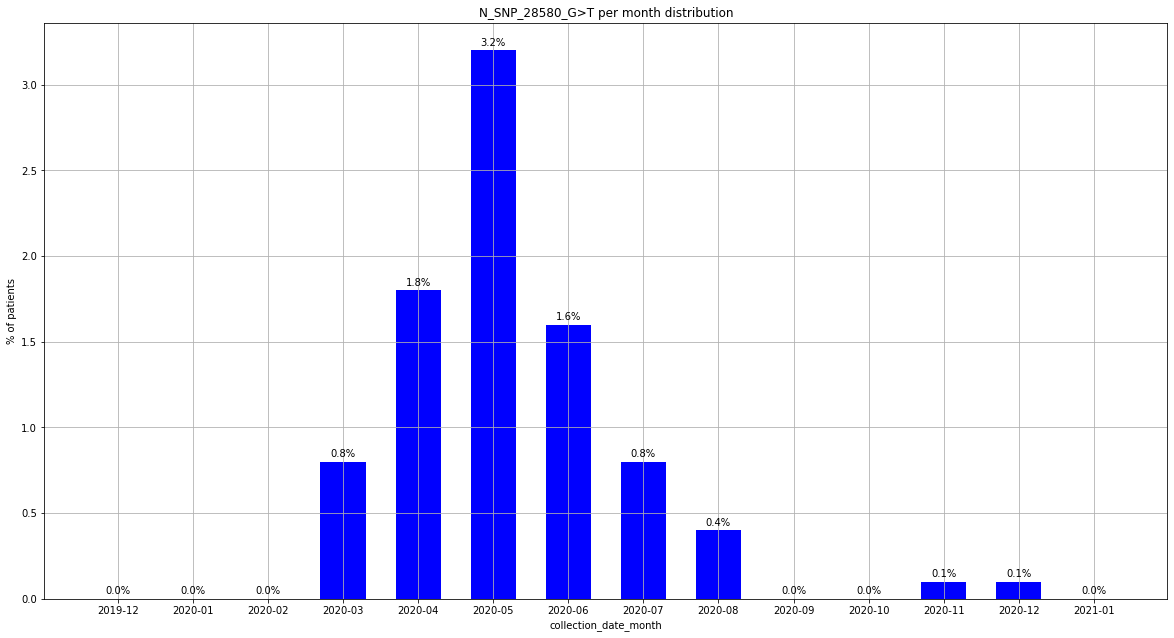

### output_20_163.png

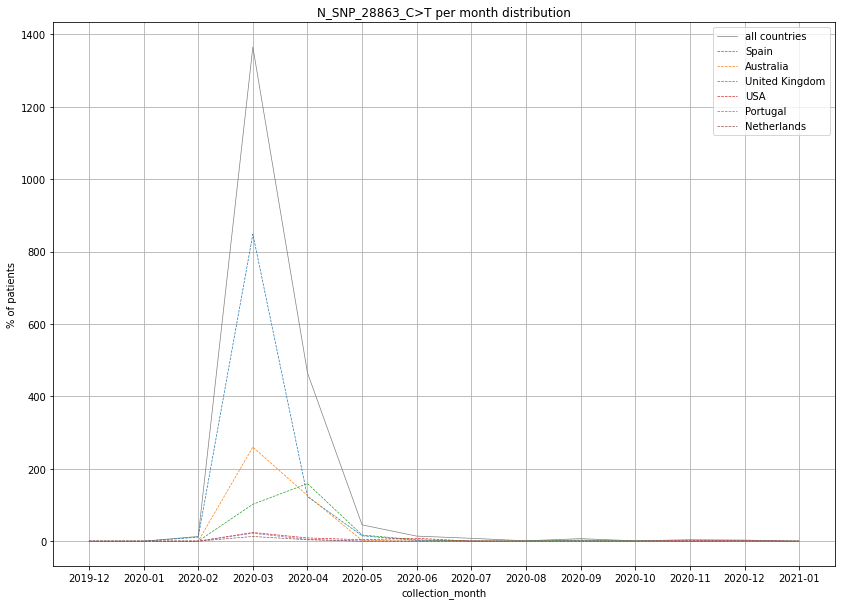

### output_20_164.png

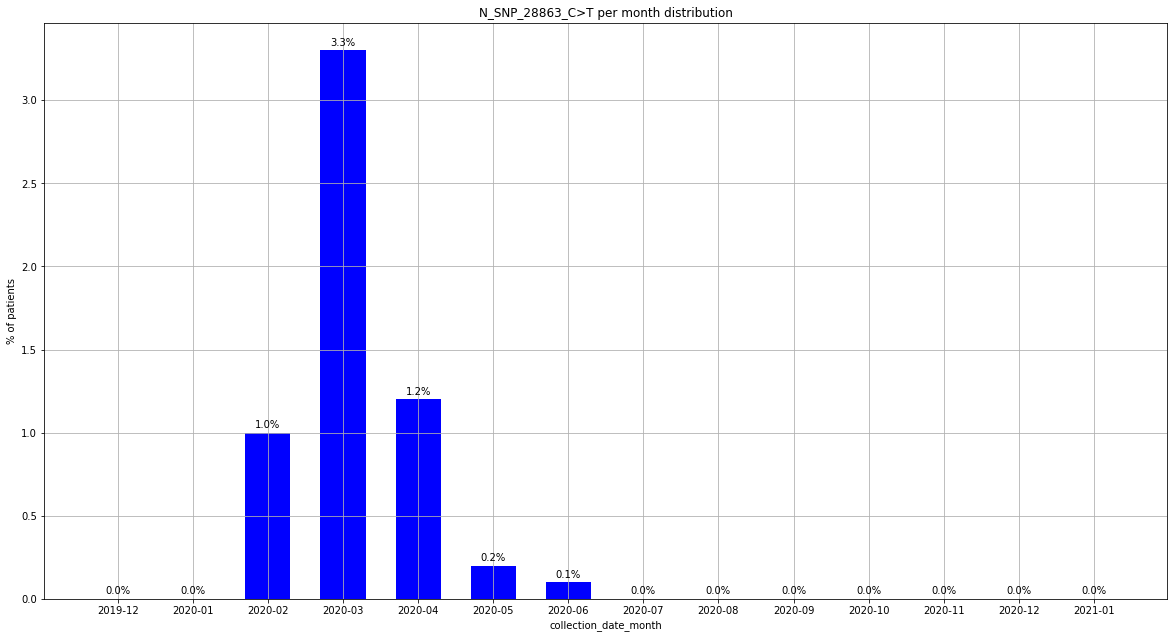

### output_20_164.png

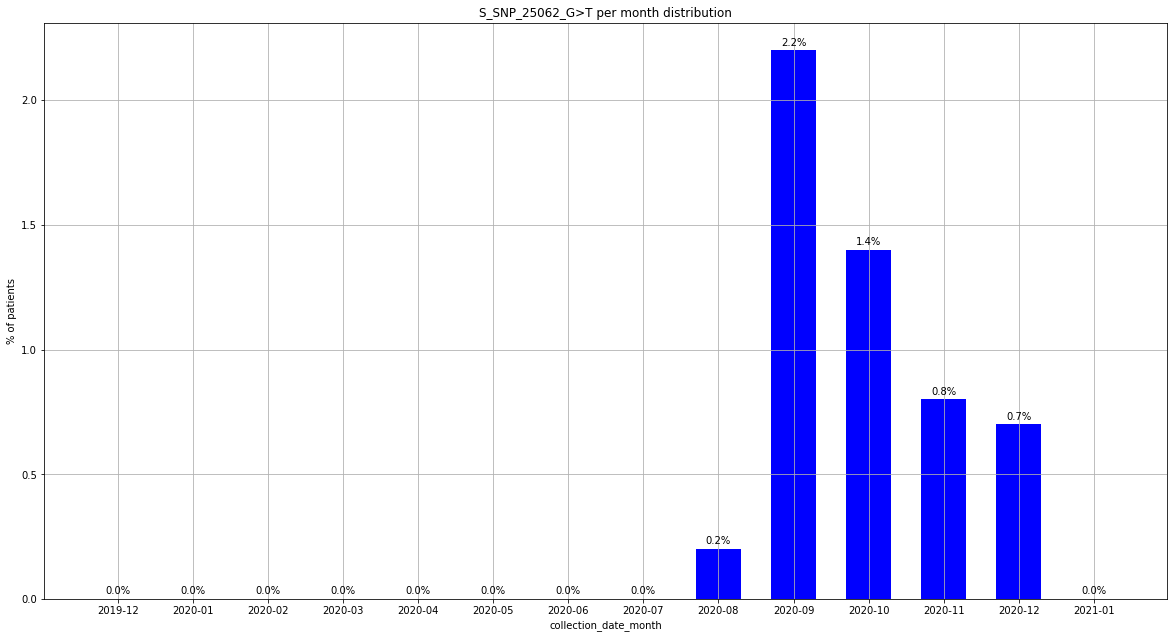

### output_20_175.png

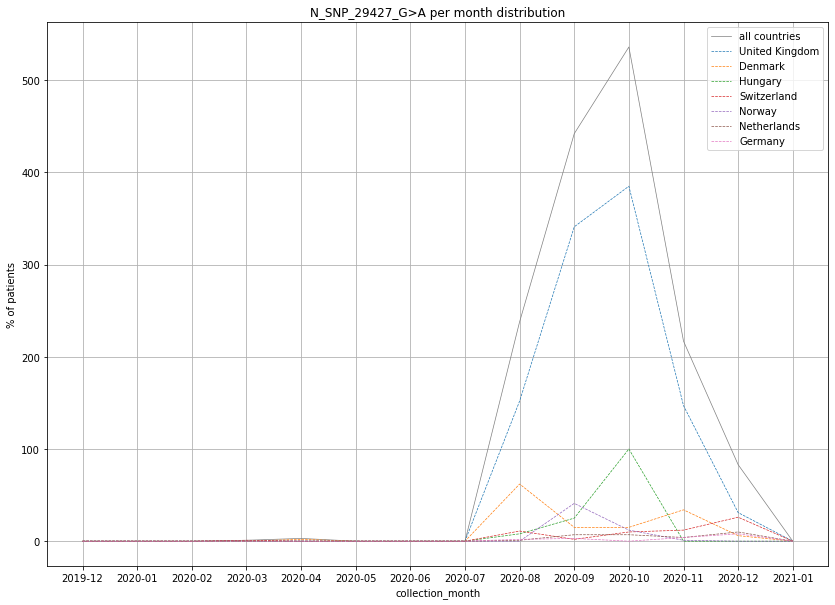

### output_20_175.png

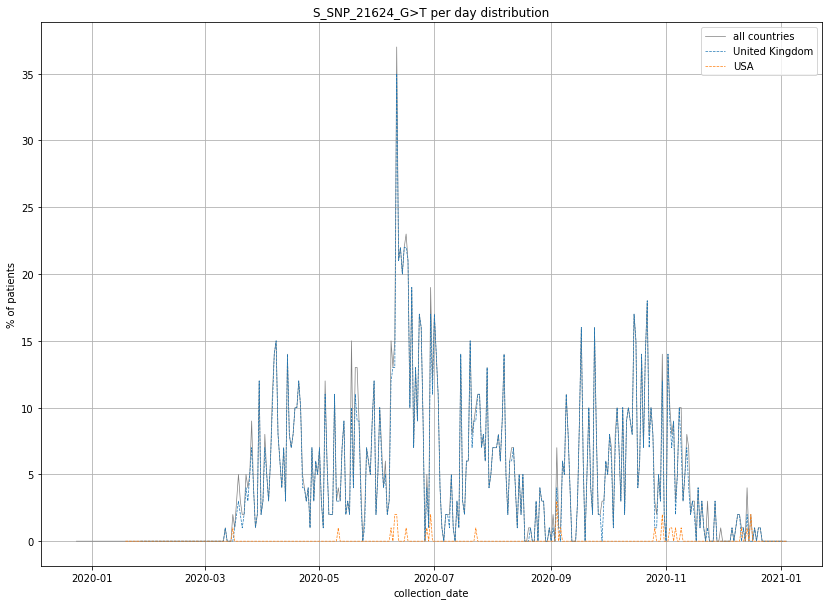

### output_20_176.png

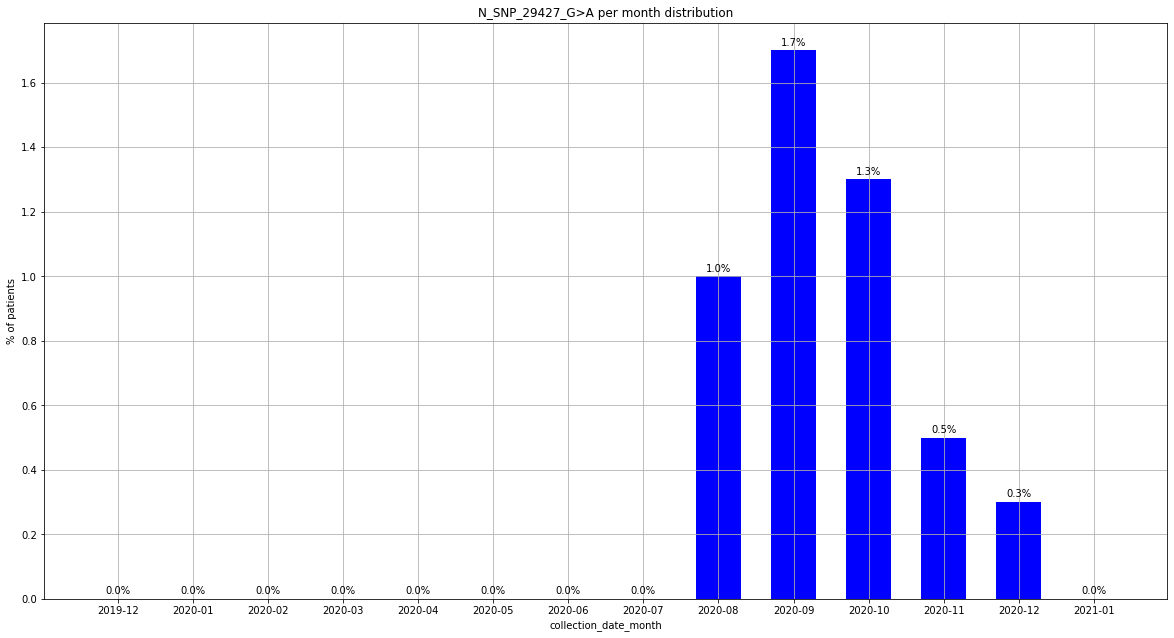

### output_20_181.png

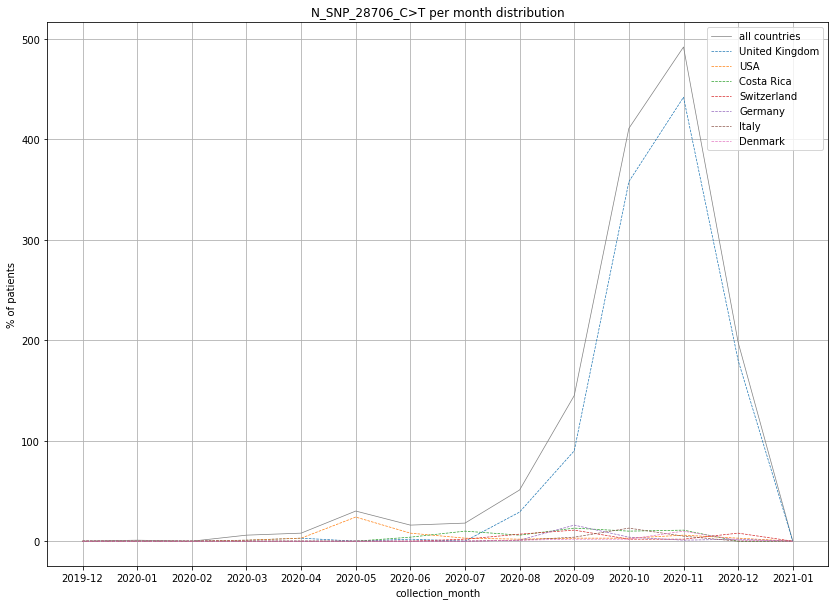

### output_20_182.png

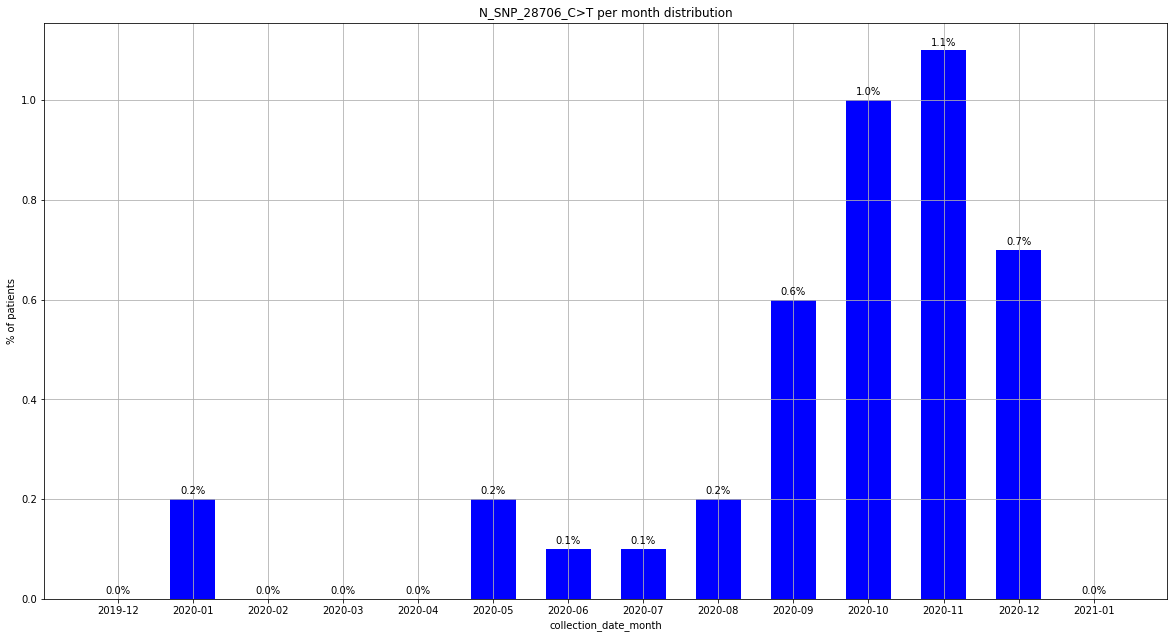

### output_20_188.png

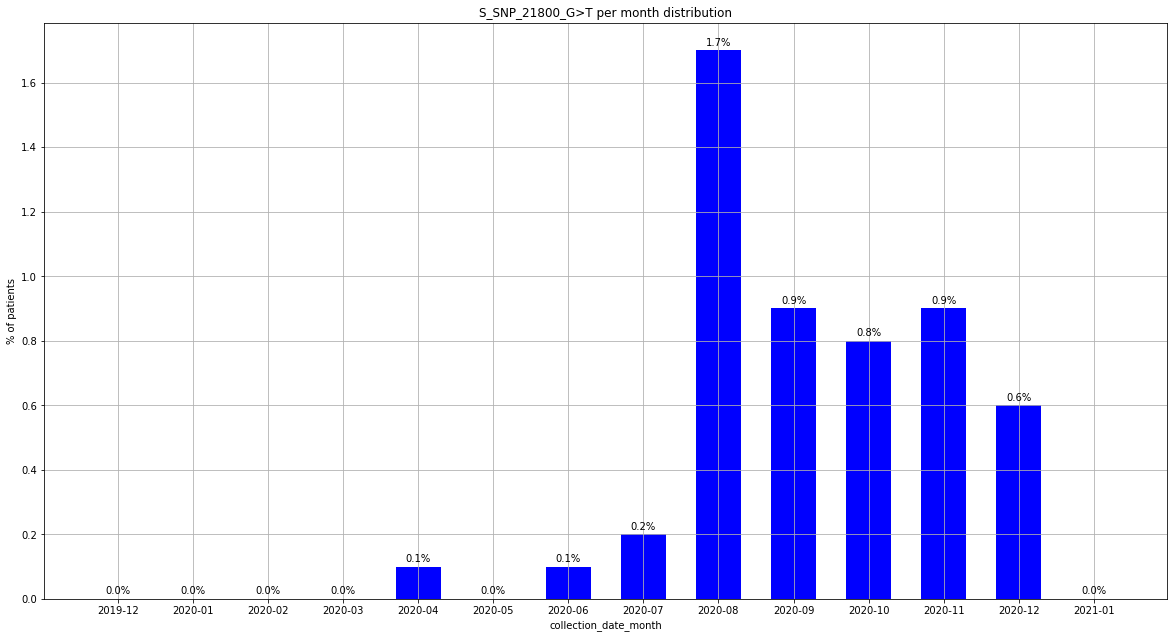

### output_20_193.png

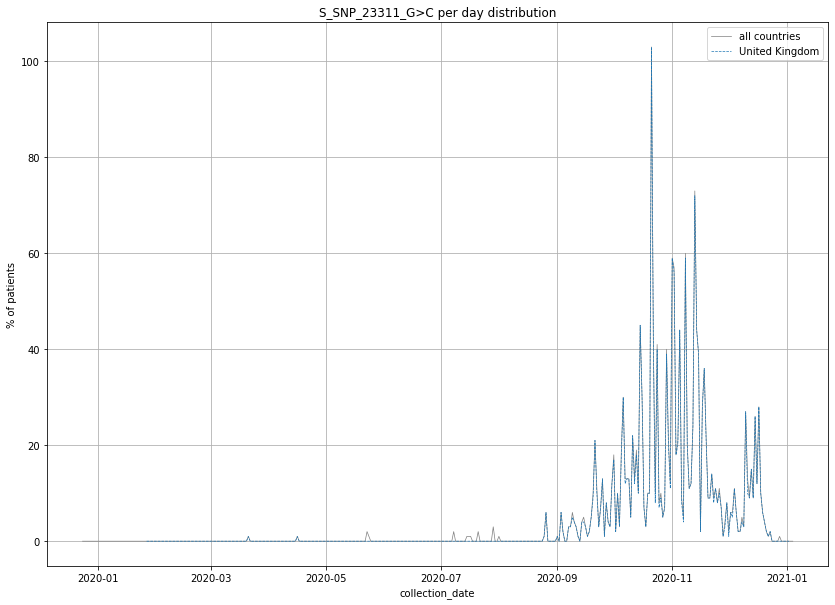

### output_20_194.png

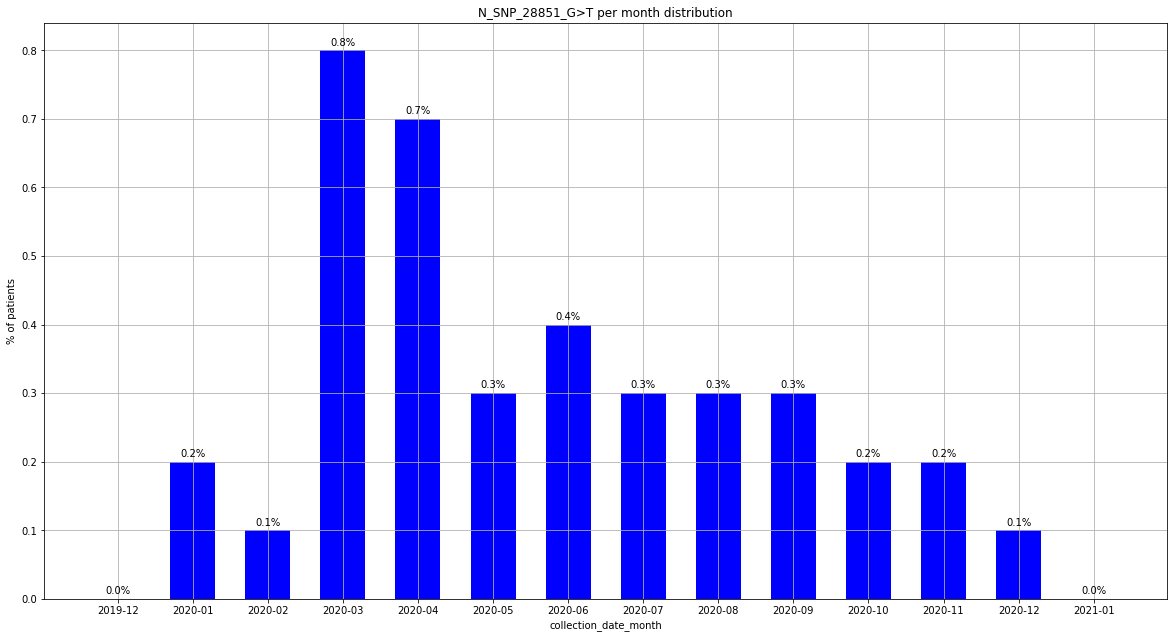

### output_20_199.png

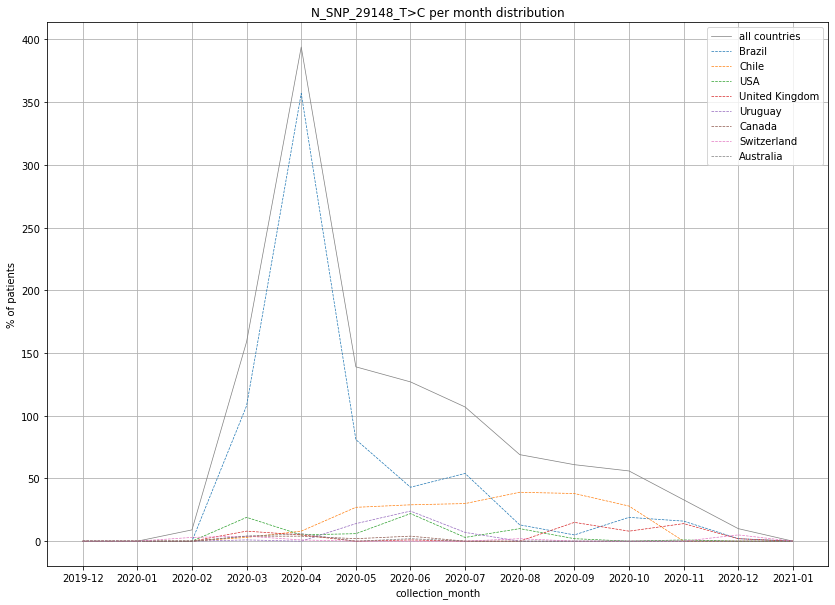

### output_20_206.png

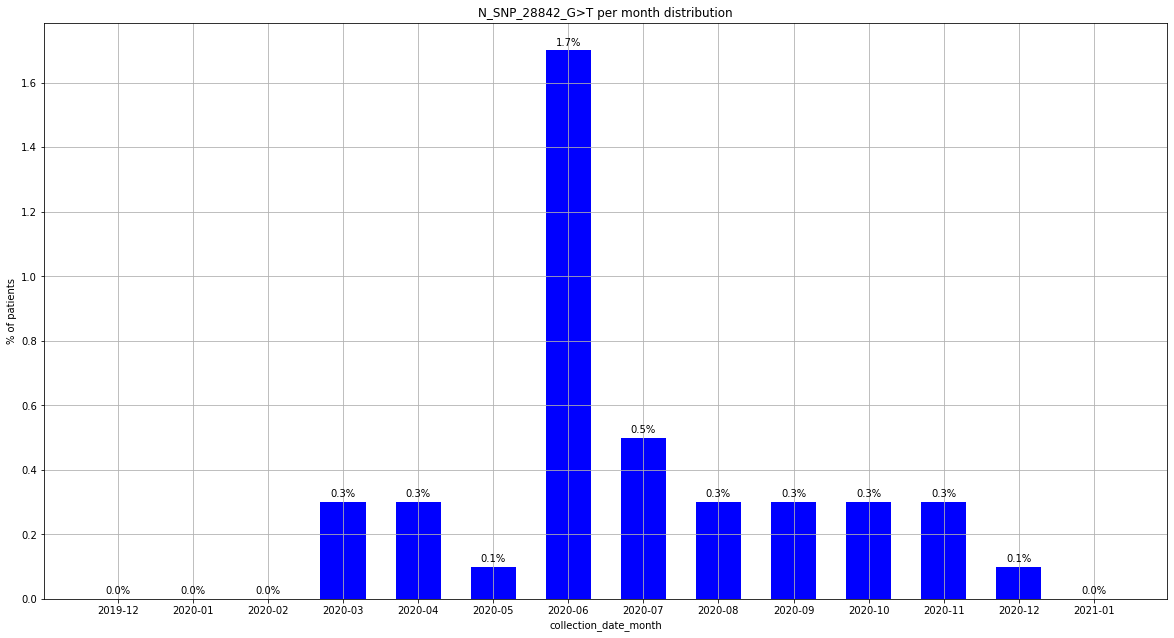

### output_20_211.png

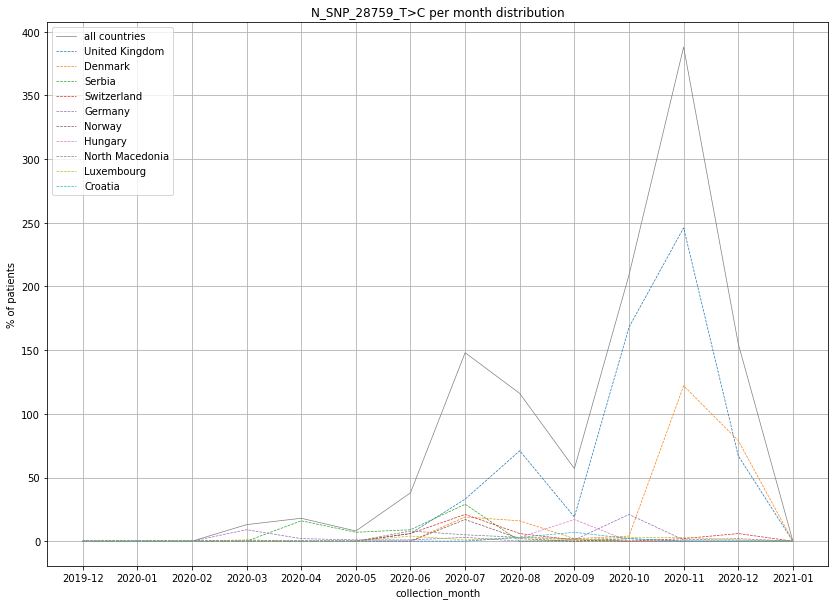

### output_20_212.png

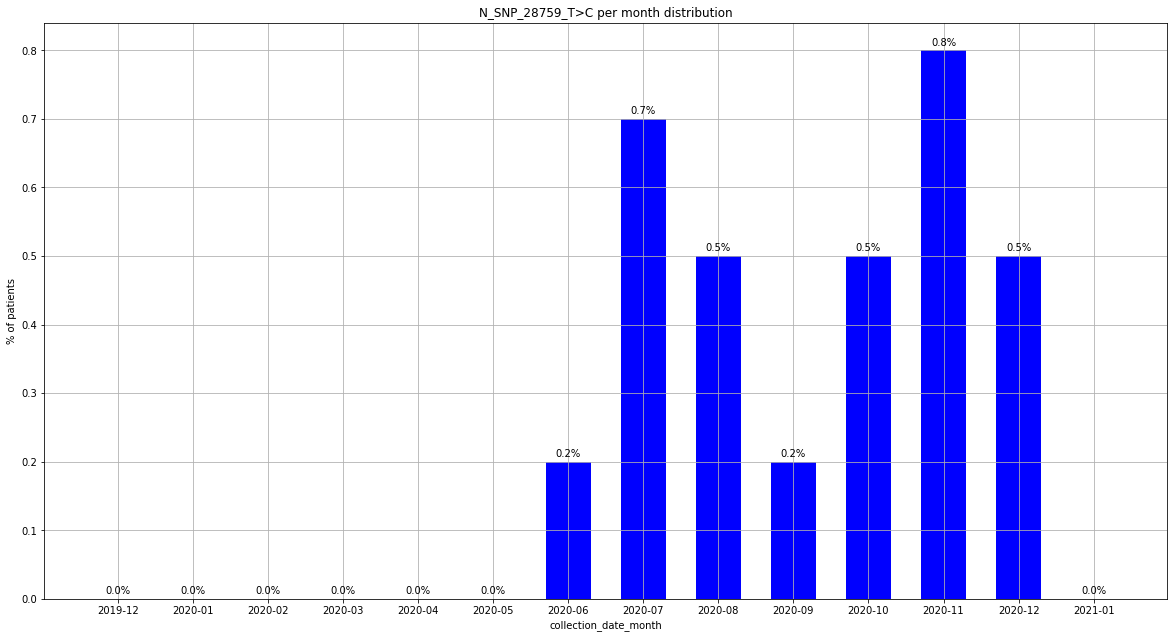

### output_20_217.png

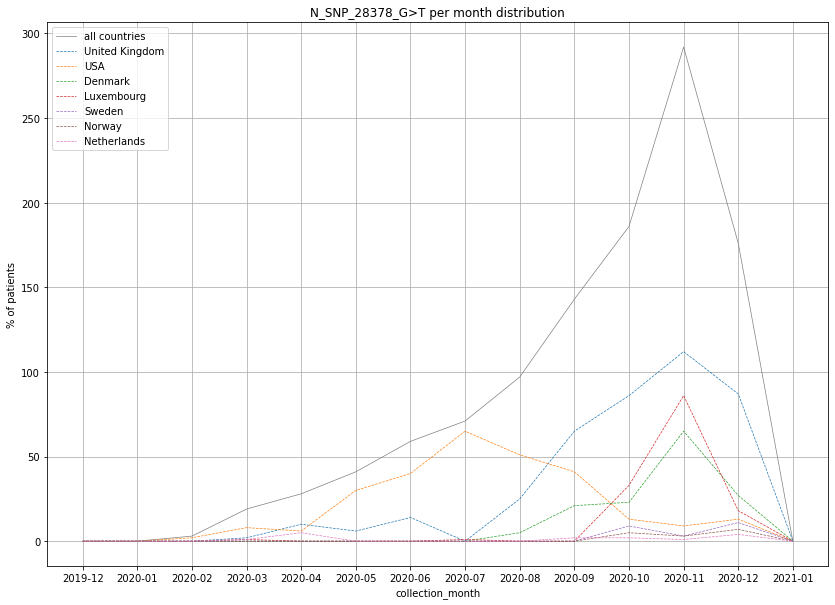

### output_20_218.png

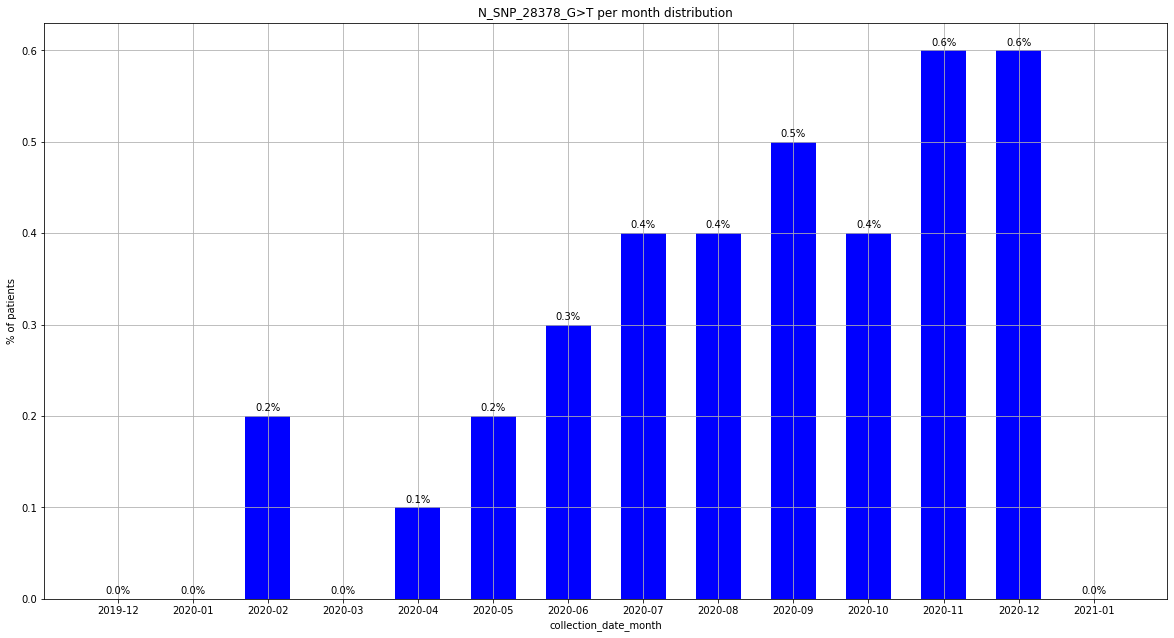

### output_20_223.png

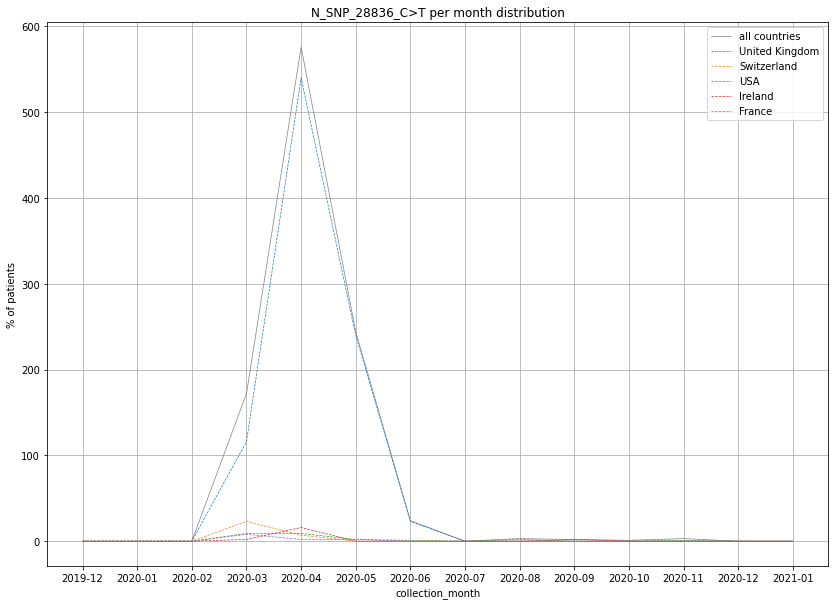

### output_20_223.png

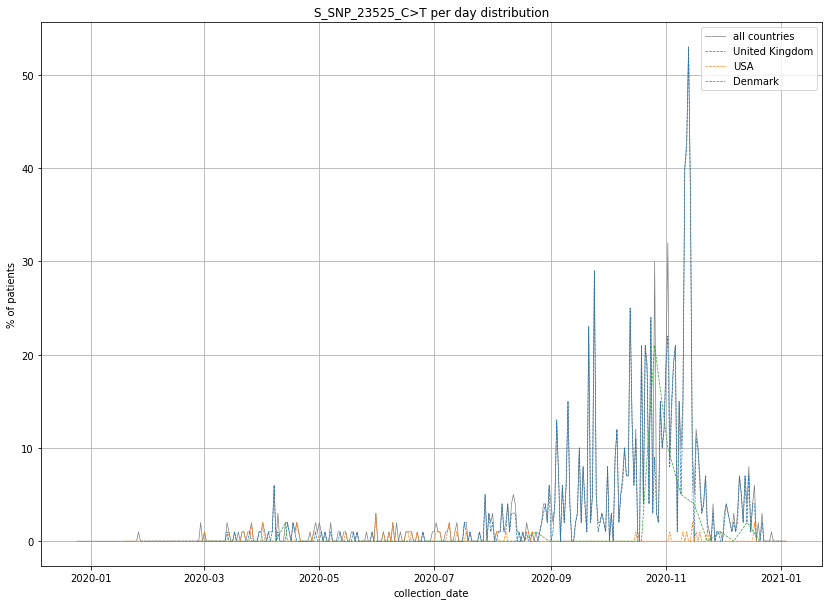

### output_20_224.png

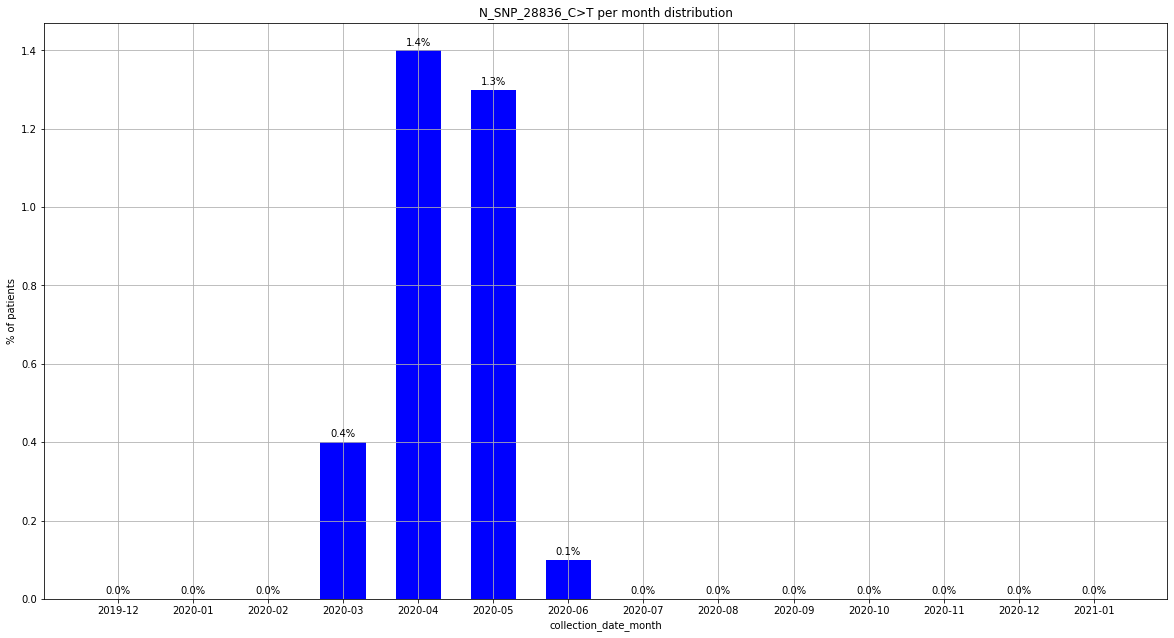

### output_20_224.png

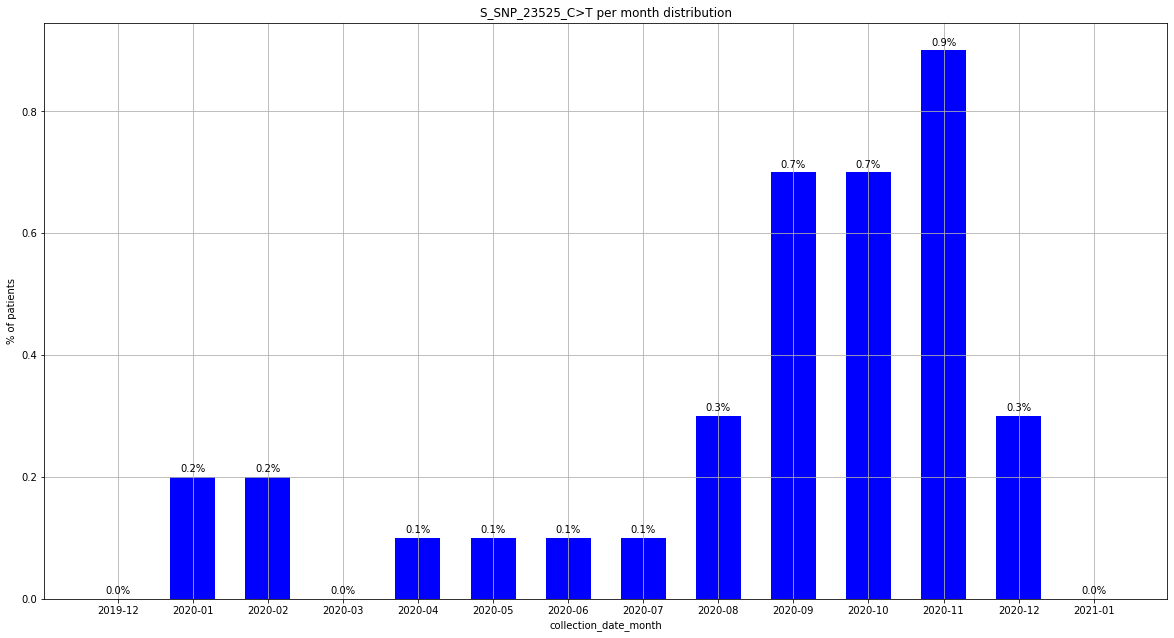

### output_20_230.png

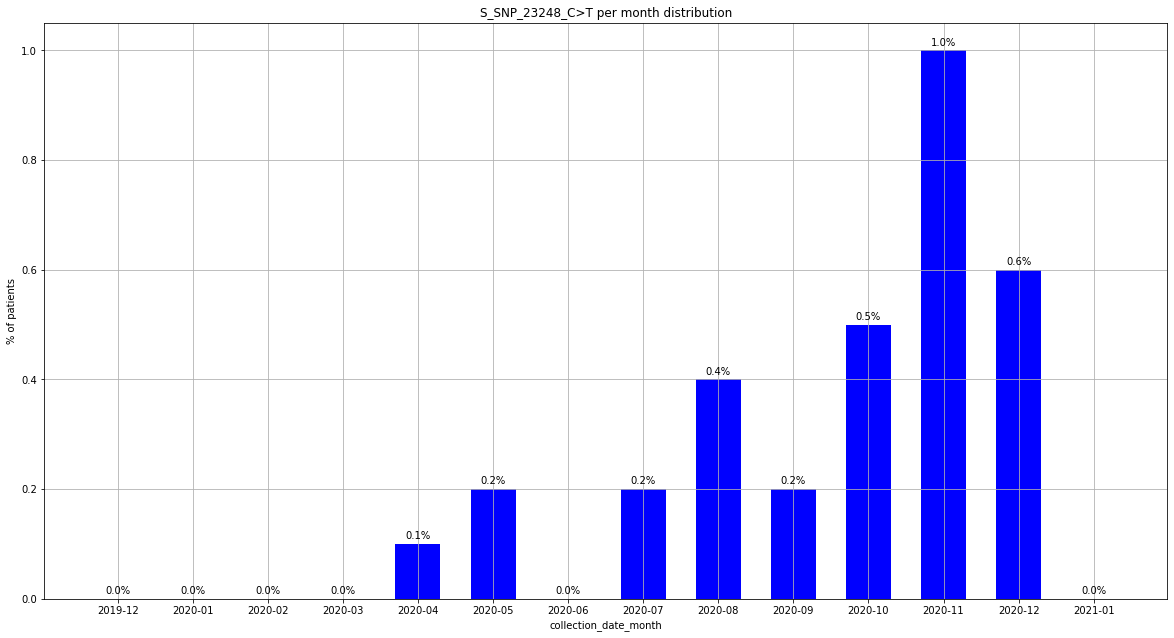

### output_20_230.png

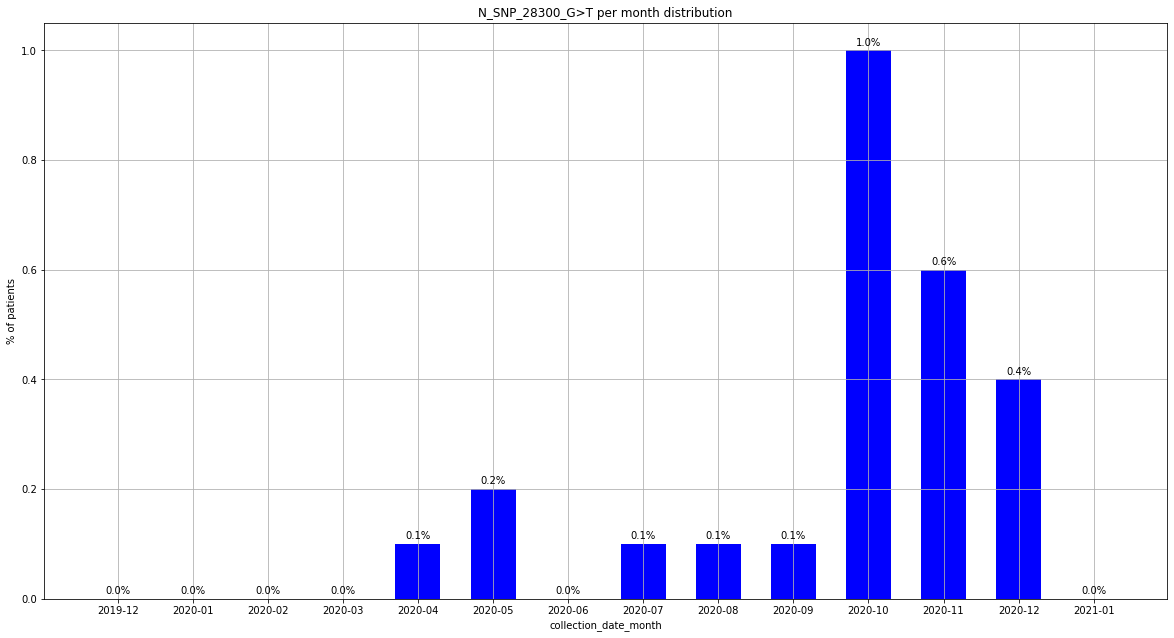

### output_20_235.png

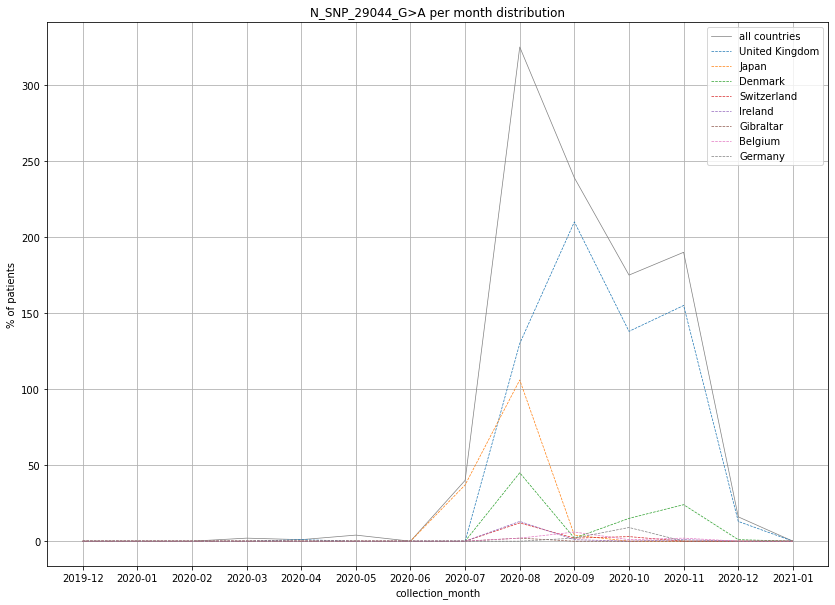

### output_20_236.png

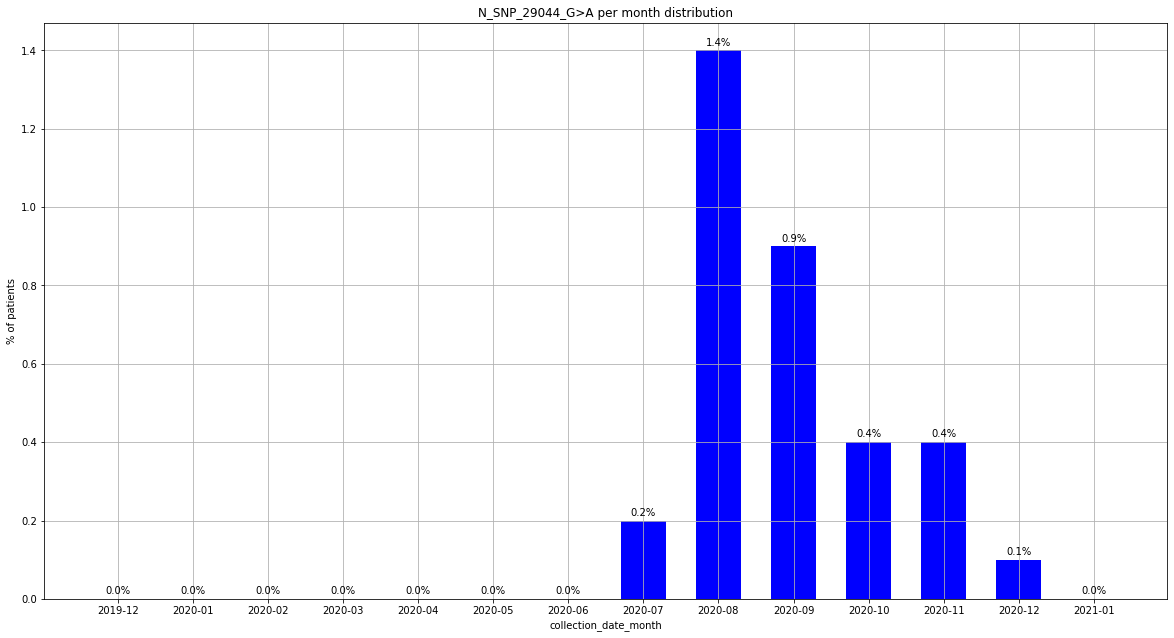

### output_20_248.png

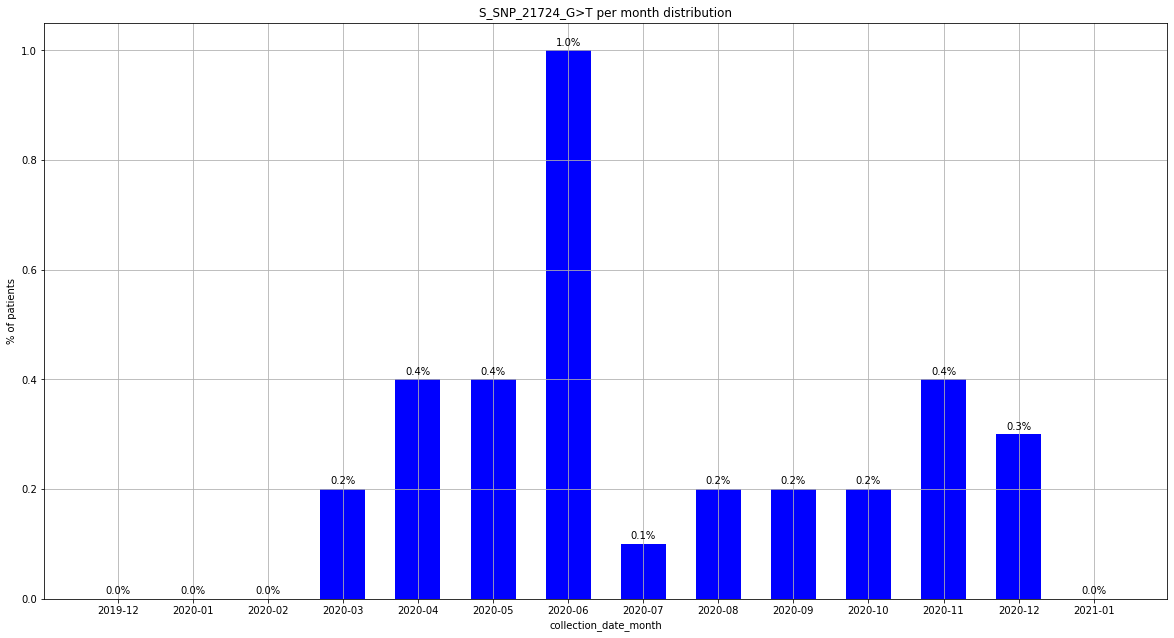

### output_20_254.png

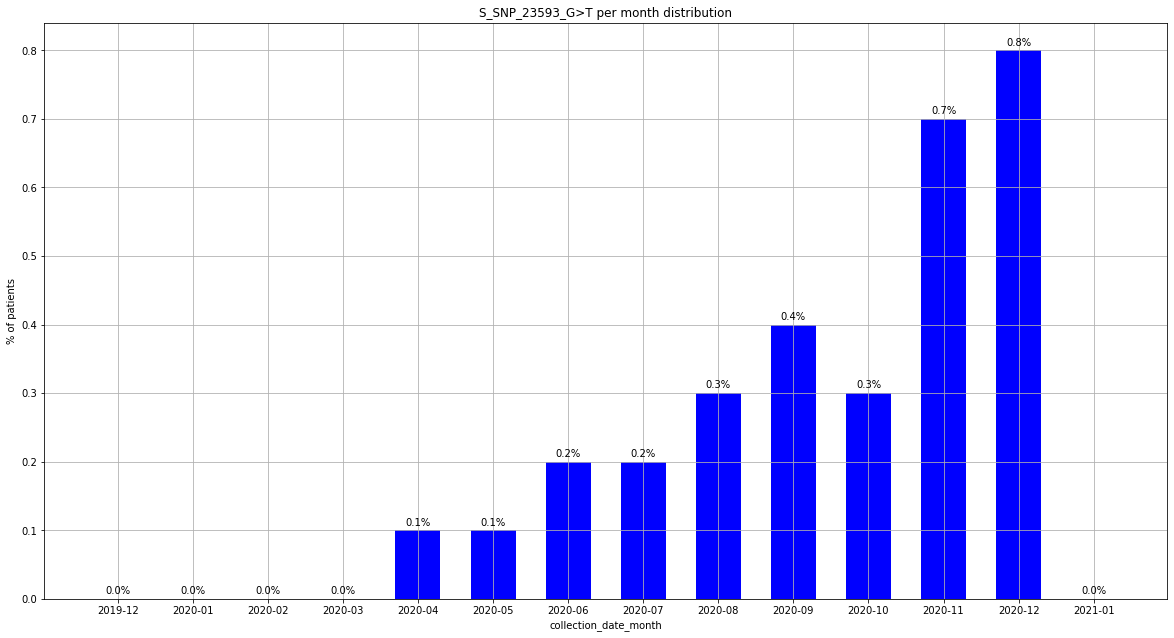

### output_20_259.png

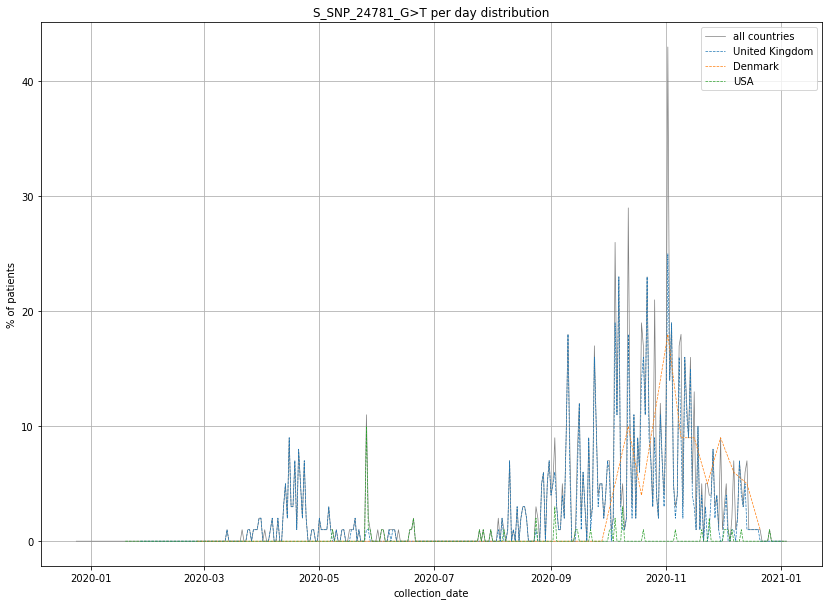

### output_20_266.png

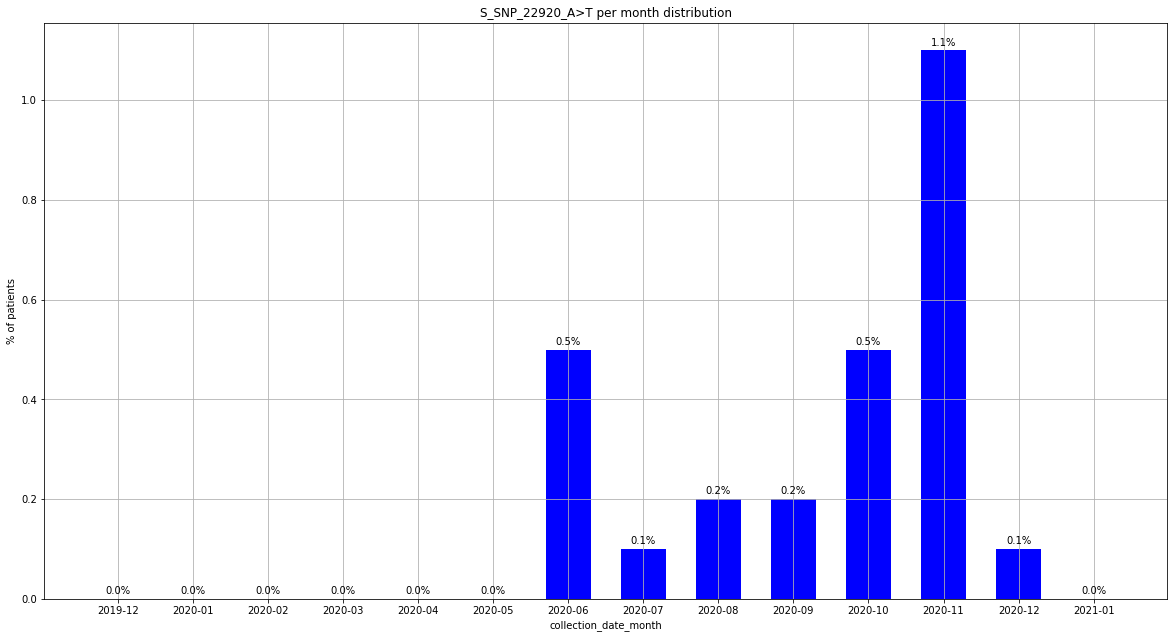

### output_20_278.png

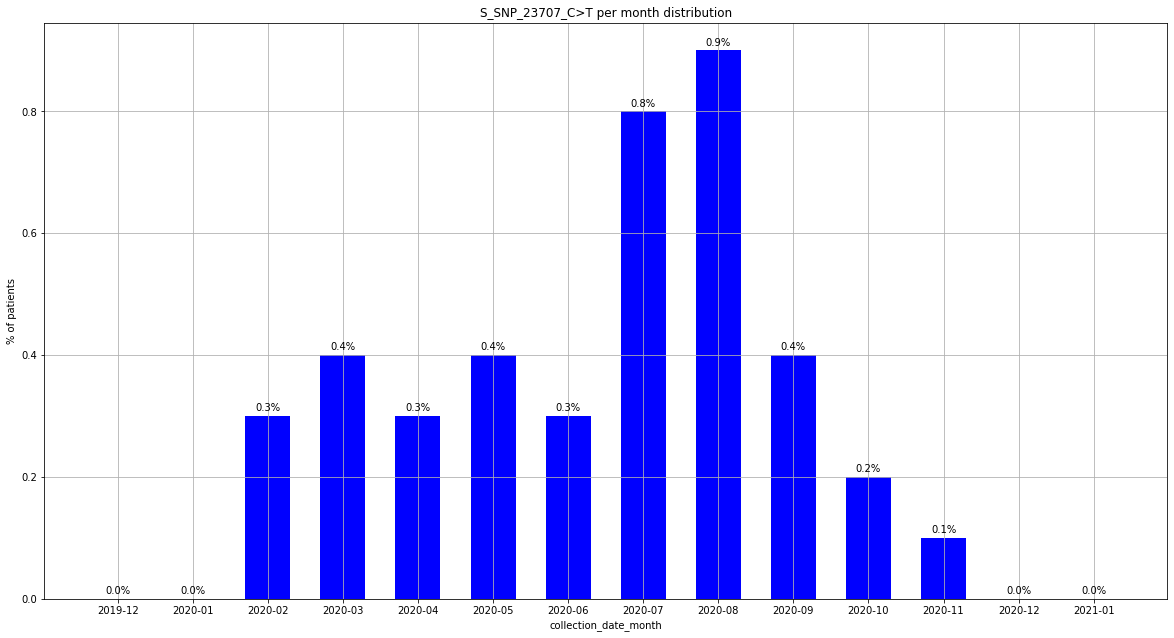

### output_20_290.png

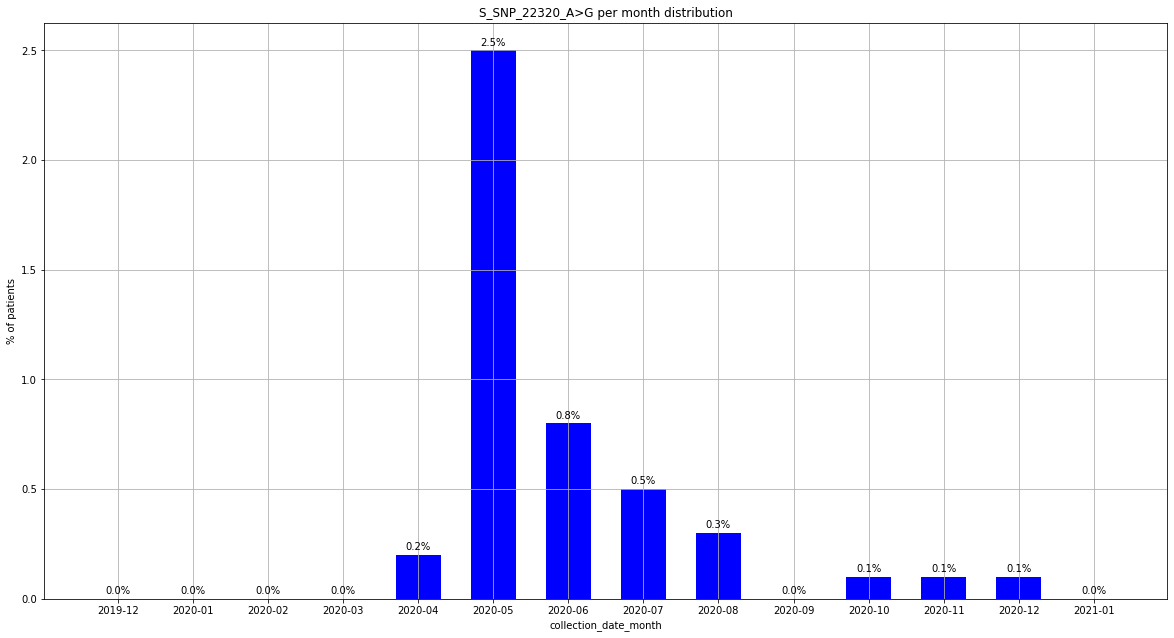

### Supplement 4

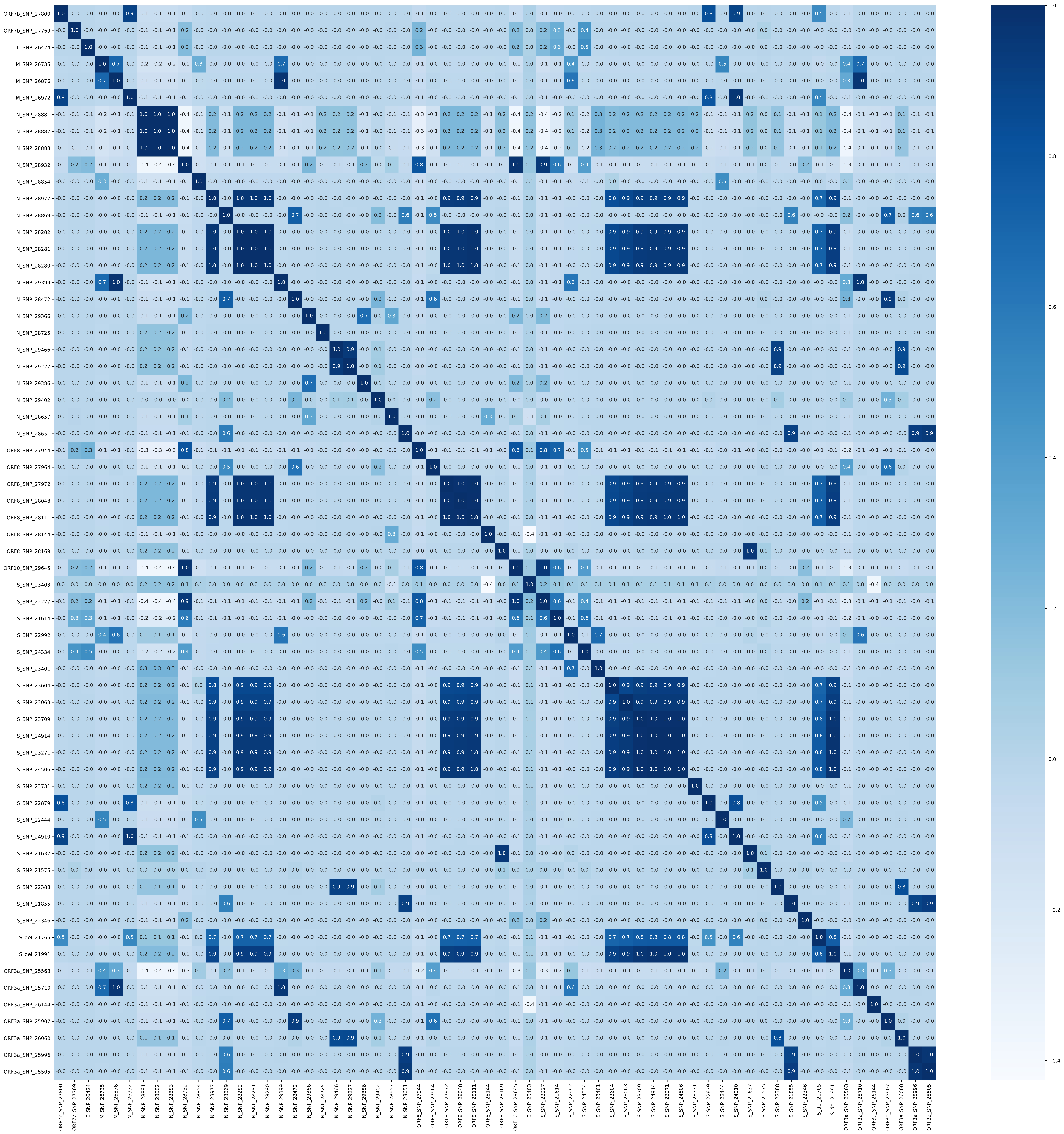
