## Supplementary figures and images for "Analysis of 329,942 SARS-CoV-2 records retrieved from GISAID database"

### loading_stats_by_collection_date.png

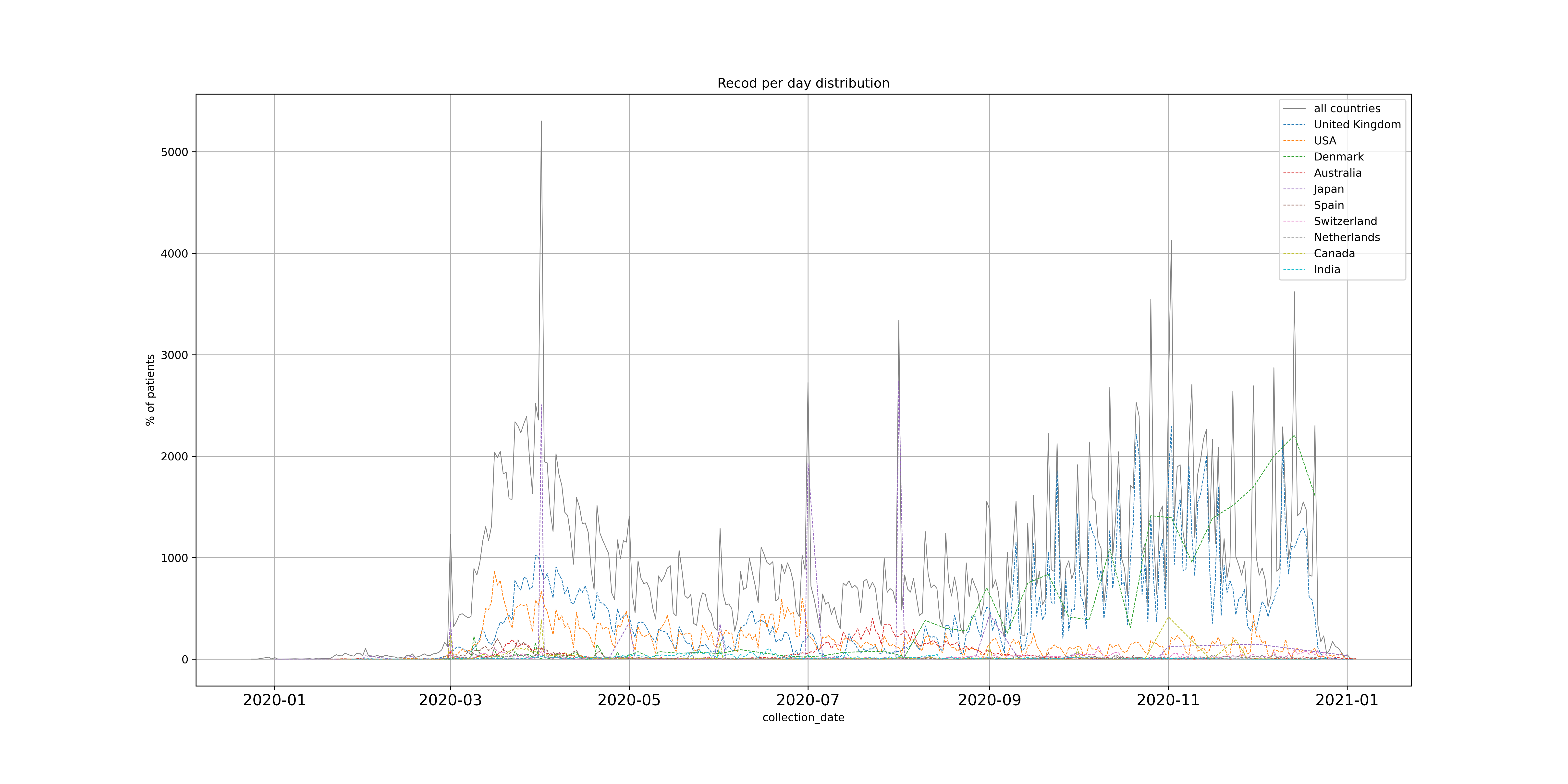

### loading_stats_by_days_ago.png

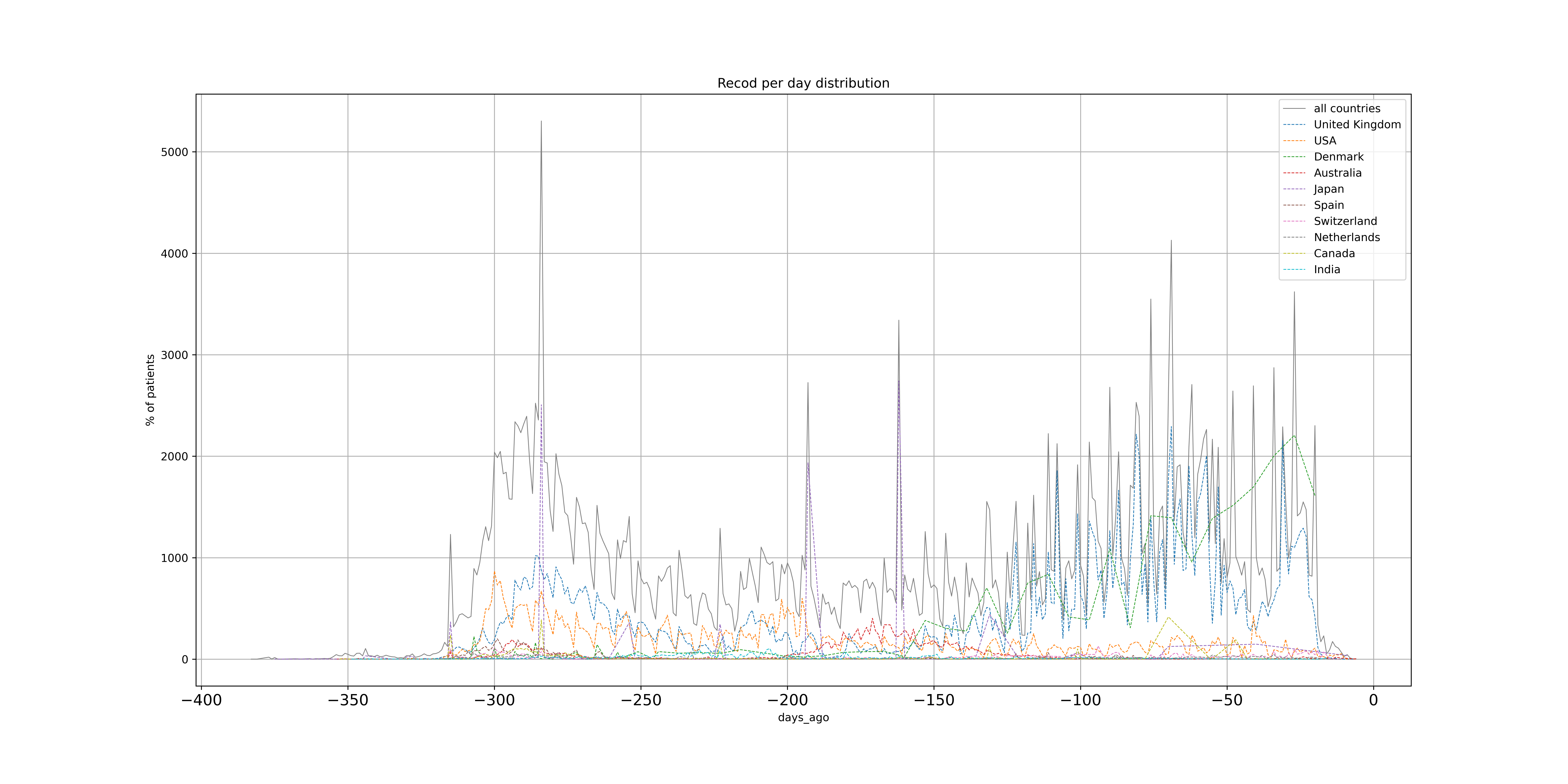

### output_19_1.png

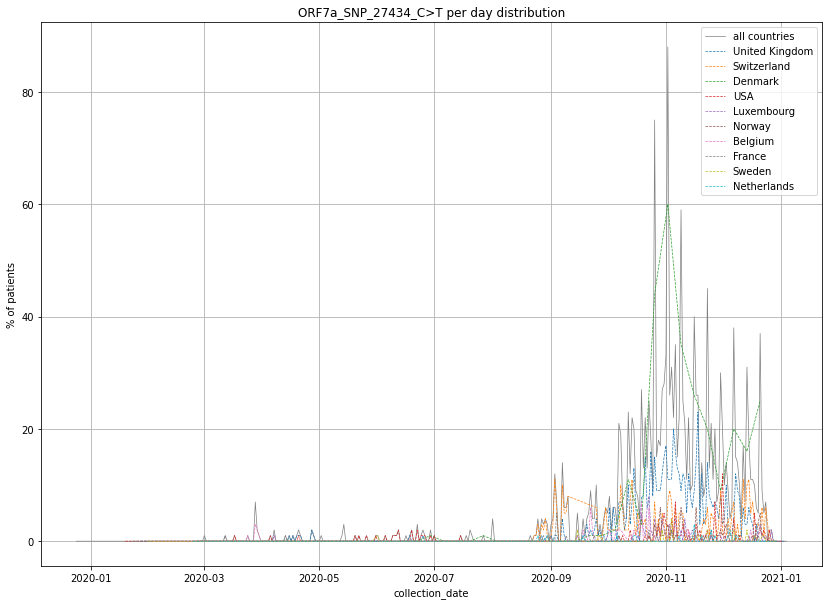

### output_19_1.png

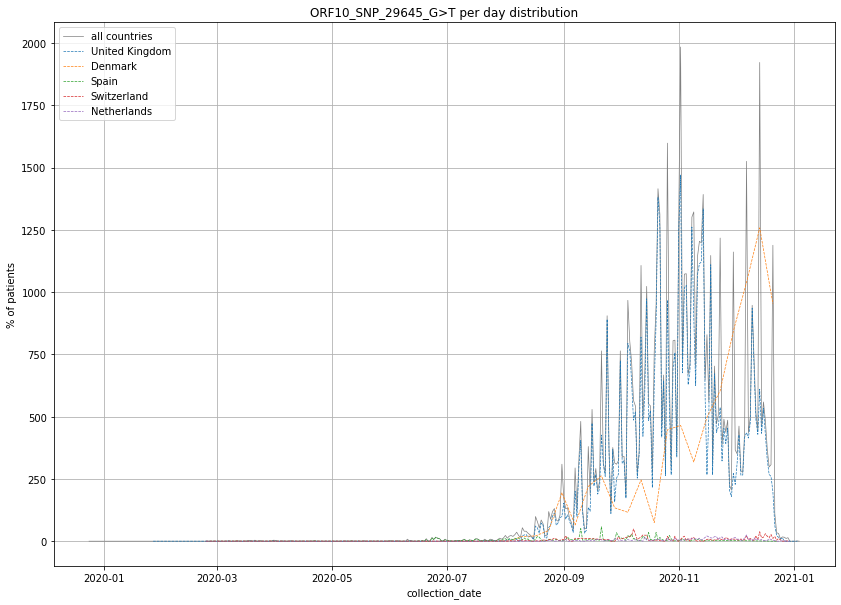

### output_19_1.png

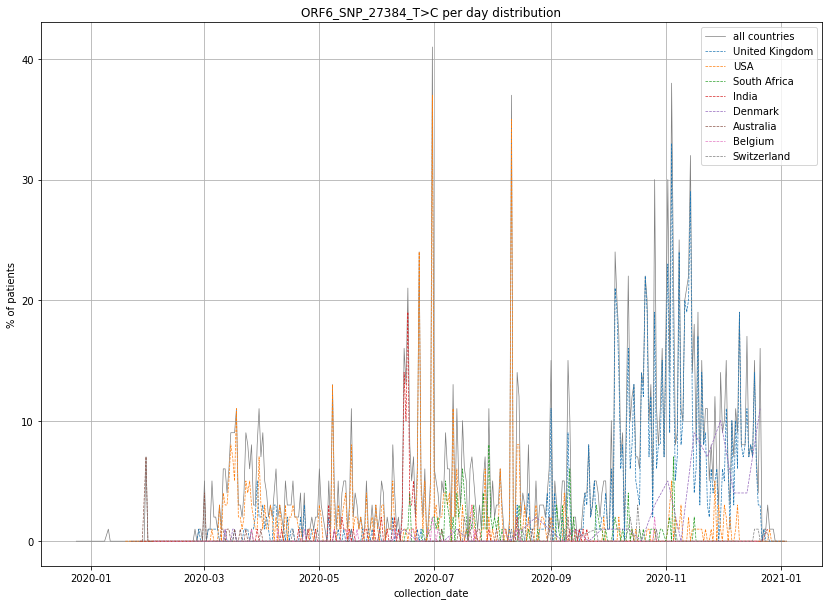

### output_19_1.png

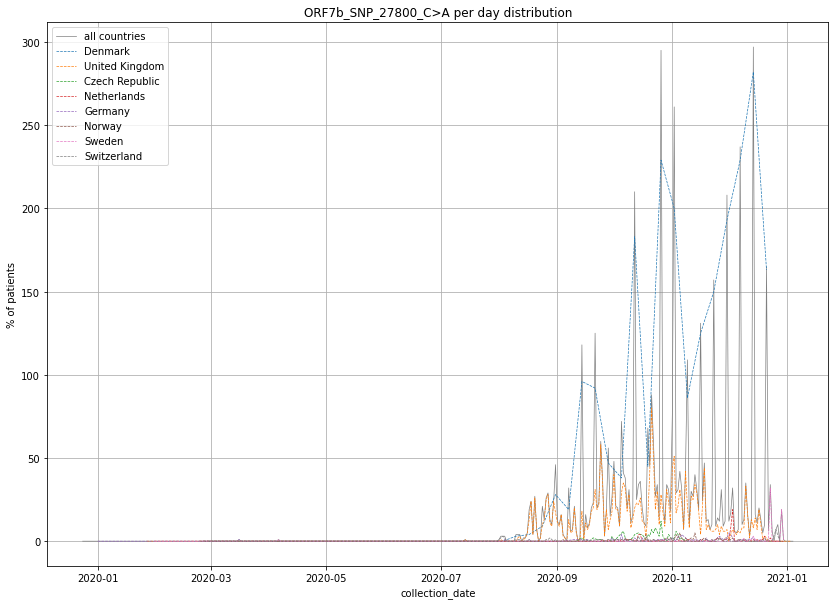

### output_19_1.png

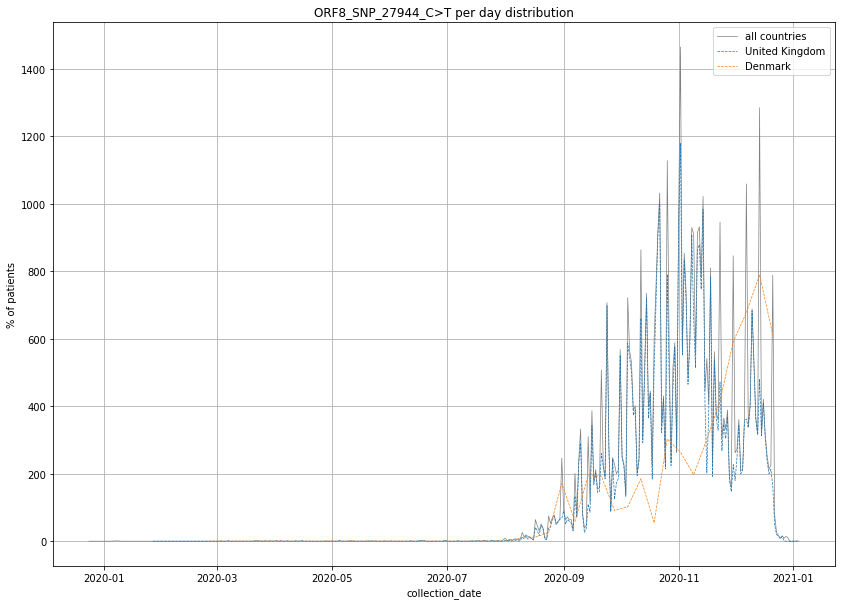

### output_19_1.png

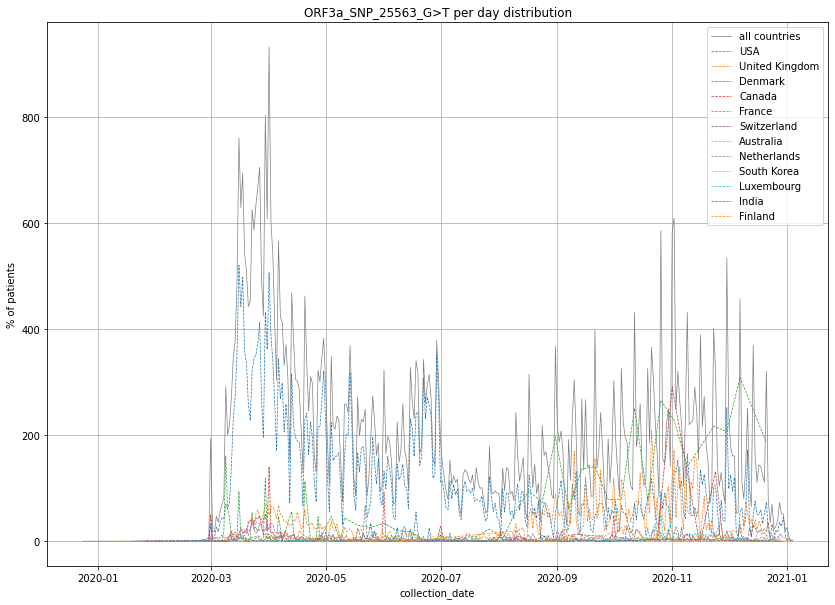

### output_19_2.png

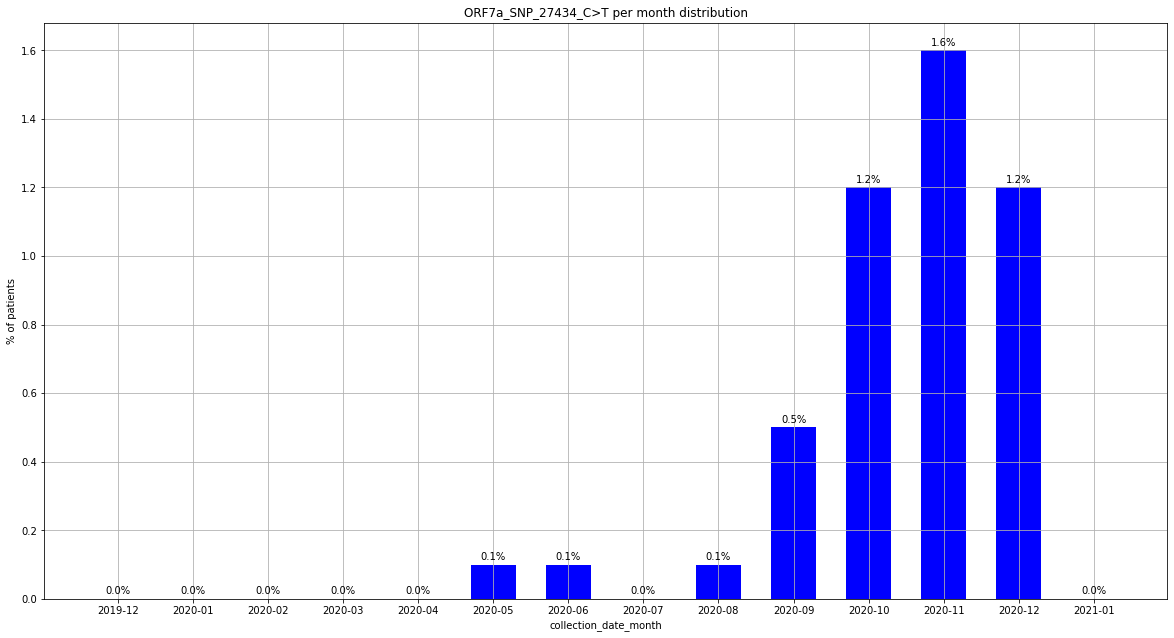

### output_19_2.png

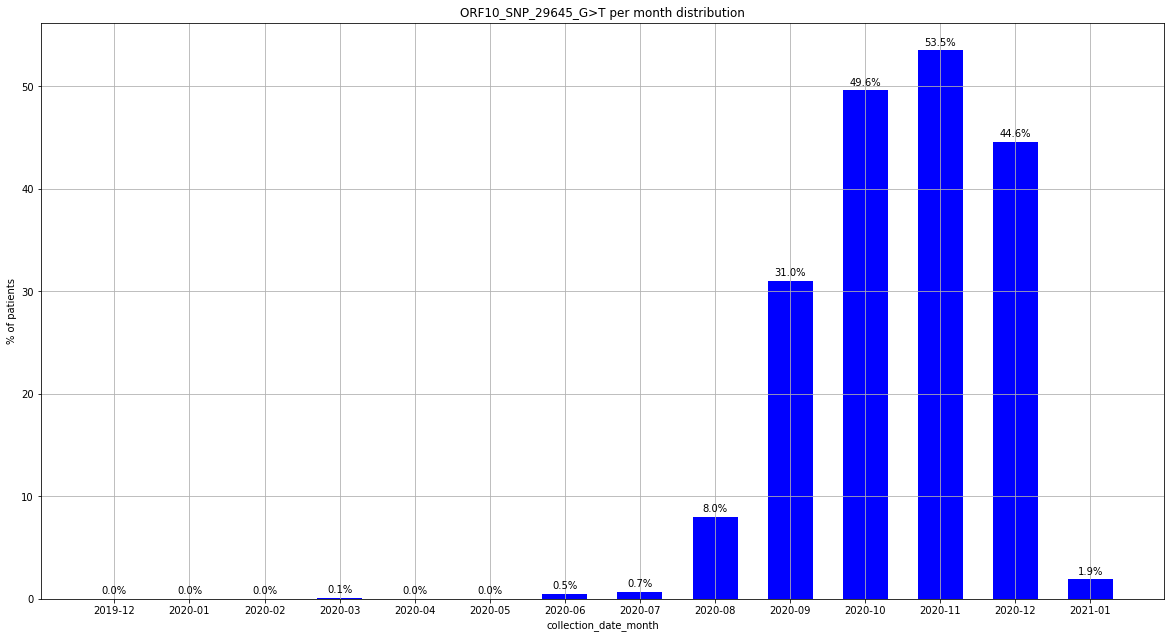

### output_19_2.png

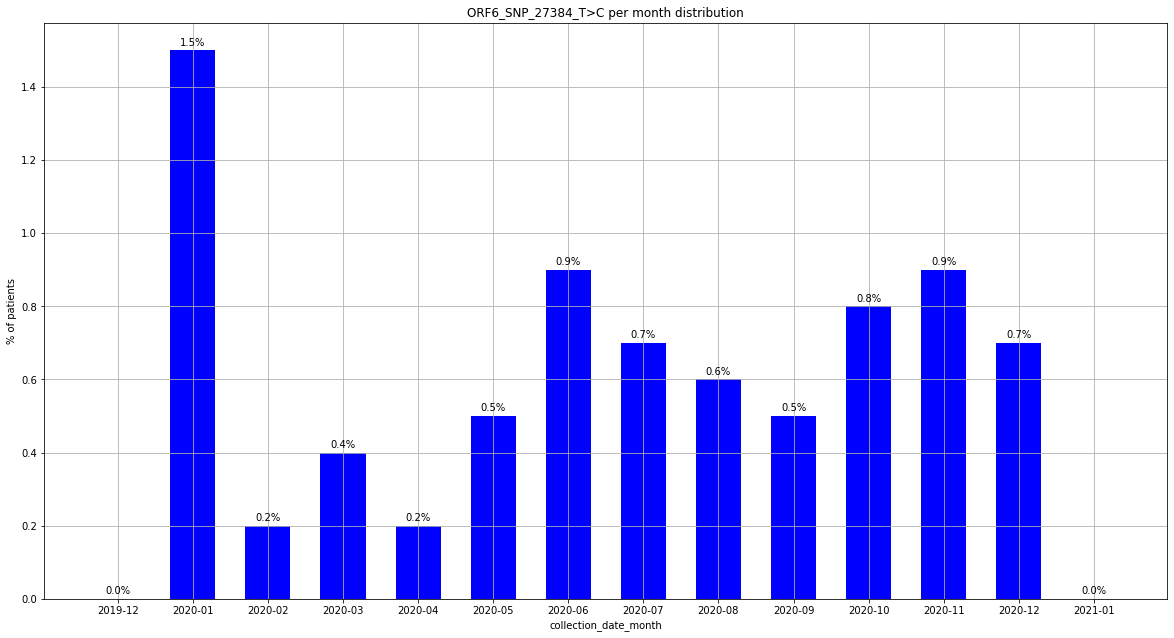

### output_19_2.png

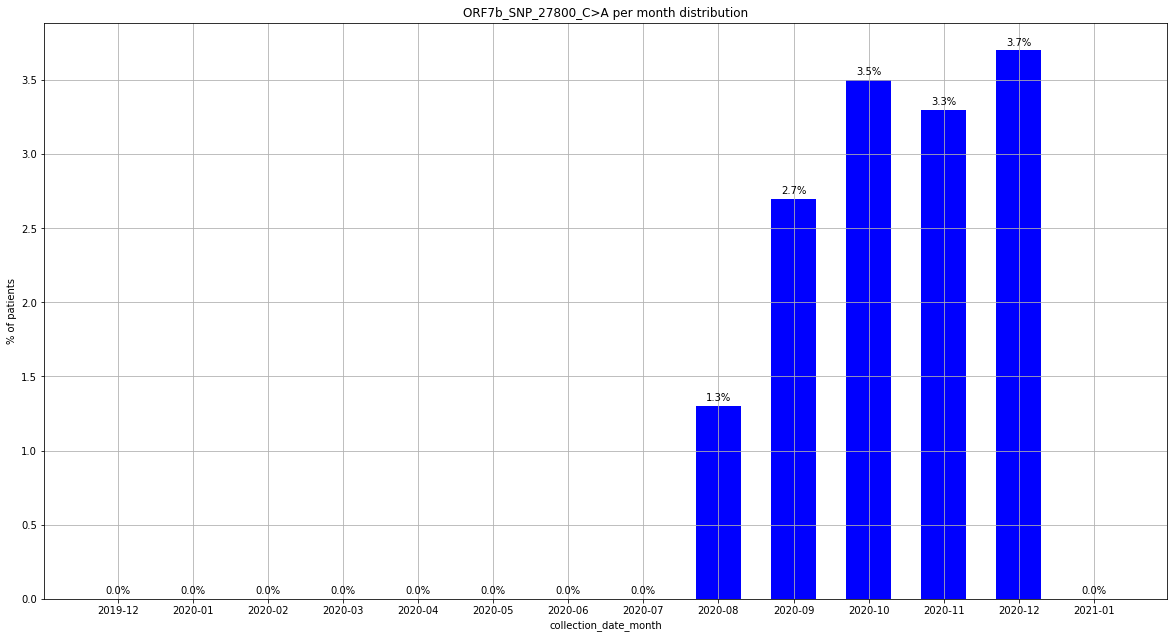

### output_19_2.png

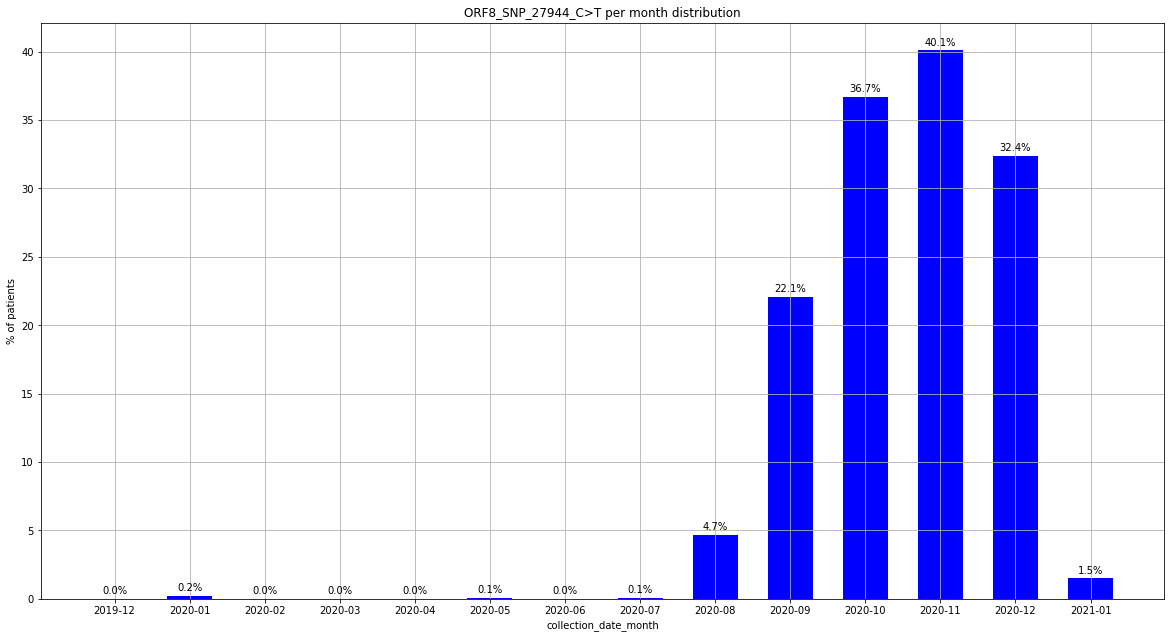

### output_19_2.png

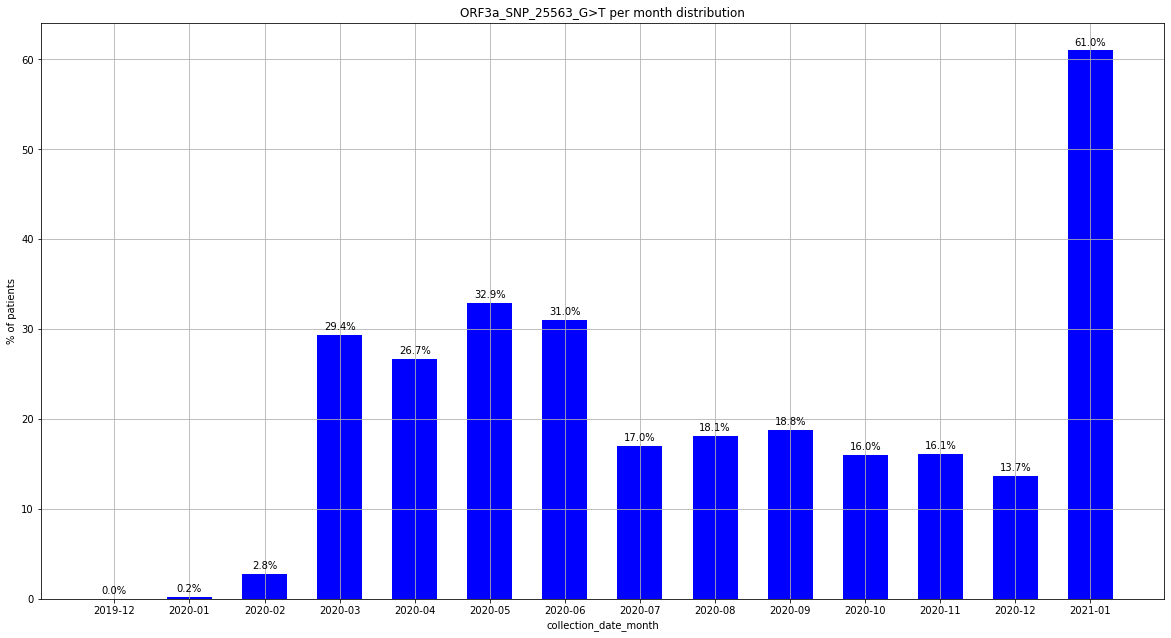

### output_19_7.png

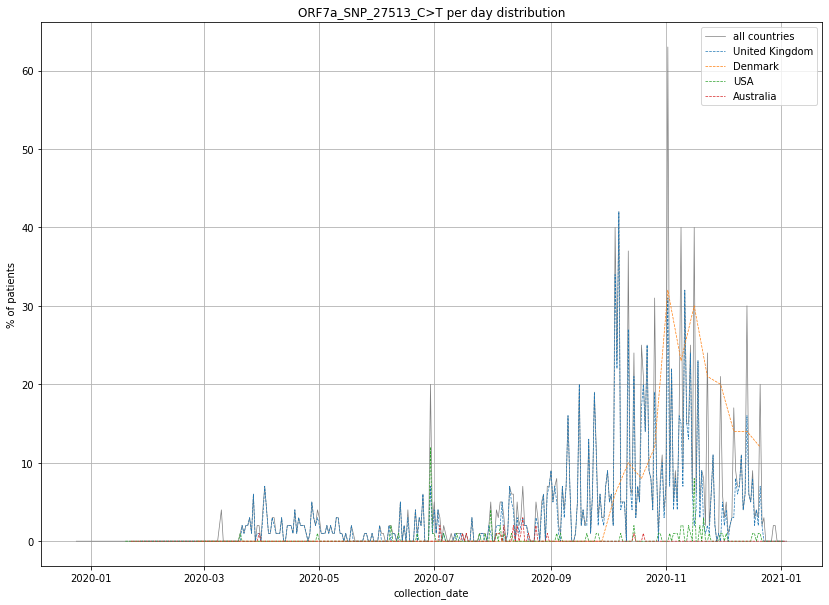

### output_19_7.png

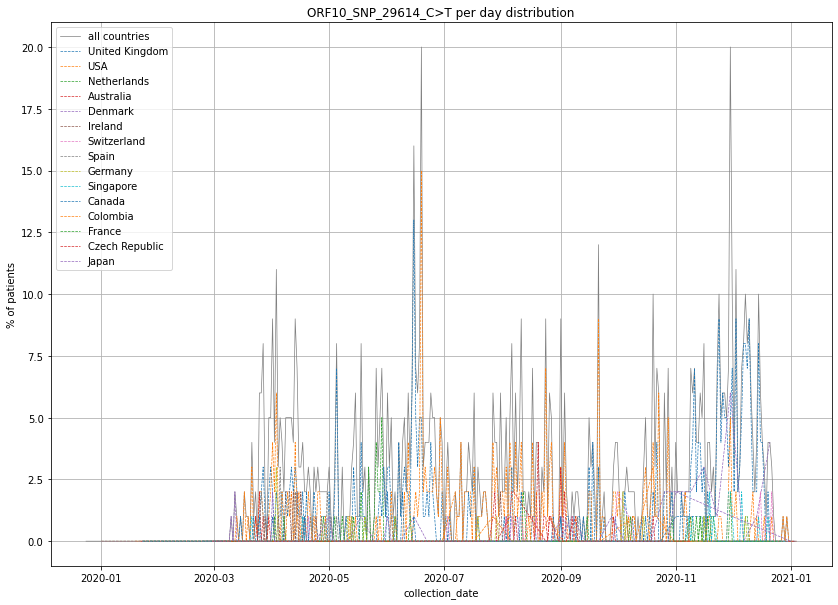

### output_19_7.png

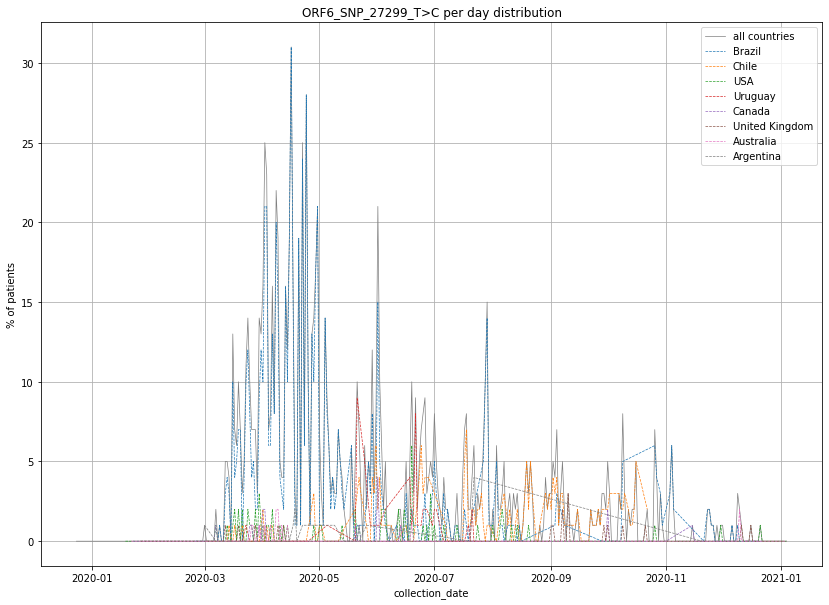

### output_19_7.png

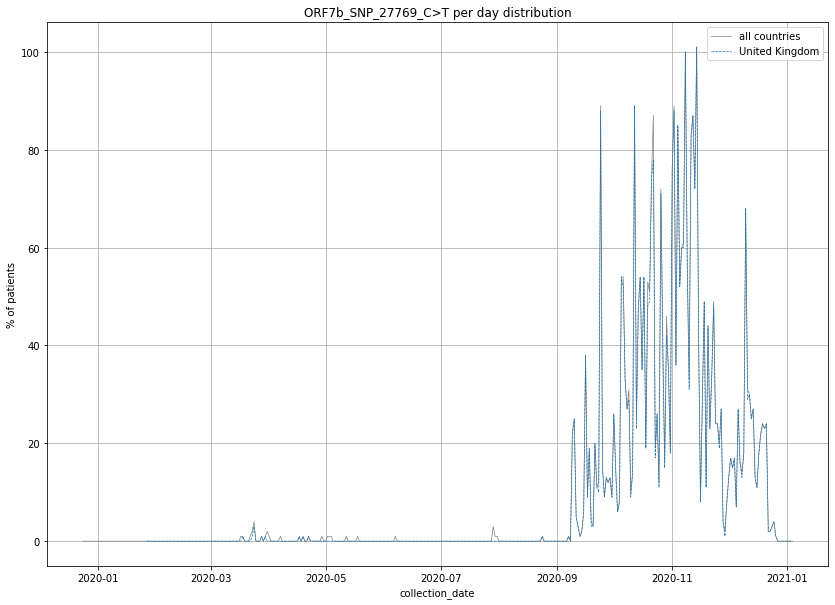

### output_19_7.png

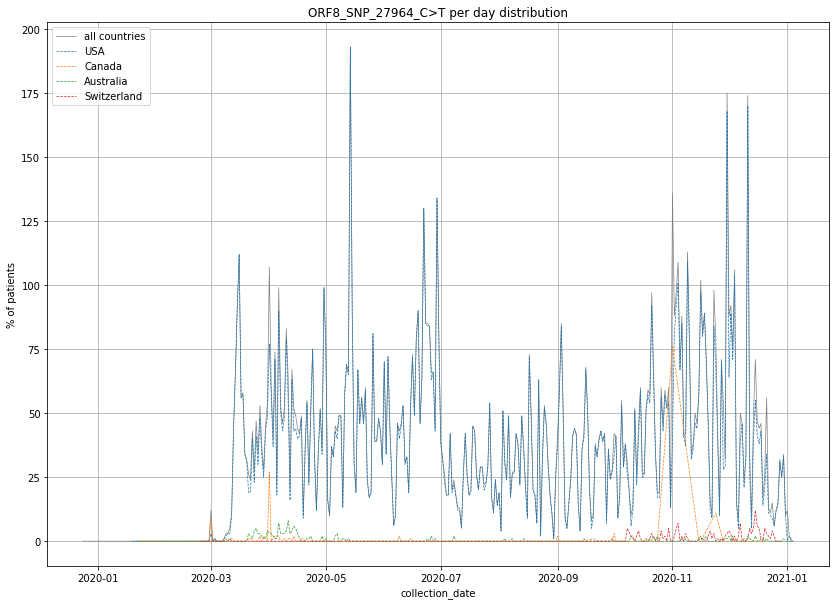

### output_19_8.png

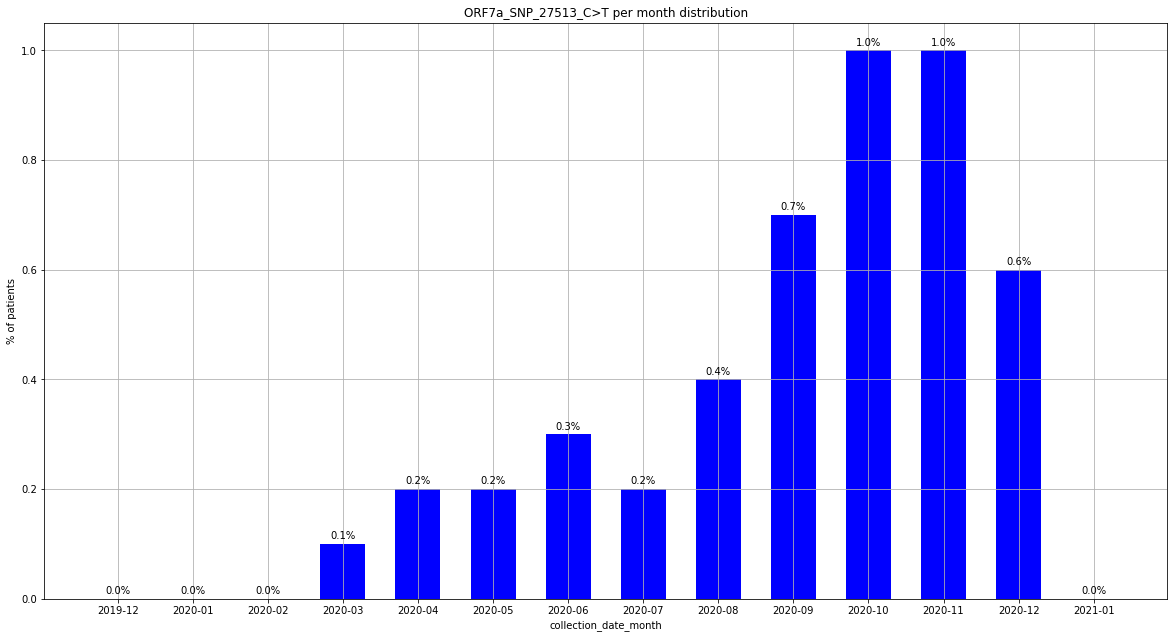

### output_19_8.png

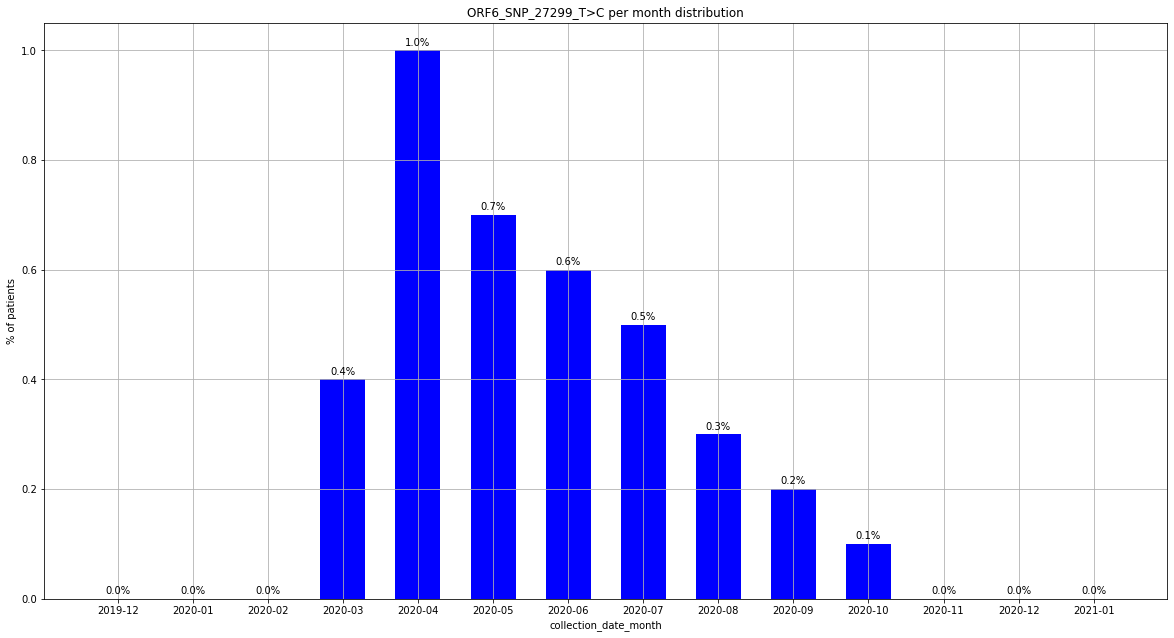

### output_19_8.png

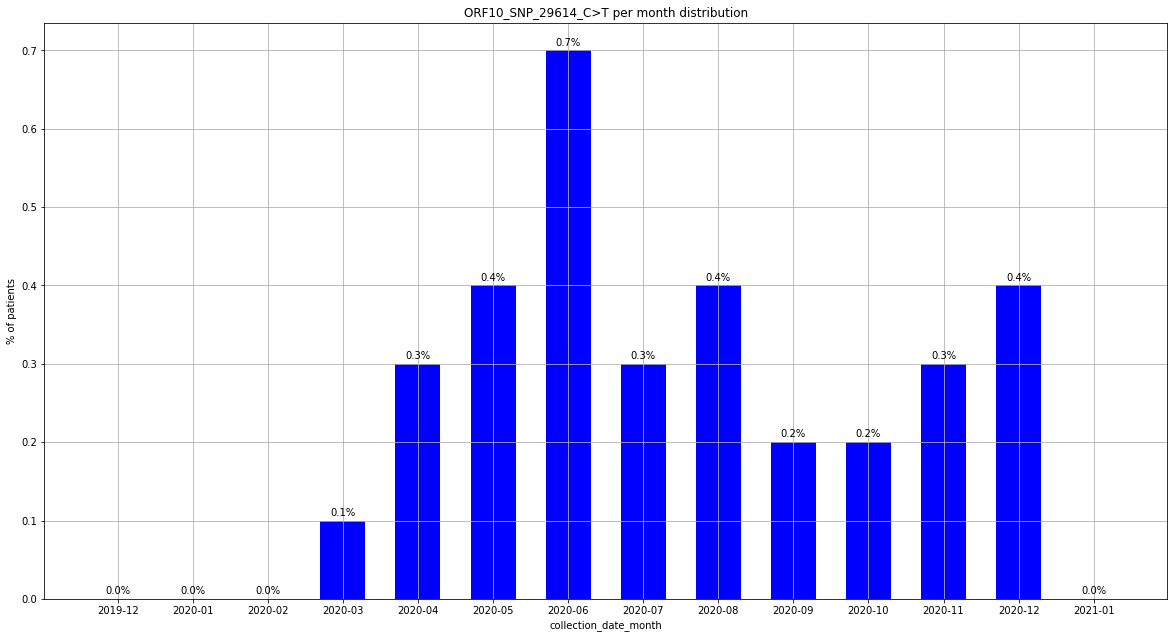

### output_19_8.png

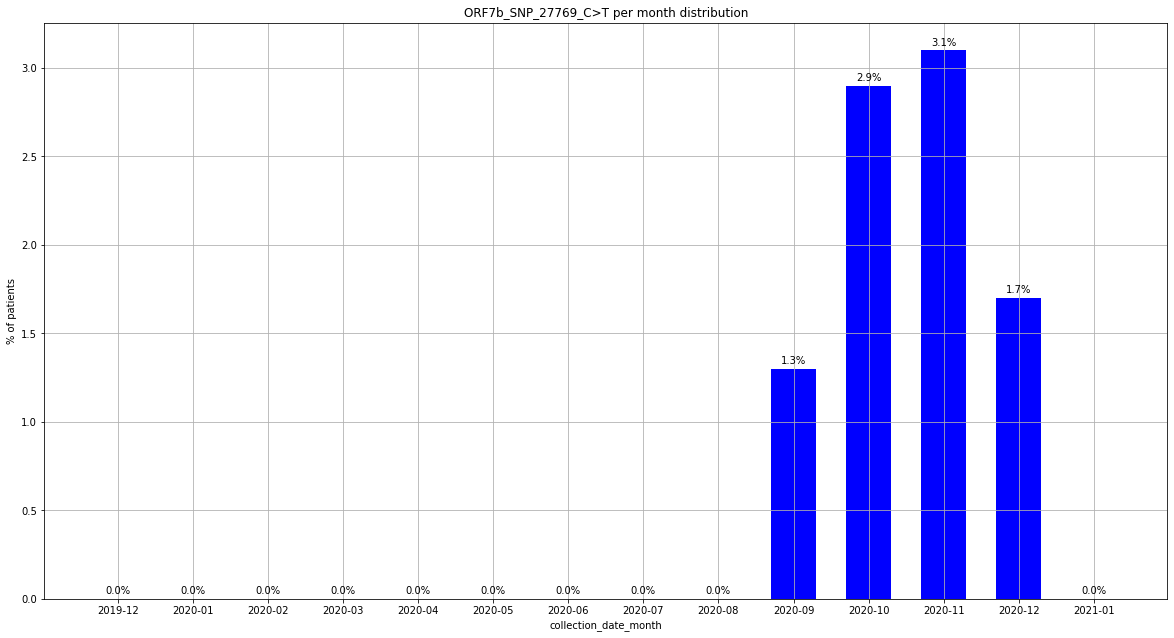

### output_19_8.png

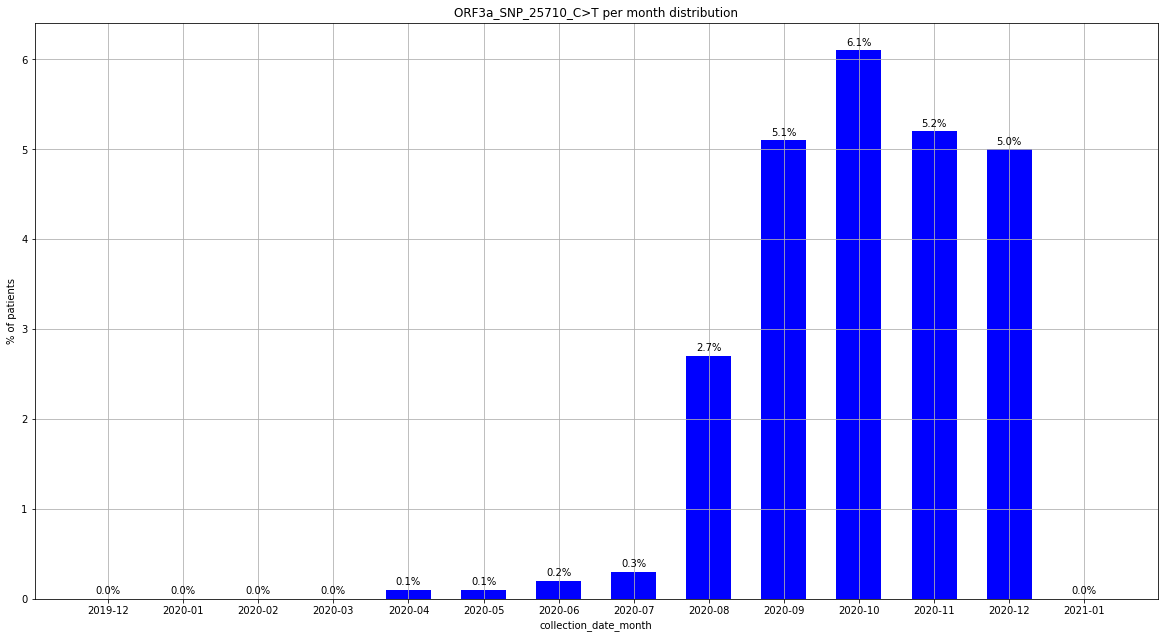

### output_19_8.png

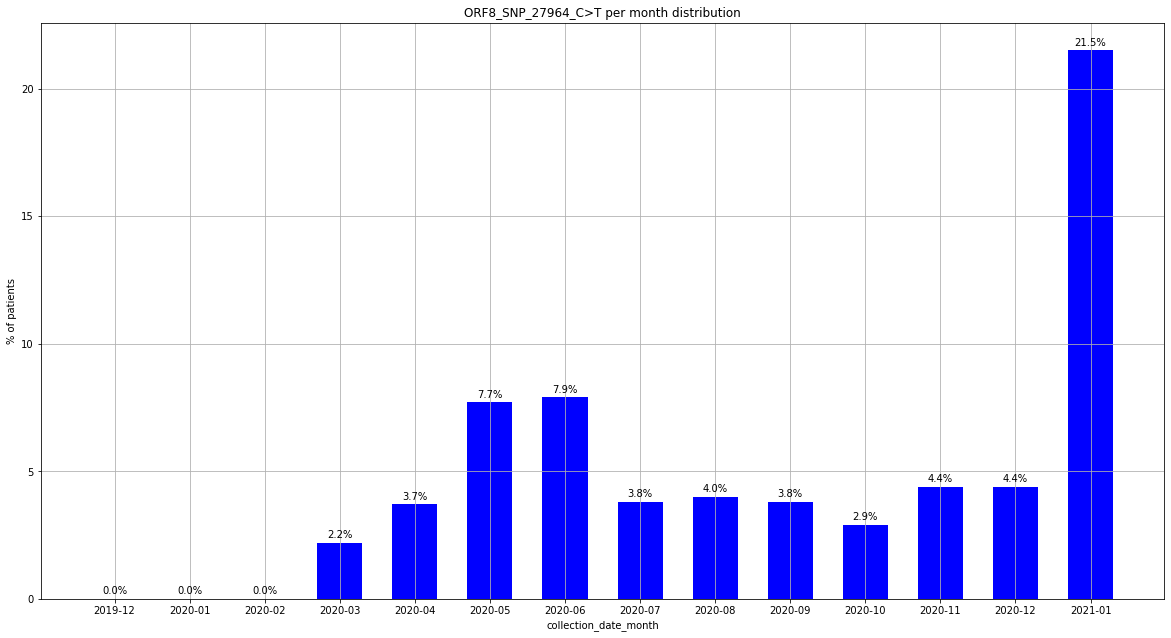

### output_19_13.png

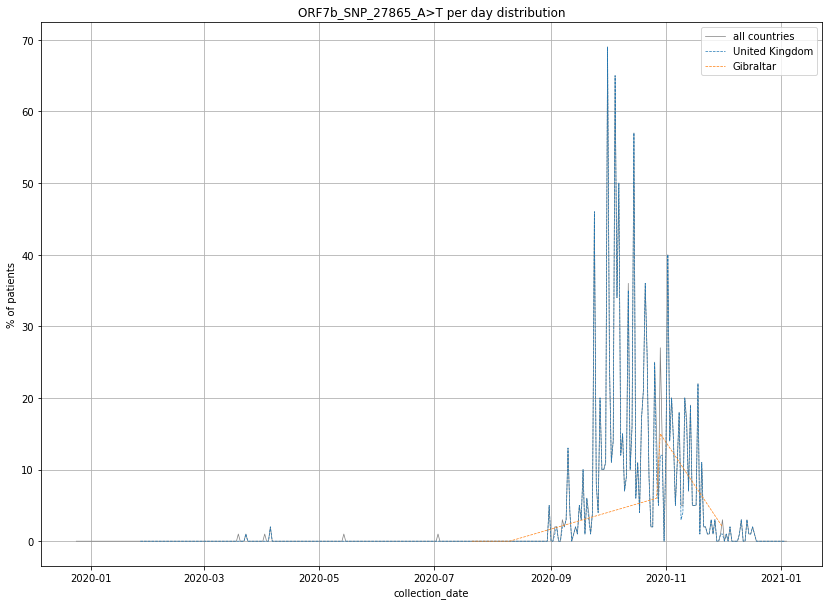

### output_19_13.png

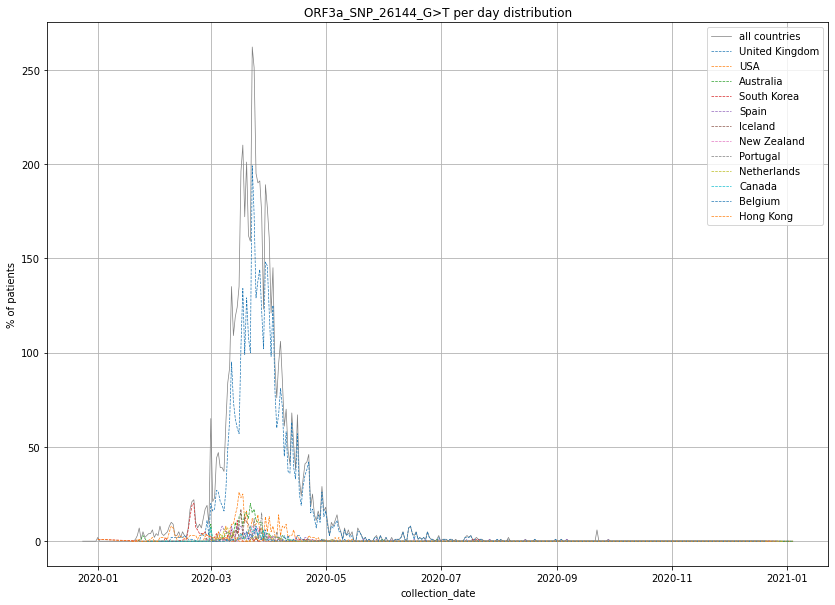

### output_19_13.png

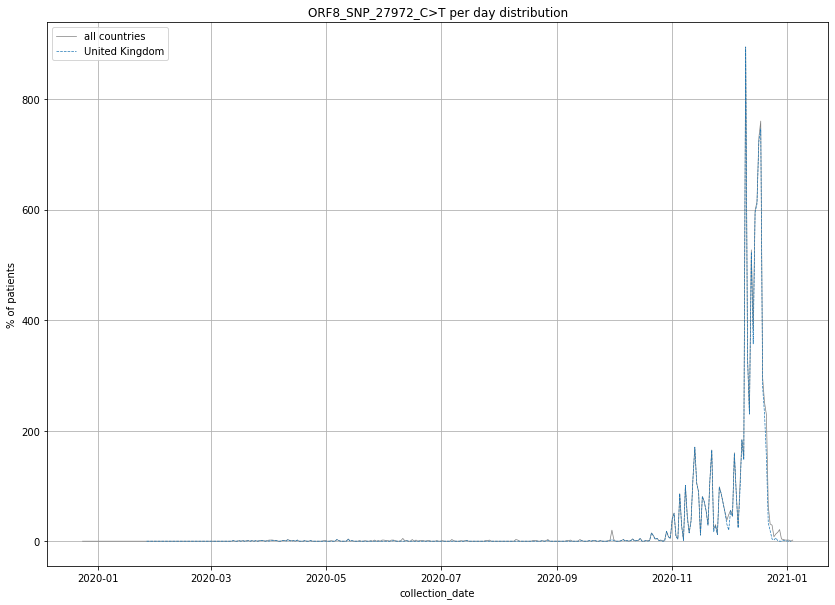

### output_19_14.png

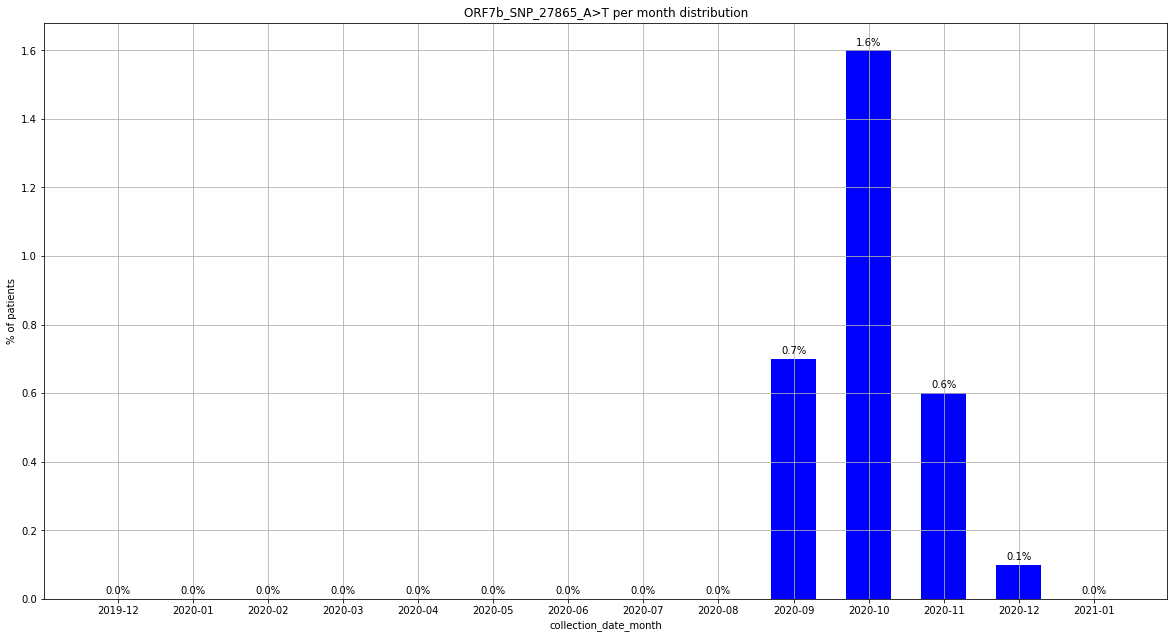

### output_19_14.png

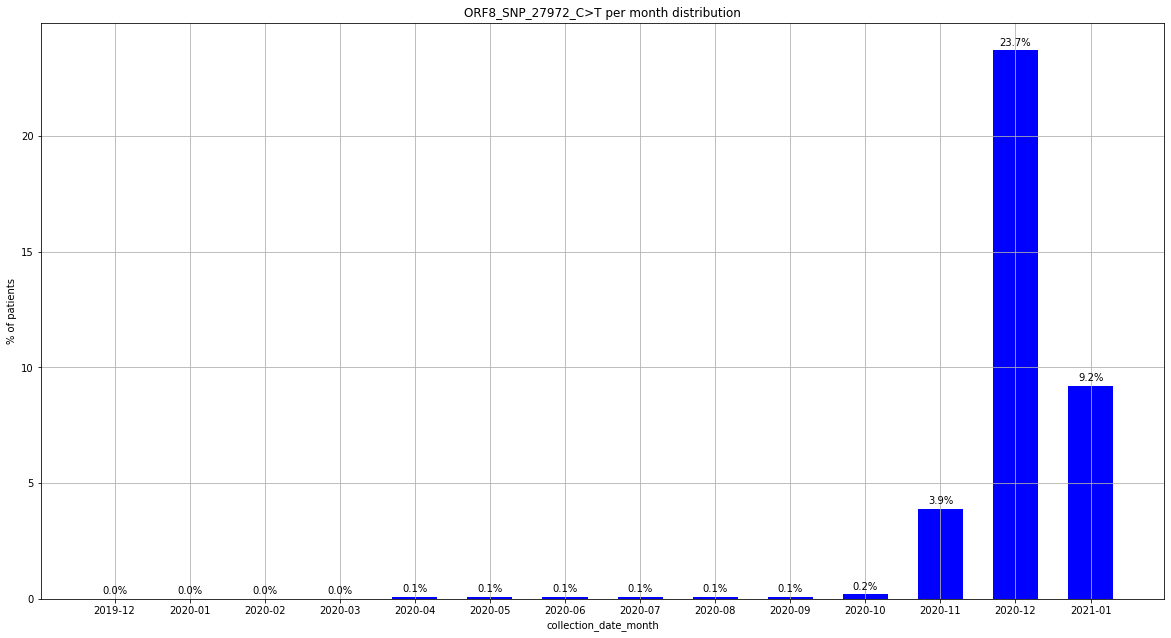

### output_19_14.png

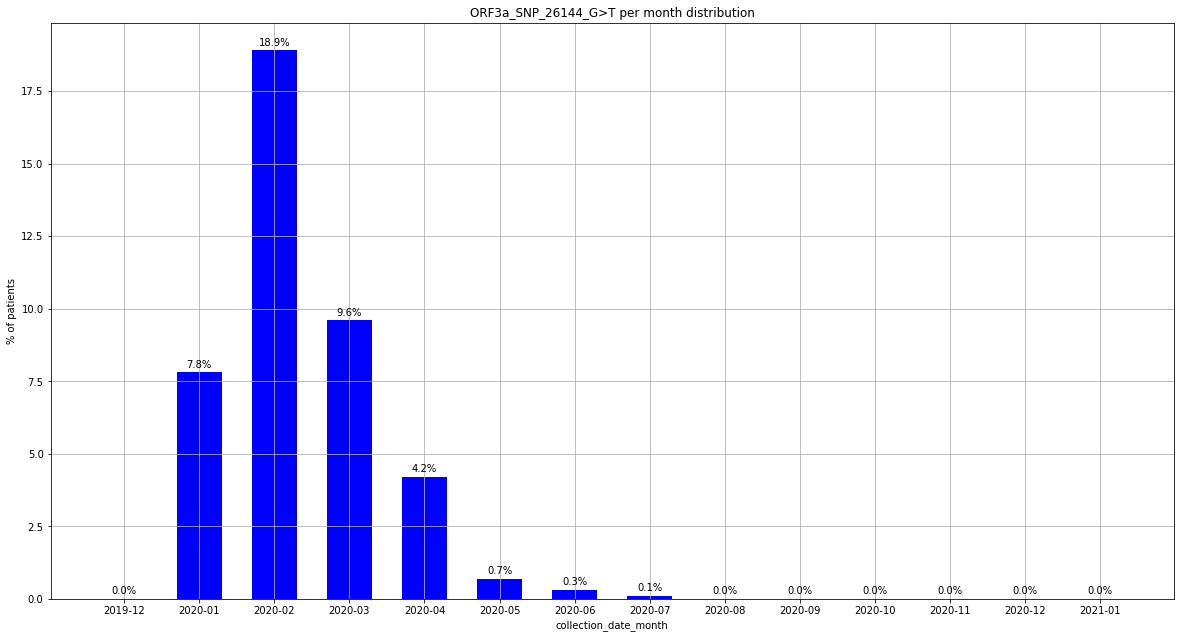

### output_19_19.png

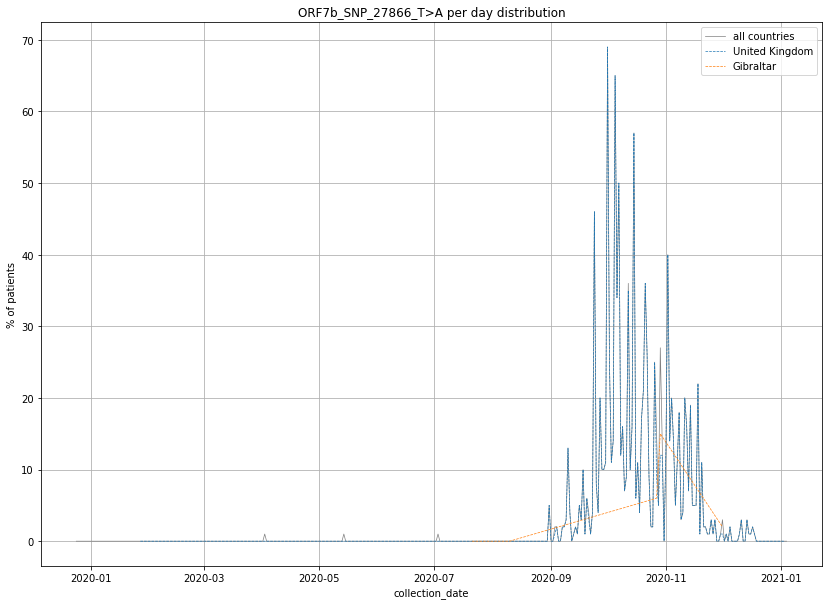

### output_19_19.png

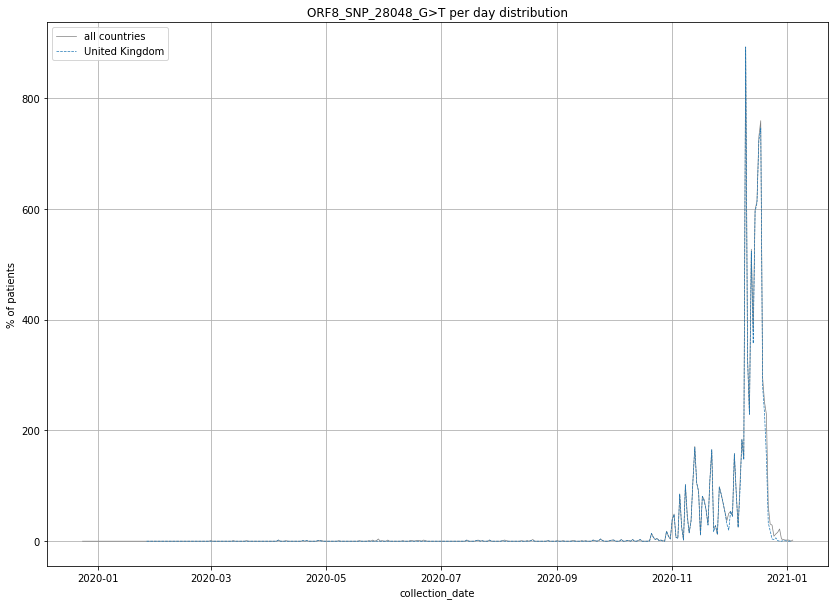

### output_19_20.png

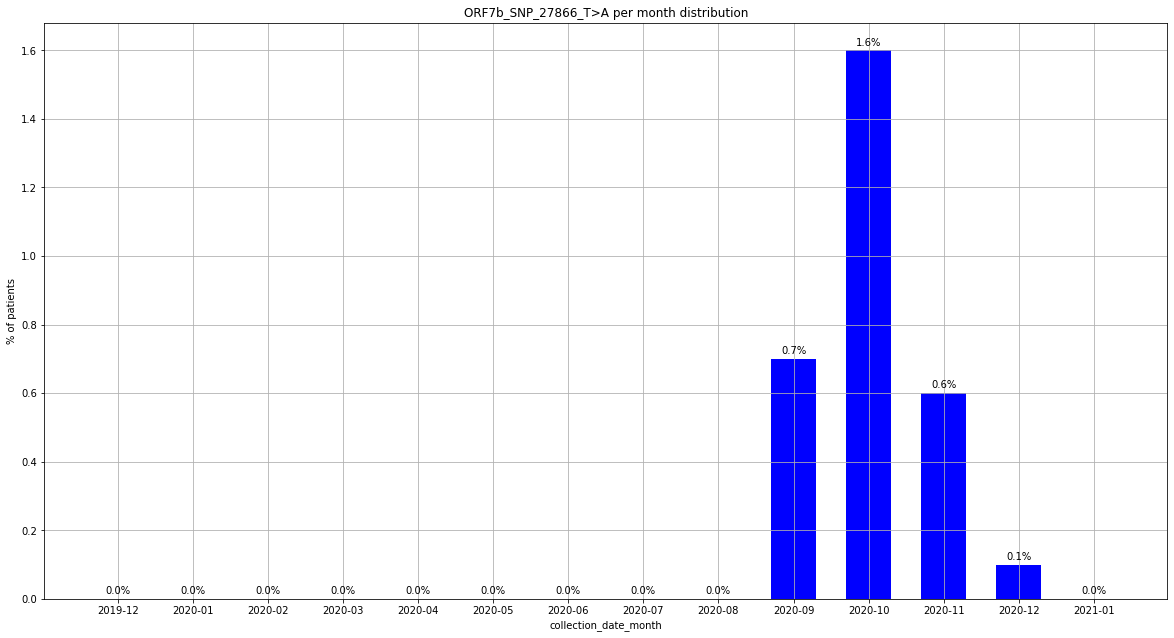

### output_19_20.png

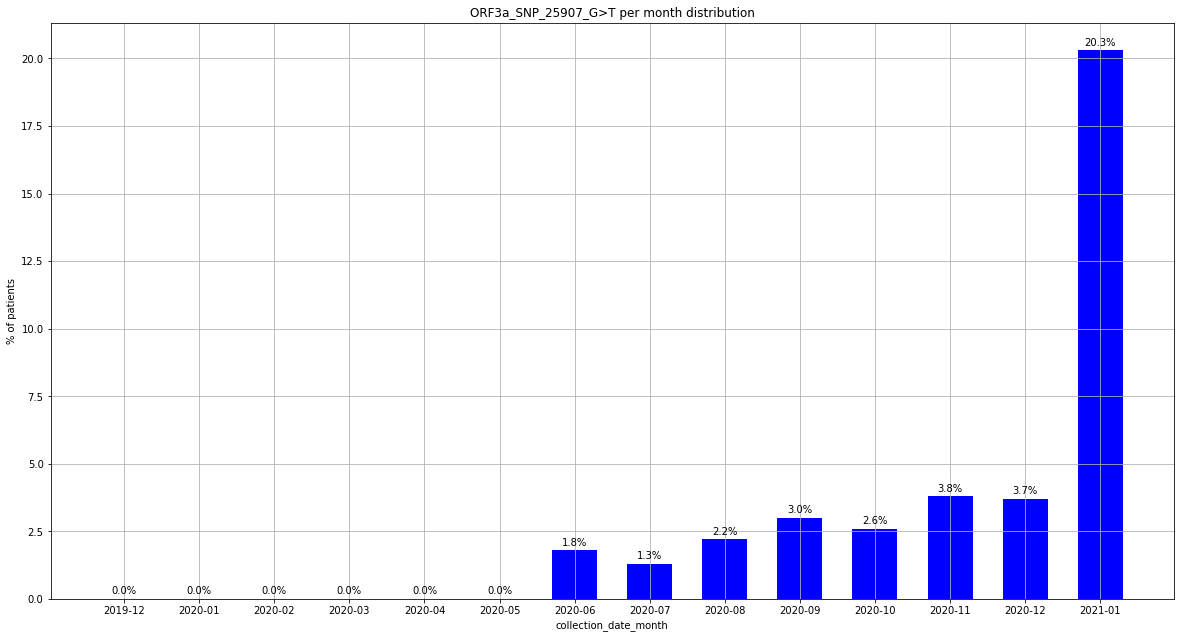

### output_19_20.png

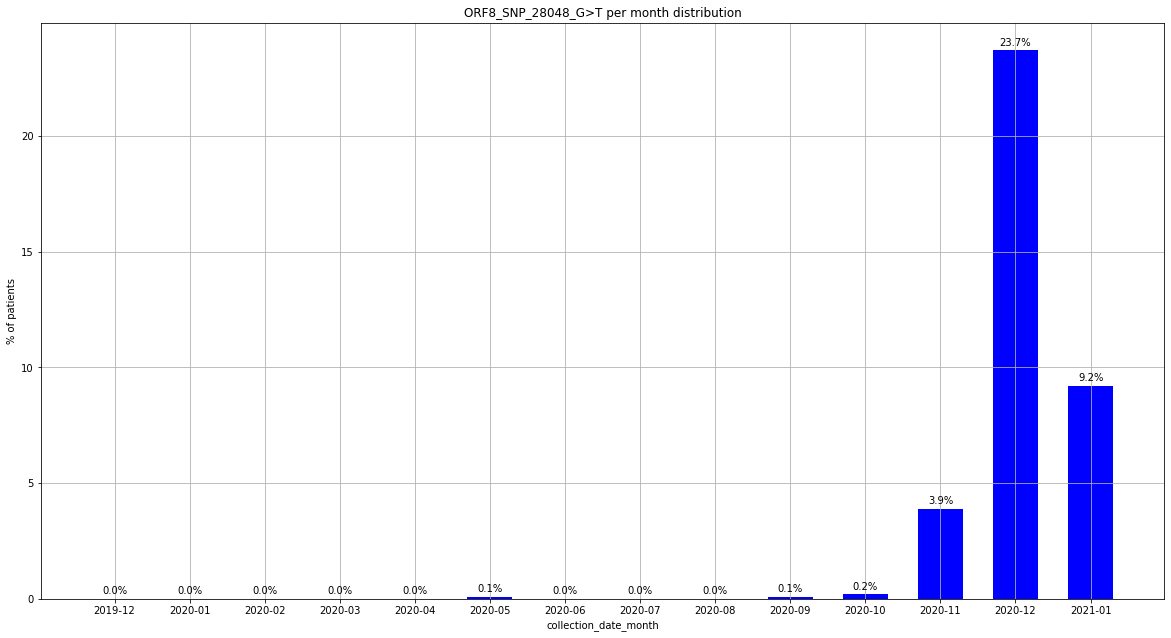

### output_19_25.png

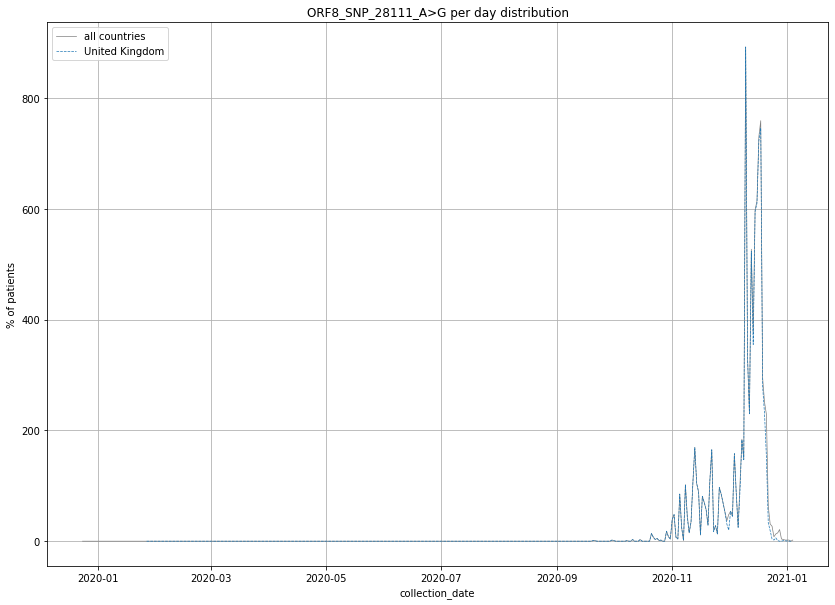

### output_19_25.png

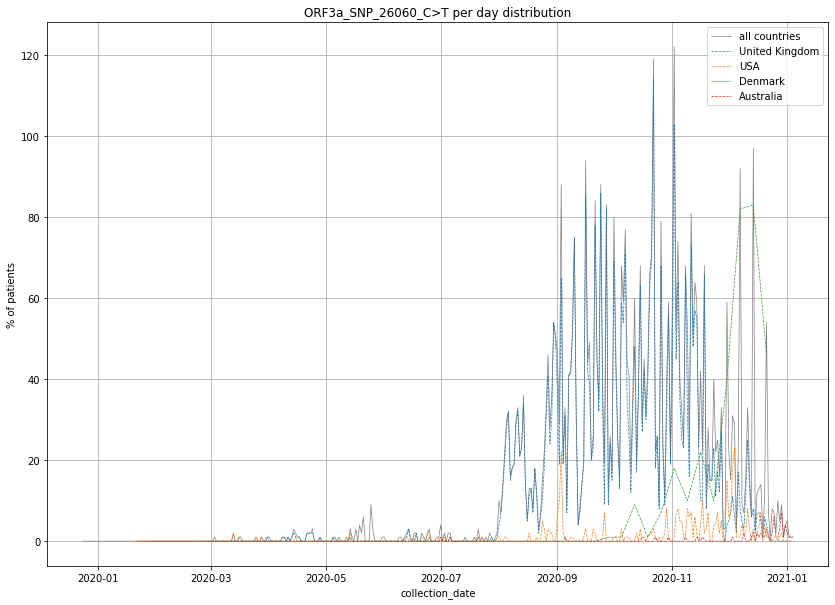

### output_19_26.png

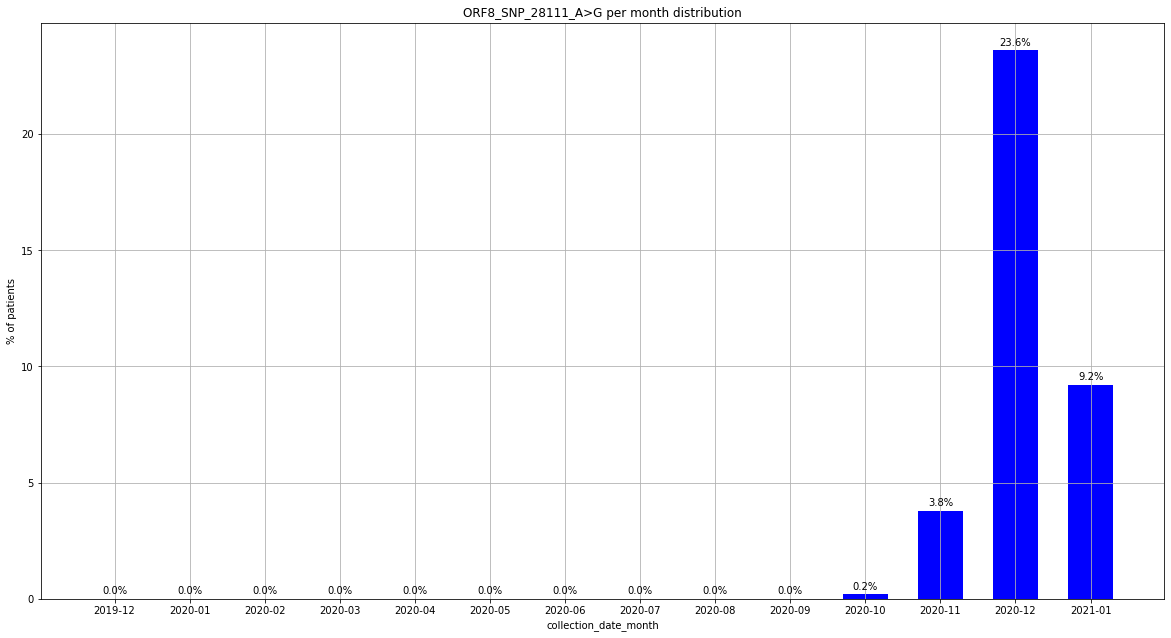

### output_19_26.png

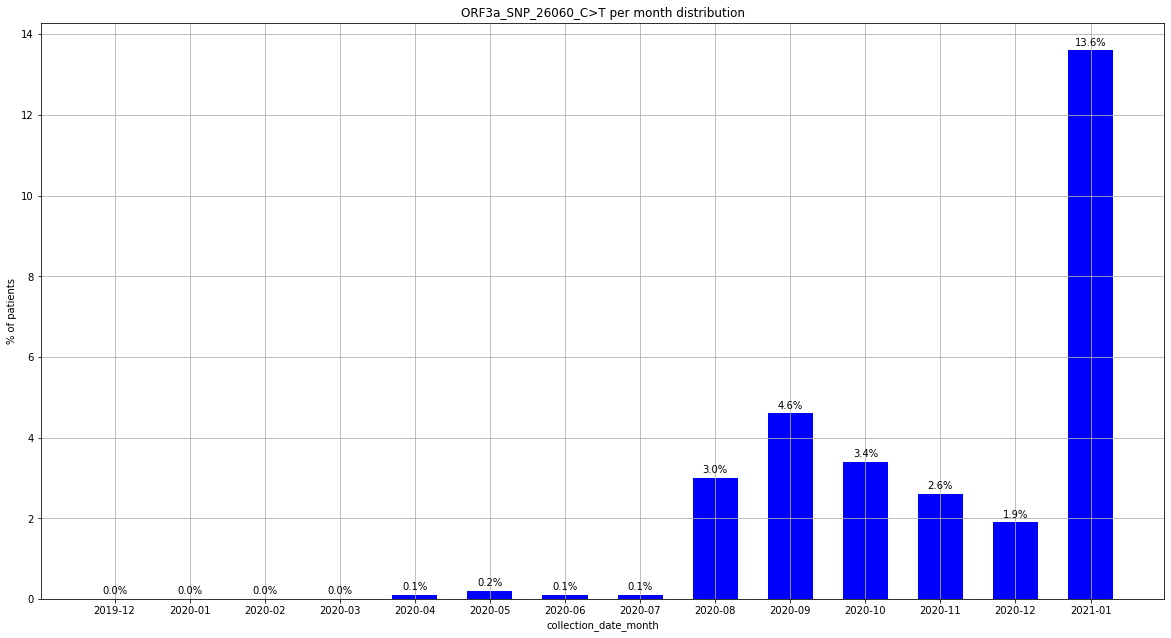

### output_19_31.png

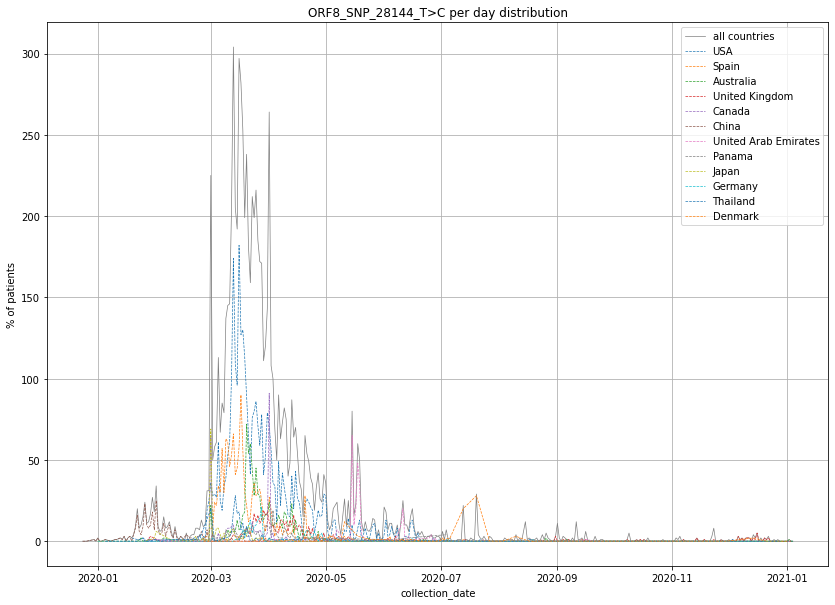

### output_19_31.png

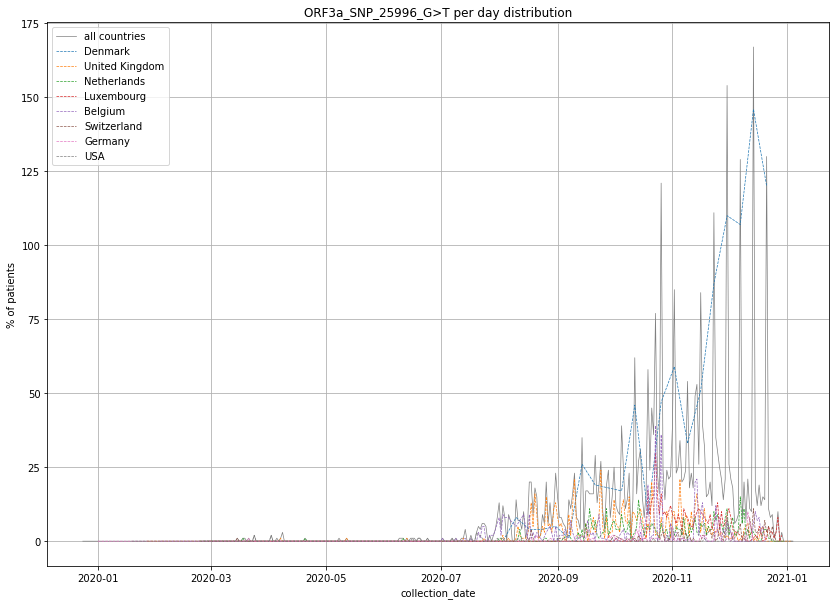

### output_19_32.png

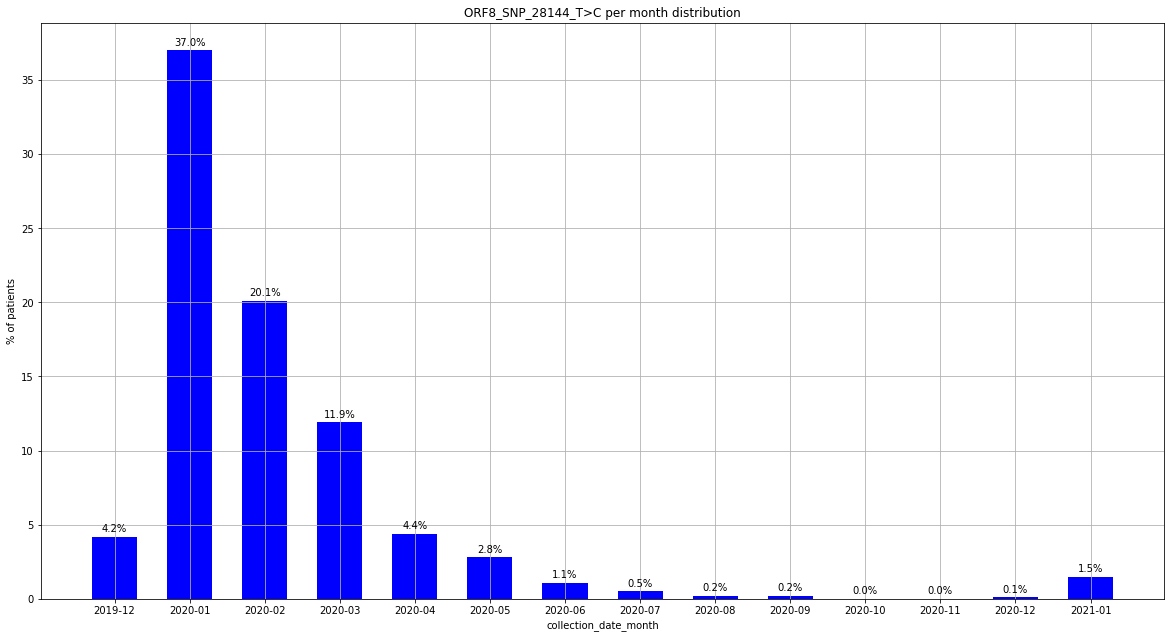

### output_19_37.png

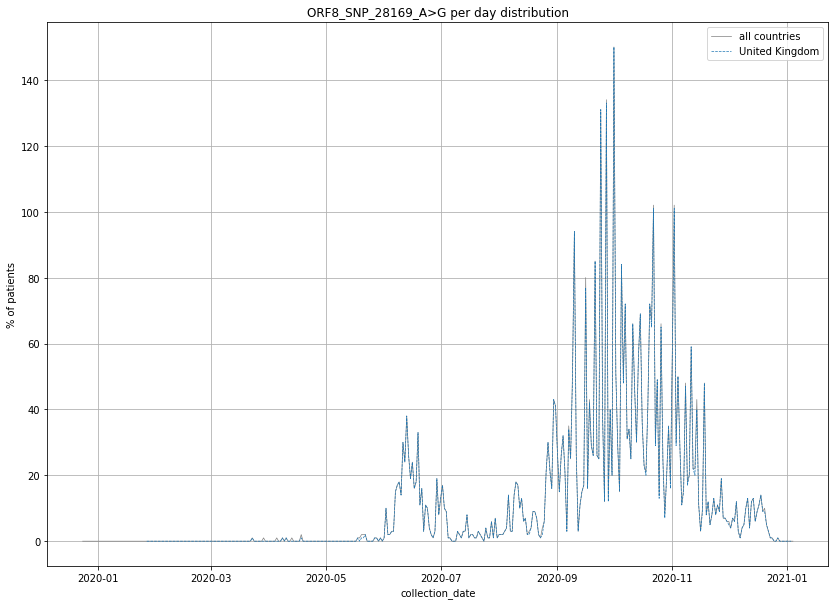

### output_19_37.png

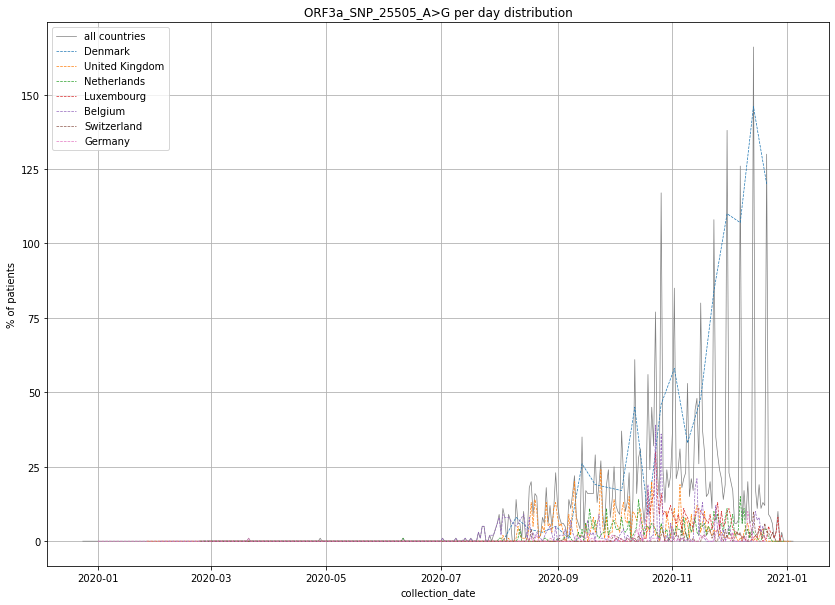

### output_19_38.png

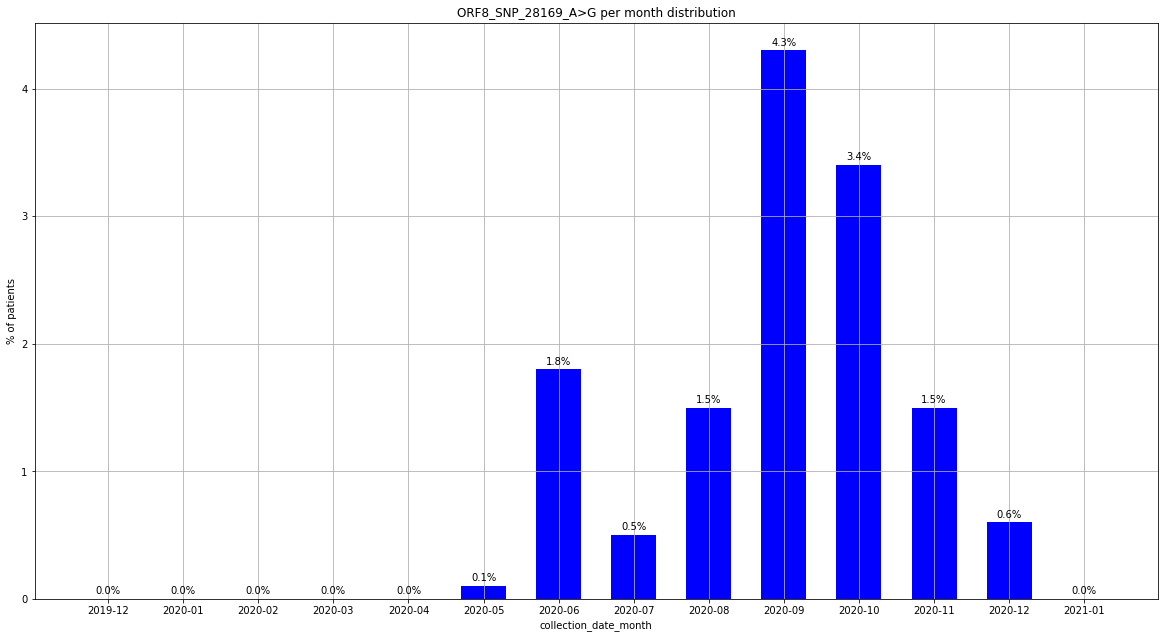

### output_19_43.png

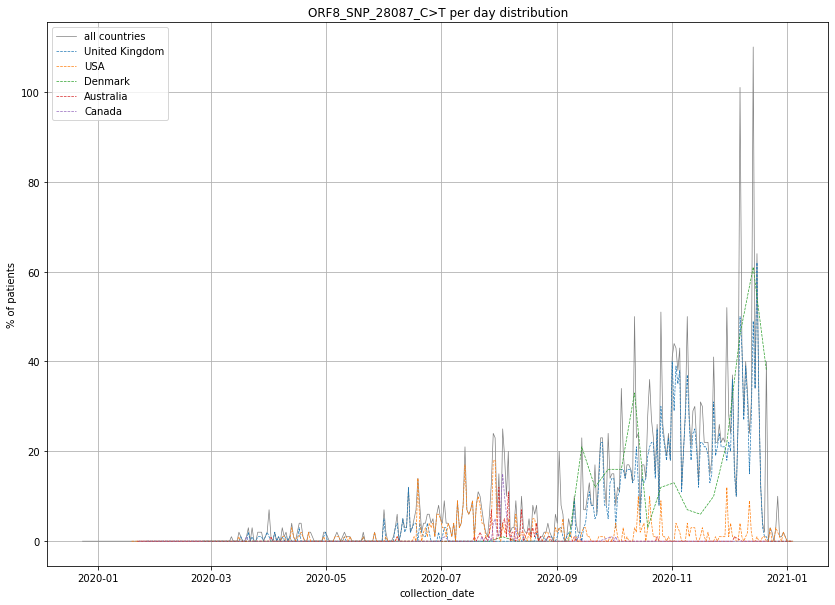

### output_19_44.png

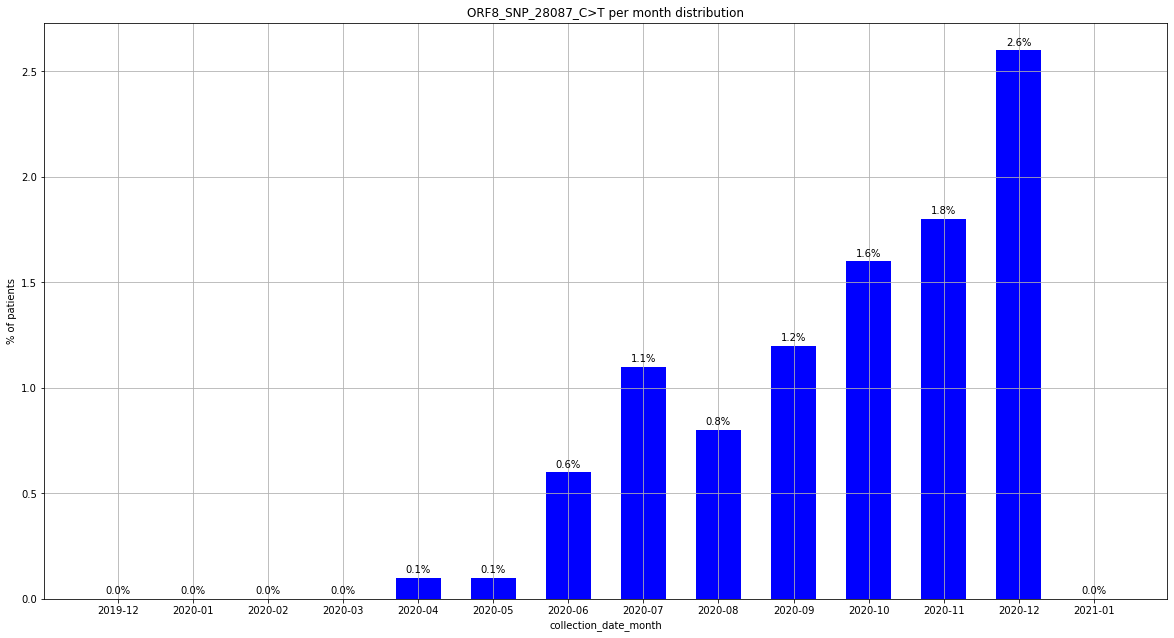

### output_19_44.png

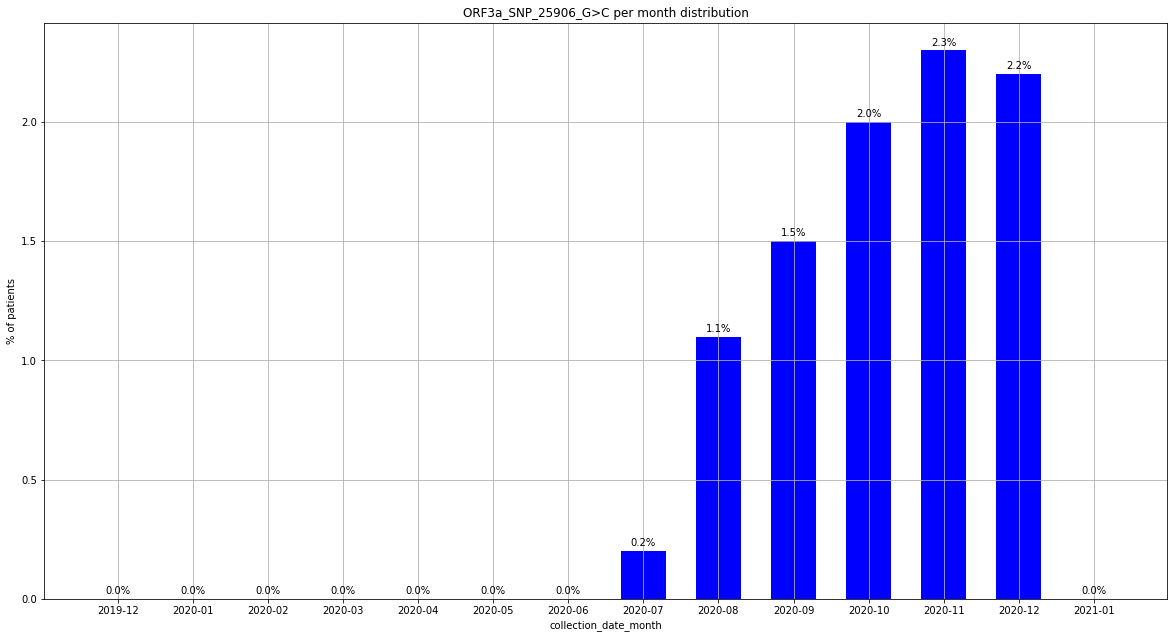

### output_19_49.png

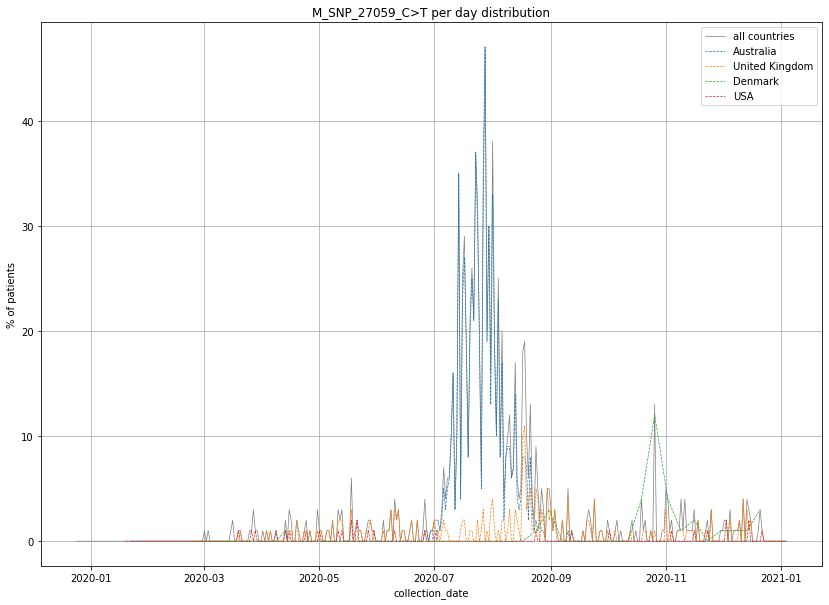

### output_19_49.png

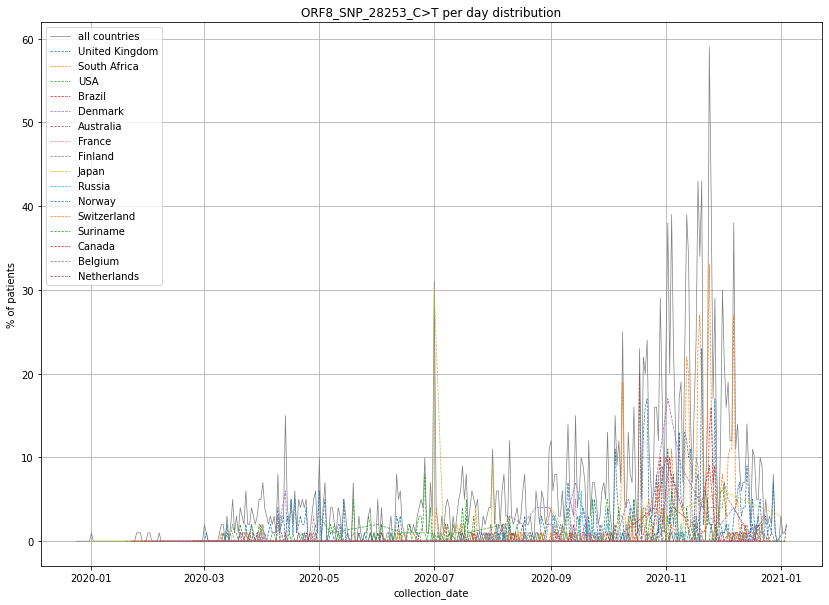

### output_19_50.png

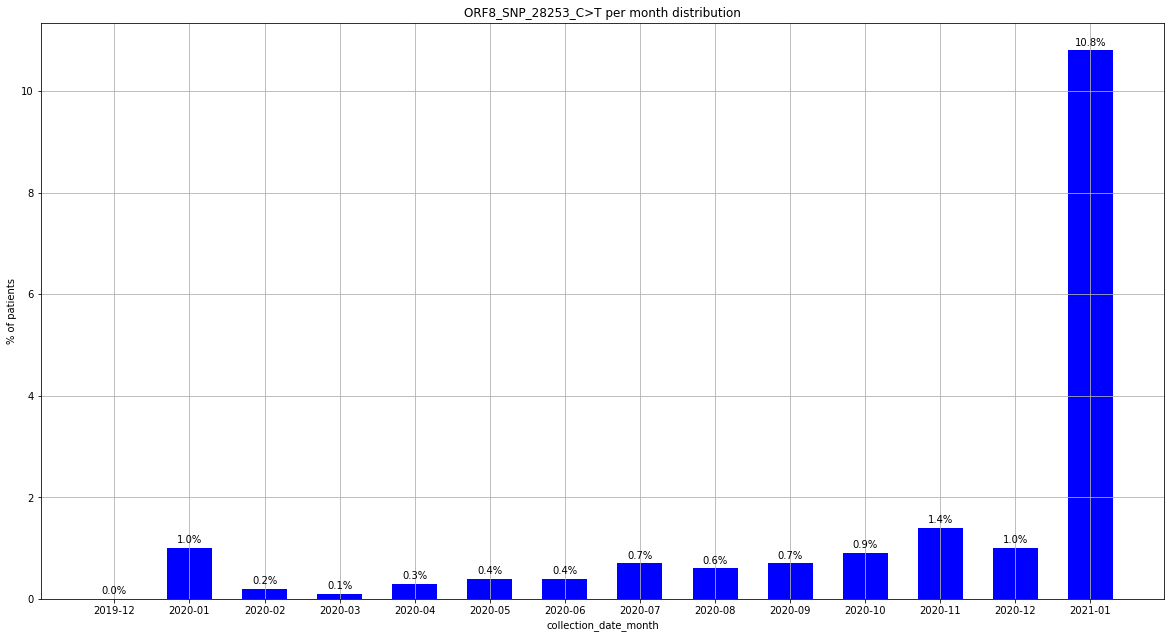

### output_19_50.png

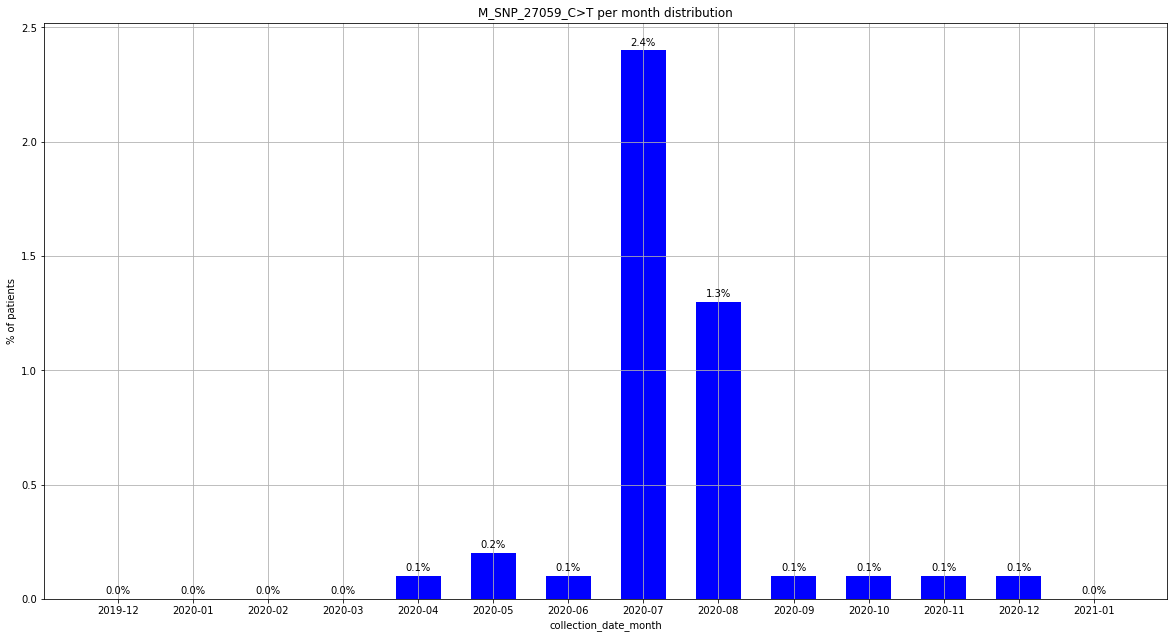

### output_19_55.png

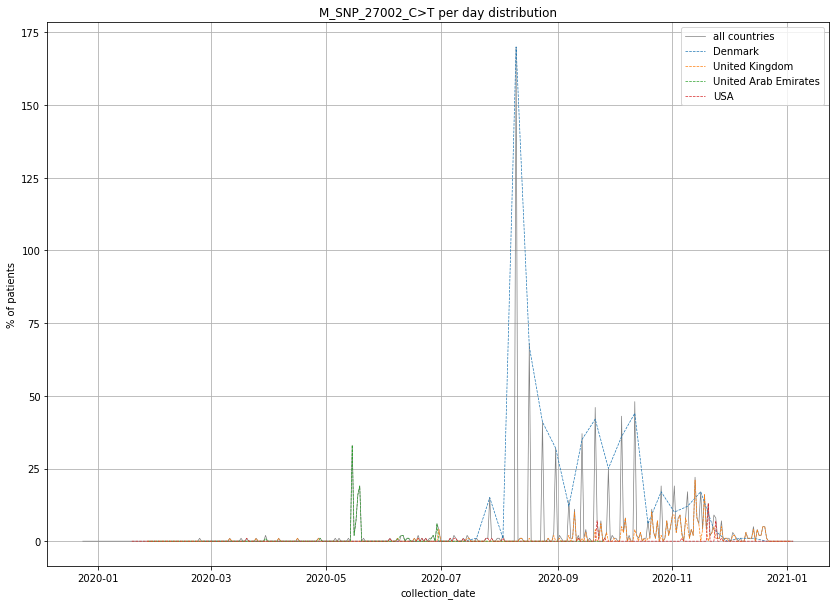

### output_19_55.png

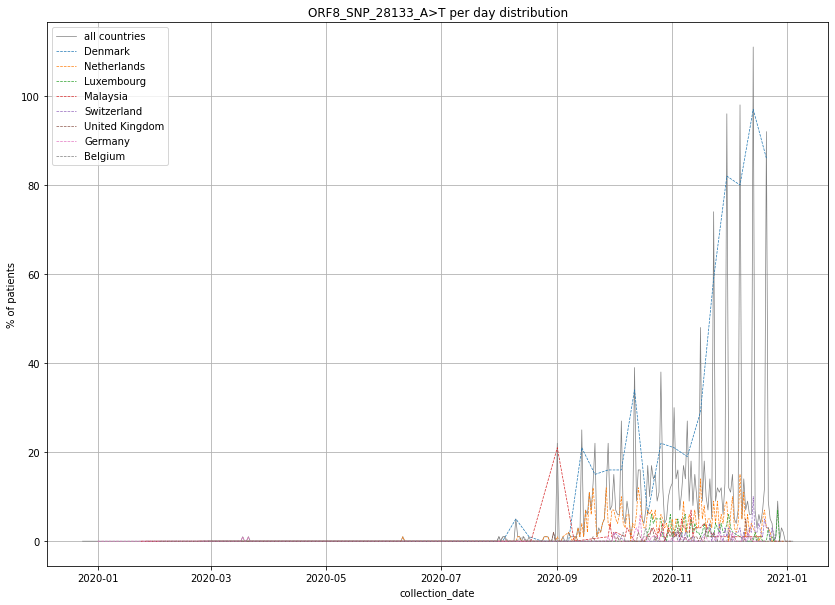

### output_19_56.png

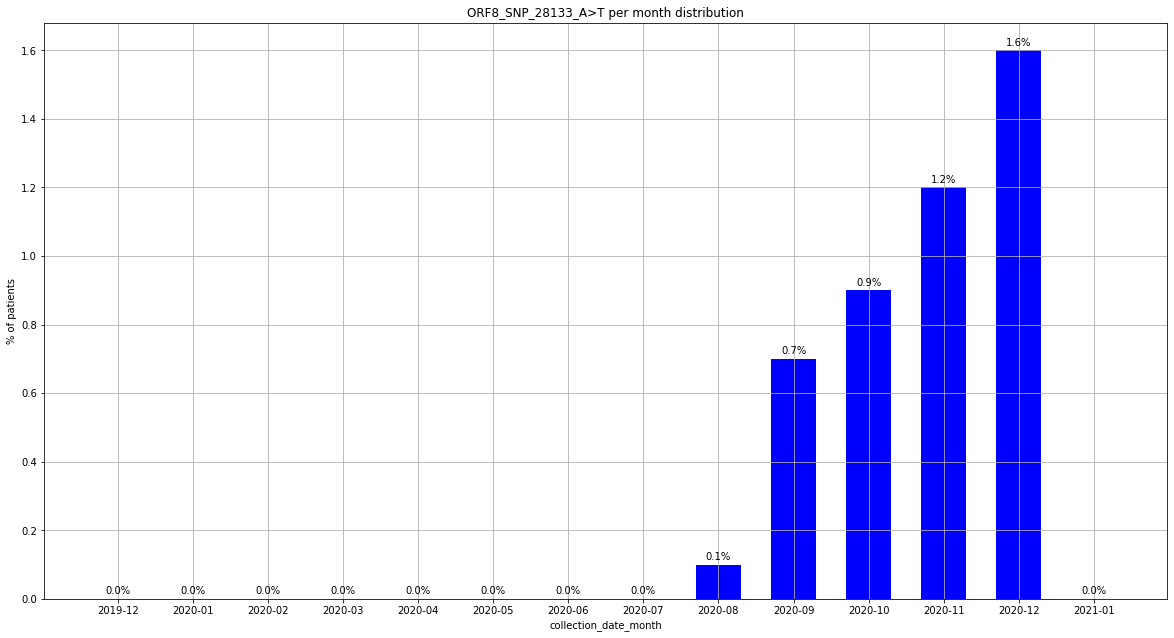

### output_19_56.png

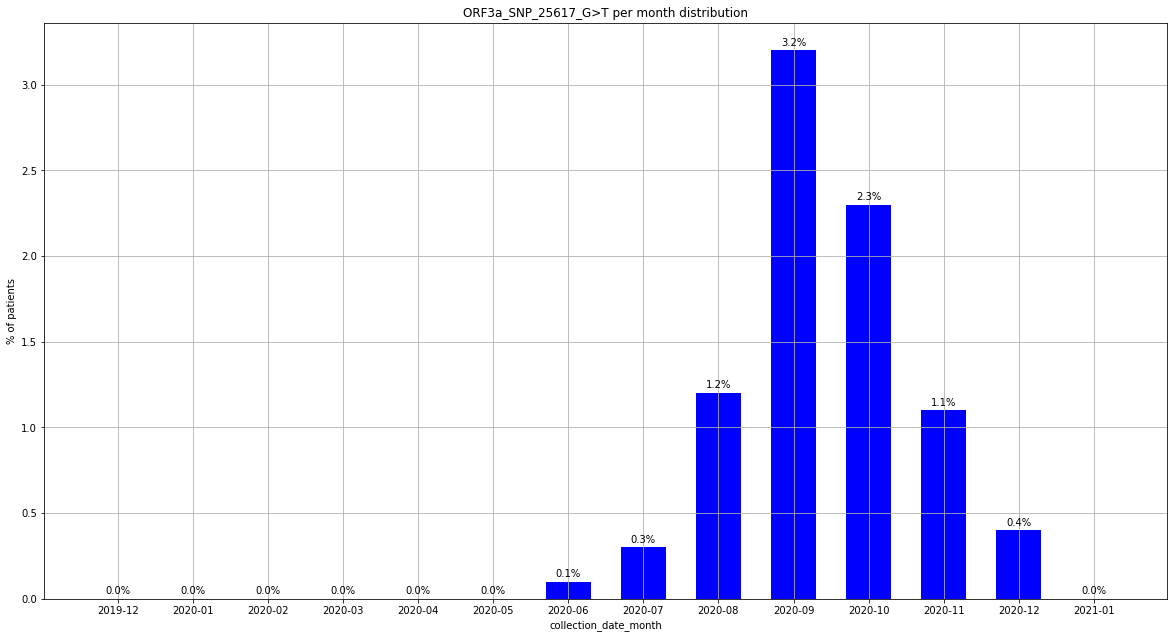

### output_19_61.png

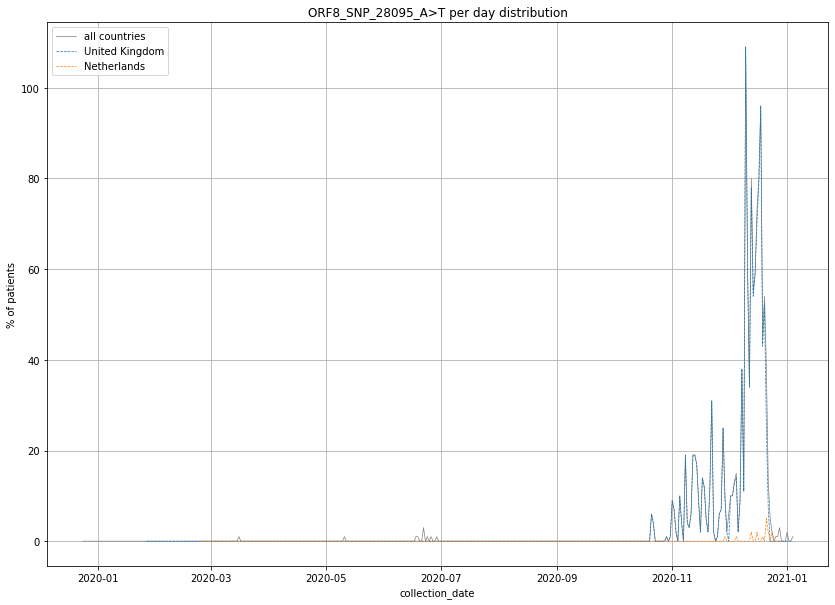

### output_19_61.png

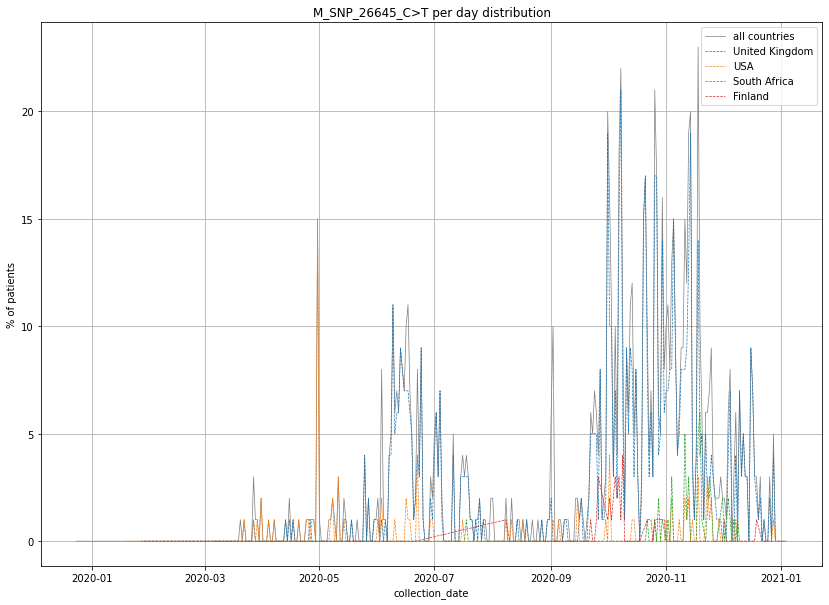

### output_19_61.png

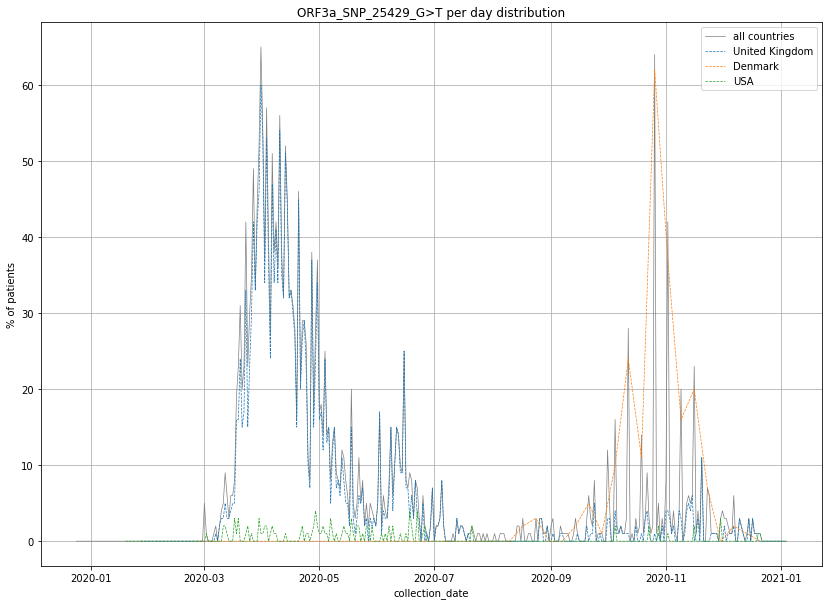

### output_19_62.png

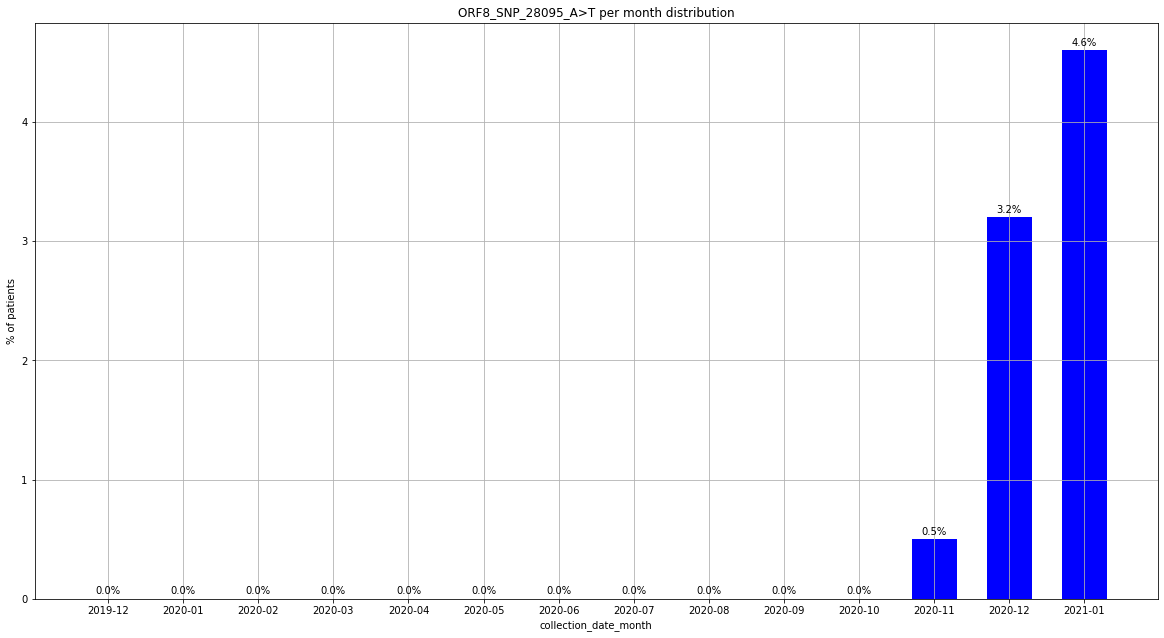

### output_19_62.png

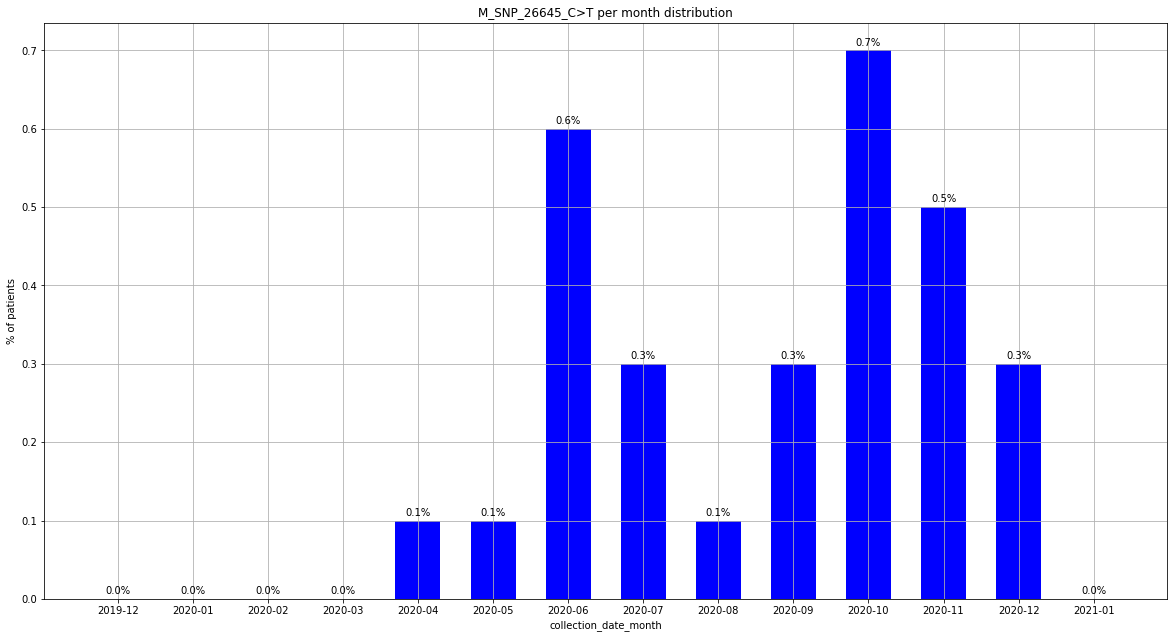

### output_19_62.png

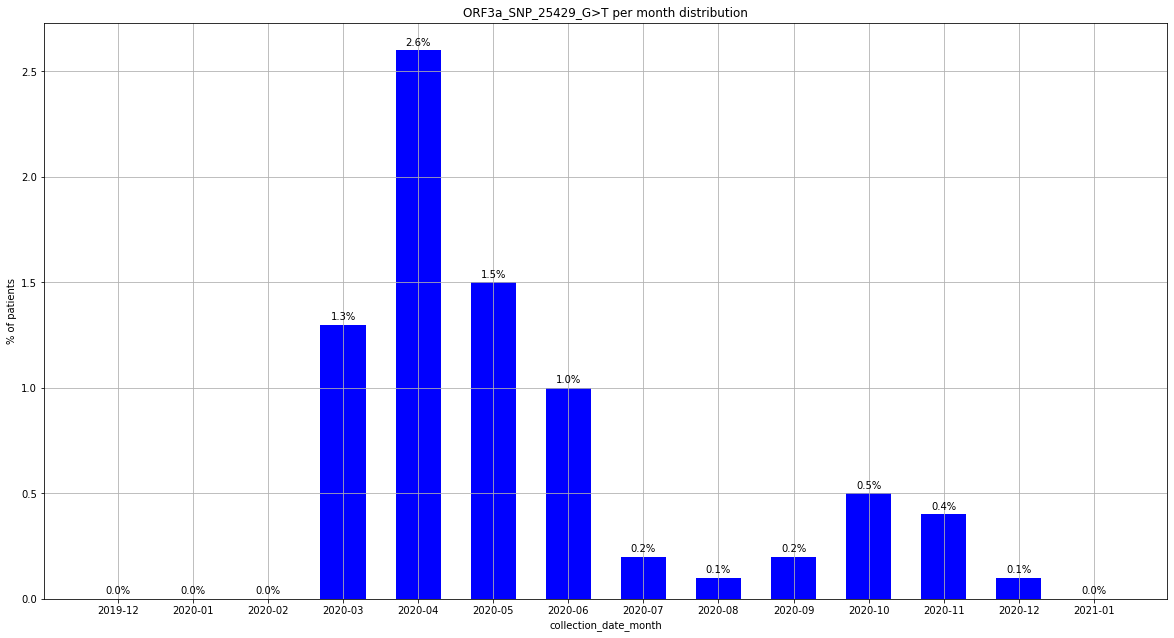

### output_19_67.png

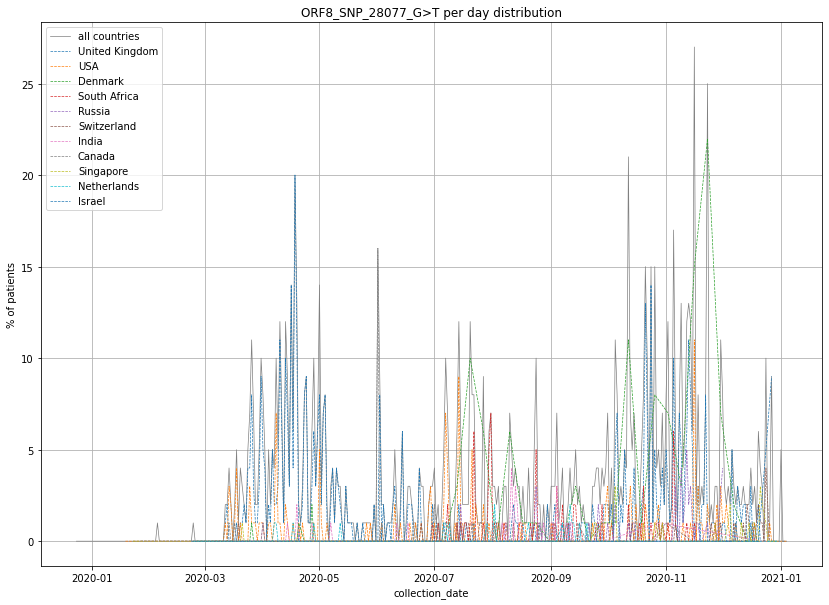

### output_19_68.png

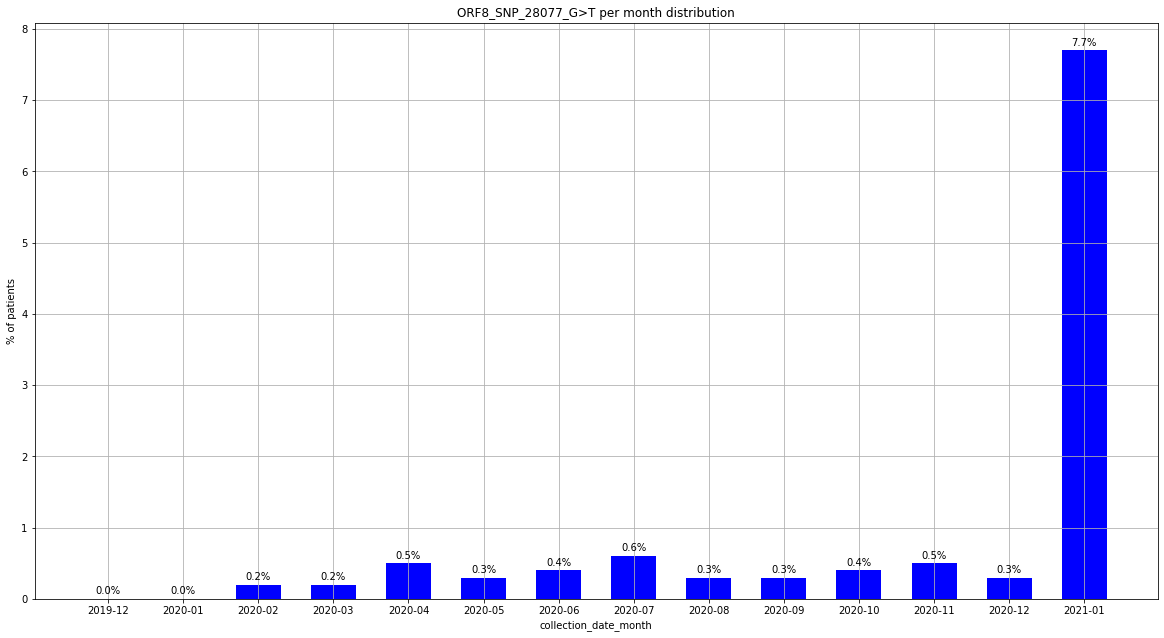

### output_19_68.png

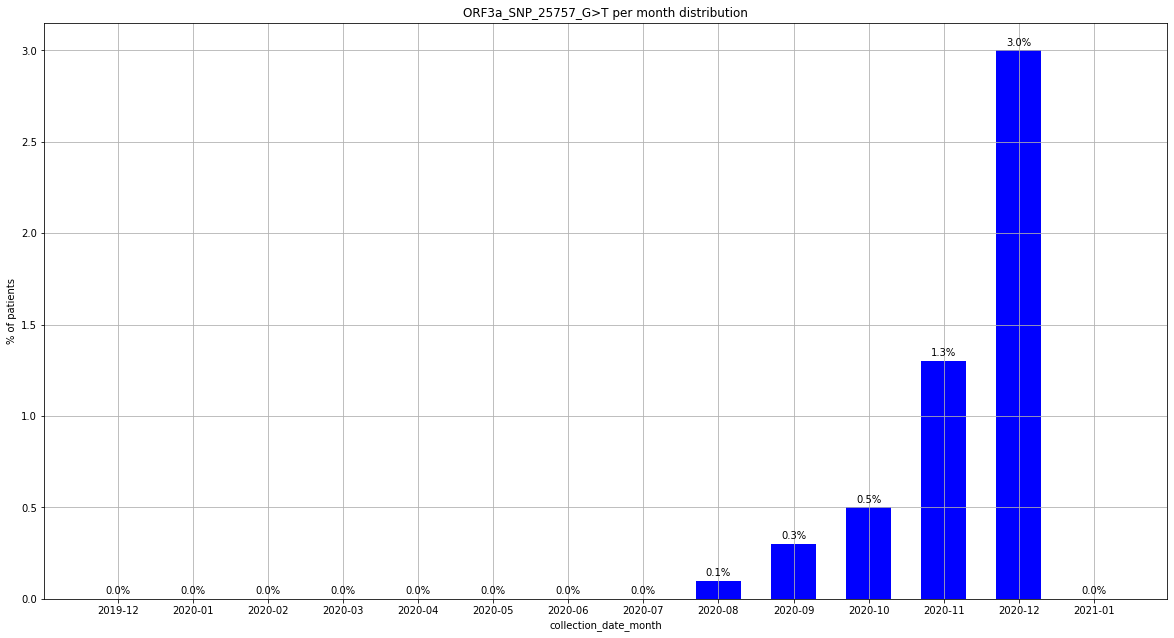

### output_19_73.png

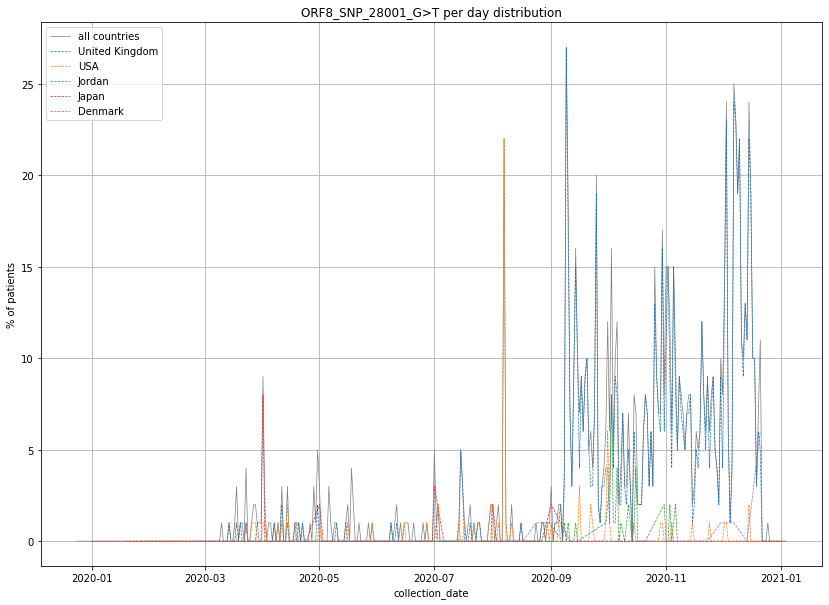

### output_19_74.png

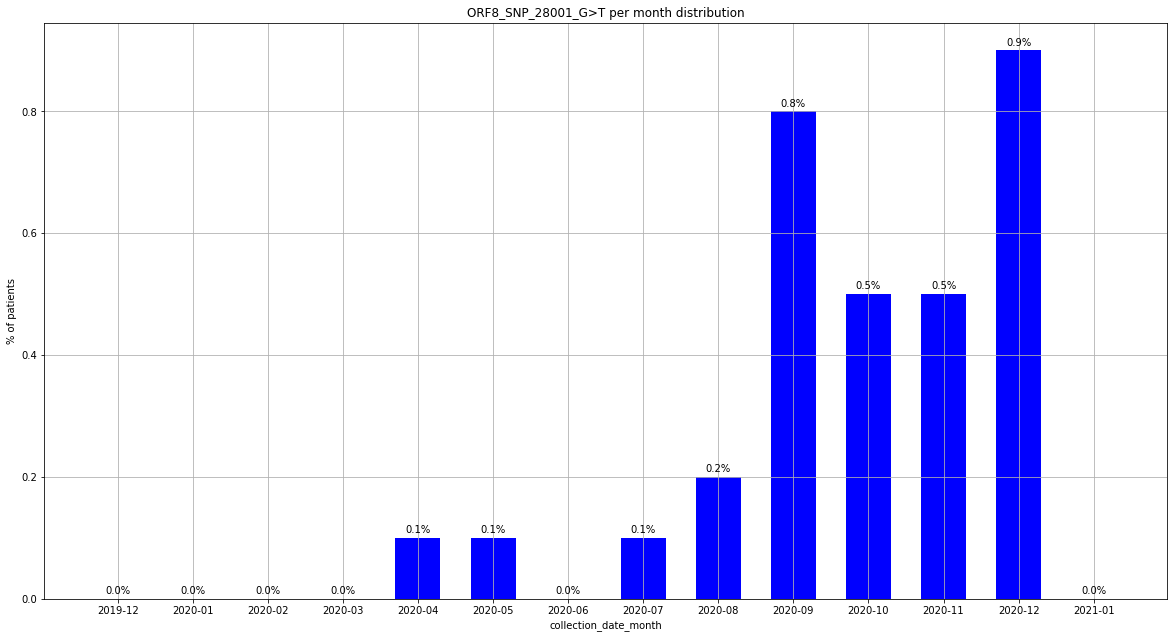

### output_19_74.png

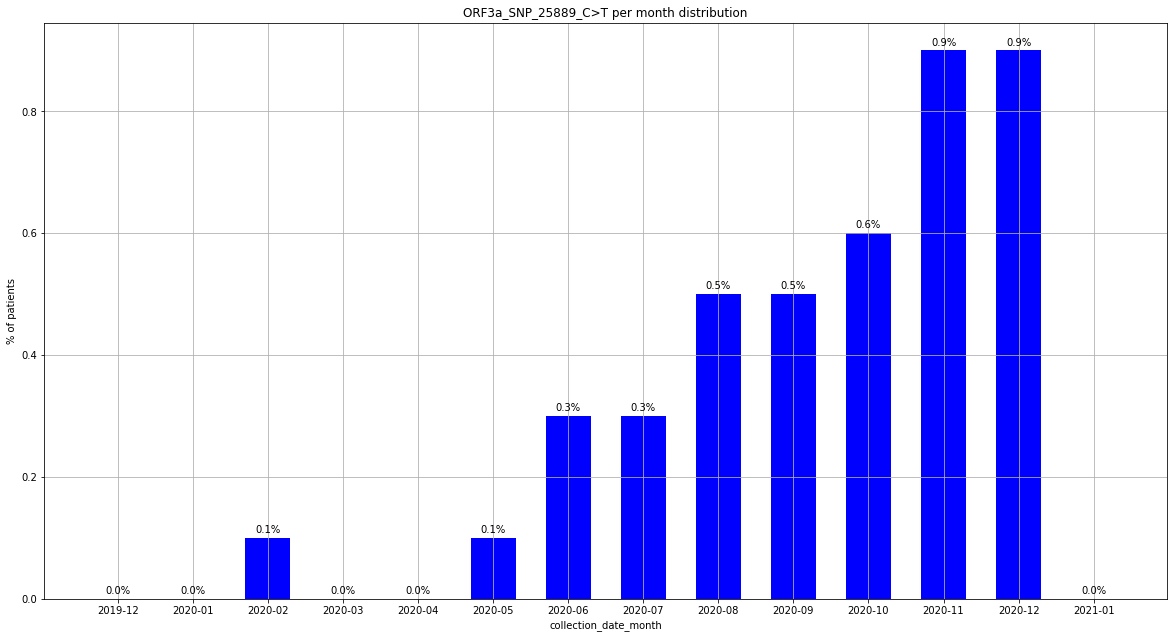

### output_19_79.png

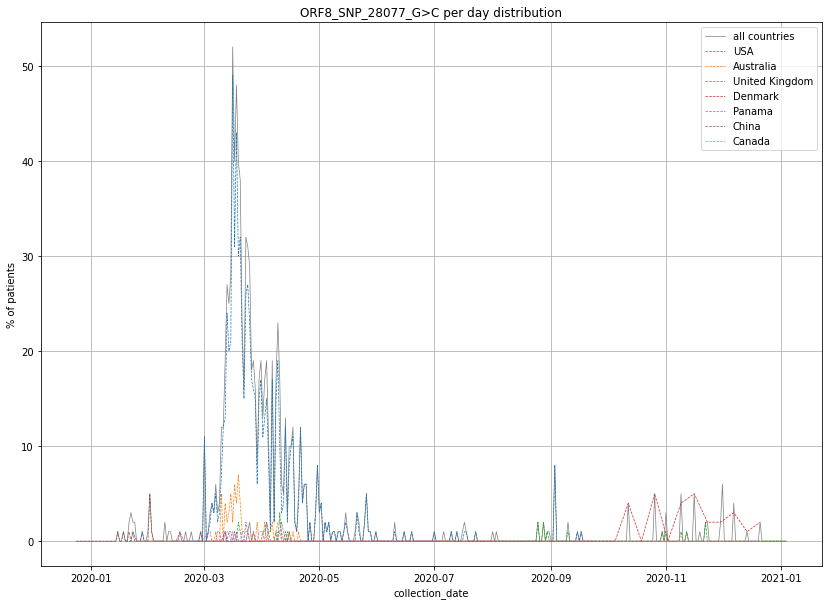

### output_19_79.png

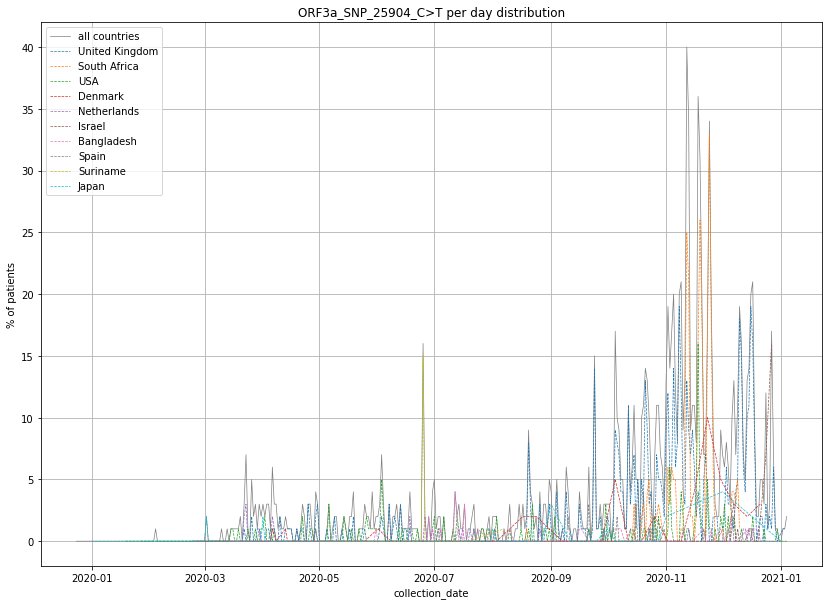

### output_19_80.png

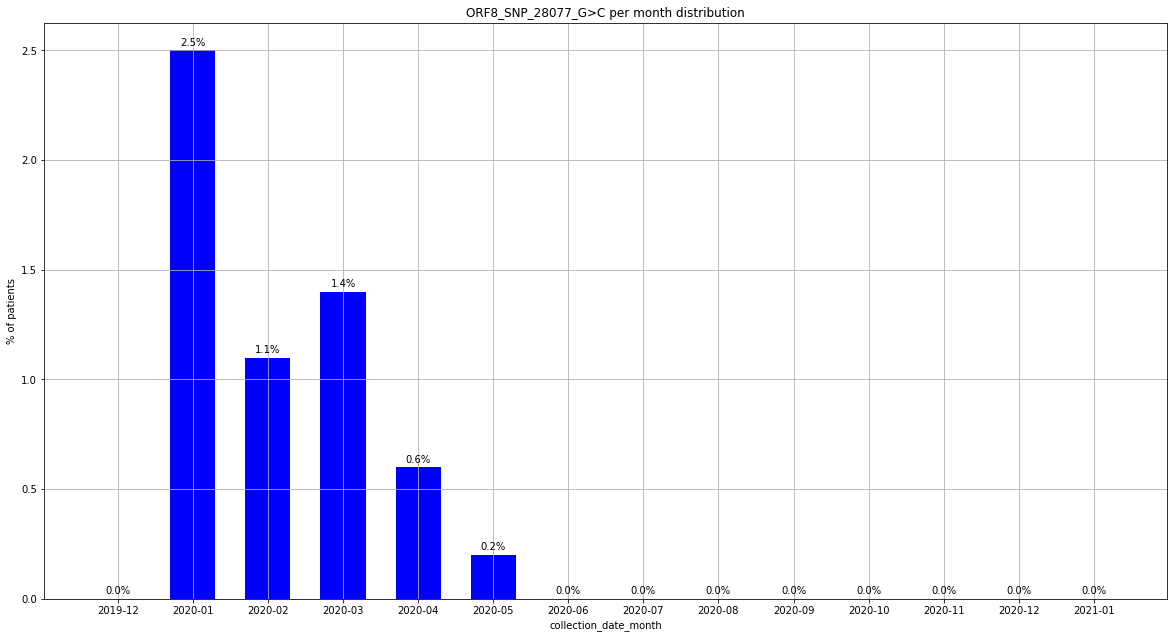

### output_19_86.png

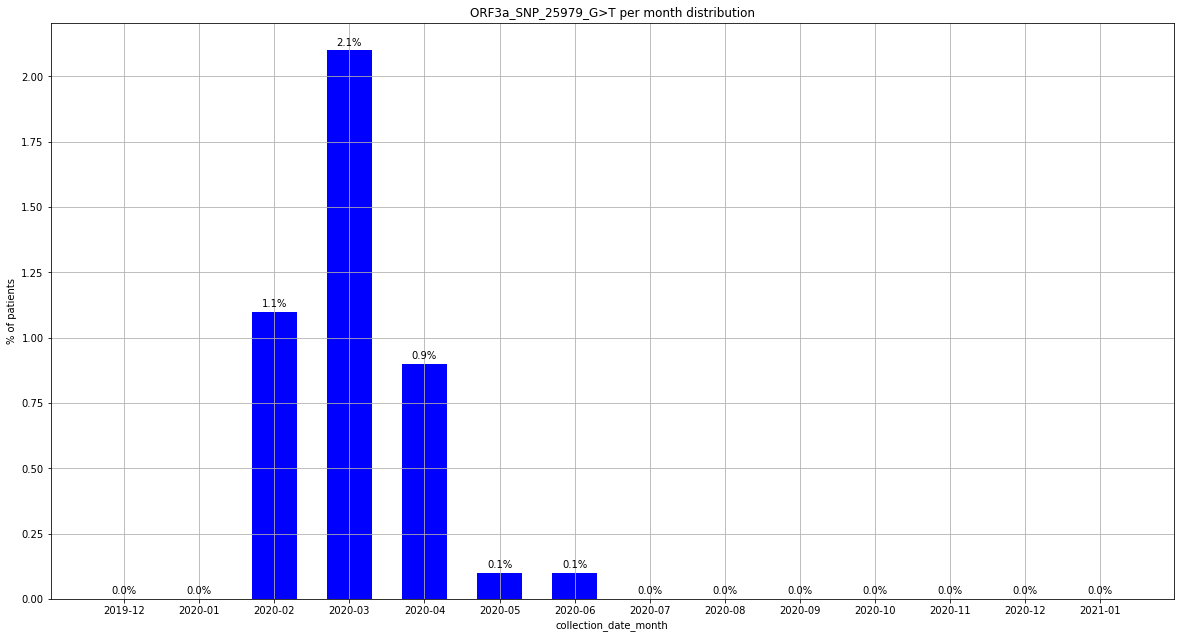

### output_19_91.png

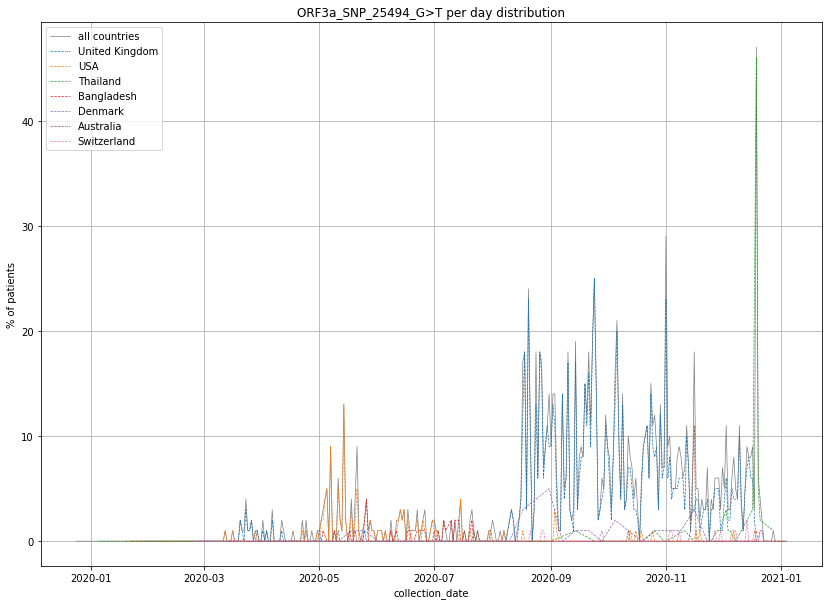

### output_19_97.png

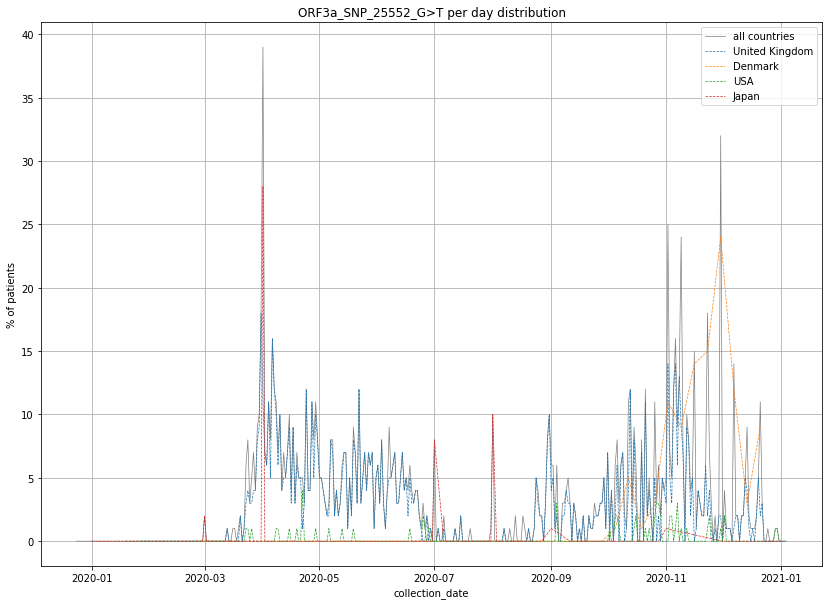

### output_19_98.png

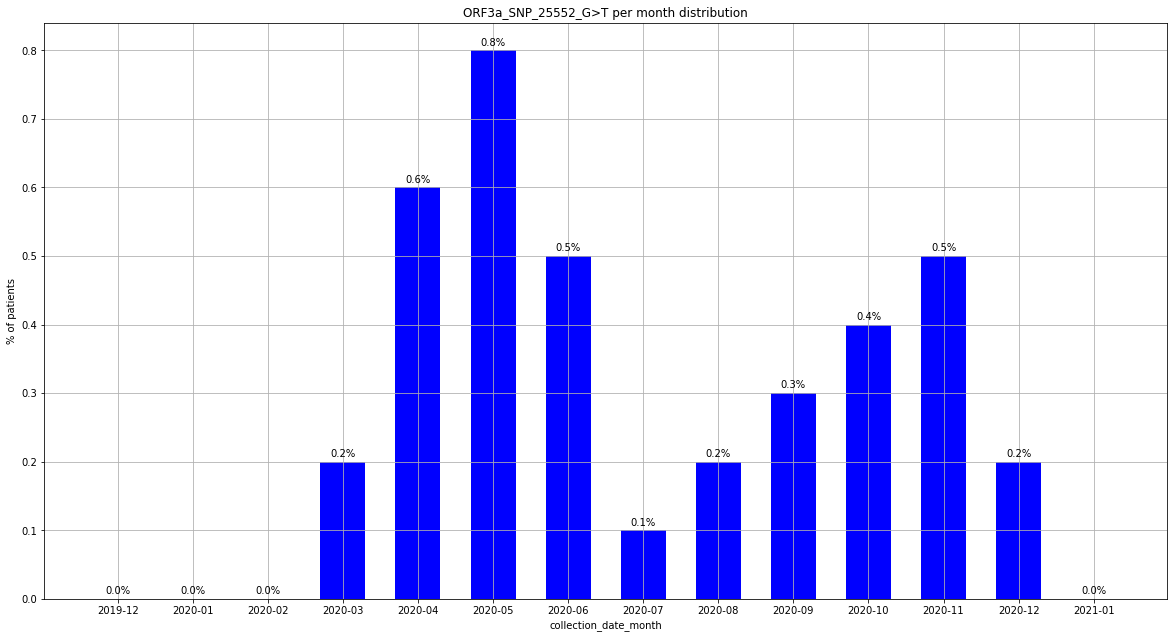

### output_19_103.png

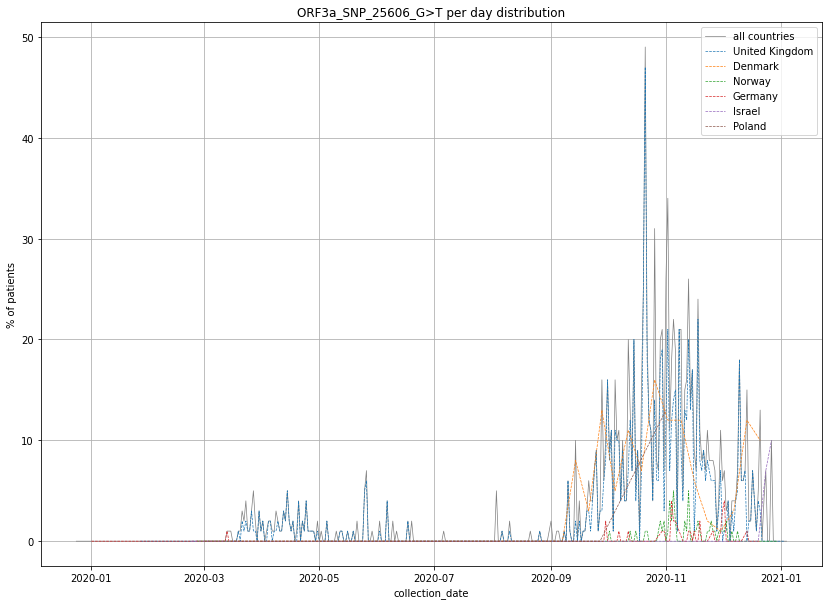

### output_19_109.png

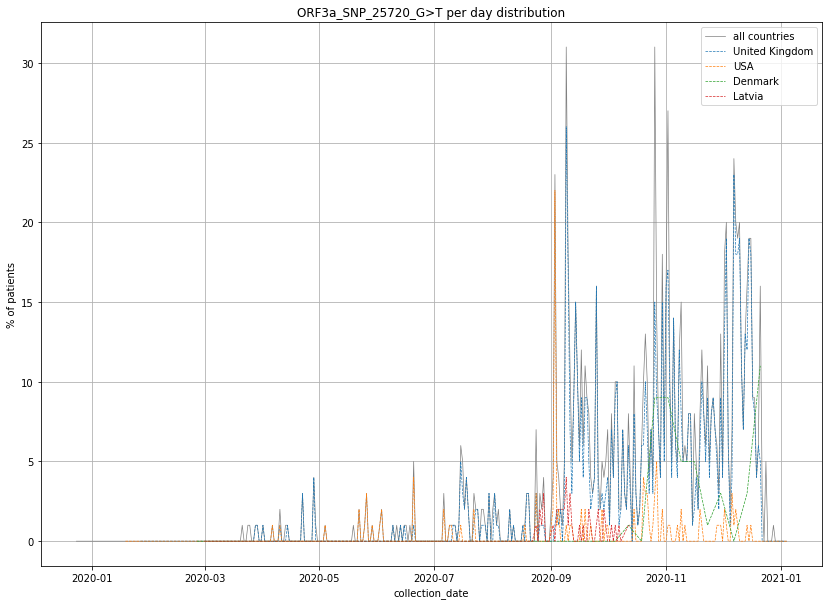

### output_19_110.png

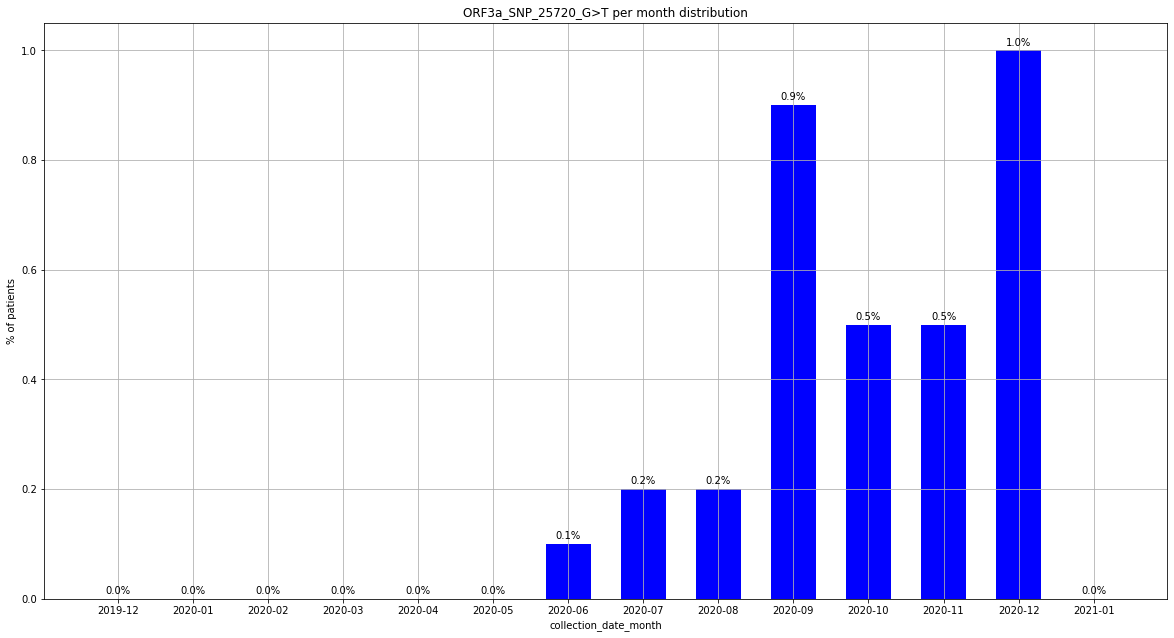

### output_19_121.png

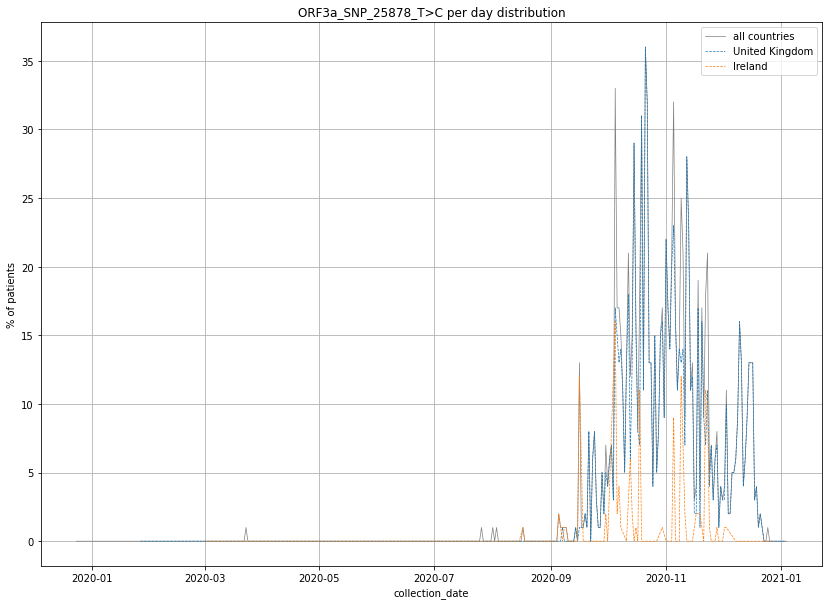

### output_19_122.png

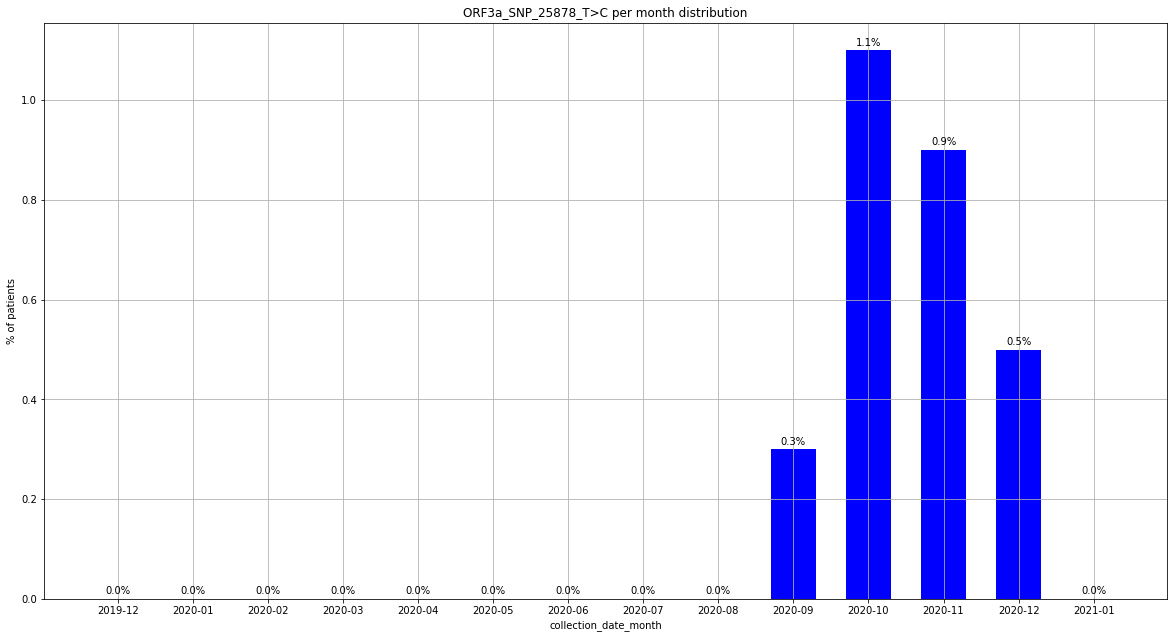

### output_19_128.png

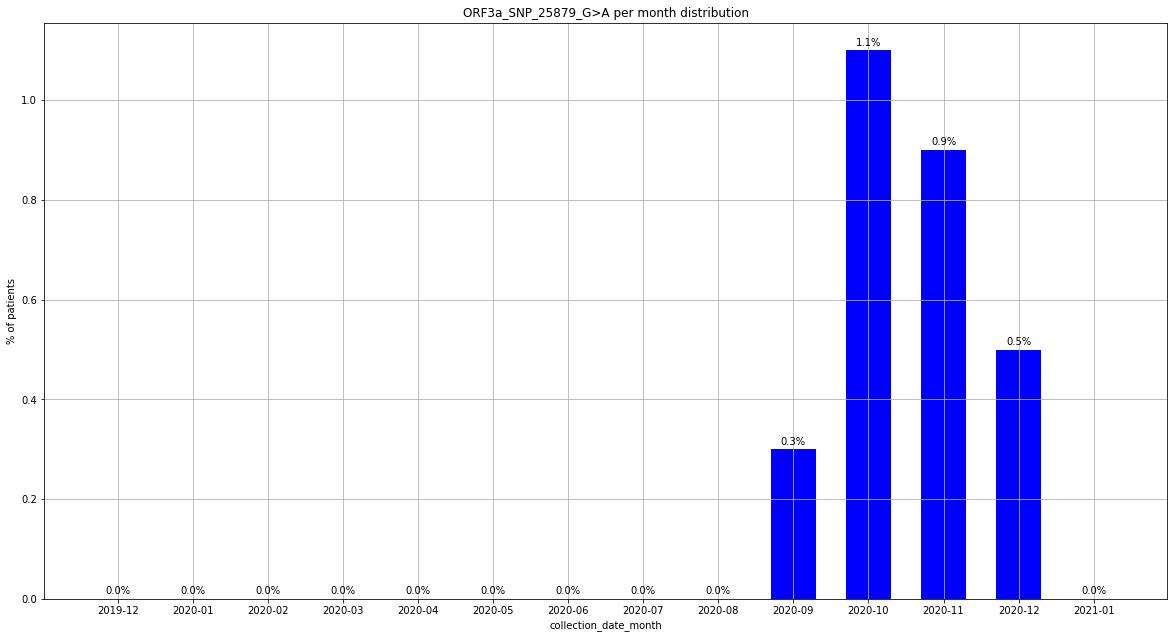

### output_19_133.png

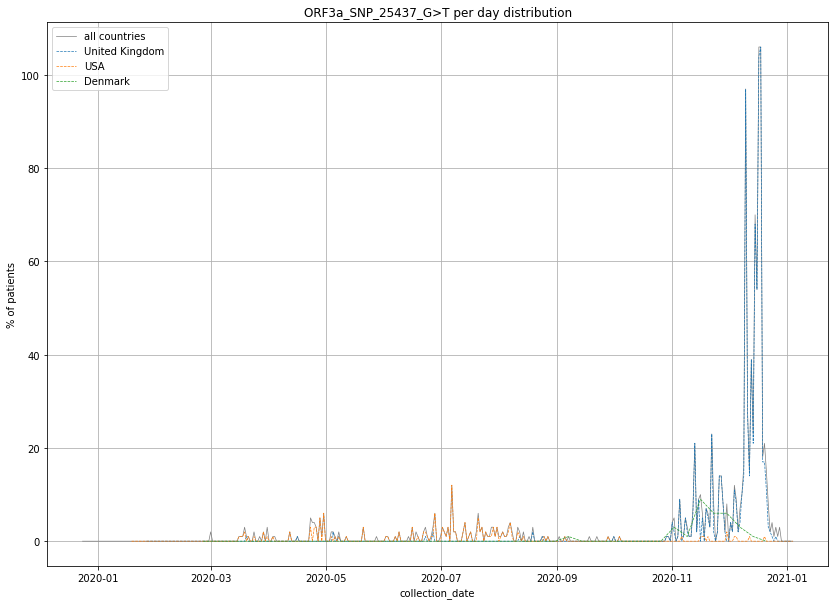

### output_19_134.png

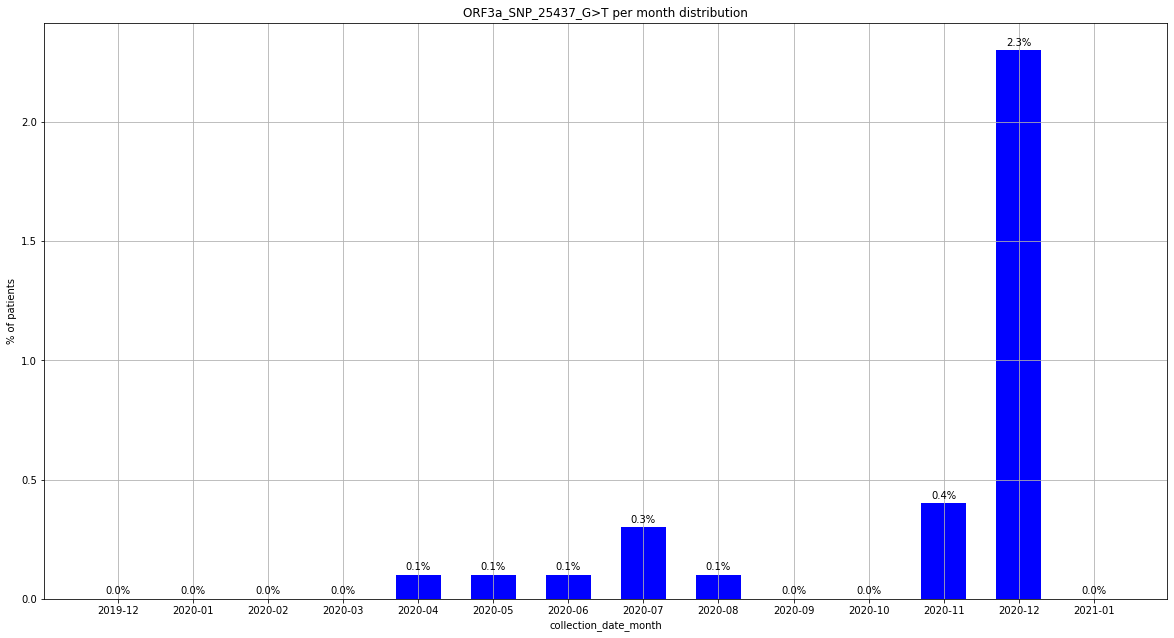

### output_19_139.png

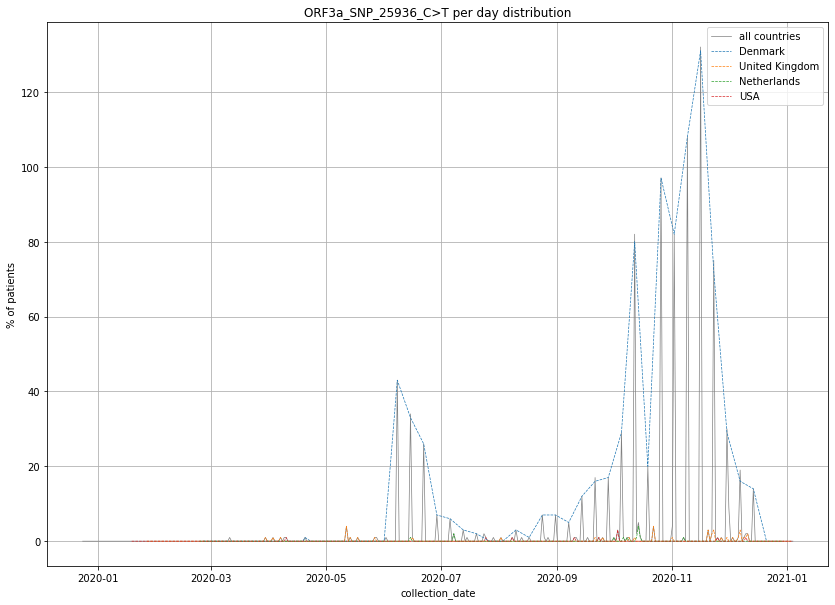

### output_19_140.png

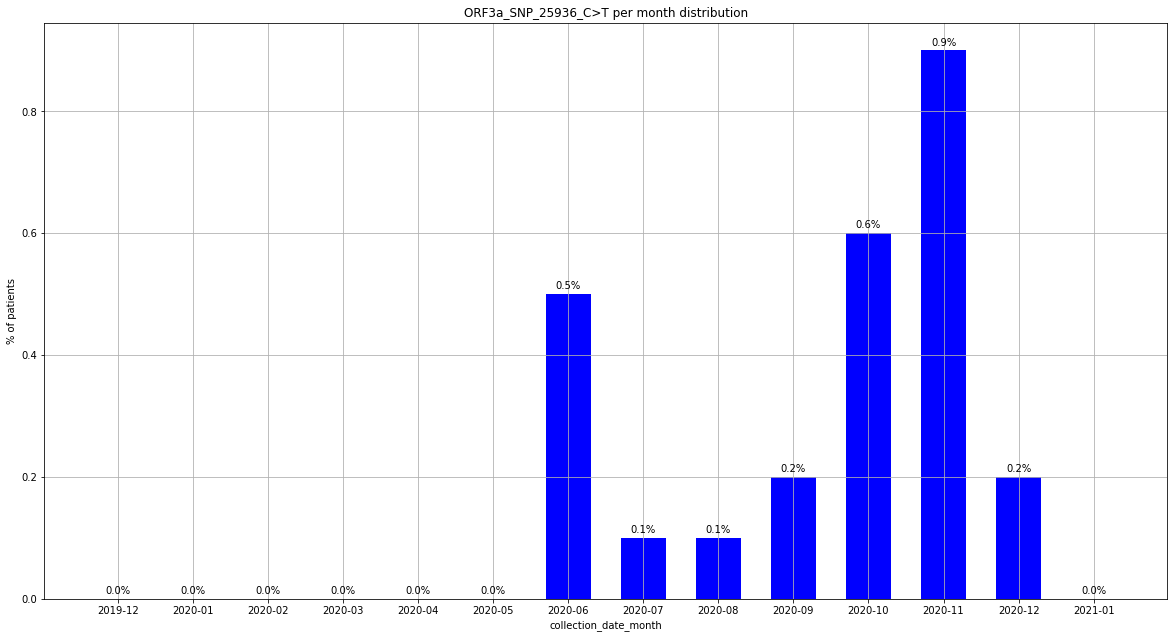

### output_19_146.png

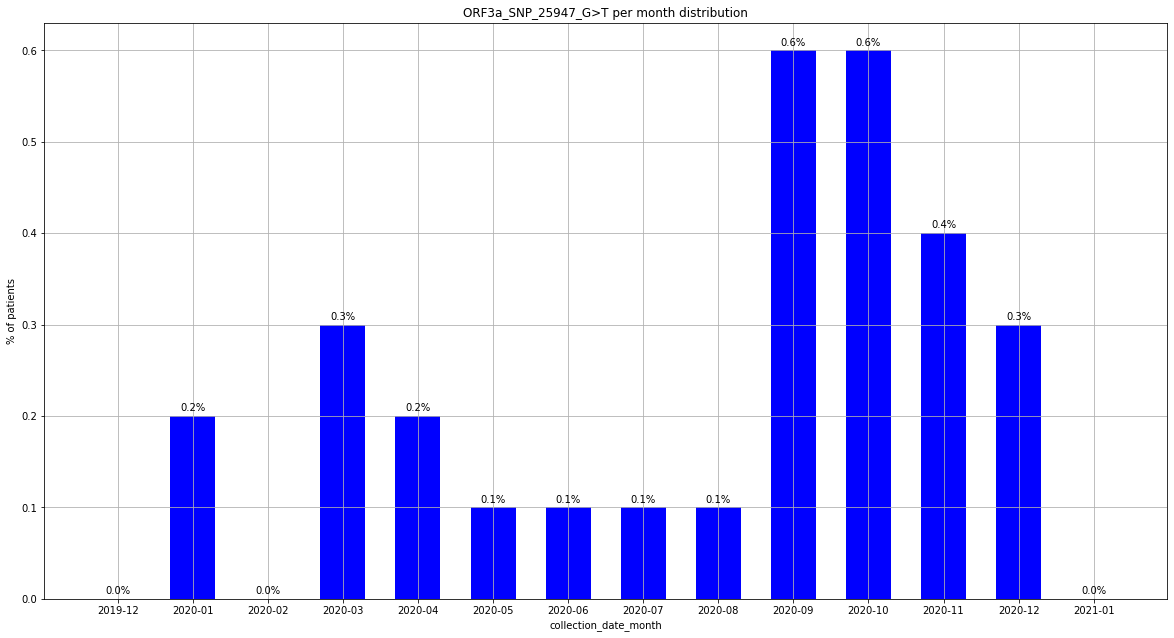

### output_20_1.png

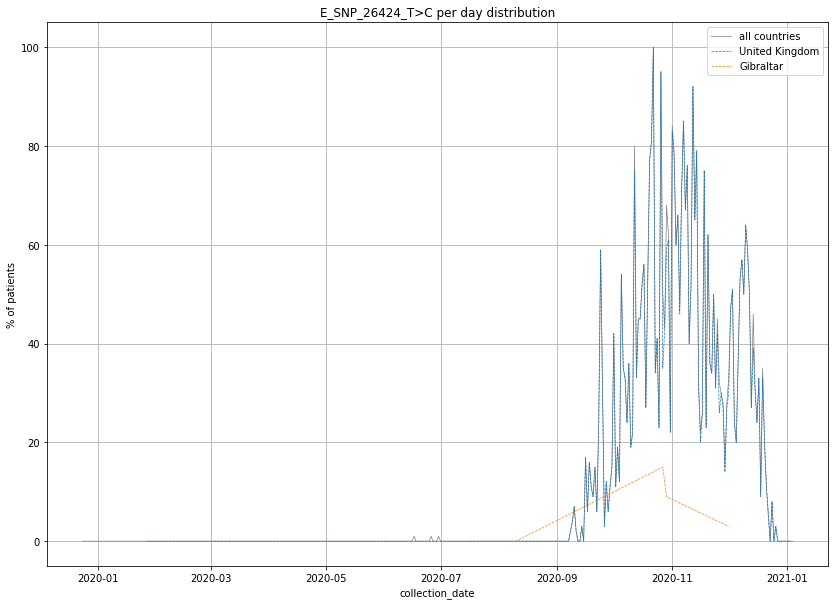

### output_20_2.png

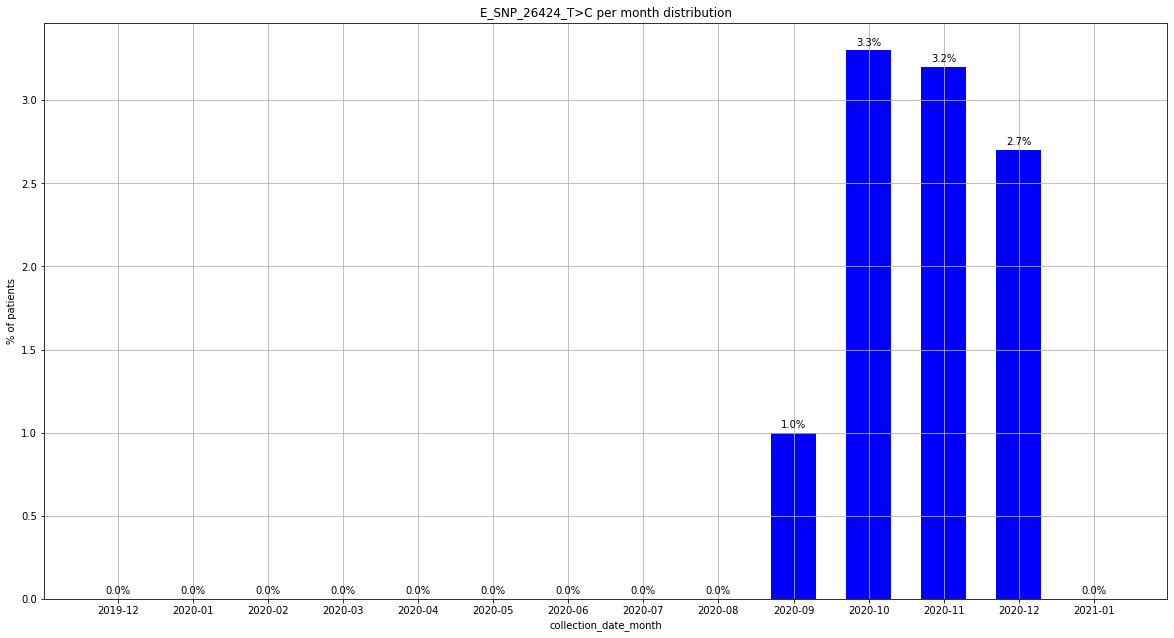

### output_20_7.png

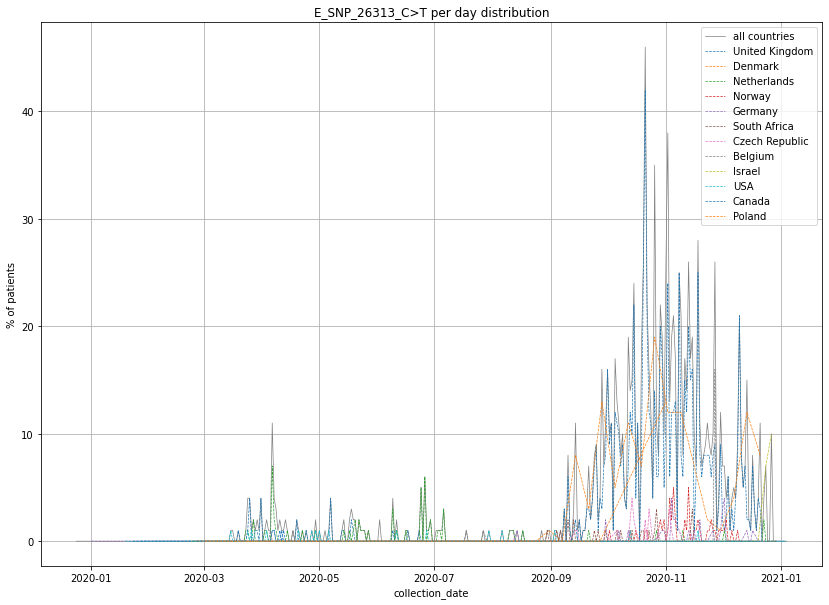

### output_20_8.png

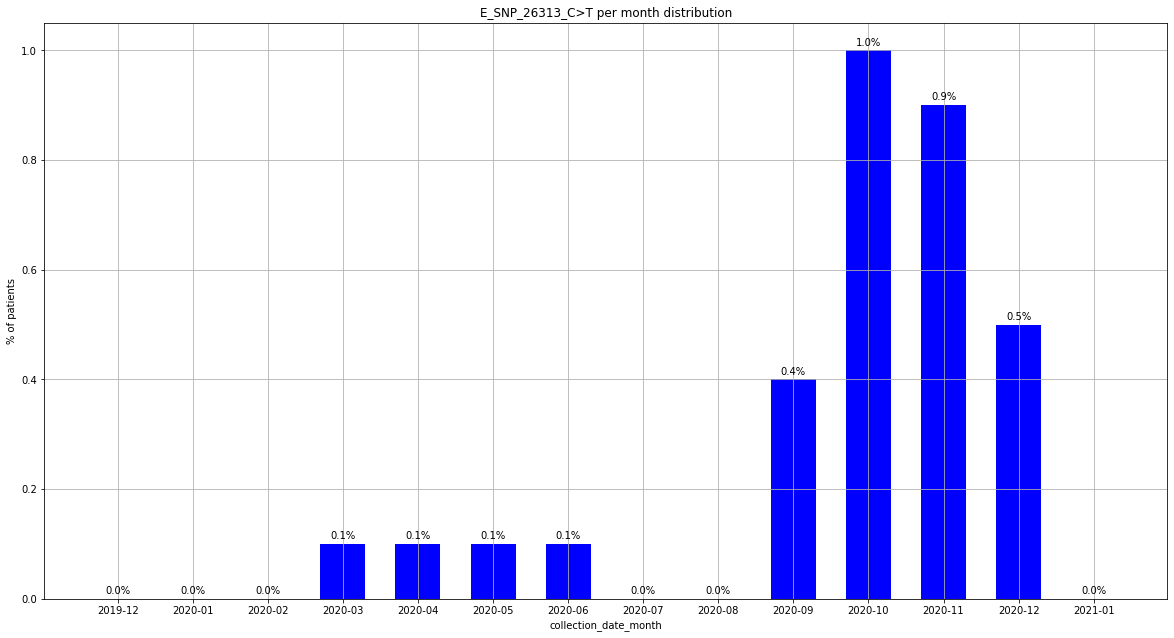

### output_20_50.png

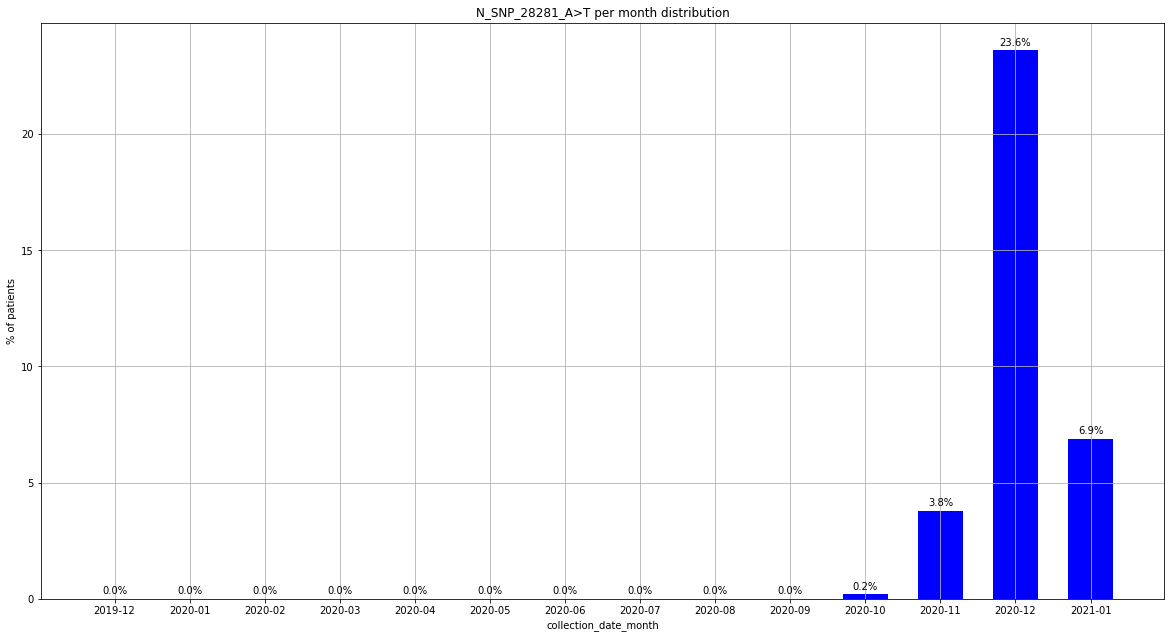

### output_20_67.png

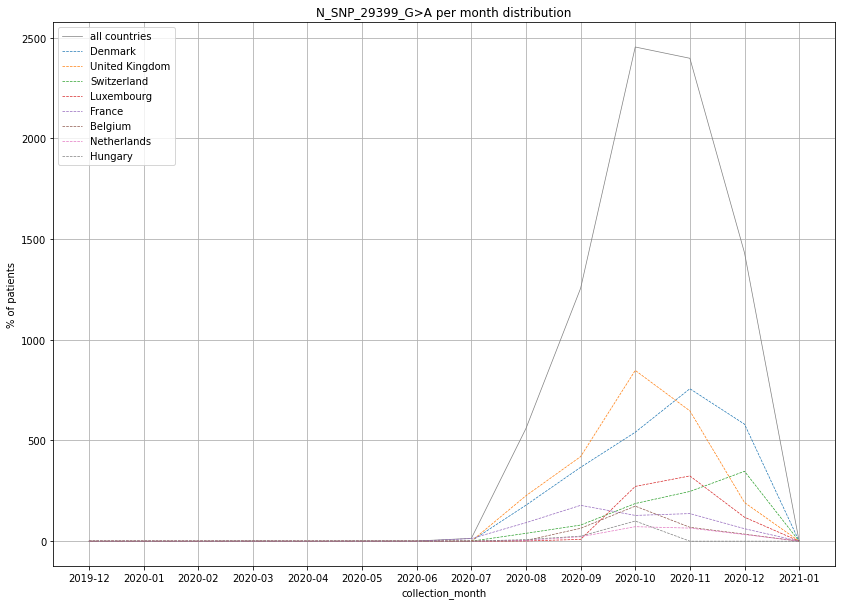

### output_20_68.png

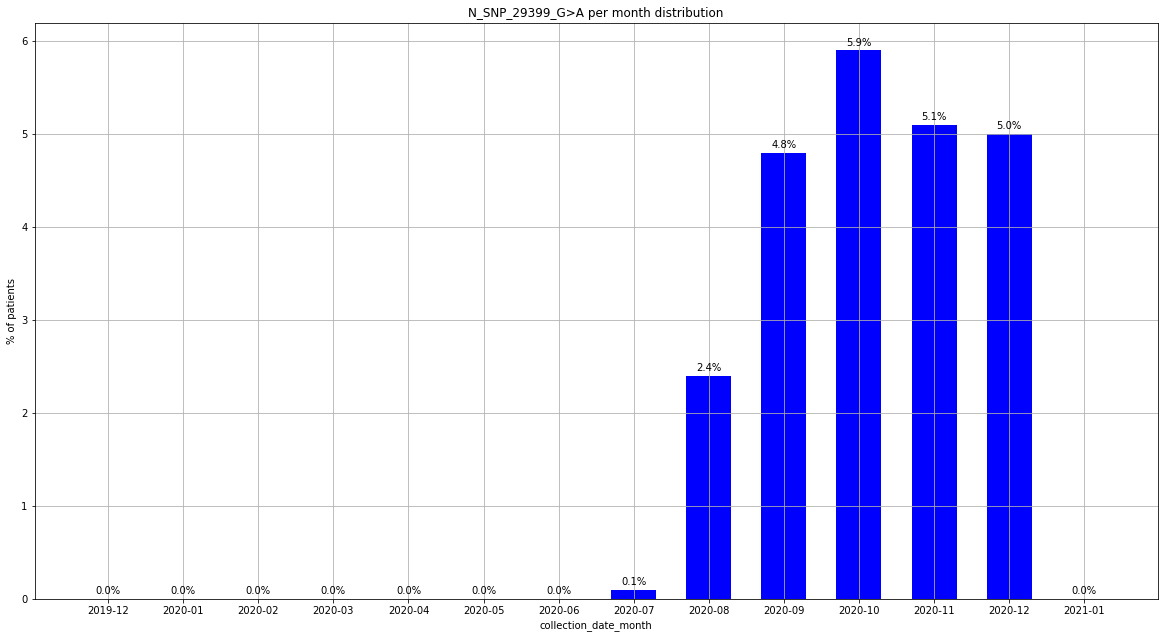

### output_20_134.png

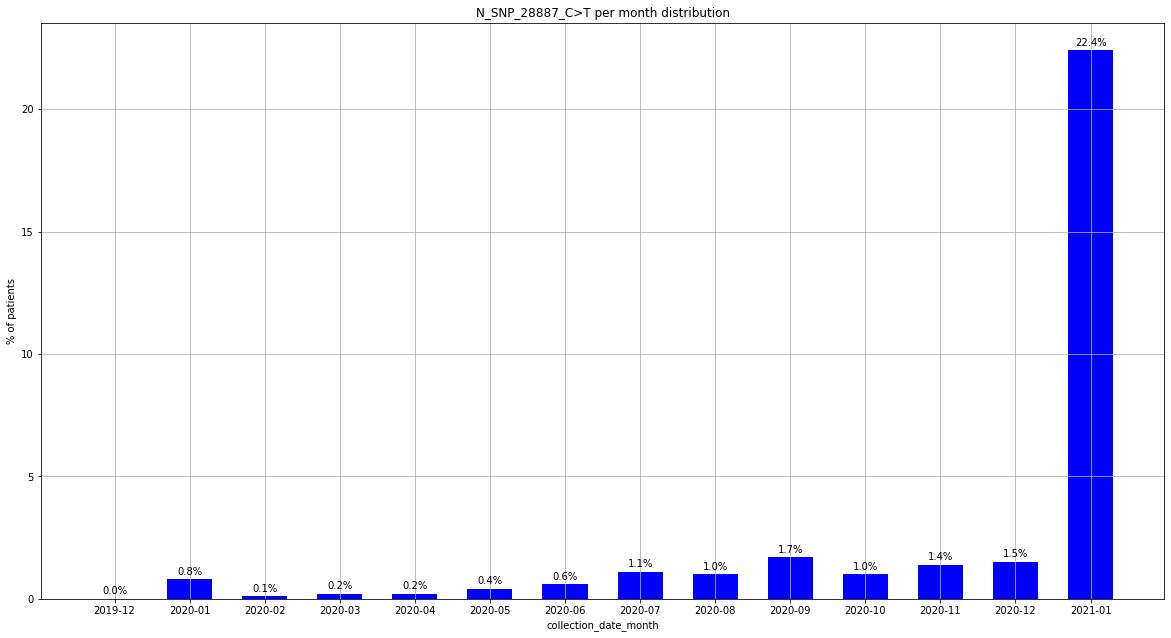

### output_20_139.png

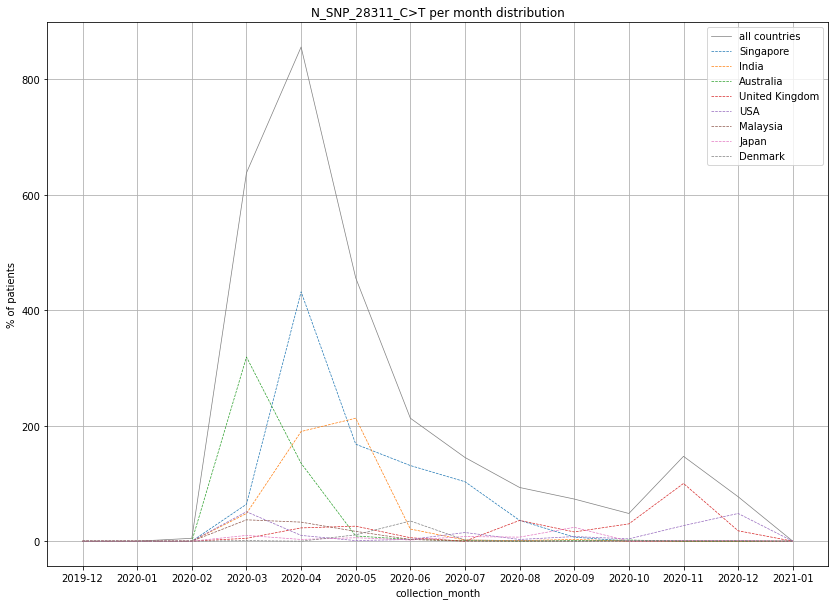

### output_20_157.png

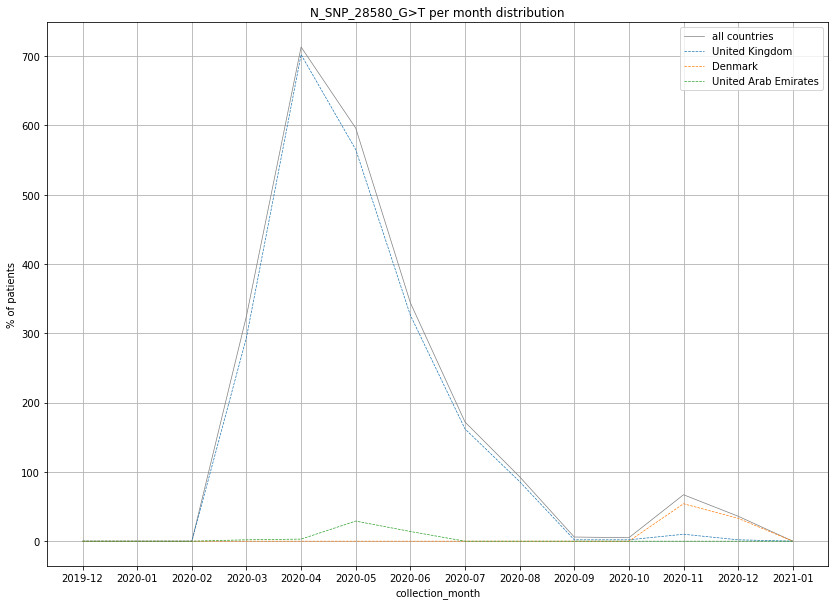

### output_20_169.png

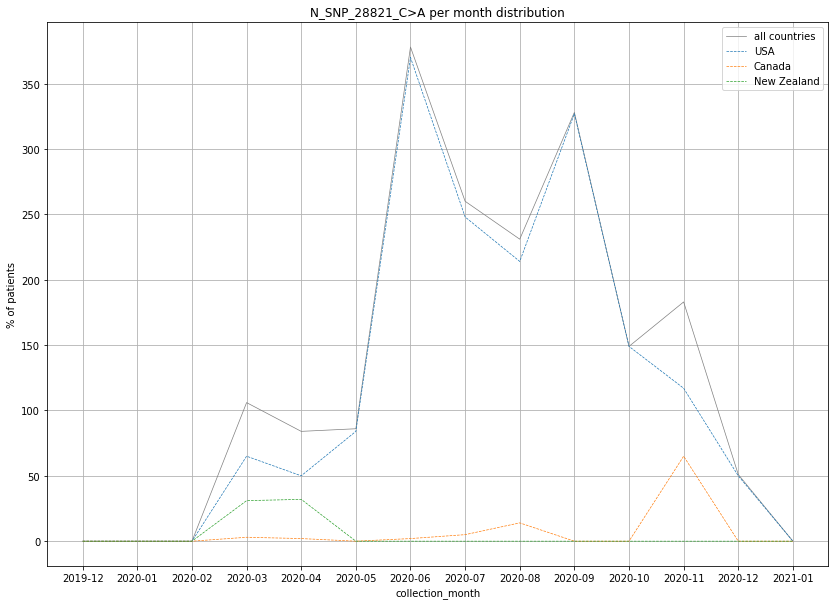

### output_20_170.png

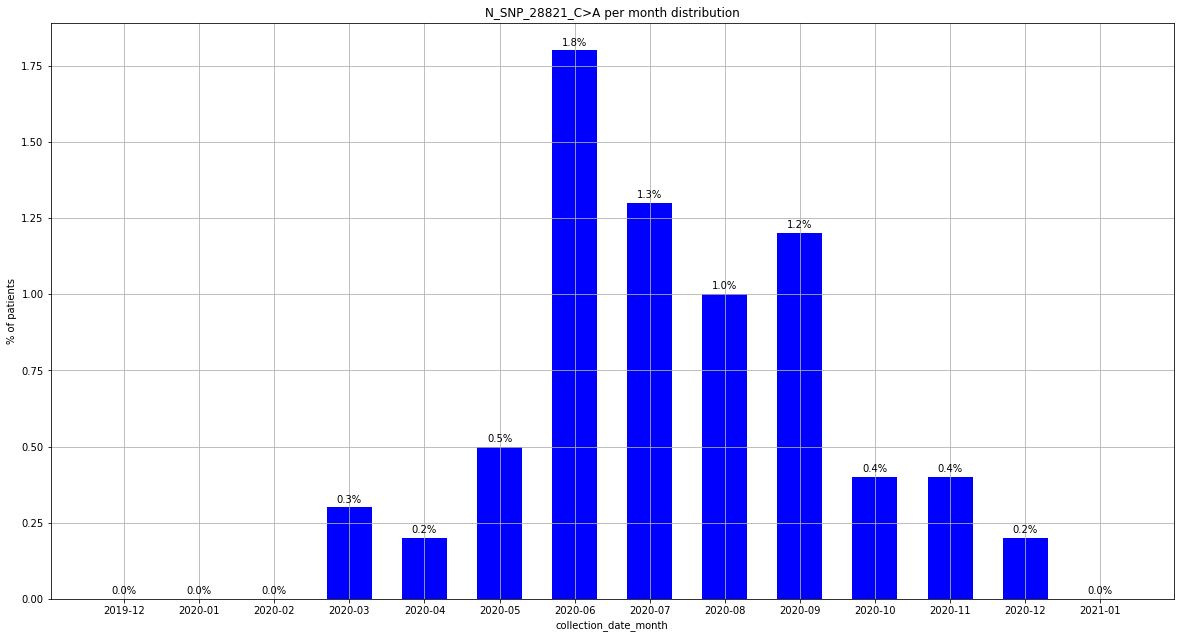

### output_20_205.png

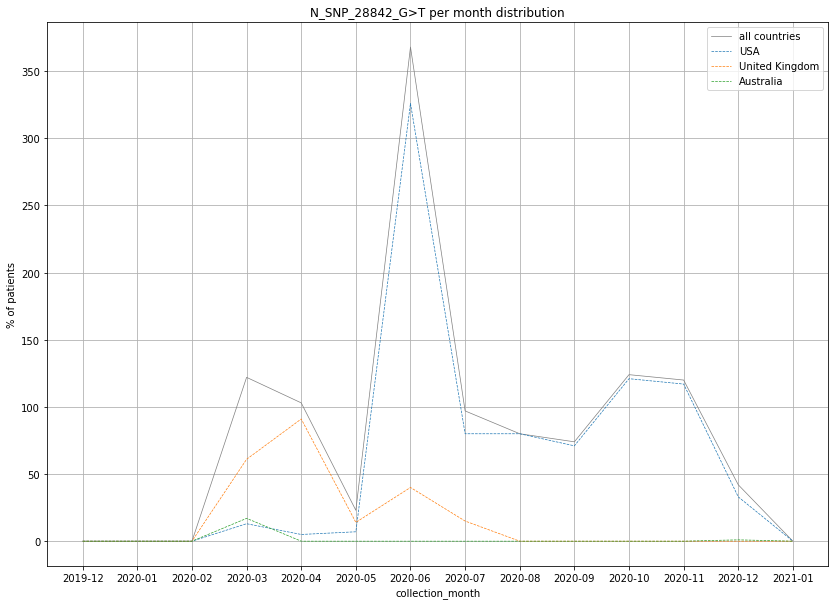
